# Language Modeling Materializes a World Model of Protein Biology

**DOI:** 10.64898/2026.06.03.729735

**Authors:** Salvatore Candido, Thomas Hayes, Alexander Derry, Roshan Rao, Zeming Lin, Robert Verkuil, Bryan Z. Wu, Jin Sub Lee, Elise S. Bruguera, Jehan A. Keval, Mykhailo Kopylov, John E. Pak, Wesley Wu, Neil Thomas, Samson Mataraso, Alvin Hsu, Ashton C. Trotman-Grant, Kilian Fatras, Allan dos Santos Costa, Rohil Badkundri, Halil Akın, Deniz Oktay, Jonathan Deaton, Elizabeth Montabana, Hrishita Sitwala, Yue Yu, Marius Wiggert, Dylan Alexander Carlin, Anthony W. Goering, Tomasz Blazejewski, McCullen Sandora, Michael Hla, Tina Z. Jia, Leon H. Kloker, Nicholas J. Sofroniew, Masatoshi Uehara, Jassi Pannu, Sharrol Bachas, Daniel S. Liu, Tom Sercu, Alexander Rives

## Abstract

Proteins are fundamental to life. The full extent of their biology is beyond our ability to characterize with experimental approaches in the physical laboratory. Accurate digital representations could accelerate the discovery of protein biology through virtual experiments. We propose language modeling to learn unified and general representations that can be scaled to all of protein biology. Building on these representations, we develop a structure prediction model that exceeds the performance of established methods for biomolecular complex prediction across benchmarks, including for the interactions of antibodies with their targets. A simple search procedure yields high experimental success rates for the discovery of proteins with nanomolar binding affinities for both miniproteins and single-chain antibodies, a modality critical for therapeutic design. Study of the concepts in the language model’s representation space reveals a systematic organization aligned with the reductionist understanding of proteins developed through empirical science. Leveraging this organization, we generate a comprehensive map of protein biology encompassing over 6.8 billion sequences and 1.1 billion predicted structures, identifying connections across known and unknown biology. As a whole, this shows language modeling as a powerful substrate for representing the biology of proteins, operating across scales from the prediction and design of protein interactions at the atomic level, to identifying properties of proteins at different levels of granularity and abstraction, to the scale of mapping connections between proteins across billions of years of evolution.

## Introduction

The sequences, structures, and functions of proteins have co-evolved over billions of years. Protein language models are trained on the outputs of evolution, and generations of ESM models have demonstrated that protein language models develop representations of the biological properties of proteins that drive evolution’s selection of amino acids across life. ESM1 found representations of biological structure and function emerge from training a transformer on protein sequences at the scale of evolution with a language modeling objective (1). The attention patterns of ESM1b contain attention heads that correlate with long-range residue-residue contacts in the protein’s folded structure (2). ESM2 showed that the representations continue to improve with scale and can be projected into atomic resolution three-dimensional structure, matching the accuracy of state-of-the-art structure predictors (3). ESM3 demonstrated that a language model of the joint distribution of protein sequence, structure, and function internalized fundamental principles of structure and function sufficient to generate a fluorescent protein highly divergent from the sequence of any other known protein (4). A number of other language models of protein sequences have been developed (5–13), reflecting the establishment of language modeling as one of the principal approaches to modeling proteins with artificial intelligence.

In every generation of ESM, improvements in the fidelity of representations were linked with the number of parameters and amount of compute used in model training. The representation of the biology of proteins is an emergent phenomenon that arises from training a model to predict the identity of amino acids in the sequence. Biological properties learned and represented by the model develop entirely without supervision.

The first ESMFold model leveraged representations from a language model to achieve atomic-level prediction of protein structure directly from sequences (3), advancing a new paradigm for structure prediction. The ESMFold approach is different from other modern structure prediction models, which, starting from the observation that statistical patterns in sequences of a protein family are predictive of structure (14–20) build on the multiple sequence alignment (MSA) as the primary input (21–24). ESMFold obtained all-atom structure prediction accuracies of RMSD_95_ *<*1.5 Å in backbone residues and *<*2 Å for all atoms without an MSA, leveraging the representations of ESM2.

Here, building on unsupervised language modeling as a substrate for representing protein biology, we present a system for modeling protein sequence, structure, and function. As the foundation of this system we trained ESM Cambrian (ESMC) with a masked language modeling objective at the 300 million, 600 million, and 6 billion parameter scale. ESMC trains on metagenomic sequences, which expands its training dataset by close to two orders of magnitude (from ∼50 million sequences to ∼2.8 billion sequences) relative to the previous-generation ESM2 model. This results in log-linear improvement in the fidelity of representations as a function of the scale of compute used for training.

We have developed ESMFold2, a model for protein structure prediction, which processes ESMC’s representations through a looped architecture and achieves state-of-the-art atomic resolution structure prediction. ESMFold2 delivers both faster and more accurate all-atom predictions of protein interactions, including antibody-antigen interactions, a critical modality for the design of medicines.

We show that it is possible to search the input space of this joint model of sequence and structure to routinely design protein interactions with high experimental success rates, including for therapeutically important molecules, achieving high affinity in a minimal number of experimental trials. Across five targets we demonstrate discovery of miniproteins and single-chain antibodies (scFvs) with nanomolar binding affinities. Increasing inference-time search increases the fraction of experimentally verified binders across modalities and targets, showing that simply increasing the number of digital experiments translates into improved empirical success rates. We performed further biophysical and functional characterization of some of the high-affinity designs. The designs exhibited specific binding to their predicted epitope and are selective for their intended target over close homologs. We also performed functional characterization of a miniprotein and antibody binder for PD-L1 and find that both demonstrate function in a range consistent with therapeutic activity.

Mechanistic investigation of the representations of the ESMC transformer with sparse coding reveals the concepts the model uses to represent proteins. We find evidence of a reduction of protein biology into a system of more basic properties and biological concepts which are organized hierarchically, across physical scales, levels of complexity, and biological abstraction. As a whole this represents a systematic compositional organization that extends from the properties of proteins to themes across evolution, linking the biology of highly divergent proteins. The identified features represent the concepts that ESMC uses internally to represent proteins. These concepts have emerged through unsupervised language model training, independent of prior knowledge. The degree of correspondence observed between this organization and the reductionist understanding of protein biology developed through empirical science suggests that the representation space recovers an organization that is intrinsic to the fundamental biology of proteins.

Using this system of representations of protein sequence, structure, and function, we mapped 6.8 billion proteins, predicting 1.1 billion structures at atomic resolution and identifying structural and functional connections between proteins at the scale of life on earth. Collectively these results demonstrate the power of language modeling for understanding proteins through artificial intelligence, from prediction and design of protein interfaces at the level of atoms, to identification of the intrinsic patterns of rules of composition making up protein structure and function, to systematic discovery of relationships across all natural proteins explored by evolution.

### A scaling law for protein language models

ESMC is a family of transformer models trained on protein sequences using the masked language modeling objective (25, 26).

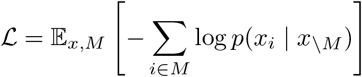

The model is trained to predict the amino acids at randomly masked positions *x*_*i*_ ∈ *M* in the contexts *x*\*M* of protein sequences across evolution. Transformer models (27) trained with this objective develop representations that contain information about the biological properties encoded by the distribution of protein sequences. The latent representations are formed by the activations of each of the neurons in each of the layers of the network as it processes a sequence. Representations are continuous, high-dimensional vectors, one for each position in the protein’s sequence. The transformer processes information through two primary mechanisms: self-attention networks which transfer information between representations of positions in the sequence, alternating with feedforward networks which process the representation at each position individually (Figure 1A). The 300 million, 600 million, and 6 billion parameter ESMC models have 16, 24, and 80 transformer layers, respectively.

**Figure 1.**
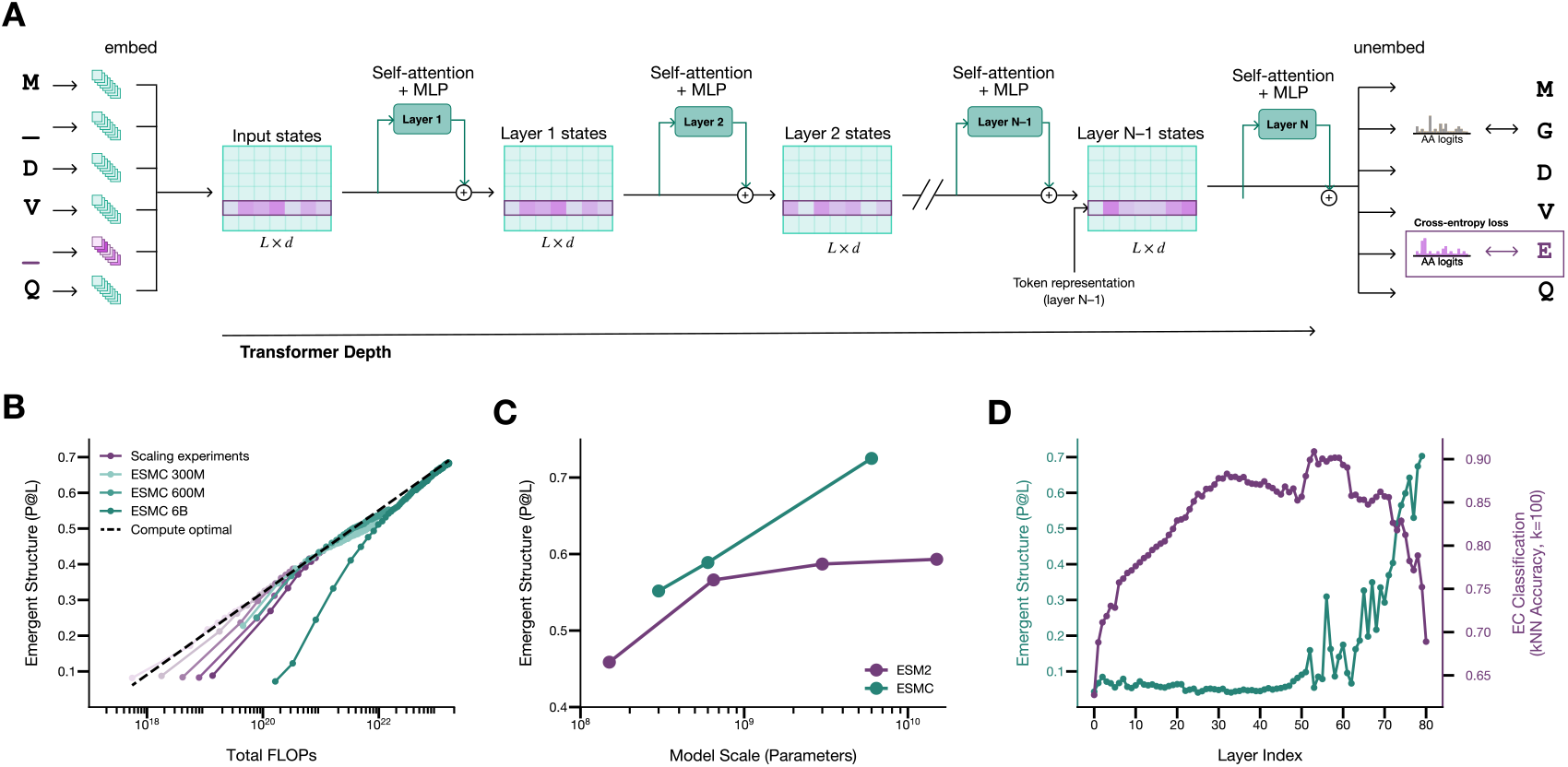
Latent representations of protein structure and function emerge and improve through scaling language modeling. (**A**) ESMC latent space. A discrete sequence of amino acids (tokens) is projected into high-dimensional vectors (2560 dimensions for 6 billion parameter ESMC), one for each amino acid. Representations are processed through successive blocks of self-attention and feedforward layers. A projection from the final layer at each position outputs an unnormalized log probability over the amino acid vocabulary at that position. During training, amino acids are masked from the sequence and the model is trained to predict the missing tokens. The activations of the neural network at every layer constitute the model’s latent representation of the protein. (**B**) Precision of predicting long range (sequence distance ≥ 24) contacts as a function of training compute (FLOPs). A log-linear relationship between training compute and emergence of tertiary structure in the model’s representation space is observed. Performance of scaled training runs (green) follow (*R*^2^ = 0.98) the projection (dashed line) fit through experiments at a small scale (yellow). (**C**) Contact precision for ESM2 and ESMC models as a function of model scale (number of parameters). ESMC shows log-linear improvements with increasing model scale without plateau. (**D**) Structure and function are represented in different layers of ESMC 6B. Organization around function (k-nearest neighbors classification accuracy) peaks between layers 50-60. Information about tertiary structure (precision at L) peaks in the final layers.

ESMC exhibits a scaling law, a log-linear relationship between training compute and the emergence of information about protein structure in the model’s representations. Emergence of this information is measured by computing the accuracy of the transformer attention heads in predicting long-range tertiary contacts in protein three-dimensional structure (2). In development experiments, a series of five models (with dimension 512 to 1536) were trained beyond their point of compute optimality. In Figure 1B, a curve is fit to the development experiments using the maximum achievable contact precision for a given compute budget (P@L-LR = 0.115 × log_10_(FLOPs) −1.98, *R*^2^ = 0.99). Based on the curve, we chose to train three ESMC models at 300 million, 600 million, and 6 billion parameters. The smaller models were then trained slightly past the predicted point of compute-optimality. The largest, a 6 billion parameter model, was trained to the predicted compute-optimal point. The empirical contact precision of the large scale training runs are accurately predicted by extrapolation from the development models (residual between ESMC 6 billion final training point at 1.63 × 10^23^ FLOPs is −0.007 P@LLR), indicating a log-linear relationship between compute and capability.

As the parameter count of the released ESMC models increases from 300 million to 6 billion parameters, contact prediction accuracy improves log-linearly (Figure 1C). This differs from ESM2, where diminishing returns from scale were observed between 650 million and 15 billion parameters. ESM2 was trained on UniRef50 which had a total of 50 million proteins. ESMC substantially increases training data on ESM2 by incorporating metagenomic sequences and an updated version of UniRef. The ESMC training data includes 156 million sequences from UniRef release 2023 02 (28), 2.029 billion from JGI (downloaded 7/2023) (29), and 621 million from MGnify version 2023 02 (30), expanding the number of training sequences to 2.806 billion. These results suggest increasing training data is sufficient for continued returns to scaling compute.

Representations of the ESMC transformer also capture information on protein function. Information about structure and function emerges at different layers of the ESMC 6 billion parameter model (Figure 1D). Long range contact precision is low in the first 40 layers of the model, and rises sharply in the later layers, peaking in the penultimate layer. The representation space also organizes proteins according to function, such that proteins sharing the same enzyme classification (EC) number (31) are mapped to the same neighborhood in the high dimensional space. Classification accuracy for enzyme class, based on the 100 nearest neighbors in a set of 8,493 proteins across 57 EC classes (Appendix A.1.4.2), rises steadily from the earliest layers, peaking in layers 50-60. The differences in per-layer accuracy on these tasks show that the model organizes and represents structure and function differently in its latent representation space.

The function of a protein is correlated with the structural topology it adopts, but ESMC also learns representations of function that are independent of a protein’s fold. The representation space of ESMC contains specific directions corresponding to conserved function that are independent of structure. We curated a dataset of enzymes where the same function (EC number) occurs in multiple distinct protein structures, CATH topologies, Sillitoe et al. (32), suggesting differing but convergent evolutionary trajectories (Appendix A.1.4.3). In this setting, function and structure appear along nearly orthogonal axes of variation (principal components) in the latent space, and linear probes fit to predict EC number on a subset of topologies generalize to predict unseen topologies. Directions that generalize across structural folds and correspond to basic biophysical properties such as stability also emerge in the latent space (Appendix A.1.4.4). These results indicate that there are directions corresponding to aspects of function that are independent of those that encode its structure.

Protein language models develop high-dimensional vector representations through pretraining that capture structural and functional properties of proteins. These representations improve predictably with scale and form a basis that can be used to understand protein biology and design novel proteins.

### Accurate prediction of protein structure from a single sequence

Using the representations of ESMC, we develop a model that obtains state-of-the-art predictions of the structures of individual protein chains and biomolecular complexes. The model, ESMFold2, has an architecture incorporating a simplified core folding module and stable recurrence in its pairwise folding state, enabling the model to leverage recurrence to a greater depth in inference (Figure 2A). Prediction of the interaction between antibodies and their targets particularly benefits from recurrent computation. The model includes the optional input of MSAs. As a whole, this approach achieves a Pareto frontier of accuracy across scales of inference compute including state-of-the-art performance across essentially all benchmarks for protein structure and biomolecular complex prediction.

**Figure 2.**
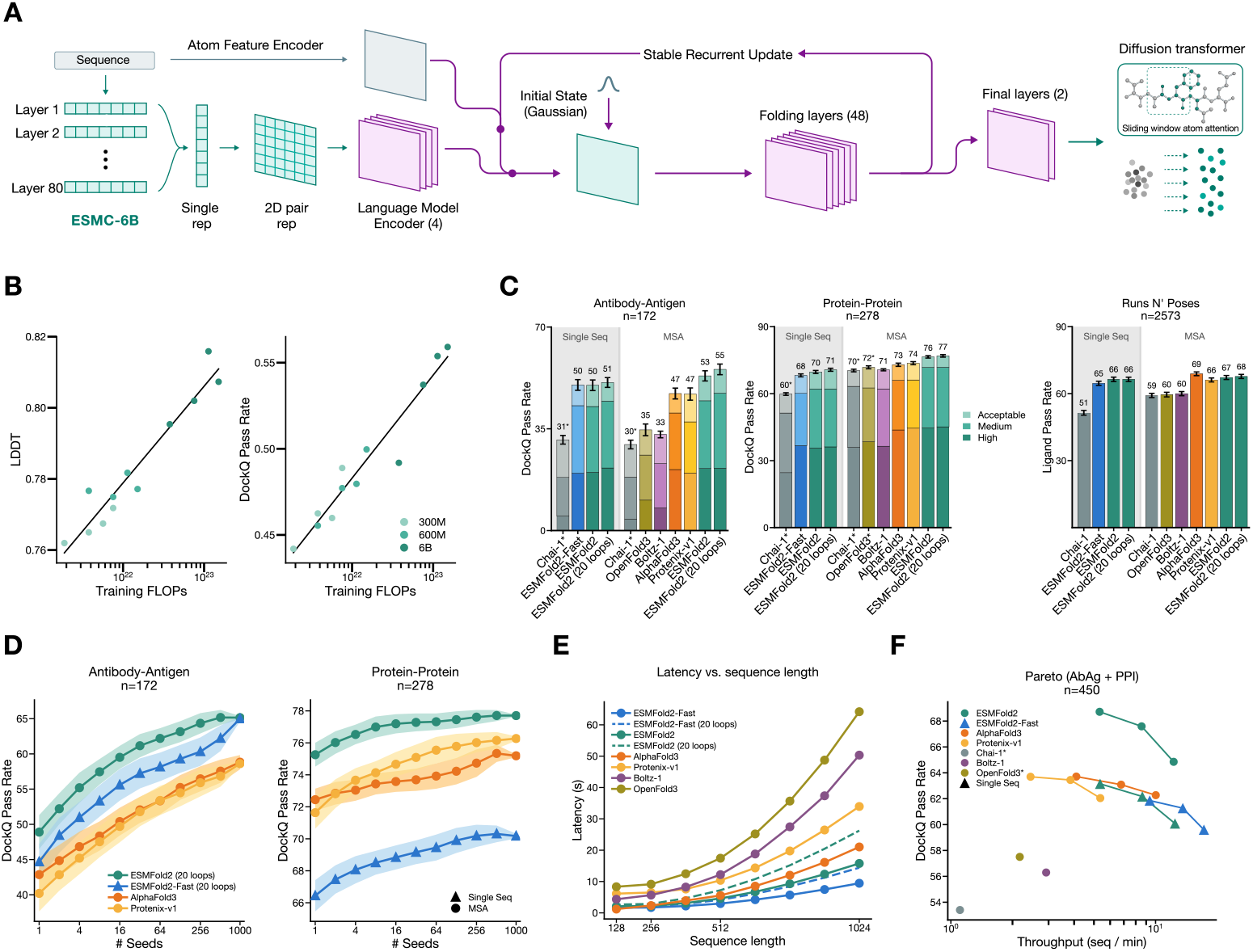
ESMC representations enable high accuracy prediction of protein interactions. (**A**) ESMFold2 architecture diagram. Representations from all layers of (frozen) ESMC 6 billion are pooled into a single representation and projected to a two-dimensional pair representation, which is refined with a 4 layer encoder. The pair state is iteratively updated by 48 folding layers (24 in ESMFold2-Fast), with a stabilized recurrent update (Appendix A.2.5), enabling backpropagation through the loop. Two final folding layers condition a diffusion transformer with sliding-window atom attention, which denoises to atomic coordinates. (**B**) Scaling of structure prediction with language model training compute. Structure predictors trained on representations from ESMC checkpoints (300 million, 600 million, 6 billion parameters) at various training FLOPs. Left: Monomer LDDT on FoldBench (33) protein monomers. Right: protein-protein DockQ pass rate (DockQ ≥ 0.23) on FoldBench protein-protein interactions. A log-linear fit of FLOPs vs. LDDT and DockQ Pass Rate is shown, demonstrating clear returns to sequence pretraining compute. (**C**) Performance on FoldBench Antibody-Antigen, FoldBench Protein-Protein, and Runs N’ Poses (34). Baseline models (23, 35–38) are run locally with the same evaluation pipeline as ESMFold2. Baseline models use 10 recycles and 200 diffusion steps. ESMFold2 is shown with 10 and 20 loops, and uses a truncated diffusion schedule with 68 diffusion steps (Section A.2.11). Error bars / shaded bands show ±1*σ* from bootstrap sampling (Section A.2.10). Performance is shown at 5 seeds and 5 diffusion samples per seed with iPTM selection. Ligand pass rate: LDDT-PLI *>* 0.8 and BiSyRMSD *<* 2Å (Section A.2.10) Single-sequence and MSA modes are separated within each panel. ^*^Denotes model did not produce predictions for all targets; results indicate performance on the subset that folded. Chai-1 folds 148/172 antibody-antigen and 251/278 protein-protein targets, results indicate success rate on this subset. OpenFold3 folds 275/278 protein-protein targets. For Runs N’ Poses, performance is shown on the intersection where all models made valid predictions. (**D**) Inference-time scaling for antibody-antigen and protein-protein folding. Models draw 5 diffusion samples per seed for 1024 seeds. Samples are ranked by iPTM and the top prediction per target is scored. Lines show mean pass rate across targets at each seed budget; shaded bands show ±1*σ* across *n*_boot_ = 300 resamples. (**E**) Inference latency vs. sequence length. All latencies measured on a single H100 GPU with matched recycle/loop/diffusion settings; MSA construction time excluded. ESMFold2 latency at 20 loops shown as dashed line. ESMFold2-Fast and ESMFold2 are the fastest and second fastest folding models, respectively. Chai-1 is excluded as it is significantly slower than other models (110s at sequence length 1024). (**F**) Accuracy-throughput pareto frontier on combined Foldbench antibody-antigen and protein-protein benchmarks. Throughput averaged across sequence lengths 128-1024 (panel E). Points along each line for ESMFold2, ESMFold2-Fast, AlphaFold3, and Protenix-v1 correspond to varying number of recycles or loops (5, 10, and 20). Triangles represent single-sequence mode; circles represent MSA mode. ESMFold2 and ESMFold2-Fast are on the upper right of the Pareto frontier.

Most current structure prediction approaches use the MSA as their primary input. The basic architecture operates on a pair state with communication patterns involving triplets of residues, and projects that state into atomic coordinates (22). For the current generation of models, this is achieved by diffusion of atoms (23). In our approach, building on the first generation of ESMFold, the primary input is the representations of a language model rather than an MSA. Representations from all layers of ESMC are combined through a learned transformation, and injected into the pair state. This is what enables the model to make a prediction of the structure directly from sequence.

In development experiments for ESMFold2, we observed a relationship between the amount of compute used to train the language model and the performance of the folding models. We trained folding models using four intermediate checkpoints from each of the three ESMC language model training runs (300 million, 600 million, and 6 billion parameters). The language model weights were frozen, and structure prediction models with identical architectures were trained on each model’s representations with identical data and amounts of compute. Log-linear relationships are observed between the amount of compute used to train the language model and the performance of the structure prediction models (Figure 2B). This is observed for both monomer structure prediction (LDDT = 0.027 × log_10_(FLOPs) + 0.181, *R*^2^ = 0.92) and protein-protein complex prediction (*P* (DockQ ≥ 0.23) = 0.060 × log_10_ (FLOPs) − 0.832, *R*^2^ = 0.93). This result establishes a direct link between language modeling and atomic-level structure prediction.

The accuracy of structure prediction improves with depth in the number of updates to the pairwise folding state during inference. Significantly faster depthwise computation of the pair state layers is made possible with a simplified module (39), which includes two operations: triangle multiplication, for passing information between pair states, and a feedforward network, which performs computation on each state independently. Notably, attention is unnecessary for pair state processing. Repeated application of pair state layers can increase depth further. Backpropagation through the recursive update enables the model to benefit from deeper recursion during inference. To stabilize this recursion, we adopt an approach developed for looped transformers (40) that projects the pair state through a contractive map, preventing the activations from growing unbounded with repeated loops. During training we backpropagate through two iterations of the folding state. Empirically we find that the models benefit from more recursive applications at inference time.

ESMFold2 predicts protein complex structure with state-of-the-art accuracy from a single sequence. On antibody-antigen complexes, ESMFold2 reaches a DockQ pass rate of 50% ± 2% (error bars on FoldBench represent ±1σ across 200 bootstrap samples of 5 seeds from 25 seeds) from sequence alone, exceeding AlphaFold3’s 47% ± 2% with an MSA (Figure 2C). From single sequences, ESMFold2 folds 70% ± 1% of protein-protein interactions and 66% ± 1% of protein-ligand complexes successfully (error bars on Runs N’ Poses represent ±1σ across 200 bootstrap resamples of the test set).

Providing ESMFold2 with MSA context yields even greater results. With MSAs, pass rates rise to 53% ± 2% on antibody-antigen and 76% ± 1% on protein-protein, surpassing AlphaFold3 in both settings (47% ± 2% and 73% ± 1% respectively). With 20 recurrent loops, antibody-antigen pass rate reaches 55% ± 2%.

ESMFold2-Fast is an inference optimized model trained for single sequence folding. Despite using half as many folding layers (24 instead of 48) and no MSA context, ESMFold2-Fast outperforms AlphaFold3 on antibody-antigen folding (50% ± 2%) and remains competitive on protein-protein (68% ± 1%) and protein-ligand interactions (63% ± 1%). This makes ESMFold2-Fast an ideal choice for high-throughput screening and design.

ESMFold2 benefits from inference time scaling. With increasing number of samples from the model, antibody-antigen pass rate rises from 49% with a single seed to 65% with 1000 samples, and protein-protein pass rate rises from 75% to 78% (Figure 2D). ESMFold2-Fast benefits even more on antibody-antigen structure prediction, with performance rising from 45% to 65% at 1000 seeds, tied with the larger ESMFold2 model. Moreover, unlike other models in Figure 2D, ESMFold2-Fast is not provided MSAs. At all compute budgets, ESMFold2 exceeds the performance of other models.

The simplified pair folding layer and several custom GPU kernels (Appendix A.2.11) reduce the cost of each recurrent loop, so ESMFold2 can run more loops at lower total cost. With 10 loops and 200 diffusion steps, ESMFold2 predicts a 1024 residue structure in 15.8 seconds, 1.3x faster than AlphaFold3. ESMFold2-Fast does so in just 9.4 seconds (Figure 2E).

ESMFold2 has learned a powerful and efficient model of protein structure and interactions. From a single sequence, it folds antibody-antigen and protein-protein complexes at or beyond the accuracy of MSA-based models, and does so faster than any other folding model. Together ESMFold2 and ESMFold2-Fast define the accuracy-throughput frontier across protein complex prediction benchmarks (Figure 2F), making them powerful tools for large scale design and discovery.

### Design of high-affinity protein interactions

Biological function is orchestrated through the formation and dissolution of biomolecular interfaces, from the assembly of complexes and the recognition and propagation of signals to the regulation of biological processes. Binding is the primary mechanism for the activity of therapeutic molecules. Together, the language model and structure predictor specify a general model of protein interactions. We have found that a simple approach, achieved by searching the input space of the model, can design atomic-resolution interfaces between protein molecules and discover proteins that bind their target with high affinity.

This search can be written as a joint optimization over a candidate binder sequence and its target-bound structure

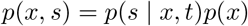

where *x* denotes the candidate binder sequence, *t* denotes the target sequence, *s* denotes the structure of the target-bound complex, *p*(*x*) is the sequence prior learned by ESMC, and *p*(*s* | *x, t*) is the conditional structural model provided by ESMFold2. ESMC guides the search toward sequences compatible with folding and stability, while ESM-Fold2 evaluates whether candidate sequences form the desired interface.

Design optimizes sequences with respect to properties evaluated on their predicted structures. This perspective is shared by a broader class of structure predictor-guided methods, where a folding model is used to guide the optimization of candidate designs (41–44). In principle, many search procedures can be used, including sampling-based methods such as MCMC (45–48). Our approach represents sequences as continuous distributions over amino acid probabilities and optimizes by backpropagation through ESMC and ESM-Fold2. Temperature annealing is used to sharpen amino acid distributions toward discrete sequences (Appendix A.3.1.1). A single-stage optimization over continuous sequences is sufficient to produce high-quality designs.

Binders are designed against five clinically relevant targets: PDGFRβ, EGFR, PD-L1, CD45, and CTLA-4. These targets span receptor tyrosine kinases, immune checkpoints, and cell-surface phosphatases, and are widely studied in oncology and immunology. Designs are of two modalities: de novo minibinders and antibody-derived single-chain variable fragments (scFvs). Minibinders are compact, entirely novel binding scaffolds, whereas scFvs are antibody-derived binders that retain a conserved immunoglobulin framework while requiring design of the complementarity-determining regions (CDRs) that mediate target recognition. CDR design is particularly challenging because these regions are highly variable and conformationally flexible, evolutionary information is limited, and small geometric errors can strongly affect binding.

ESMFold2’s efficiency makes it tractable to search large regions of the design space *in silico* before selecting candidates for experimental evaluation. We used a simple protocol: generate many candidate binders, predict each target-bound complex with multiple ESMFold2 replicas, and rank candidates by their average ipTM, a confidence score for the predicted interface. Sets of candidates for each target–modality pair were generated with two scales of inference compute. More compute was applied in two ways: first, by generating more candidates, and second, by ranking candidates with scores from a larger ensemble of folding models. For minibinders, the smaller and larger sets contained ∼15,000 and ∼67,000 candidates on average, requiring ∼500 and ∼2,400 H100-hours, respectively. For scFvs, the sets contained ∼28,000 and ∼117,000 candidates on average, requiring ∼1,800 and ∼7,700 H100-hours, respectively (Appendix S16). In the higher-compute setting, the ranking ensemble increased from 4 to 19 critics. Eighty-four designs were selected and experimentally evaluated for each target, modality, and compute scale.

High hit rates were achieved for all targets and modalities, and more compute improved experimental outcomes (Figure 3A, left). Hit rates were assessed by biolayer interferometry (BLI, Figure S17, Table S11). A hit is defined by a stringent cutoff for high-confidence binding (Appendix A.3.3). A more inclusive threshold, aligned with the recent literature, is shown for reference. Cohorts of designs from the higher compute experiments achieved hit rates of 36–88% for minibinders and 15–29% for scFvs. Increasing inference compute improved hit rates in 9 of 10 target–modality pairs (Figure 3A, right). Averaged across targets, minibinder hit rates increased from 53.8% to 70.0% and scFv hit rates increased from 12.1% to 21.0%, approaching a twofold improvement in the more challenging antibody-design setting. These results demonstrate that ESMC and ESMFold2 capture the determinants of binding with high fidelity and that scaling computational search improves the quality of solutions.

**Figure 3.**
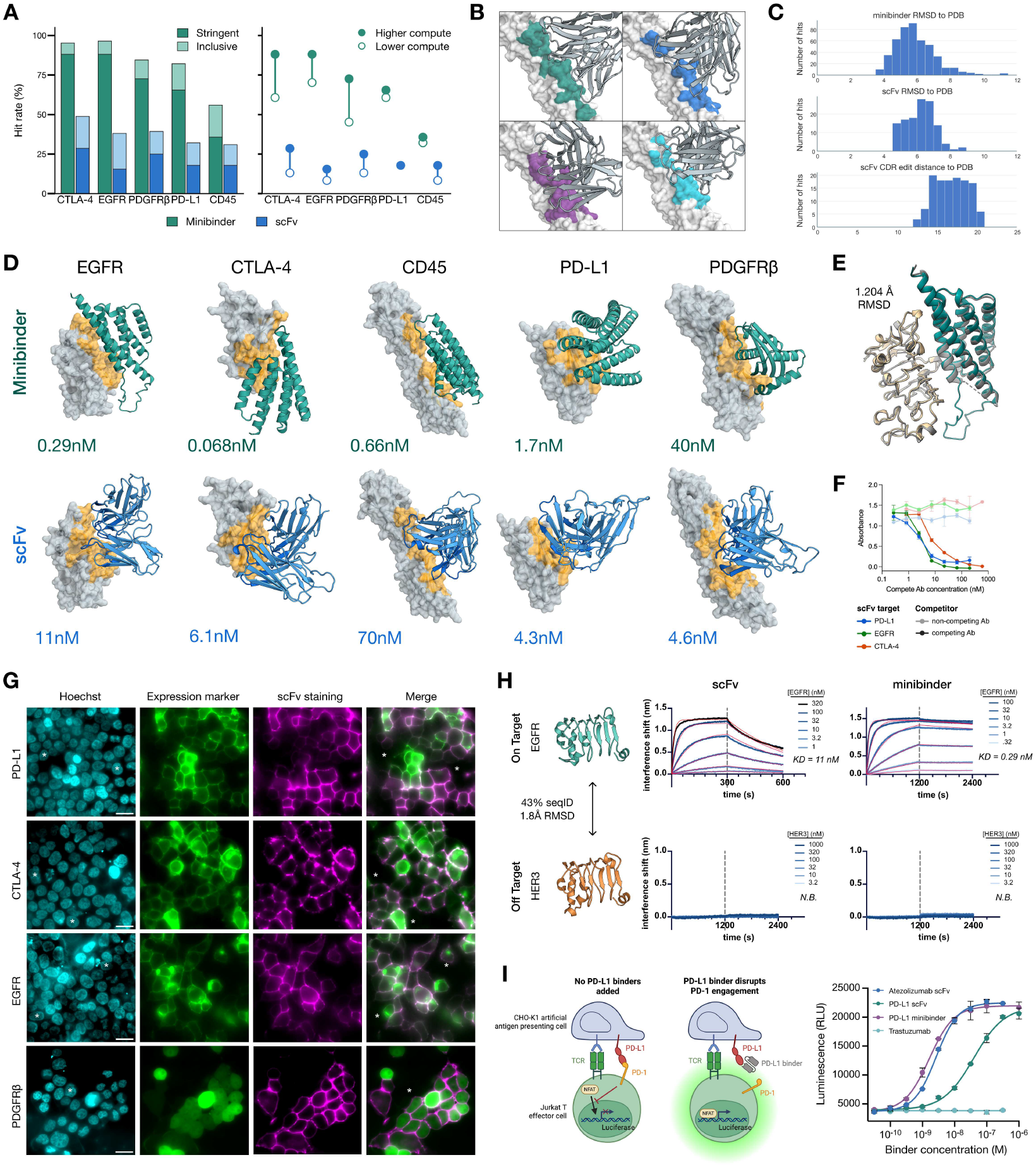
Searching ESMC representations yields high-success-rate therapeutic binder design. (**A**) Laboratory hit rates for designed binders. Left: BLI-measured hit rates for minibinders and scFvs against CTLA-4, EGFR, PDGFR*β*, PD-L1, and CD45. Shading indicates success at different thresholds (light: inclusive weak-binder threshold, dark: stringent threshold). Right: Effect of inference-compute used for generating and ranking design candidates. Open circles indicate lower-compute design pools, and filled circles indicate higher-compute pools generated with ∼4.5× greater total GPU-hours. (**B**) Diverse binding modes among validated CD45-binding scFvs. Predicted structures show designs engaging CD45 through distinct binding orientations and interfaces. (**C**) Novelty analysis of validated binders. Top, middle: distribution of minimum RMSD to nearest PDB structure. Bottom: Edit distance of CDR regions to nearest PDB antibody. (**D**) High-affinity binders for each target and modality. Target proteins are shown as gray surfaces, designed minibinders and scFvs are shown as cartoons, and modeled interface regions are highlighted in orange. BLI-measured affinities are indicated for each design. (**E**) Structural determination of EGFR-minibinder complex by cryo-electron microscopy (cryo-EM) to 3.8 Å resolution. EGFR domain III (left) and the minibinder (right) are shown here in ribbon cartoons. Gray: Experimentally determined structure. Tan and teal: The computational model aligned to the experimental data via EGFR domain III. Pairwise atomic RMSD between the experimental and computational minibinder models is indicated and shows high fidelity between the predicted and realized complex structure. (**F**) Competition enzyme-linked immunosorbent assay to characterize scFv epitopes. Designed scFvs bound surface-coated antigens in the absence or presence of therapeutic biosimilar antibodies targeting the designed epitopes: atezolizumab (anti-PD-L1), cetuximab (anti-EGFR), ipilimumab (anti-CTLA-4), or non-competing trastuzumab (anti-HER2). Error bars indicate 1 SD across duplicate measurements. Reduced signal in the presence of the competing antibody supports intended epitope engagement. (**G**) Live-cell immunofluorescence staining by designed scFvs for characterization of binding specificity. Human embryonic kidney 293T cells were transfected with constructs harboring the indicated targets tagged with green fluorescent protein (GFP, green). Cells were then stained with Hoechst 33342 (cyan) and Twin-Strep-tagged scFvs complexed with Strep-Tactin XT-DY649 (magenta), followed by imaging on an epifluorescence microscope. The GFP variant is fused to the target of interest in all cases except for PDGFR*β*, where it is translated separately via an internal ribosome entry site. The fourth column is a merge between the green and magenta channels. White asterisks denote untransfected cells or cell clusters, which show no detectable scFv staining. Scale bars, 20 *µ*m. (**H**) Characterization of binding selectivity by BLI. Binding affinities of an anti-EGFR scFv and minibinder were measured against the intended antigen, EGFR domain III, and the homologous antigen HER3. Left: Comparison of structure, sequence identity and backbone RMSD between the homologous domains. Top right: on-target BLI sensorgrams and measured dissociation constants (*K*_*D*_). Bottom right: off-target BLI sensorgrams. N.B. denotes no binding. (**I**) Cellular immune checkpoint blockade assay. Left: Schematic of the assay mechanism. Addition of PD-L1 binders that can disrupt its binding to PD-1 enables the nuclear factor of activated T-cells (NFAT) to enter the nucleus in Jurkat T-cells and drive the transcription of a luciferase reporter. Right: Cells were treated with the indicated PD-L1 binders or trastuzumab (negative control). The designed PD-L1 minibinder and scFv restored signaling in a dose-dependent manner, with estimated functional IC_50_ values of 1.6 nM and 39 nM, respectively, compared with 2.6 nM for an atezolizumab scFv control. Luminescence is plotted as relative light units (RLU). Error bars, 1 SD across duplicate measurements.

Validated binders are diverse in sequence, structure, and binding interface (Appendix A.3.4), indicating that the model recovers multiple distinct solutions within each target–modality pair rather than converging on a single binding mode. This diversity was observed for both minibinders and scFvs, spanning a range of sequence lengths, structural topologies, binding orientations, and interfaces. Figure 3B shows predicted structures for representative confirmed anti-CD45 scFvs, illustrating varied modes of engagement with the same target.

The designs are distinct from known proteins and antibody structures (Figure 3C). Search of minibinder sequences against `nr` using BLASTP (49) identified significant matches (E-value *<* 10^−3^) for just 5 of 438 binders, all with sequence identities *<* 34%. Similarly, all validated scFvs are at least 10 edits from their nearest antibody in the PDB (50) when comparing concatenated CDRs from each variable domain against all such concatenations in the PDB. Structural searches further showed that the designed interfaces differ from known PDB interfaces (Appendix A.3.4).

Therapeutic utility depends on more than biophysical binding. Therapeutic candidates must combine multiple properties, including robust expression, stability and solubility, binding affinity and specificity, engagement of the intended epitope, and functional modulation of the target. We evaluated designs across biochemical, biophysical, and functional axes. Minibinders and scFvs that demonstrated high apparent affinities during screening were expressed in *E. coli* or by *in vitro* translation and purified for further characterization. Size-exclusion chromatography demonstrated that the purified binders are homogeneous and non-aggregated (Figure S18).

For all targets and design modalities, we designed proteins that achieve high-affinity binding (Figure S22). Designs for each target and modality are shown in Figure 3D, with BLI-measured affinities ranging from 68 pM to 70 nM. Low-nanomolar affinities are generally considered therapeutically relevant, as they correspond to pharmacokinetically achievable concentrations *in vivo*. Binders within this regime were observed across all targets and both modalities.

Designs engage the intended epitopes. Structural validation by cryo-EM confirmed the modeled binding pose of an EGFR minibinder (Figure 3E, Figure S19). The experimentally determined complex aligned closely with the predicted target-bound structure, with an RMSD of 1.204 Å, within the experimental uncertainty of the 3.8 Å reconstruction. In epitope competition assays, binders for PD-L1, CTLA-4, and EGFR exhibited reduced antigen binding in the presence of biosimilar therapeutic antibodies targeting the designed epitope: atezolizumab for PD-L1, ipilimumab for CTLA-4, and cetuximab for EGFR (Figure 3F; Figure S20). These results indicate that the design process realizes target binding at the intended epitope, consistent with the specified interaction geometry.

Binders specifically engage their intended cell-surface targets. In live-cell immunofluorescence assays designed scFvs produced strong staining of target-expressing cells with no detectable staining of untransfected cells (Figure 3G), suggesting minimal reactivity to the cell surface proteome. Similar target-dependent staining was observed for minibinders across the same target panel (Figure S21). Specificity was further tested against close homologs of the design targets. Anti-EGFR and anti-CTLA-4 binders bound their intended antigens with nanomolar affinity by BLI, but showed no detectable binding to the corresponding homologous domains in HER3 and CD28, respectively. Representative anti-EGFR scFv and minibinder results are shown in Figure 3H, with corresponding anti-CTLA-4 results in Figure S22. Together, these results show production of binders with target-specific recognition, engaging the intended antigen in its cellular context while discriminating against closely related homologs.

Disruption of the PD-1/PD-L1 immune checkpoint is a highly successful treatment option for lung cancer, melanoma, and an increasing number of cancers. Atezolizumab, for example, is an FDA-approved biologic targeting PD-L1. Our PD-L1 minibinders and scFvs target the same epitope as atezolizumab, so we wondered whether these designed binders produce functional activity consistent with this therapeutic mechanism. These binders were assessed in a cellular functional assay using a co-culture system in which PD-L1 engagement of PD-1 suppresses T cell receptor signaling in Jurkat T cells, measured through an NFAT-driven luciferase reporter. The PD-L1 minibinder and scFv restored signaling in a dose-dependent manner, with estimated IC50 values of 1.6 nM and 39 nM, respectively, compared with 2.6 nM for an atezolizumab-derived scFv control (Figure 3I). All three reached similar maximal levels of pathway activation. These results demonstrate that the designed binders not only engage PD-L1, but also relieve PD-1-mediated suppression of T cell signaling.

ESMC and ESMFold2 provide a system for searching and designing protein interfaces. This search procedure yields minibinders and scFvs that bind with high affinity, engage intended epitopes, and discriminate close homologs. This is achieved across targets and modalities using only the target sequence as input and without task-specific retraining, which indicates that the model captures general determinants of molecular recognition. Inference-time scaling further improves design outcomes, showing that the efficiency of ESMFold2 can be converted into higher-quality experimental candidates through expanded computational search.

### The organization of the ESMC latent space

The selection process that determines the amino acids that occur at each position in every protein is driven by structural and functional constraints—the creation of specific physical and biochemical environments that make reactions energetically favorable, stabilize particular structures or interactions, or facilitate transport to particular locations in the cell. By learning to predict the amino acids evolution selects across differing functional contexts, ESMC could develop representations of the more basic properties that give rise to function. Using the techniques of mechanistic interpretability, developed to understand the workings of large language models of text, we find evidence within the latent representation space for a reduction of protein biology into more fundamental principles across levels of biological complexity.

The concepts used by language models to organize their representation space are difficult to identify since they are represented through superposition in the high dimensional activations of the model. Language models encode concepts using directions in the representation space (51), but the individual dimensions of the representations are polysemantic: individual coordinates activate non-specifically and diffusely because they participate in multiple superposed features (52–54).

The field of mechanistic interpretability has developed techniques to identify the monosemantic features that organize the representation space of language models. These techniques rely on the sparse coding to identify a basis in which a few dimensions are sufficient to approximately reconstruct most representations (55). Sparse autoencoders (SAEs) are a sparse coding method that is effective for identifying monosemantic features corresponding to interpretable concepts in large language models (56–62) and have also been applied to find biological concepts in protein language models (63–65). Since sparse coding techniques are fully unsupervised, the organization they reveal is inherent in the model.

We train SAEs on the representation vector of each amino acid in its context, projecting the vector into a sparse higher-dimensional feature space. To represent a protein as a single vector, the individual feature vectors are aggregated by taking the maximum activation of each feature across the sequence. SAEs are trained for every layer of the 300 million, 600 million, and 6 billion ESMC models, using 8 billion tokens drawn from the same data distribution used to train the language model. A separate SAE is trained for every layer across a range of dimensions of the feature space (2^13^, 2^14^, 2^15^, 2^16^, 2^17^) and sparsity levels (8, 16, 32, 64, and 128 features active for each amino acid). Features activate at a range of specificities across proteins depending on the dimension of the feature vector, the activation threshold, and the sparsity (Appendix A.4.2.1). Our analysis focuses on the SAE with 2^14^ features and 64 active per amino acid trained on representations after the 60^*th*^ layer in the transformer (3/4th depth), near the peak for enzyme classification in Fig. 1D.

Features from the SAEs reveal intrinsic concepts that compose ESMC’s representation space. SAEs are unsupervised, so after training, an agentic system is used to hypothesize the biological concepts to which they correspond. Agents make inferences from the known structure and function of 195 thousand non-redundant proteins that have been characterized in SwissProt. For each feature, agents are provided a set of proteins, annotations, and sequence positions on which the feature activates, and follow a five-fold hypothesis generation and verification procedure to develop text summaries (Appendix A.4.3.1). These descriptions are used for classification and interpretation of the features throughout.

Features span concepts across the hierarchy of protein biology, from primary sequence to complex function. Figure 4B shows examples of features classified by an agent based on their text descriptions (Appendix A.4.3.2). At the level of primary structure (Figure 4B, first row), there are 88 features that activate on specific amino acids (e.g. tryptophan, threonine, cysteine) or represent classes of amino acids (e.g. aromatic or negatively charged side chains). The model has also learned the local interactions between residues which stabilize protein backbones into their two primary low-energy configurations, alpha helices and beta sheets (Figure 4B, second row). The feature space captures more than just the presence of a particular secondary structure class, but almost 2,000 specific components that encode their context. For example, features capture distinct backbone configurations at helix end-caps, the edges of beta strands, and periodic signals at positions stabilized by hydrogen bonds. The loops and turns that form the transitions between ordered secondary structures, providing local backbone flexibility and enabling the global topology to form, are also seen clearly in the feature space.

**Figure 4.**
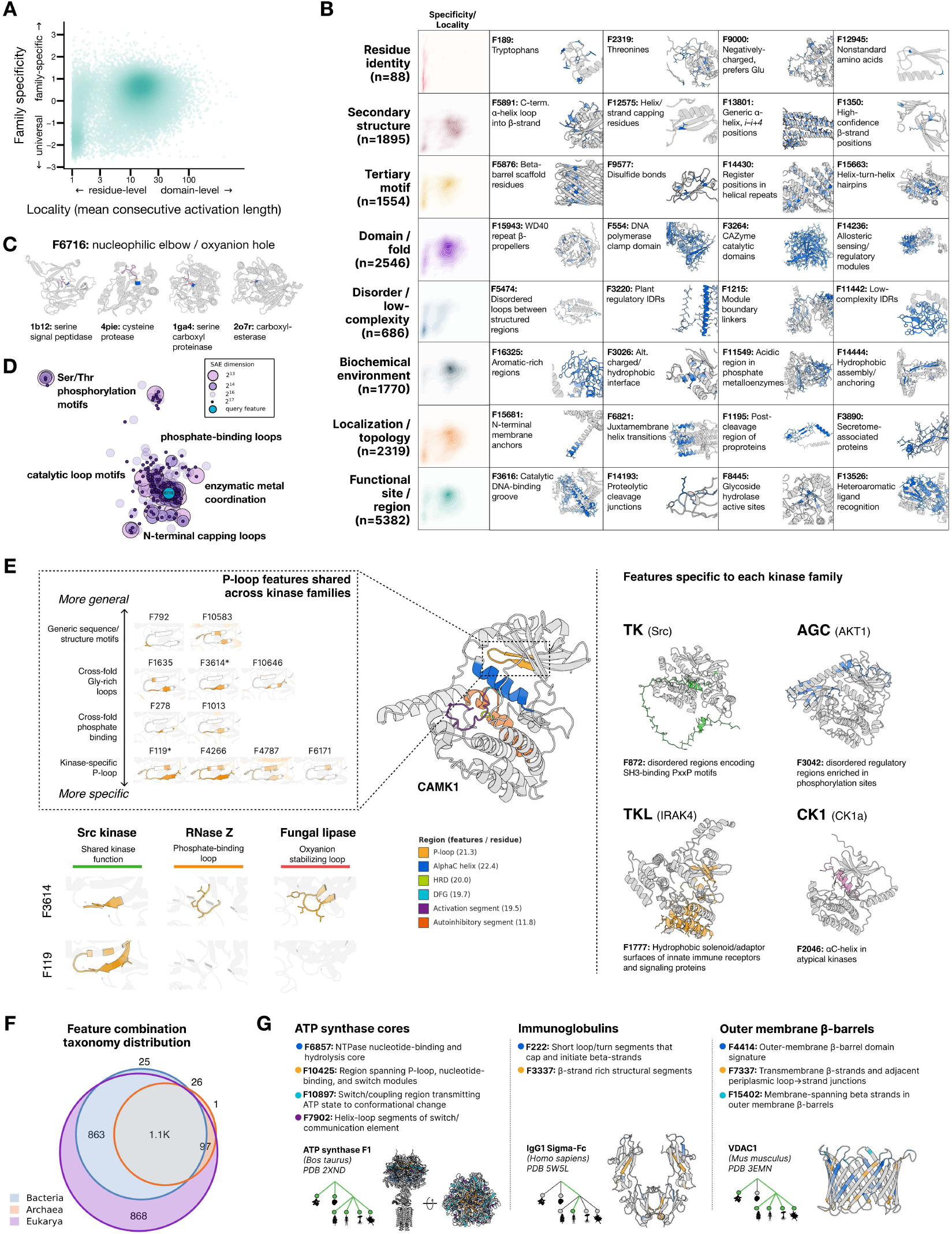
The ESMC latent space contains a reduction of protein biology to biochemical, structural, and functional principles across levels of biological complexity. (**A**) Sparse autoencoders (SAEs) trained on ESMC representations learn features corresponding to primary directions in the latent space. All 2^14^ = 16, 384 features from an SAE trained on layer 60 representations of ESMC 6 billion are plotted according to their specificity (universal to family-specific) and the local granularity of the activation within a sequence (residue-level to domain-level). (**B**) Examples of features spanning levels of biological complexity (top to bottom). Features in each category occupy different regions of the specificity-granularity space (left). (**C**) Four proteins which activate F6716, the nucleophilic elbow catalytic motif, across diverse topologies, reactions, and nucleophile residues. (**D**) Feature neighborhood around F6716 in the SAE decoder space for multiple feature dimensionalities: 2^13^, 2^14^, and 2^16^, shown as different size dots. (**E**) Feature compositionality in protein kinases. Top: 11 features with varying specificity in the phosphate-binding P-loop. Middle: Key functional regions of calcium/calmodulin-dependent protein kinase type 1 (CAMK1, ESMFold2-predicted structure). Kinase-related features (*n* = 531, across 35 proteins, 7 kinase classes) are concentrated in conserved catalytic regions. Bottom: Cross-protein activation of two P-loop features: F3614 (general, flexible loops in diverse proteins); F119 (specific, kinase P-loop). Right: Family-specific features for four kinase subfamilies, largely capturing non-catalytic regulatory elements. (**F**) Distribution of feature combinations across domains of life based on 81 reference proteomes. (**G**) Topic analysis showing feature combinations used across different branches of life. Activation of the highest-weighted features in each are shown on PDB structures. Tree schematics show the taxonomic distribution of each. Left: universal (ATP synthase); Middle: lineage-specific (immunoglobulins); Right: cross-lineage (outer-membrane *β*-barrels).

At a higher level of complexity, features represent tertiary interactions where residues or substructures far apart in sequence are brought together in three-dimensional space. Locally, these are represented as chemical interactions such as disulfide bonds and salt bridges, metal coordination sites and stabilizing hydrogen bond networks (Figure 4B, third row). A large fraction of the 1,554 features in this category capture different types of helix packing, where specific residues on opposite helices come together to form tight bundles. At the level of domain topology—the specific three-dimensional arrangement of helices and sheets—folds which are repeatedly used across biology such as beta-propellers and helix-turn-helix DNA binding domains have features which activate across the entire domain (Figure 4B, fourth row).

There are also features (Figure 4B, fifth row) which capture the absence of structure: regions of intrinsic disorder that do not form a consistent ordered structure and range from just a few residues to entire domains. The functions of these disordered regions are often uncertain, but are widespread in protein biology and are essential for signaling, regulation, and mediating binding interactions (66–68). SAEs capture the diversity of these regions, using 686 individual features (5–10% of the budget of features) to represent disordered regions with different properties.

Protein function is mediated not only by topology, but by the formation of local biochemical environments that enable their activity and interactions (69, 70). These are represented by 1,770 features, including hydrophobic or aromatic regions that facilitate binding, alternating polar and non-polar residues in amphipathic helix packing interfaces, acidic patches, or charge-dense stretches that create favorable electrostatic environments (Figure 4B, sixth row). These features are among the most difficult to characterize based on database annotations, as they may not be associated with a specific function but simply amino acid composition and structural context.

There are features for more complex mechanisms. Almost 50% of the total features correspond to conserved functional sites and substructures that are found across biology due to their common biological roles, such as DNA or carbohydrate processing, energy production and transduction, signaling, or structural scaffolding. Many of these (n=2,319) capture the features which determine a protein’s location or transport within the cell, such as organellar localization signals, or its orientation with respect to membranes or other cellular structures (Figure 4B, seventh row). Others (n=5,382) reflect post-translational modification processes such as the sites of phosphorylation or cleavage by proteases during protein maturation. Still others reflect general classes of function such as transferases or molecular transporters, independent of the specific molecules they act on or the structural fold (Figure 4B, eighth row).

Features form a basis for the model’s representation of proteins. The feature space is organized hierarchically where simpler features are more universal and specificity increases as complexity increases (Figure 4B, density plots), and each protein uses features from all levels of complexity (Appendix A.4.3.2). Even complex features are shared across protein families that have a common underlying mechanism. An illustrative example is the nucleophilic elbow, a conserved catalytic motif which precisely positions a nucleophilic residue (serine or cysteine) relative to a substrate in such a way as to destabilize the ground state and initiate nucleophilic attack. This local biochemical motif appears to have evolved many times across diverse proteases and hydrolases in distinct structural contexts (71). Nonetheless, a single feature activates on the nucleophilic elbow on 75 out of 99 relevant enzymes, spanning 25 distinct folds (Figure 4C, Appendix A.4.3.3).

Each SAE feature corresponds to a direction in ESMC’s representation space, and similar directions represent similar concepts. Figure 4D shows the neighborhood of all features whose directions have a dot product of at least 0.1 with the nucleophilic elbow feature, projected into two dimensions using UMAP (72). The neighborhood is biologically coherent, containing other flexible catalytic loops and phosphorylation motifs. Features from SAEs trained on the same representations with larger or smaller feature dimensions show substantial overlap between their directions, indicating that all SAEs capture similar important axes of variation in the model’s representation space. Depending on the size of the feature space, a different number of features may be allocated to cover these directions—in the top right, one feature in the 2^13^ space corresponds to ten in the 2^16^ space.

Complex functional mechanisms are represented through the composition of many features. Kinases are an important class of signaling molecules whose dysfunction is linked to cancer and autoimmune diseases. The core catalytic machinery, which transfers a phosphate group from ATP to the substrate, is made up of several important subcomponents (Figure 4E, middle). These include a flexible loop for binding the phosphate (the “P-loop”), two long activation loops which contain key sequence motifs (HRD and DFG) that must be positioned correctly for catalysis, and a regulatory αC helix which acts as a conformational switch.

Each of these components is represented by many features from across the complexity scale, from universal to kinase-specific concepts (Figure 4E, top left). For example, the residues of the P-loop activate features capturing generic loops and coils (F792, F10583), glycine-rich loops which confer local flexibility across many families (F1635, F3614, F10646), phosphate-binding in diverse proteins (F278, F1013), and finally kinase-specific P-loops (F119, F4266, F4787, F6171). The difference in specificity can be seen directly by comparing two of these features, F3614 and F119 (Figure 4E, bottom left). The more universal feature, F3614, activates on similar flexible loops across diverse families such as kinases, ribonucleases, and lipases, while F119 activates only on kinases. Similar composability is observed in the other kinase subregions, enabling the model to represent complex functional machinery with a relatively small number of concepts.

The activation of individual features can be directly correlated to activity in some proteins. In Src kinase, just 108 features are sufficient for a linear model to predict the local activity landscape around wild-type with high accuracy (Spearman ρ = 0.74), and many of these features activate in mechanistically important regions. Single point mutations that disrupt the enzyme’s function often reduce the activation of these features, for example every mutation of G277 in the P-loop reduces the activation of P-loop feature F278, and the magnitude of that reduction correlates strongly with the mutation’s effect on kinase activity (ρ = 0.73). Features can also respond to mutations many residues away (e.g. F10351 and F8957), reflecting 3D contacts and long-range allostery (Appendix A.4.4.2).

Across the other major eukaryotic kinase families, the same set of core features fire on the catalytic domain while additional features are used to represent each family’s unique functional context (Figure 4E, right). These features primarily correspond to regulatory context rather than catalytic activity. Tyrosine kinases such as Src are regulated by their SH3-binding domains (F872), AGC kinases like AKT1 are regulated by phosphorylation in their C-terminal region (F3042), and tyrosine kinase–like proteins such as IRAK4 are critical mediators of innate immunity (F1777). Casein kinase 1α (CK1α) does uniquely activate features in its catalytic domain that other kinases do not, particularly in the αC helix (e.g. F2046). This may reflect the fact that CK1 is constitutively active and does not rely on the activation loop phosphorylation and αC helix conformational switching mechanisms of most kinases (73). Through composing structural, biochemical, and regulatory features, the model captures the subtleties of functional modulation both within and across families.

Feature co-occurrence patterns capture biological associations between proteins as well as within them. Proteins do not act in isolation, but instead are components of intricate cellular systems where they interact with diverse molecules such as other proteins, DNA and RNA, metabolites, and ions. Feature pairs identified in the PDB can be used to predict interactions in a held-out dataset of proteins from high-confidence protein-protein interaction experiments despite low sequence similarity (AUROC 0.855; Appendix A.4.4.3), with many capturing characterized interactions such as the TCR-MHC I interaction (F14846, F12376) that initiates the immune system’s T cell activation pathway.

Systematically analyzing feature activation patterns at scale identifies features which are commonly composed with one another to produce specific functions. Using non-negative matrix factorization, we identify ∼3000 weighted combinations of features that together reconstruct the functional space of 81 reference proteomes (74) from across the domains of life. Many of these combinations are shared across all domains (n=1103; Figure 4F), representing universal functions such as ATP synthase catalytic machinery (Figure 4G, left). Others reflect more specific functions such as immunoglobulins, a key component of vertebrate-specific adaptive immunity (Figure 4G, middle). Some combinations reflect key evolutionary events, including the ancient endosymbiosis event between an archaea-like organism and a bacteria from which all eukaryotes originated. The record of this event is found in eukaryotic organelles, mitochondria and chloroplasts, which exhibit a unique set of proteins that are otherwise only found in bacteria. This is also captured in the SAE space by 863 feature compositions, including many found in β-barrel outer membrane proteins (Figure 4G, right). For each of these examples, the feature space provides an interpretable decomposition into subcomponents which contribute structurally or biochemically to the overall functional protein.

The organization of biology is hierarchical and can be described at different scales of complexity and levels of abstraction. The ESMC representation space contains features that map to each level of the hierarchy from primary to tertiary structure, from local biochemistry to complex protein functions, and extending to interactions between proteins and functional themes across evolution. The patterns of co-occurrence between features across this hierarchy are consistent with the way in which complex structure and function are realized through the composition of more basic biological building blocks. The representation space thus reflects a systematic reduction of protein biology which closely mirrors the understanding developed through decades of molecular biology research, despite emerging through the training of a large language model on protein sequences alone.

### An atlas of protein structure and function spanning life

Using ESMC we construct an atlas mapping the structure and function of 6.8 billion unique protein sequences, resolving 1.1 billion structures, and identifying structural and functional relationships between proteins at the scale of life on Earth. The atlas contains a comprehensive view of protein sequences spanning the three branches of life, from vertebrates to micro-organisms, as well as metagenomic samples across a wide range of biomes.

The sequences that make up the atlas are sourced from eight publically available databases: the UniProt archive (UniParc (75)), the Searchable Planetary-scale mIcrobiome Resource (SPIRE, (76)), MGnify (30), the Unified Human Gastrointestinal Genome (UHGG (77)), and four datasets from JGI’s Integrated Microbial Genomes (IMG) sequencing and metagenome-assembled genomes (MAGs) (78, 79). The bulk of the data, 5.6 billion sequences, is from metagenomic samples in SPIRE and MGnify, which are dominated by human and animal gut, aquatic, and soil microbiomes but also contain a variety of less frequently sampled environments, both natural and human-associated, such as wastewater, agriculture, and the built environment (Fig. 5A). Taxonomy is known for the remaining 1.2 billion sequences based on the source database metadata. The majority are prokaryotic and are drawn from a variety of sources, including UniParc, UHGG, and IMG. Among eukaryotic samples, most come from UniParc while metagenomics databases contribute a smaller fraction of sequences but many of which are sampled from less well-characterized organisms. Almost all viral sequences are metagenomic, largely derived from bacteriophages.

**Figure 5.**
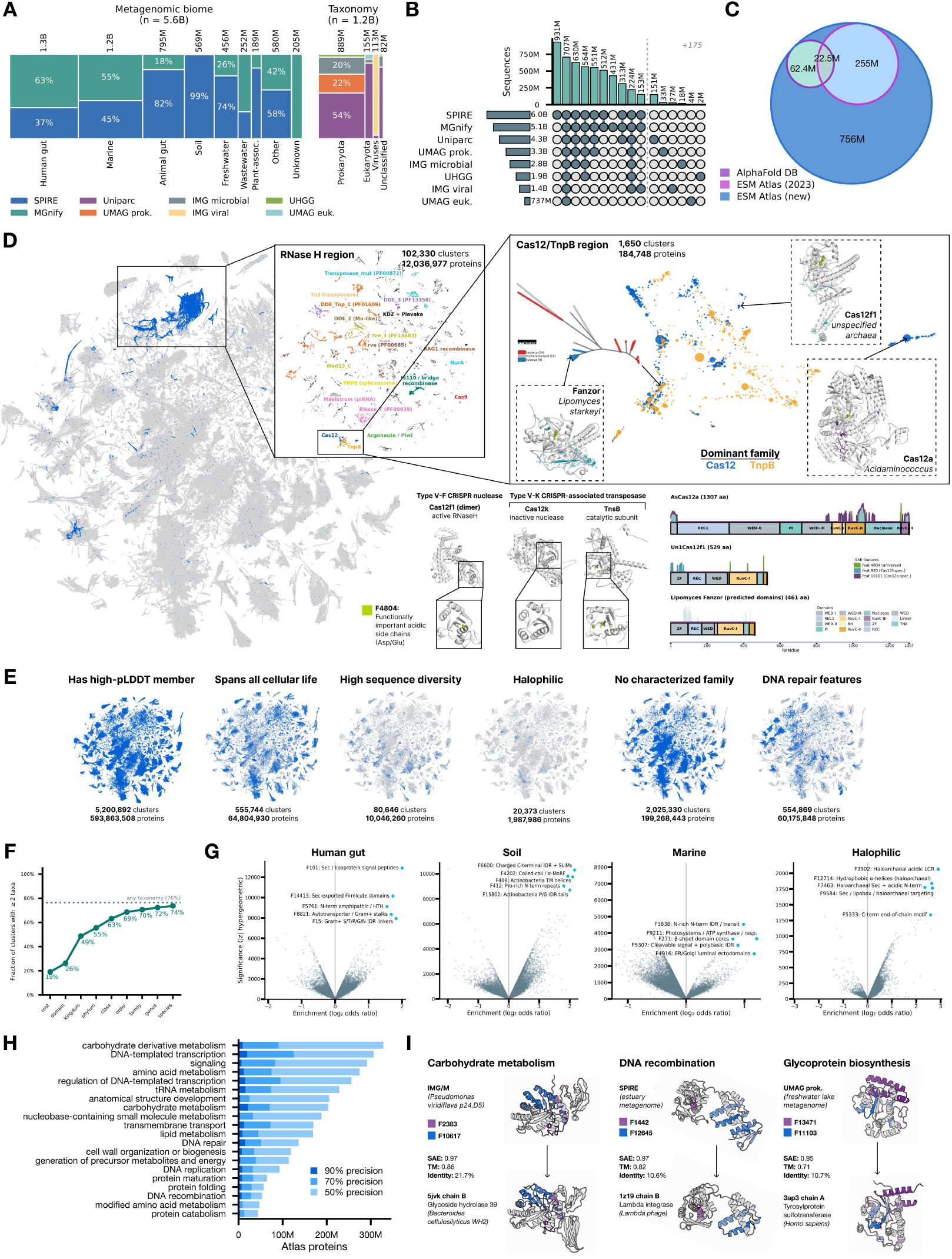
An atlas of protein structure and function spanning life. (**A**) Sequence sources for the ESM Atlas. SPIRE and MGnify are grouped by biome and the remaining six databases are grouped by taxonomic domain. (B) Sequence clusters overlap between source databases at 70% identity. Databases are shown in rows, with the marginal sequence counts as horizontal bars. Each column represents a specific combination of databases, indicated by the filled circles, and the vertical bars show the number of sequences in clusters that contain each of the databases in the specified combination. The top 10 combinations are shown along with the single-database cluster counts (175 combinations not shown). (**C**) Structure coverage in the new ESM Atlas compared to AlphaFold Database and the previous ESM Atlas from Lin et al. (3). Circles represent the number of clusters with at least one sequence present in each predicted structure database. (**D**) Biological organization according to the ESMC latent space. Left: Representatives of 7.7 million SAE similarity clusters with ≥ 50 members projected into 2D with UMAP. Clusters associated with the ribonuclease H Pfam clan (CL0219) are highlighted blue. Middle top: Projection of RNase H clusters colored by the most common Pfam family annotation. Right top: Projection of the region containing Cas12/TnpB families. ESMFold2 predicted structures are shown for representatives of highlighted clusters: a Fanzor/TnpB cluster (with phylogenetic tree shown for all 53 cluster members) as well as clusters containing canonical Cas12f1 and Cas12a. Structures are highlighted with the activations of features 4804 (green, universally active in region), 920 (cyan, specific to Cas12f), and 10161 (purple, specific to Cas12a). Middle bottom: Activation of feature 4804 (acidic residues in the DDE triad) on Cas12f1 (active), Cas12k (inactive), and TnsB (active). Middle right: Domain architectures and feature activation patterns of F4804, F920, and F10161 on each of Fanzor, Cas12f1, and Cas12a. (**E**) Biological themes visualized across the atlas. From left to right: SAE clusters with at least one folded structure with mean pLDDT *>* 0.7; clusters with a lowest common ancestor at the cellular root level; clusters with a mean pairwise sequence identity of less than 30%; clusters with majority of members from high-salinity biomes; clusters where no member is annotated with a Pfam domain of known function; and clusters containing features associated with the DNA repair biological process at 70% precision. (**F**) The taxonomic breadth of SAE clusters, plotted as the proportion which contain proteins from at least two distinct taxa at each level of the taxonomy tree. (**G**) SAE feature enrichment in selected biomes relative to the full atlas. Statistical significance (based on hypergeometric z-score) is plotted against the effect size (log-odds ratio) and the top five features by combined score (significance × enrichment) are highlighted. (**H**) Number of proteins in clusters with high activation of features associated with specific biological processes from the Gene Ontology. Features are selected at 50%, 70%, and 90% precision for each process on Swissprot and then used to identify clusters in the Atlas. (**I**) Examples of 90%-precision features that activate on both unannotated metagenomic proteins in the Atlas and known proteins from the PDB. One example from each of three processes, selected by SAE similarity to annotated clusters, is searched against the PDB using Foldseek and the top hit is shown with TM-score and sequence identity.

Source databases contain substantial overlap, but they also individually contribute to the biodiversity of sequences contained in the atlas. Figure 5B shows the numbers of sequences that share a 70% sequence cluster with a sequence from another database (Appendix A.5.2). The majority of clusters contain sequences from more than one database, and 708 million sequences are in clusters with members from all eight. Each individual database contributes millions of sequences that do not cluster with any sequence from other sources. Even the two large metagenomic databases that sample from similar distributions of biomes, SPIRE and MGnify, are differentiated.

We predict structures for the entire 1.1 billion representative sequences with ESMFold2. This substantially expands the structural coverage of the protein universe, exceeding the cumulative number of predicted structures in AlphaFold Database (80) and the original ESM Atlas (3) by 835 million and 336 million, respectively, with 418.5 million of these structures predicted with high confidence (mean pLDDT *>* 0.7). In total, the atlas contains structure predictions for 756 million proteins in sequence clusters that are not covered by AlphaFold Database or the original ESM Atlas (Figure 5C).

The ESMC latent space is used to identify structural and functional similarities across the atlas. SAE features are computed for all proteins, and are used to create a single vector representation for each protein by taking the maximum activation across the sequence dimension (Appendix A.4.5.1). The 6.8 billion proteins in the atlas are clustered using these vector representations, where members of a cluster have Jaccard similarity of at least 0.6 to the cluster center (Appendix A.5.4). Above this threshold, proteins tend to share function—the median similarity between pairs of domains annotated with the same family by Pfam (81) is greater than 0.6 for over 88% of families, and 50% exceed this threshold for all pairs (Appendix A.4.5.2). The SAE clustering produces 230 million clusters with at least five members, and 7.7 million clusters with at least fifty members.

Distance in the sparse feature space reflects structural and functional similarity. SAE similarity achieves higher precision for the retrieval of proteins with shared topology and enzyme class than search using sequence or structure alone, particularly when sequence similarity is below 40% (Appendix A.4.5.3). SAEs also exhibit higher performance than sequence and structure based methods for detecting remote functional orthologs (Appendix A.4.5.4) and assigning pathway annotations to recently annotated human mitochondrial proteins (Appendix A.4.5.5).

Figure 5D visualizes the SAE feature space of the atlas for 7.7 million representative sequences (of 818 million unique proteins) from clusters with more than 50 members by projecting them into two dimensions using UMAP. Figure 5D (center inset) shows clusters that belong to protein families containing a structural motif characterized by a mixed α*/*β topology with a central five-stranded β-sheet flanked by α-helices, found in diverse nucleic acid-processing enzymes including ribonuclease H (RNase H), retroviral integrases, and transposases. Proteins are broadly organized according to the class of nucleases to which they belong, and in local regions of the projection closely related families such as Cas12 and its evolutionary ancestor TnpB (82) are located together.

ESMC groups this set of proteins due to shared nuclease-related features across the entire feature space, but the activation of specific features also separates proteins along established functional lines due to mechanistic differences. For example, feature 4804 activates on acidic residues in catalytically important sites across families, including the active sites of Cas-family nucleases which rely on a DDE motif for catalysis. Variants which have lost the DDE motif, and therefore the nuclease function, do not activate this feature. In contrast to defense-associated Type V CRISPR systems, the Type V-k CRISPR-associated transposase complex includes a Cas12k subunit that has lost its catalytic function and serves solely to direct DNA location while its TnsB subunit performs the transposition chemistry (83). Both catalytically active proteins, Cas12f and TnsB, activate F4804 while the inactive Cas12k does not.

Many clusters contain proteins from highly diverged branches of the tree of life, brought together by similarity in the ESMC latent space. One such cluster contains a number of RNA-guided DNA endonucleases including orthologues of eukaryotic Fanzor (84) and prokaryotic TnpB, the evolutionary ancestor of Cas12 (85). This cluster contains nine yeast Fanzors identified by Jiang et al. (86) that cluster with TnpBs, reflecting their known shared function despite low sequence homology. Comparison identifies features that are specific to Cas12/TnpB/Fanzor proteins relative to other RNase H–like proteins (F14197, which activates on the catalytic residues), as well as features that are specific to either Cas12a (F10161, which activates on the extended RuvC domain) or Cas12f/TnpB/Fanzor (F920, which activates on the ZF domain). Separation of Cas12a from Cas12f and TnpB reflects differences between them: Cas12a is a full-length CRISPR-associated nuclease with different domain structure from the highly compact Cas12f and TnpB systems.

Other clusters contain protein sequences that are not annotated by any Pfam hidden Markov model. For example, one 477-member cluster in the TnpB region has 461 members with no Pfam coverage (97%). The cluster centroid has 0.87 Jaccard similarity in SAE feature space and a TM-score of 0.59 to the canonical ISDra2 TnpB, despite a sequence identity of only 13.9%. This cluster contains a small number of proteins with Pfam coverage, but there are 315 additional clusters that only contain unannotated proteins but have high SAE feature similarity (Jaccard *>* 0.6) to clusters annotated with either Cas12 or TnpB domains. The closest five clusters to the Fanzor characterized by Saito et al. (84) contain 249 unannotated proteins with sequence identity as low as 26.7% to any known Fanzor (Appendix A.5.5).

Themes across protein biology can be identified and mapped through the atlas (Figure 5E). Of the 7.7 million clusters with more than 50 members, more than five million contain at least one high-confidence predicted structure. Half a million clusters contain proteins whose lowest common ancestor, taxonomically, is at the root of all cellular life. There are 20 thousand clusters in which the majority of proteins come from halophilic environments. Over 80 thousand clusters have high sequence diversity. (A mean pairwise sequence identity of less than 30%, of which 87.5% share a conserved structural fold.) Many can be associated with known proteins by remote homology, while others are clustered together for reasons that remain to be explored. A large fraction of clusters in the atlas are not well understood, with over two million containing no members with a Pfam annotation (excluding domains of unknown function). Features associated with proteins involved in DNA repair are found in over half a million clusters representing 60 million proteins. The atlas provides the foundation for analysis of functional and evolutionary patterns at massive scale.

Clusters span varying levels of taxonomic specificity in the tree of life. Some clusters represent protein functions that are conserved across life while others represent more specialized functions found only in specific lineages (Figure 5F). Of the clusters which contain proteins with taxonomy information (76%), the largest fraction of clusters (over 20%) have a lowest common ancestor at the domain level, reflecting broadly conserved protein functions. A further 19% span multiple domains, including the Fanzor and TnpB example shown above, making connections between proteins which are evolutionarily distant but associated by the sparse feature space.

The differential enrichment of features in specific biomes sampled by metagenomic sequencing reflect functional differentiation according to the unique needs of each environment (Figure 5G). For example, the human gut microbiome is enriched for gram-positive bacterial secretion machinery, soil is enriched for actinobacteria and interaction/signaling motifs, and marine biomes are enriched for photosystems and lumenal domains that are common in algae and cyanobacteria. Extreme biomes provide a unique insight on functional adaptation, and we see this reflected as well in the feature space—in halophilic environments, there is significant enrichment for haloarchaeal signatures and acidic regions which are necessary to maintain stability in high-salt conditions.

Some features are specific to high-level biological programs that are used across organisms and can be used to map the prevalence of proteins involved in particular processes across the atlas. Analyzing proteins with Gene Ontology (87) annotations in Swissprot, we identify 4,113 features spanning 61 biological processes for which high activation identifies a process with precision of greater than 0.5 (Appendix A.5.6). Even at a very high precision of 0.9 we find features specific to 49 unique processes. For each of these processes, tens to hundreds of millions of proteins belong to clusters whose representative activates a process-specific feature above its precision-tuned threshold (Figure 5H). The largest categories represent fundamental processes of life, including those associated with transcription, translation, cellular signaling, and metabolism.

The feature space maps functional themes across biology without requiring proteins to map directly to human-curated annotation systems. Millions of proteins that activate high-precision features for a biological process have no existing families annotated by Pfam (up to 70% of retrieved clusters). The features identify shared mechanisms or structural components that sequence similarity is unable to discover. Figure 5I shows three such examples where uncharacterized metagenomic proteins activate features associated with carbohydrate metabolism (F2383: polysaccharide-binding cleft; F10617: carbohydrate-binding groove), DNA recombination (F1442: tyrosine recombinase active site; F12645: recombinase/integrase DNA-binding module), and glycoprotein biosynthesis (F13471: PAPS-dependent sulfation domains; F11103: phosphate-binding motif). Comparing the predicted structures to the nearest structures in the PDB shows high overall cosine similarity in the SAE feature space and similar activation patterns of the key process-related features. This is corroborated by high structural similarity as measured by TM-score, despite very low sequence identity of 10–22%.

The atlas maps sequence, structure, and function across billions of proteins from all biomes and domains of life. Proteins are organized according to their similarity in ESMC’s representation space, enabling connections to be made between proteins which have significantly diverged or independently arrived at shared characteristics through different evolutionary trajectories, and providing a mechanism for interpreting these connections through biologically relevant features.

## Conclusion

The representations learned through language modeling of protein sequences provide a basis for state-of-the-art modeling of protein sequence, structure, and function. They enable prediction of protein structure and interactions at the atomic level, and can be inverted to design proteins and binding interfaces. Their organization captures the fundamental properties of proteins, recovering the reductionist understanding developed by more than a century of empirical science, and identifies structural, functional, and evolutionary relationships among billions of proteins.

There is a sense in which the language modeling task could not be more simple: given a protein sequence, predict the amino acids that have been masked from it. But there is another sense in which this task contains all the complexity of biology. The amino acid that evolution selects at each position in a protein is shaped and constrained by the many and varied roles that proteins play across life, and by the more basic structural, chemical, and physical properties that give rise to function. The distributional structure in the patterns of amino acids across evolution as a whole is thus a reflection of the deep underlying biology of proteins. By learning to predict amino acids across billions of sequences, protein language models internalize the variables and constraints that shape evolution.

From this standpoint, language modeling materializes a world model of protein biology within the model’s representation space. This world model supports the prediction of the structure of proteins directly from the representations, and the design of new proteins by searching over inputs that satisfy specific constraints on these predictions. It is general and comprehensive, representing the biology of proteins developed through natural evolution and generalizing beyond it. Together, these capabilities suggest the model has developed a representation space that is true to biology.

Accurate representations of proteins open up scientific questions to investigation at scales far beyond the physical limitations of laboratory science. The structures of 1.1 billion proteins were computed in approximately two weeks and can be used to answer questions about the organization of protein biology as a whole. Hundreds of thousands of antibodies can be screened in a few hours of computational time to discover ones that bind therapeutic targets in the laboratory.

## Acknowledgements

We thank Bridget Carragher, Aashish Manglik, David Agard, and Eshan Mehrotra for input on biophysical characterization; Eric S. Lander, Manuel Leonetti, Vamsi Mootha for discussions on biological interpretation; Shu-Hsien Sheu for scientific discussions and assistance with fluorescence microscopy; Mohammed AlQuraishi for discussions on structure prediction evaluations; Donghui Li for review of the manuscript; Alexander Sorkin (University of Pittsburgh), Mien-Chie Hung (China Medical University Hospital), and Hirofumi Nakatomi (Riken Center for Brain Science) provided plasmids for EGFR, PD-L1, and PDGFRβ, respectively; Finally we thank the entire Biohub team, particularly the contributors to the platforms, atlas web application, release of models, documentation, and other materials integral to disseminating the science of this paper.

## Appendix A: Materials and Methods

### A.1. ESM Cambrian Models

#### A.1.1 Architecture

The ESM Cambrian (ESMC) protein language models are based on a Transformer architecture adapted for protein sequences. We train three model variants at different scales: ESMC 300M, ESMC 600M, and ESMC 6B, with approximately 300 million, 600 million, and 6 billion parameters, respectively.

Each model consists of a token embedding layer, a stack of transformer blocks, and a regression output head. Input amino acid sequences are tokenized to a one hot embedding, which includes the 20 standard amino acids as well as special tokens. The token embedding maps each input token to a dense vector of dimension *d*_model_.

Each Transformer block follows a pre-norm architecture and consists of two sub-layers: a multi-head self-attention (MHA) module and a feed-forward network (FFN), each with a residual connection. Layer normalization is applied before each sub-layer. The attention module uses Rotary Position Embeddings (RoPE) (88) to encode positional information, enabling the model to generalize to varying sequence lengths. We apply layer normalization to the query and key projections prior to computing attention scores, which stabilizes training at scale. All attention heads use a fixed head dimension of 64 across all model sizes. The feed-forward network uses the SwiGLU activation function. All linear layers throughout the model are bias-free.

To improve training stability at depth, we apply a residue scaling factor of 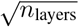 to the outputs of both the attention and feed-forward sub-layers before adding them to the residual stream. A final layer normalization is applied after the last Transformer block.

The output head is a two-layer MLP with a GELU activation and intermediate layer normalization, projecting from *d*_model_ back to the vocabulary size to produce per-position token logits.

We release three model variants: ESMC 300M, ESMC 600M, and ESMC 6B. Table S1 summarizes the architectural hyperparameters for each model variant.

### A.1.2. Pretraining Data

The ESMC models are trained on protein sequences drawn from three large-scale databases: UniRef (75), MGnify (30), and the Joint Genome Institute (JGI) Metagenome database (29). Together, these sources span both curated reference proteomes and metagenomic sequences sampled broadly across the tree of life. To reduce redundancy, all sequences are clustered at 70% sequence identity, yielding approximately 83M clusters from UniRef, 372M clusters from MGnify, and 2B clusters from JGI.

Training proceeds in two stages with distinct data mixtures and context lengths (Table S2). In Stage 1, the model is trained for 1M steps with a context length of 512 tokens. The data composition in this stage is weighted toward metagenomic sequences, with JGI and MGnify together comprising approximately 65% of the training data and UniRef accounting for 36%. In Stage 2, training continues for an additional 500K steps with an extended context length of 2048 tokens. The data mixture is shifted to emphasize UniRef sequences, which constitute approximately 63% of the training data, while JGI and MGnify are reduced to 31% and 6%, respectively. This two-stage strategy allows the model to first learn broad sequence statistics from the large and diverse metagenomic corpus at shorter context lengths, before refining its representations on higher-quality curated sequences at longer context lengths.

#### A.1.3. Training Details

All ESMC models are trained with a masked language modeling (MLM) objective (25). At each training step, 15% of input tokens are randomly selected and replaced with a special [MASK] token. The model is trained to predict the original identity of the masked positions. The training loss is a per-sequence weighted cross-entropy, ensuring that each sequence contributes equally to the gradient regardless of its length or number of masked positions.

We optimize all models using AdamW with *β*_1_ = 0.9, *β*_2_ = 0.95, and *ϵ* = 10^−8^. Weight decay is applied selectively, excluding bias terms and layer normalization parameters. Gradient norms are clipped to a maximum of 1.0. The 300M and 600M models are trained in bfloat16 mixed precision; the 6B model is trained in FP8 mixed precision using TransformerEngine.

Learning rates are scaled across model sizes using Maximal Update Parameterization (*µ*P). We calibrate a base learning rate and weight decay on a small proxy model (*d*_model_ = 512, *n*_layers_ = 16) and transfer these hyperparameters to larger models by scaling the learning rate inversely with width and with the square root of depth, while adjusting weight decay to maintain a constant learning rate–weight decay product. This allows us to train all model sizes without per-model hyperparameter tuning.

We use a Warmup-Stable-Decay (WSD) learning rate schedule. The learning rate increases linearly over a 1,000-step warmup, remains constant at the peak through Stage 1, and decays over all of Stage 2 (500,000 steps, ≈1/3 of the 1,500,000-step total) to a minimum value of 0.1× the peak learning rate.

As described in A.1.2, training proceeds in two stages. In Stage 1, models are trained for 1M steps with a context length of 512 tokens. In Stage 2, training continues for an additional 500K steps with an extended context length of 2048 tokens. All model sizes use approximately 4.2M tokens per batch across both stages: Stage 1 uses batch size 8,192 at sequence length 512, and Stage 2 uses batch size 2,048 at sequence length 2,048. Table S3 summarizes the key training hyperparameters.

**Table S1.** Architectural hyperparameters for the ESMC model family.

| Hyperparameter | ESMC 300M | ESMC 600M | ESMC 6B |
| --- | --- | --- | --- |
| Layers ($n_{\text{layers}}$) | 30 | 36 | 80 |
| Model dimension ($d_{\text{model}}$) | 960 | 1,152 | 2,560 |
| Attention heads ($n_{\text{heads}}$) | 15 | 18 | 40 |
| Head dimension ($d_{\text{head}}$) | 64 | 64 | 64 |
| FFN hidden dimension | 2,560 | 3,072 | 6,912 |
| Vocabulary size | 64 | 64 | 64 |

**Table S2.** Pretraining data sources and stage-dependent sampling proportions.

| Source | Clusters | Stage 1 | Stage 2 |
| --- | --- | --- | --- |
| UniRef | 83M | 36% | 63% |
| MGNify | 372M | 11% | 6% |
| JGI | 2B | 54% | 31% |

**Table S3.**
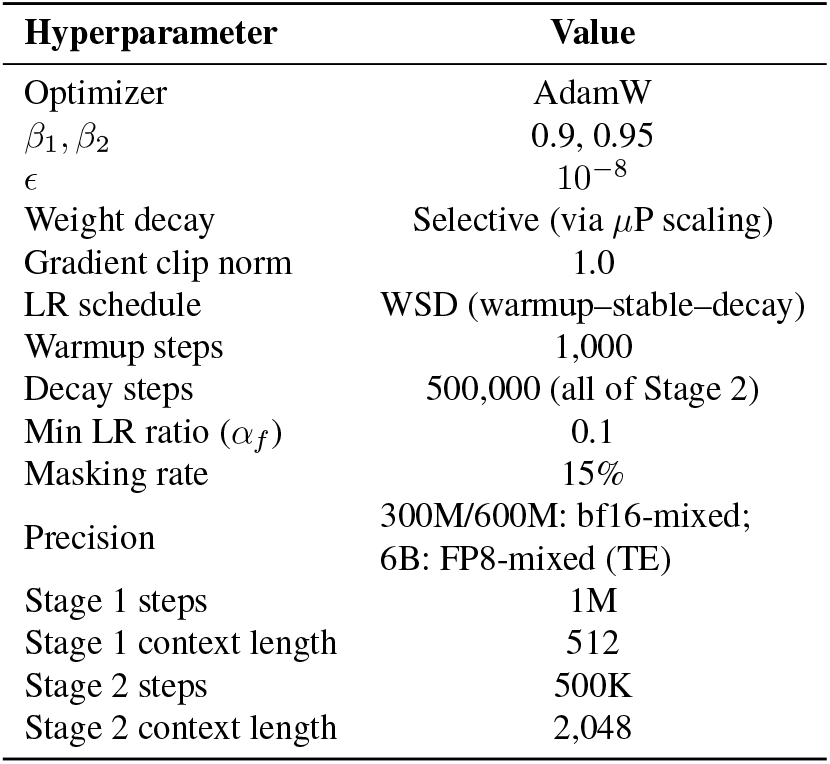
Training hyperparameters for the ESMC model family.

| Hyperparameter | Value |
| --- | --- |
| Optimizer | AdamW |
| $\beta_1, \beta_2$ | 0.9, 0.95 |
| $\epsilon$ | $10^{-8}$ |
| Weight decay | Selective (via $\mu\text{P}$ scaling) |
| Gradient clip norm | 1.0 |
| LR schedule | WSD (warmup–stable–decay) |
| Warmup steps | 1,000 |
| Decay steps | 500,000 (all of Stage 2) |
| Min LR ratio ($\alpha_f$) | 0.1 |
| Masking rate | 15% |
| Precision | 300M/600M: bf16-mixed;<br>6B: FP8-mixed (TE) |
| Stage 1 steps | 1M |
| Stage 1 context length | 512 |
| Stage 2 steps | 500K |
| Stage 2 context length | 2,048 |

#### A.1.4. Evaluations

We evaluate the quality of ESMC representations using unsupervised contact prediction, a standard probe of whether protein language model attention maps encode structural information without explicit supervision.

##### A.1.4.1 Unsupervised Contact Prediction

Following Rao et al. (2), we assess whether the self-attention maps of ESMC capture residue–residue contacts. For each model, attention maps are extracted from all layers and a logistic regression is trained on the symmetrized attention weights to predict binary C*β* contacts (distance *<* 8 Å) from a small set of 20 training structures. Model performance is evaluated on single chain structures from the PDB (downloaded `2024-02-28`), clustered at 40% sequence identity (`mmseqs linclust --min-seq-id 0.4 -c 0.8 --cov-mode 1 --cluster-mode 2 --kmer-per-seq 100`). For each protein of length *L*, the precision of the *L* most confident predictions (P@L) for all residue pairs with sequence separation ≥ 24 is reported. Structures with fewer than *L* true contacts are filtered from the evaluation set. For evaluation speed, sequences with length *>* 510 are truncated to the first 510 residues, this affects 3,425 chains. After filtering, a total of 20,775 chains are evaluated on.

###### Scaling experiments

A series of five masked language models are trained at increasing scale. Each model shares the same architecture described in A.1.1. Model sizes range from 50 million parameters (depth 16, width 512) to 1.36 billion parameters (depth 48, width 1536) (see Table S4).

**Table S4.** MLM Architecture and hyperparameters. Scaling sweep (top) and ESMC models (bottom)

| Model | Depth | Width | Parameters | Steps |
| --- | --- | --- | --- | --- |
| 50M | 16 | 512 | 54.9 M | 80,000 |
| 170M | 24 | 768 | 170.7 M | 80,000 |
| 400M | 32 | 1024 | 412.4 M | 80,000 |
| 790M | 40 | 1280 | 814.8 M | 80,000 |
| 1.4B | 48 | 1536 | 1.36 B | 80,000 |
| 300M | 30 | 960 | 333.0 M | 1,000,000 |
| 600M | 36 | 1152 | 575.0 M | 1,000,000 |
| 6B | 80 | 2560 | 6.35 B | 1,000,000 |

Models are trained for 80,000 optimizer steps with a batch size of 2^22^ tokens at context length 512. Total training FLOPs for each configuration are computed as

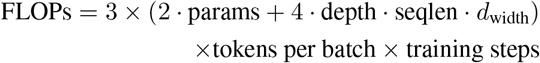

A line in Figure 1B is fit by least-squares to the running-max Pareto frontier of (log_10_(FLOPs), P@L-LR) across all checkpoints from the five scaling experiments, then extrapolated to ESMC 300M, 600M, and 6B. The ordinary-least-squares fit gives P@L-LR = 0.115 · log_10_ (FLOPs) − 1.982, *R*^2^ = 0.99. Evaluating the fit at the FLOPs of every ESMC checkpoint on the ESMC running-max Pareto frontier checkpoints from 300M, 600M, and 6B training trajectories recovers the empirical precision with *R*^2^ = 0.98 and RMSE = 0.013 P@L-LR.

###### Comparison of ESMC with ESM2

Figure 1C shows the performance of ESMC models compared with the prior ESM2 models. ESMC 300M (P@L-LR = 0.552 ± 0.002) achieves contact prediction performance comparable to ESM2-650M (0.566±0.002) despite roughly half the parameters. ESMC 600M (0.589 ± 0.002) is statistically indistinguishable from ESM2-3B (0.587 ± 0.002) and approaches ESM2-15B (0.593 ± 0.002). ESMC 6B (0.725 ± 0.002) outperforms the largest ESM2 models by 0.132 ± 0.002 P@L-LR. All ranges are 95% confidence intervals (paired bootstrap, *n*_boot_ = 5000).

##### A.1.4.2 Layerwise Evaluations

The extent to which information about function appears in each layer is assessed using an unsupervised clustering approach to classify Enzyme Commission (EC) Numbers. To evaluate layerwise embeddings, a dataset was first constructed from SwissProt and The Encylopedia of Domains (TED) (89), including proteins with unambiguous fourth-level (x.x.x.x) EC and CATH annotations. The full procedure for dataset construction is described in A.1.4.3. This results in a dataset of 10,371 proteins (8493 unique sequences) with sequence length *<* 1024, representing 57 EC classes. The 2560-dimensional hidden states at each layer of ESMC 6B are then mean-pooled across the sequence dimension to construct a layer-specific embedding. For each layer, the embeddings are projected down to 10 dimensions using Principal Components Analysis (PCA) to keep the most relevant dimensions for clustering. A k-nearest neighbors (kNN) classifier with k=100 is then trained on all of the labeled data, and the same dataset is re-annotated with the predicted EC number via majority vote. Figure 1D reports the accuracy of the classifier at each layer.

##### A.1.4.3 EC-CATH Evaluations

Dataset construction. To identify whether the model’s latent space contains representations of function that are independent of structure, we construct a benchmark dataset for enzyme function prediction containing only enzyme classes which span multiple distinct structural topologies. For enzyme class, we use the Enzyme Commission (EC) number classification system (31), and for structures we use the CATH classification system (32), both of which have a four-level hierarchical numbering scheme (e.g. x.x.x.x). This EC-CATH dataset focuses on the fourth (most specific) level of the EC hierarchy and the third level of the CATH hierarchy to ensure that the structures across splits differ in overall topology.

The dataset was constructed by integrating curated functional annotations, structure predictions, and domain classifications from three primary sources: SwissProt for enzyme function (as of December 6, 2024), AlphaFold DB (AFDB) v4 (90) for protein structures, and TED for domain topology annotations. Only proteins with length ≤ 512 residues and exactly one fully-annotated CATH domain and one fully-annotated EC number were used to remove confounding by multi-domain proteins or proteins with ambiguous labels (e.g. 3.5.-.-). The dataset in this form was used for the layerwise evaluations in A.1.4.2.

To construct the evaluation dataset, we sample positive and negative examples from each (EC, CATH) group. To ensure sufficient support in each (EC number, CATH topology) group, pairs with less than 6 unique proteins are removed. The proteins within a group serve as positives for predicting that EC number, and structurally controlled negatives are sampled by randomly drawing up to 1000 proteins from the TED dataset which have the same CATH topology but a different EC number. Only proteins annotated with at least one EC number are included, ensuring that the negatives are also enzymes and not simply missing EC number annotation. A separate task dataset is created for each structural holdout (in which one topology is held out at a time per EC number), resulting in 99 unique (EC, CATH) train-test splits across 42 unique EC numbers. Most EC numbers have only two unique topologies, but some have 3–5, e.g., 3.1.26.5, 3.1.3.48, 3.2.1.4, 3.1.3.16, 5.3.1.6.

Inspection revealed that some of tasks are simple to solve by sequence similarity due to closely related enzymes, high similarity between topologies, or misannotation, e.g., identical sequences annotated with different CATH topologies in the TED dataset. We manually removed 10 EC numbers that have greater than 50% sequence identity between any two proteins across topologies: 1.11.1.7, 2.3.1.16, 2.5.1.141, 2.7.7.6, 3.1.11.6, 7.1.1.2, 7.1.1.9, 2.7.7.7, 3.4.19.12, 7.2.1.4. The final dataset contains 5,829 unique proteins as positive examples and 9,211 unique structure-matched negatives, distributed across 73 leave-one-CATH-out train-test splits across 32 EC numbers and 42 CATH topologies.

Figure S1 shows a subset of this data where every EC appears on ≥ 2 topologies and every CATH topology has ≥ 2 EC numbers organized according to the latent space of ESMC (cosine distance on the mean-pooled embeddings from layer 60, close to the peak in Figure 1D). Structural topology appears to be a primary organizing principle in the latent space, but within each topology the local geometry shows clear organization by EC number.

**Figure S1.**
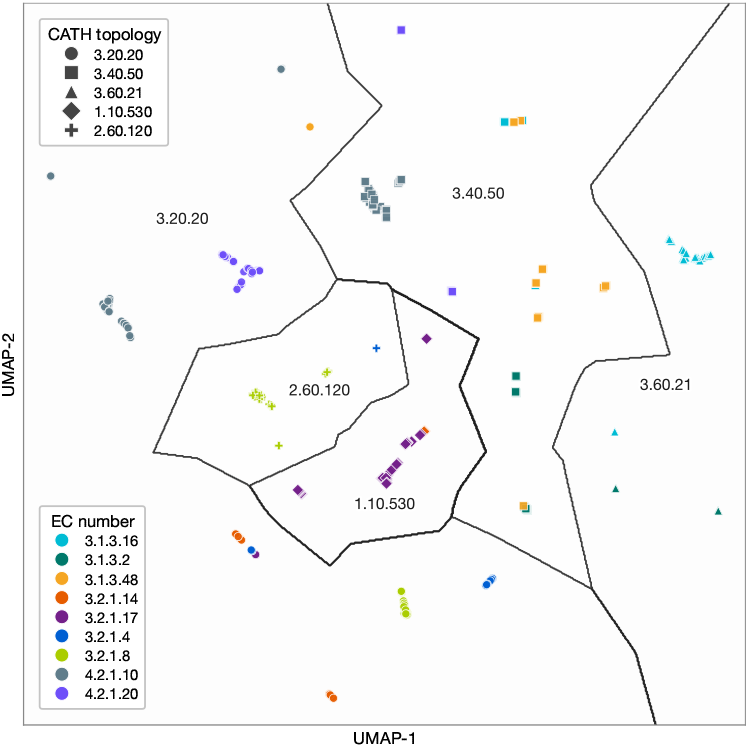
Subset of EC-CATH dataset in the ESMC representation space. UMAP projection of a subset of the EC-CATH benchmark in the ESMC layer-60 latent space. Each point is one protein, colored by EC number (function) and shaped by CATH topology (structure). Voronoi cells partition the UMAP by nearest CATH topology.

###### Unsupervised evaluation

We use principal component analysis (PCA) to additionally probe how enzyme class and structural topology separate in the latent space of the ESMC 6B. The four combinations shown in Figure S2 are derived from the subset of the data where all combinations of at least two EC numbers and at least two CATH classes are present, redundancy-reduced within each class by clustering at 50% sequence identity. The mean-pooled hidden states from every layer are concatenated into one large embedding for each sequence, thus mapping the latent space of the entire network. Figure S2 shows that there are high-variance principal components that encode both EC and CATH, often in orthogonal subspaces. Performing the same analysis with a model of the same architecture with randomly initialized weights shows that this organization emerges during training.

To quantify the functional organization of the latent space of ESMC in a structure-controlled manner, we measured the cosine similarity between proteins of the same EC class across the topology pairs in the dataset and compared to the similarity between proteins of different EC classes. Concretely, for each EC number in the EC-CATH dataset, we consider all pairs of proteins that differ in CATH topology. Some of these pairs have the same EC number (the “positives” used to construct the dataset) and some have different EC numbers (the “negatives”). The cosine similarity between representations of each pair of proteins that share the same EC number across these topology pairs is z-score normalized against the background distribution defined by pairs with different EC numbers but the same pair of topologies, to control for structural variation as much as possible.

Figure S3A shows these normalized similarity distributions for four example EC numbers from the EC-CATH dataset using representations from each of the ESMC model scales. The distributions are computed from the layer that achieves the best mean z-score within each EC number, i.e., the layer that best separates EC number across CATH topologies. These distributions show that proteins with the same enzymatic function cluster together more closely in the latent space than proteins with different enzymatic functions, and also that this separation is greatest in the 6B parameter model. Due to z-score normalization, this difference is not driven simply by differences in the embedding dimension.

The same procedure is performed using intermediate models throughout training to explore how information emerges during the training process. Figure S3B shows the mean cosine similarity z-score for positive pairs across all EC numbers. Separation by EC class emerges early in training, in the first 250k steps, before rapidly plateauing in the smaller models (300M and 600M parameters). The 6B-parameter model continues to improve as training continues, with little evidence of diminishing returns up to 1.5M steps. As shown in Figure S3C, this improvement is largely driven by improved sequence understanding (pseudoperplexity). Below an average pseudoperplexity of about 3.0 (computed across this dataset, the two smaller ESMC models do not achieve pseudoperplexities below 3), we see a step change increase in the degree to which functions share directions in the latent space.

###### Supervised evaluation

Linear probes fit on the representations are one way to identify directions in the latent space that are predictive of enzyme class and generalize across structural topologies. We train a logistic regression model for each of the 73 EC prediction tasks defined in the ECCATH dataset, holding out an entire topology as a validation set. Ridge regression models with an L2 penalty ranging from 10^−3^ to 10^20^ were fit for each task and mean-pooled representations from each layer of the model, with classes weighted according to their relative proportion in the training dataset. The regularization parameter for each model is chosen based on the Matthews Correlation Coefficient (MCC) on the held-out CATH topology. We use MCC due to the wide range of class imbalances in the test data, which includes tasks with both high fraction of positives and high fraction of negatives. For added robustness on small and noisy datasets, we fit five models with different random seeds for each task and choose the best-performing model on the held-out data. For final evaluation, we select the best-performing layer for each task and report MCC on the held-out topology. This procedure is designed primarily to discover directions in the latent space that generalize across CATH topologies, not to train a generally optimal EC prediction model.

**Figure S2.**
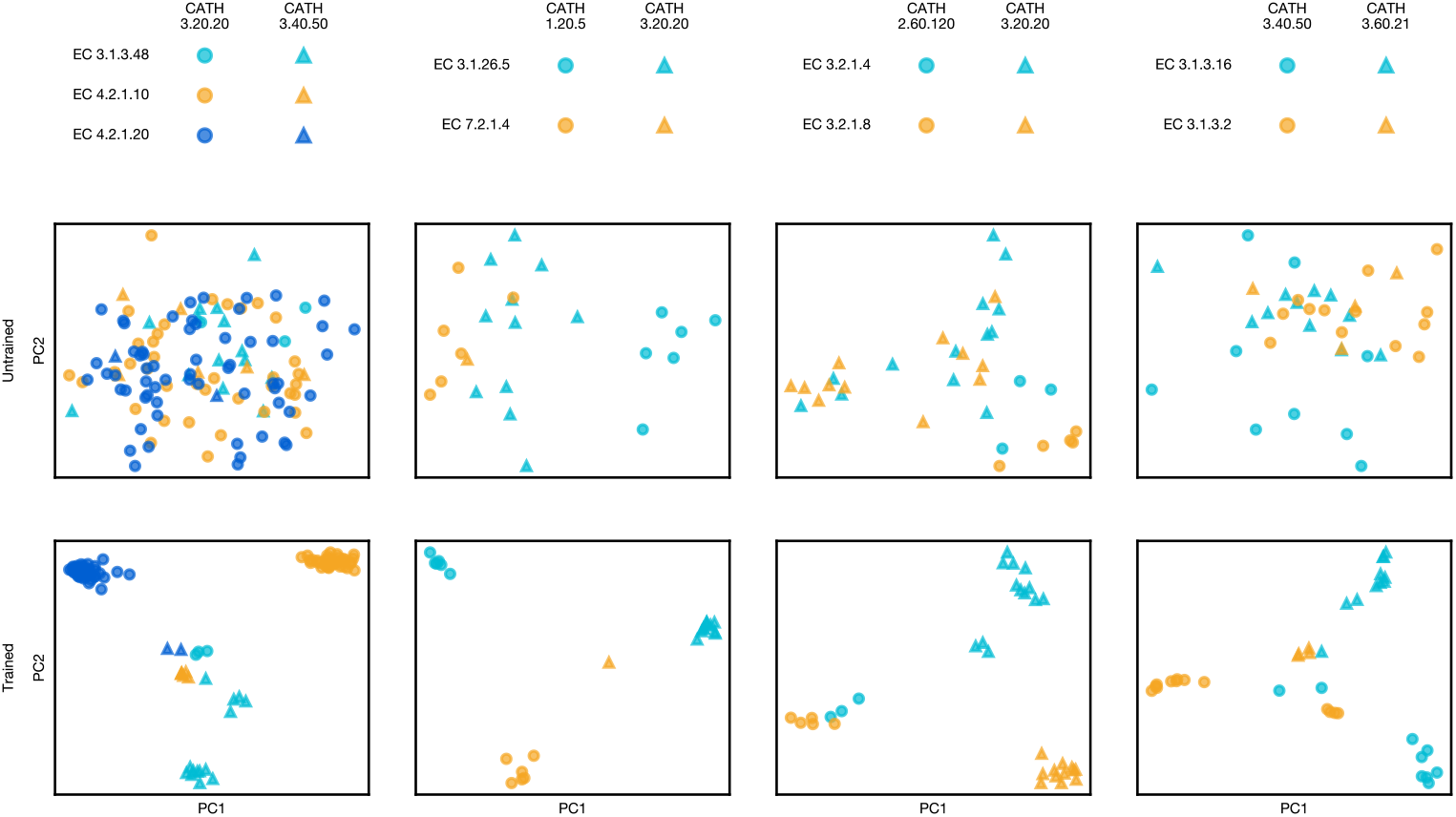
Principal components analysis on the EC-CATH dataset. Principal components analysis of concatenated layer representations of ESMC 6B before and after training. A separate PCA is fit to each of the four subsets shown from left to right, with EC numbers shown as colors and CATH topologies shown as shapes.

Figure S4A shows that the performance of all prediction tasks increases as the models scale. Across all model sizes, trained ESMC representations outperform untrained baselines, reinforcing that unsupervised training causes these function-specific directions to emerge. Comparison to untrained baselines suggests that the returns to scale arise from improved organization of concepts concordant with biology in the representations, rather than from increased capacity alone.

When the models are ranked by performance on each task, they sort by model scale, with the 6B model achieving the best performance on 53 / 73 tasks (72.6%) Figure S4B. No untrained baseline achieves higher performance than any trained model on any task. The improvement in performance with scale holds across all thresholds of MCC, with the 6B-parameter model achieving the best task-level success rate at each threshold Figure S4C.

##### A.1.4.4 Stability Prediction Evaluations

We investigated whether the representation space contains information on basic biophysical properties of proteins by predicting the thermodynamic stability of protein folding.

###### Methods

We evaluate on the Megascale (91) dataset, which comprises 776,000 data points corresponding to 331 natural and 148 *de novo* designed protein domains 40 to 72 amino acids in length. For each domain, the dataset includes all single mutants, and in some cases selected double mutants, around the wild-type sequence. Each mutant is labeled with its thermodynamic folding stability Δ*G* as measured by a high throughput experimental assay.

A meaningful and challenging data split was constructed based on structural similarity. First, following the split methodology from ProteinDPO (92), all predicted domain structures were clustered with Foldseek “easy-cluster” (93) with a 0.5 coverage threshold (with the flag `-c 0.5`). No structural clusters were permitted to span across train, validation, and test. Non-singleton clusters, i.e., structural clusters with multiple families, were placed in the training dataset, keeping a single family as a representative. Singleton structural clusters were placed into validation and test sets.

**Figure S3.**
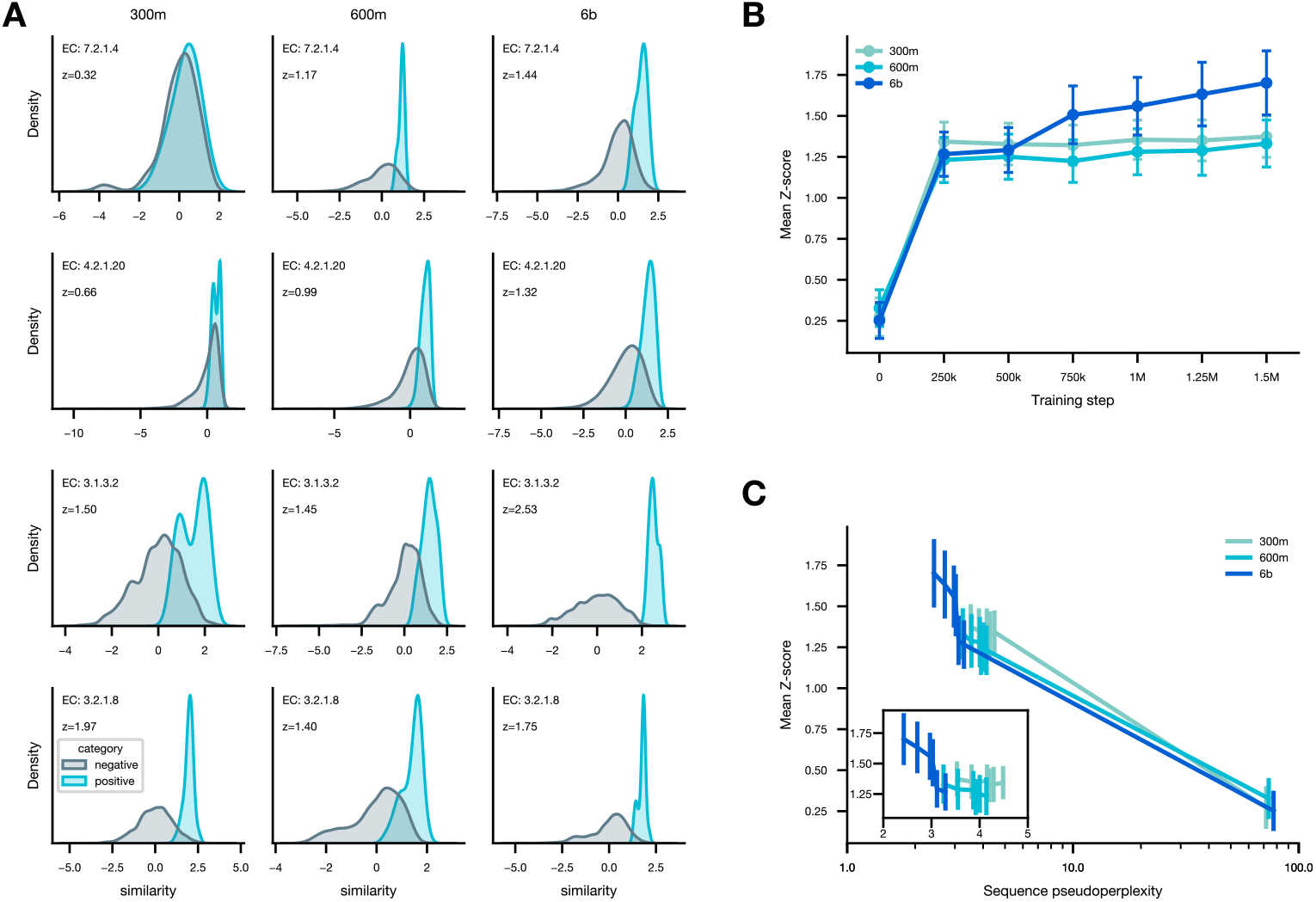
Unsupervised analysis of EC-CATH dataset. **(A)** Normalized cosine similarity distributions for four sampled EC numbers in the dataset across contrastive pairs of CATH topologies. **(B)** Normalized mean z-score for matching EC numbers over the course of training, as a function of training step. Error bars are standard error computed over the 32 unique EC numbers in the dataset. **(C)** Normalized mean z-score for matching EC numbers over the course of training, as a function of sequence pseudoperplexity on the proteins in the EC-CATH dataset. The inset shows the same data with the untrained model (step 0) removed for clarity. Error bars are standard error computed over the 32 unique EC numbers in the dataset.

**Figure S4.**
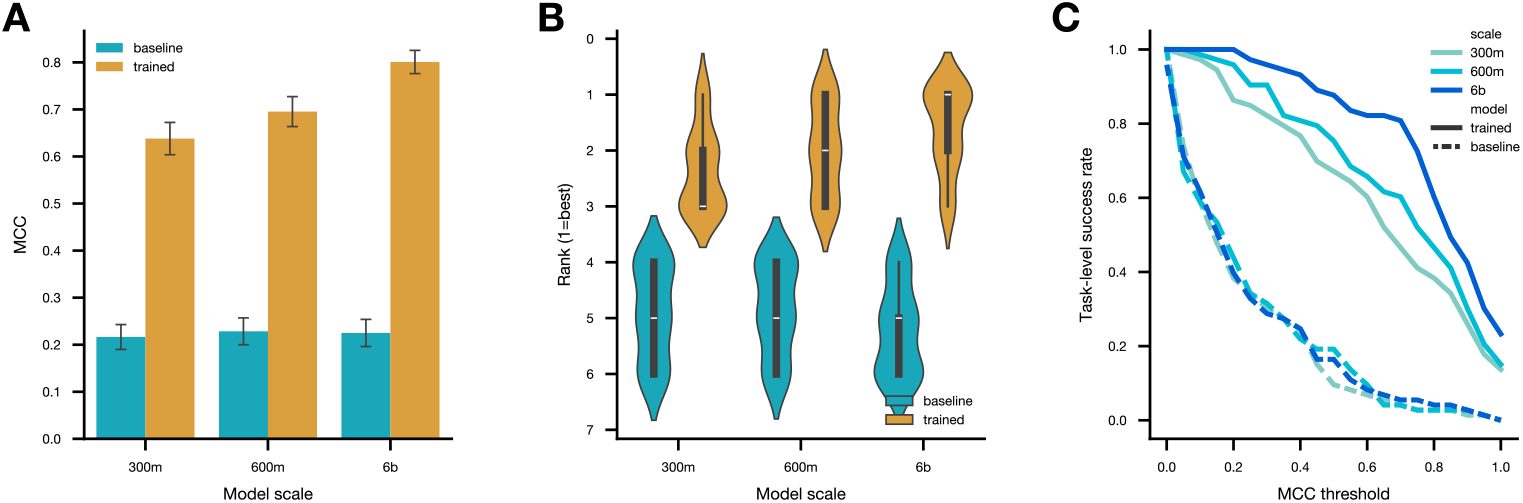
Supervised evaluation on the EC-CATH dataset. **(A)** EC prediction performance across all tasks in terms of Matthews correlation coefficient (MCC), comparing ESMC models at each scale before and after pretraining. We show the average over the best-performing layer for each task, with error bars representing the standard error over tasks. **(B)** Task win rate (proportion of tasks where model achieves best performance) over a range of training data subsamples from 10% to 100%. Reported results are for the best performance achieved over five subsamples for each model. **(C)** Task-level success rate, defined by MCC above threshold, across the full performance distribution.

As observed in other protein prediction applications such as mutation effect prediction (94) and protein structure prediction (95), we find that the performance of stability classifiers depends on the language model pseudoperplexity (pppl) of the wildtype sequence (Figure S5A). In order to remove perplexity confounding across splits, proteins were stratified into train, validation, and test sets by pseudoperplexity under the ESMC 6B model at three levels: low (pppl 0–2), medium (2–8) and high (8–20). This resulted in 47 families in train, 13 in validation, and 50 in test (Table S5).

**Table S5.** Train, validation, and test dataset statistics for stability prediction.

| pppl (ESMC 6B) | Train | Val | Test |
| --- | --- | --- | --- |
| Low [0–2] | 10 | 3 | 10 |
| Medium [2–8] | 32 | 8 | 32 |
| High [8–20] | 5 | 2 | 8 |
| Total | 47 | 13 | 50 |

To predict Δ*G*, a ridge-penalized linear regression from `sklearn` was trained on top of ESMC representations, mean-pooled across the sequence dimension. The ridge penalty was tuned on the validation set, searching between 10^−3^ and 10^12^. Comparing performance across layers, the second to last embedding from the ESMC transformer stack consistently performed best (Figure S5B), significantly outperforming the final layer (which is projected to final sequence logits).

###### The ESMC latent space contains directions that correspond to Δ*G*

To assess whether directions in the ESMC representations correspond to stability, a ridge regression model was trained on embeddings from the penultimate layer to predict Δ*G* on the Megascale training dataset.

To compare data efficiency between models, the training dataset was subsampled to subsets of the 47 training families: 1, 4, 23, and 47 families. While zero-shot performance is strong (0.56) for all ESMC models with respect to local stability within protein families (ΔΔ*G*; Figure S5C), predictions continue to improve with additional training data, reaching Spearman correlation of 0.8. There are mild but not significant returns from scale, with ESMC 6B slightly outperforming ESMC 300M and ESMC 600M on average.

The returns to model scale become more apparent when evaluating global stability (Δ*G*), computed across diverse protein families that are structurally distinct from the training dataset (Figure S5D). This implies that directions corresponding to stability can be learned from less data with larger models, and larger models see compounding benefits from increasing dataset size. Zero-shot prediction of global Δ*G* on 50 families fails to show predictive value. However, with training data, Δ*G* prediction performance improves steadily with the addition of training data from 1 to 47 training families (*x*-axis). In addition, Δ*G* performance improves as a function of ESMC model scale, with larger models showing improved performance at the same training data scale. The best model, ESMC 6B, achieves Spearman of 0.68 with all training data. While trained models continue to improve with training data, baselines models with randomly initialized weights decrease with additional training data beyond four families.

###### Learned stability directions generalize to new datasets

To assess the generalization of these stability probes, we evaluated them on an entirely held out experimental dataset: Fireprot (96), specifically the homolog-free version created in ThermoMPNN (97) which was filtered for sequences that are similar to the Megascale training data at a sequence identity cutoff of 25%.

Fireprot differs from the Megascale dataset in several ways. First, Megascale only has short proteins up to length 75, while Fireprot varies between 50–300. Second, Megascale uses a single high-throughput sequencing based screen to measure every variant, while Fireprot aggregates data across a variety of experiments (enriched for alanine scans). Third, Fireprot only includes natural proteins, while Megascale includes some de novo designed proteins.

To evaluate performance, we use stability directions fit on Megascale using the full 47 family training dataset. Fireprot sequences were then embedded using the same layer (second-to-last) and the dot product between the learned stability direction and the embedded Fireprot sequence was computed. Performance on ΔΔ*G* prediction was evaluated on a per-family basis. Figure S5E shows that the learned directions have excellent predictive capability on the held-out dataset. These learned directions are also significantly better predictors of stability than pseudoperplexity (Wilcoxon paired signed-rank test, *p* = 1.3 × 10^−5^), achieving an average Spearman close to 0.5 across 37 landscapes compared to ∼0.2 for zero-shot pseudoperplexity. Figure S5F shows that ESMC stability prediction performance is on par with structure-based predictors such as FoldX (98) and Rosetta (99), despite only requiring protein sequence as input.

### A.2. ESMFold2

#### A.2.1. Architecture

During training, ESMC learns the implicit determinants of protein structure from pretraining on sequences. ESM-Fold2 integrates frozen representations of ESMC to create an atomistic model of protein, nucleotide, and molecular interactions. The representations *x*_*i*_ from ESMC are converted to a 2D pair representation *z*_*ij*_ (for each pair *x*_*i*_, *x*_*j*_), and processed with triangle operations which pass information between representations *z*_*ij*_, *z*_*ik*_, and *z*_*jk*_ (22, 23).

**Figure S5.**
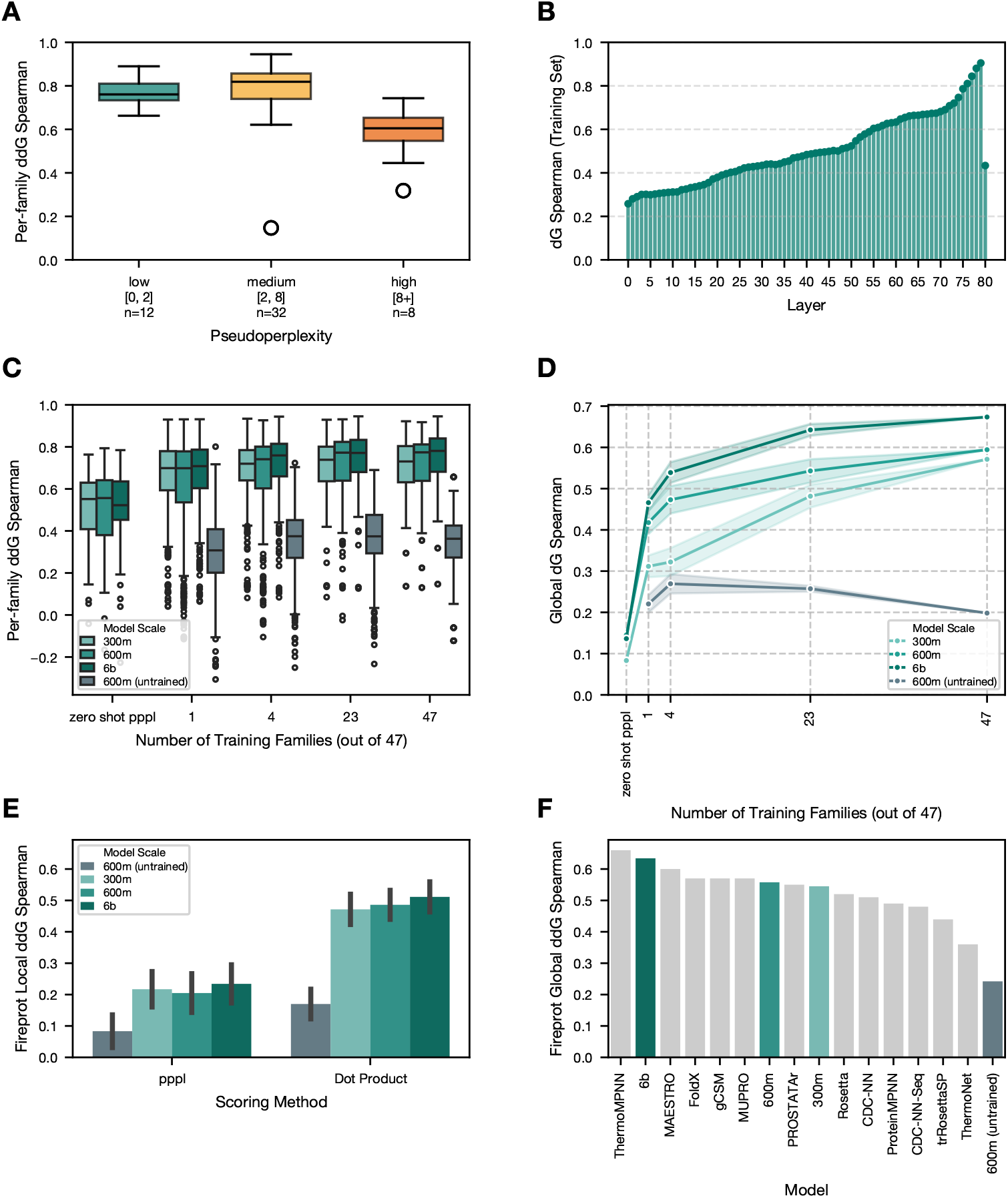
Stability prediction with ESMC representations. **(A)** Performance of stability (Δ*G*) prediction stratified by pseudoperplexity. Each point represents a single landscape in the test set, with the *y* axis showing Spearman correlation of the stability predictor for a given landscape. **(B)** The performance of each layer of ESMC 6B when used as input to a Ridge regression model to predict stability. **(C)** Spearman correlation of Ridge regression stability predictors with experimentally measured ΔΔ*G* (i.e. Δ*G*(variant) − Δ*G*(wildtype)) on 50 structurally held-out families, measured per-family. Each point represents the performance of a predictor on a single landscape. At each training dataset size, 3 separate bootstrap samples are taken and models are trained independently. **(D)** Spearman correlation of Ridge regression stability predictors with experimentally measured Δ*G* on 50 structurally held-out families. Error bars show 95% confidence intervals for 3 random subsets of the training data at the indicated dataset size. Three model scales of ESMC (300M, 600M, and 6B) as well as an untrained 600M baseline (grey) are shown. **(E)** Spearman correlation of zero-shot pseudoperplexity and stability directions on Fireprot dataset. Error bars are ±1 standard error across 37 landscapes with *>* 10 data points. **(F)** Performance on predicting global ΔΔ*G* with Fireprot data (*N* = 2,578). Methods in light grey require structure prediction, while ESMC-based predictors are sequence-only. ESMC predictors are trained on Δ*G* data, and are converted to ΔΔ*G* predictors by taking ΔΔ*G*_predicted_ = dot(direction, embedding_mutant_) − dot(direction, embedding_wt_).

Inference time compute can be scaled by passing the representations from the end of the pair folding layers back into the beginning. In ESMFold2, this update is constructed to be contractive, which prevents the activations in the residual stream from growing unbounded (40). This construction enables stable gradients through the recurrent folding loop.

At a high level, ESMFold2 processes inputs through four learned components: an input embedder that produces a token-level feature **f**_inputs_, recurrent pair only folding layers that produce the trunk pair representation **z**, a diffusion sampler that turns **z** and reference atom features into 3D coordinates, and a confidence head that scores the prediction. The full inference flow is given in Algorithm 1; each component is detailed in the following subsections.

- Input Encoding (A.2.3): Full, uncropped protein sequences {**x**_*c*_} for each protein chain in the input are passed independently to ESMC 6B and representations are extracted and projected to a pair state **z**_**l**m_ (A.2.2). Atom-level features **f**_atom_ (reference 3D positions **x**_ref_, element, charge, atom name) are encoded via sliding-window atom transformer and aggregated to token level, then concatenated with residue features **f**_seq_ and MSA features **f**_msa_ (profile and deletion mean, if MSAs are provided) to form **f**_inputs_ and **z**_**i**nit_; the token-bonds encoding **f**_bond_ is added to the pair stream. If MSAs are provided, the MSA encoder is used to process the MSAs and update **z**_**i**nit_.
- Recurrent Folding Layers (A.2.5): A series of pair folding layers that update the pair representation **z** over *T* recurrent loops. Each pair folding layer uses a streamlined architecture introduced by Ouyang-Zhang et al. (39), consisting of a triangle multiplication update and a feed forward network. After passing through all layers, representations are fed back into the initial layer. This recurrent update is stabilized by constraining each update to be contractive (40), which enables backpropagation through the recurrent update. At the end of the recurrence, two pair folding layers produce the final representation, from which distogram logits **z**_**d**isto_ are read.
- Diffusion Sampling (A.2.6): An atom-level diffusion module conditioned on the pair representation **z** and the initial single representation **f**_inputs_ produces predicted atomic coordinates **x**_pred_. Sliding window atom attention is used for atom-level updates. Inference uses a truncated noise schedule that skips the very-high-noise regime where signal-to-noise ratio is negligible (Algorithm 4).
- Confidence Head (A.2.7): Four pair folding layers which take as input the final pair representation, the predicted distogram, and the diffusion sample to predict per-token pLDDT **x**_plddt_, predicted distance error **z**_**p**de_, and predicted aligned error **z**_**p**ae_.

The simplified pair folding layer (which removes the single representation and triangle attention) substantially reduces the compute and memory requirements of ESMFold2. We also release an inference optimized ESMFold2-Fast variant, with 24 folding blocks instead of 48, and without MSA conditioning for single-sequence inference. Table S6 summarizes the architectural hyperparameters for the two models.

##### A.2.2. Language Model Integration

ESMFold2 uses a frozen ESMC 6B language model to provide rich per-token representations to the folding trunk. The language model parameters are never updated during structure prediction training.

###### Full-sequence encoding

ESMC processes full protein sequences. During training, the folding layers receive a cropped window of a complex, but the language model receives the full protein sequence (up to 2046 tokens). This means ESMC can provide contextual representations from residues outside the crop window, giving the model access to long-range sequence context that would otherwise be lost to cropping. When multiple protein chains are present, each chain **x**_*c*_ is encoded separately by ESMC, so that each chain is processed in-distribution for the pretrained model. This is particularly important for homomeric structures, where including both chains in the same context degrades representation quality due to copying.

###### Representation extraction

Protein sequences are tokenized and passed through the frozen ESMC model. Following Lin et al. (3), hidden states from all transformer layers, including the initial embedding layer, are extracted, producing a tensor of shape [*B, L, N*_layers_ + 1, *D*_lm_], where *B* is batch size, *L* is sequence length, *N*_layers_ is the number of transformer layers, and *D*_lm_ is the language model hidden dimension.

###### Projection to pair space

Each layer’s hidden states are independently projected via a shared LayerNorm followed by a linear projection from *D*_lm_ to *d*_*z*_ = 256. A learnable weight vector **w**_*z*_ of length *N*_layers_ + 1 is combined via softmax to produce per-token representations. These are then expanded to pair space via an outer-sum operation, followed by an MLP and LayerNorm, yielding a pair contribution **z**_**l**m_ that is added to the initial pair representation. Four pair folding layers refine the language model contributions before it is added into the residual stream.

**Table S6.** Architectural training hyperparameters for ESMFold2 model variants.

| Component | ESMFold2-Fast | ESMFold2 |
| --- | --- | --- |
| Pair folding layers | 24 | 48 |
| Pair dimensions | 256 | 256 |
| Diffusion layers | 12 | 12 |
| MSA encoder layers | - | 4 |
| MSA encoder dimension | - | 128 |
| LM encoder layers | 4 | 4 |
| Confidence head layers | 4 | 4 |

###### Algorithm 1

ESMFold2 Forward Pass.

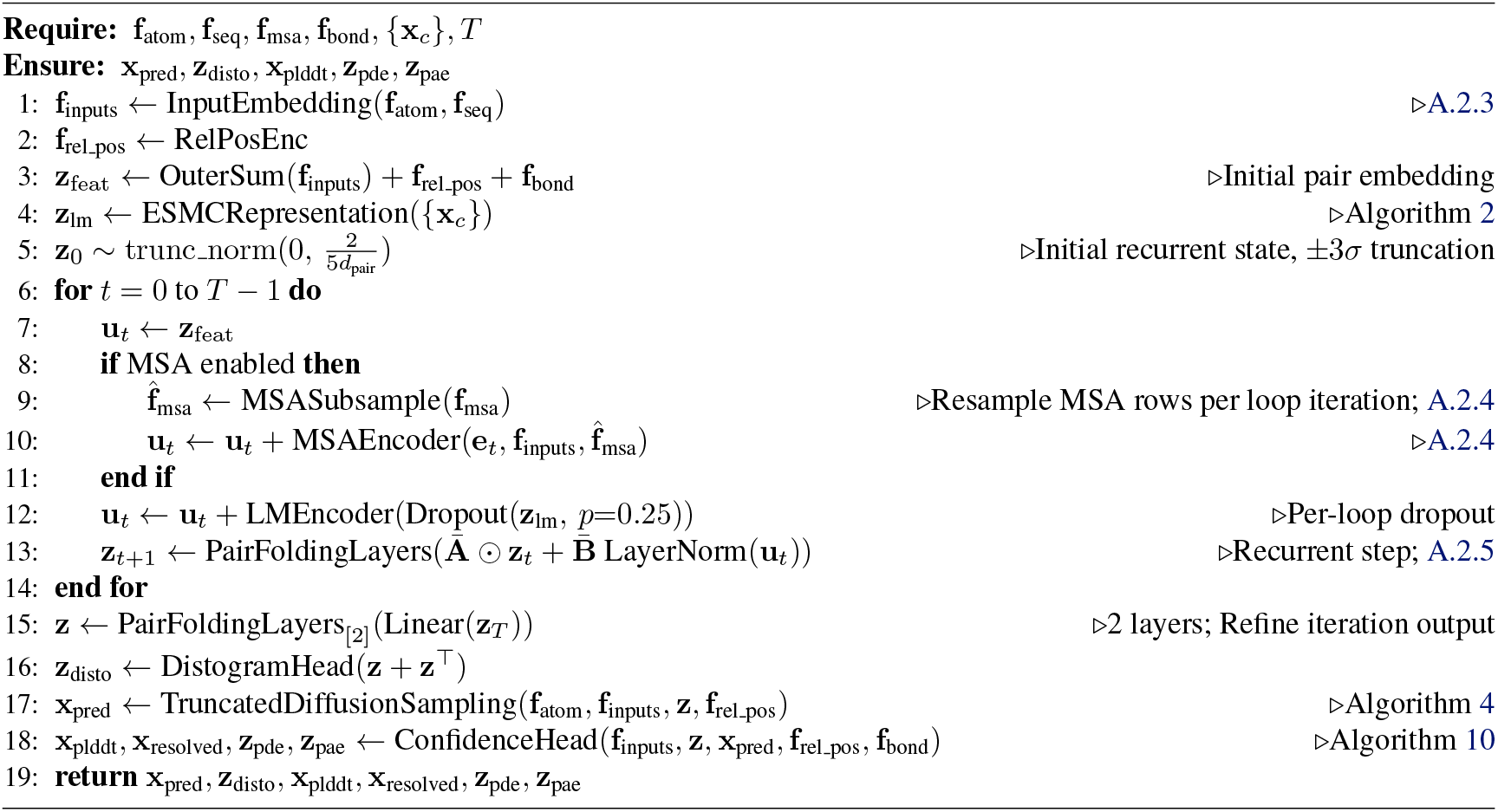

###### Non-protein tokens

Only protein tokens are processed through the language model. Non-protein tokens (nucleic acids, ligands) are not embedded via ESMC. At non-protein positions, the LM pair contribution **z**_**l**m_ is zero-filled before being added to the pair representation. Non-canonical amino acids are presented to the folding model using one token per atom. However, to ESMC, each non-canonical residue is presented as a single unknown token ‘X’ per residue. The ESMC representation is then expanded to one vector for each atom in the non-canonical residue.

###### Per-loop dropout

During training, 25% element-wise dropout is applied to the LM pair contribution **z**_**l**m_ each time it is added to the pair representation. A new dropout mask is sampled at each recurrent loop iteration; the LM forward and the projection to pair space are not recomputed. During inference, 30% dropout is used unless otherwise stated.

##### A.2.3. Input Embedding

The input embedding module converts raw atomic and sequence features into initial token-level representations.

###### Atom feature embedding

For each atom, the feature vector **f**_atom_ is constructed by concatenating: reference 3D coordinates **x**_ref_ (3 dims), formal charge (1 dim), attention mask (1 dim), element type one-hot (128 dims), and atom name characters one-hot (4 × 64 = 256 dims), for a total of 389 dimensions. This is projected to *d*_atom_ = 128 via a linear layer followed by layer normalization.

###### Algorithm 2

ESMC Representation Integration.

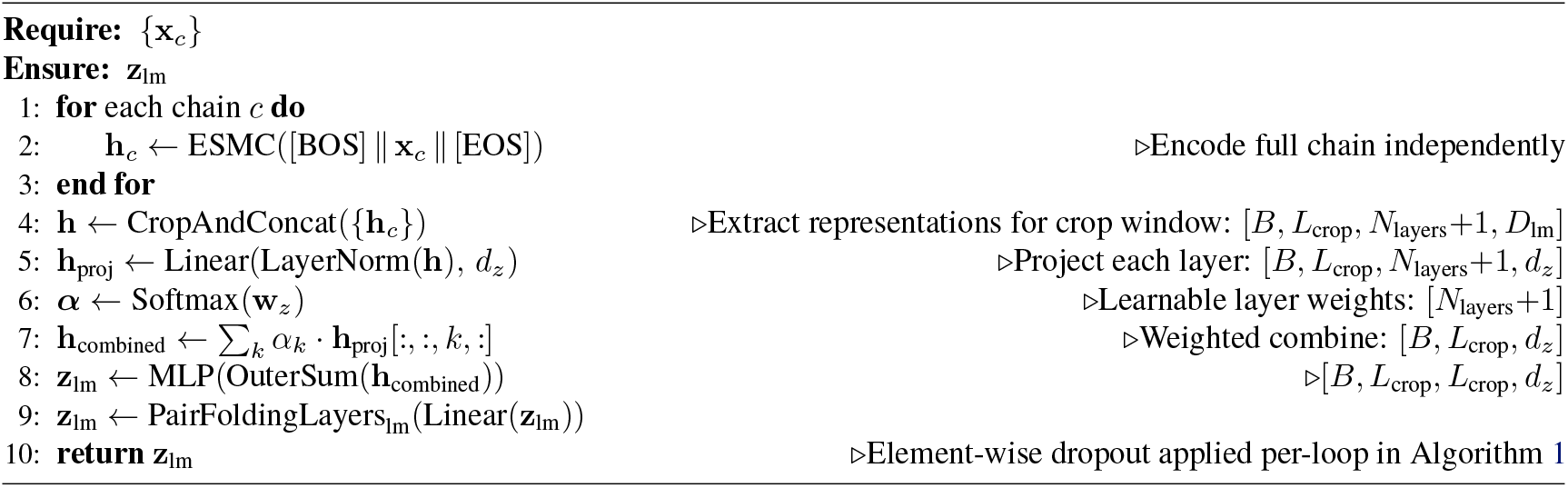

###### Atom attention encoder

The projected atom features are processed by a 3-block sliding-window atom transformer (A.2.6). In the input embedding context, no conditioning is applied—the atom encoder receives only the reference atom features, without noise conditioning or coordinate inputs.

###### Token aggregation

Atom-level representations are aggregated to token-level via a linear projection to *d*_token_*/*2 = 384 dimensions, followed by ReLU and scatter-mean over atoms belonging to each token.

###### Input feature concatenation

The token-level atom encoding is concatenated with the residue type **f**_seq_ and the (optional) MSA features **f**_msa_ (profile and deletion mean) to produce the input feature vector **f**_inputs_ of dimension *d*_inputs_.

#### A.2.4. MSA Module

ESMFold2 may optionally be conditioned on MSAs. In ESMFold2-Fast, the MSA module is disabled entirely, and MSA profile and deletion mean inputs are not provided.

The MSA encoder consists of 4 layers, each operating on MSA representations of dimension *d*_msa_ = 128 with 8 attention heads. In each layer, `OuterProductMean` is computed first, followed by pair-weighted averaging and feed-forward network on the MSA residual stream, then triangle multiplication (outgoing and incoming) and feedforward network on the pair stream. Triangle attention layers are also removed in the MSA encoder, as they do not have an observable effect on performance.

##### MSA dropout

During training, 15% row dropout is applied to the MSA track and 25% row/column dropout is applied to the pair track within the MSA encoder. In 10% of training iterations, we also dropout all non-query rows, which was found to be beneficial for single-sequence performance in ESMFold2.

##### MSA subsampling

During training, the MSA depth is randomly subsampled: a random number of sequences between 1 and min(available, max depth) is selected, always including the query sequence. We use max depth = 1024 across all stages. During inference, MSAs are subsampled to min(available, max depth) at every loop, which is vital for improved performance across recurrences.

##### MSA column masking

To supplement MSA subsampling, we also apply MSA column masking to increase seed diversity during inference. Rather than replacing uniformly sampled columns with unknown (X) (100), we found it more beneficial to randomly mask out columns in the `OuterProductMean`, thereby restricting coevolutionary information flow from input MSAs in the pair state. We apply 10% MSA column masking only at inference-time; training is unaffected.

#### A.2.5. Recurrent Folding Layers

ESMFold2 processes the pair representation through a series of folding layers, which takes the majority of the compute. To enable inference time scaling, the layers can be run iteratively in a loop, as described in Algorithm 1.

The initial pair embedding is computed as:

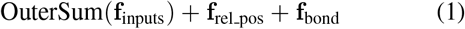

The recurrent pair state is initialized independently as **z**_**0**_ ∼ trunc_norm(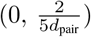) with ±3σ truncation.

To enable inference time scaling, the MSA is subsampled to 1024 sequences 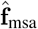, and the LM embedding **z**_**l**m_ is dropped out with probability 0.25 at each iteration. The pairwise update is computed as 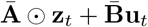.

Each pair folding layer is presented in Algorithm 3. Similar to (39), we find that the removal of sequence-wise residual stream does not degrade performance, and that triangle attention is unnecessary when models are compute matched. These choices enable a faster and simpler pair processing layer.

The standard residual update found in (22, 101, 102) **z** _*t*+1_ = **z**_***t***_ + *R*(**z**_***t***_) is difficult to optimize for large *T*. Prairie et al. (40) treats this as a discretized linear dynamical system:

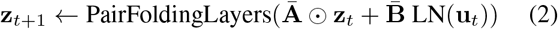

where Ā = exp(−Δ ⊙ A) is a Zero-Order-Hold-discretized negative-diagonal matrix and 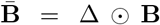 is Euler-discretized, with Δ, A, and B as learnable channel-wise parameters. Constraining Ā to have eigenvalues in (0, 1) prevents residual-state explosion across long recurrent unrolls. During training, the total number of loops *T* is sampled from a Poisson distribution with mean 3 (clamped to [1, 6]), and gradients are backpropagated only through the last ⌈3*/*2⌉ = 2 loops. At inference, *T* can be increased to scale test-time compute. After the loop, the final pair state **z**_***T***_ is passed through a linear readout followed by two pair folding layers before being handed to the diffusion module.

#### A.2.6. Diffusion Module

The diffusion module predicts atomic coordinates from the pair representation. The full inference sampler is given in Algorithm 4. It includes the truncated noise schedule which saves 33% of total steps.

##### Diffusion Conditioning

Conditioning signals are prepared from the pair layer outputs and noise level. Since ESMFold2 operates without a single representation, the single conditioning signal s_cond_ is derived solely from the input features **f**_inputs_ rather than from a processed single state as in AlphaFold 3. The noise level σ is embedded via a Fourier embedding of dimension 256 and added to s_cond_. The pair conditioning **z**_**c**ond_ is derived from the pair representation and relative position encoding.

##### Token Diffusion Transformer

The token-level diffusion transformer consists of 12 blocks with 16 attention heads operating at *d*_token_ = 768. Each block applies gated multi-head attention with pair bias followed by a conditioned transition block (SwiGLU FFN and an expansion factor of 4). Adaptive layer normalization conditions each block on the noise level through the single conditioning signal.

##### Denoising

The denoised prediction at noise level *t* with data scaling σ_data_ is given by:

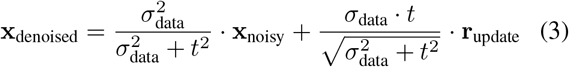

where r_update_ is the raw network output normalized by the input scaling factor 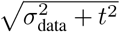, and *σ*_max_ = 16. During training, noise is sampled from a log-normal distribution: σ = σ_data_ · exp(*µ* + σ_std_ · *z*), where *z* ∼ 𝒩 (0, 1), *µ* = −1.2, σ_std_ = 1.5.

At inference, we apply EMA weights to the full model and solve for **x**_pred_ using the Euler-Maruyama sampler with modifications (23). We set noise scale λ = 1.003, step scale η = 1.5, γ_min_ = 1.0, and γ_0_ = 0.8.

##### Truncated Diffusion Sampling

The diffusion process uses *N* sampling steps with the noise schedule defined by σ_data_, *s*_max_, *s*_min_, and ρ. The schedule generates sigma values:

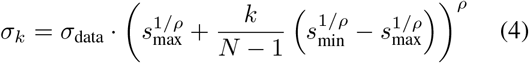

for *k* = 0, …, *N* −1, followed by σ_*N*_ = 0.

The full schedule spans σ ∈ [σ_data_ · *s*_min_, σ_data_ · *s*_max_]. However, at inference, the schedule is truncated via σ_max_: all schedule values exceeding σ_max_ are discarded and the initial sigma is capped at σ_max_. This skips the initial high-noise portion of the schedule where σ ≫ σ_data_ and the signal-to-noise ratio is negligible. At these extreme noise levels, the denoising network’s prediction is dominated by the scaling prefactor 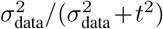, which is near zero, making these steps effectively wasted computation. We find σ_max_ = 256 did not degrade performance, and at default hyperparameter effectively results in 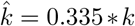. We use *N* = 100 which reduces to 68 sampling steps; we observed no benefit of increased sampling steps for structural quality (Figure S13).

##### Sliding Window Atom Attention

Atom-level diffusion expands the context length substantially, since each token (amino acid or nucleotide) contains multiple atoms whose positions must be predicted. This increases both the computional and memory requirements for atom-level diffusion far beyond the rest of the network. One approach to limiting the cost of atom diffusion is block local attention, where query atoms are partitioned into fixed-size blocks, each block attends to a local key window, and an explicit per-pair atom-pair tensor is constructed and added as attention bias (23). We identify this as a key bottleneck of the diffusion module, as the attention is implemented as a dense matrix multiply with pair bias and is poorly optimized by modern kernels.

ESMFold2 replaces the entire stack with a standard sliding-window attention transformer (window *w*=128) implemented via FlashAttention’s native sliding-window support, conditioned via AdaLN-Zero (103). The pair bias is removed entirely; positional information instead enters the queries and keys through 3D RoPE (A.2.6). The sliding window transformer is used in both the input embedding (A.2.3) and the diffusion module. In the diffusion module, the encoder additionally receives noisy atom positions. Algorithm 6–Algorithm 8 give the encoder, decoder, and shared transformer.

###### Algorithm 3

PairFoldingLayer.

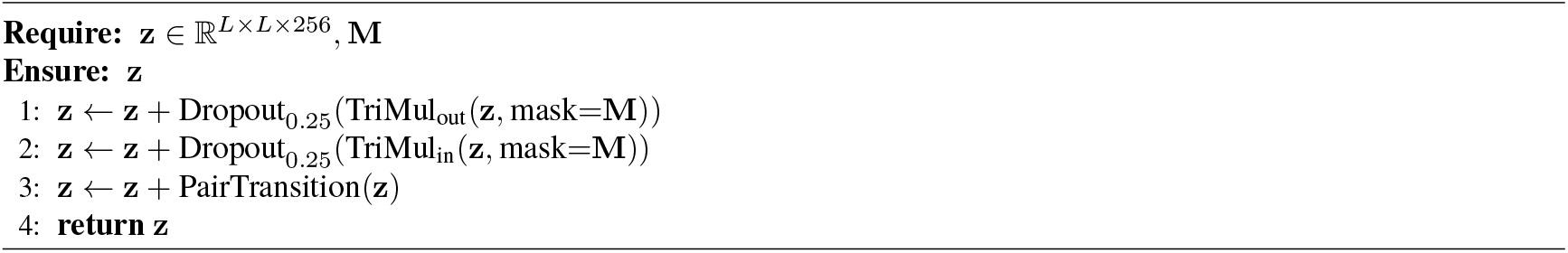

###### Algorithm 4

Truncated Diffusion Sampling.

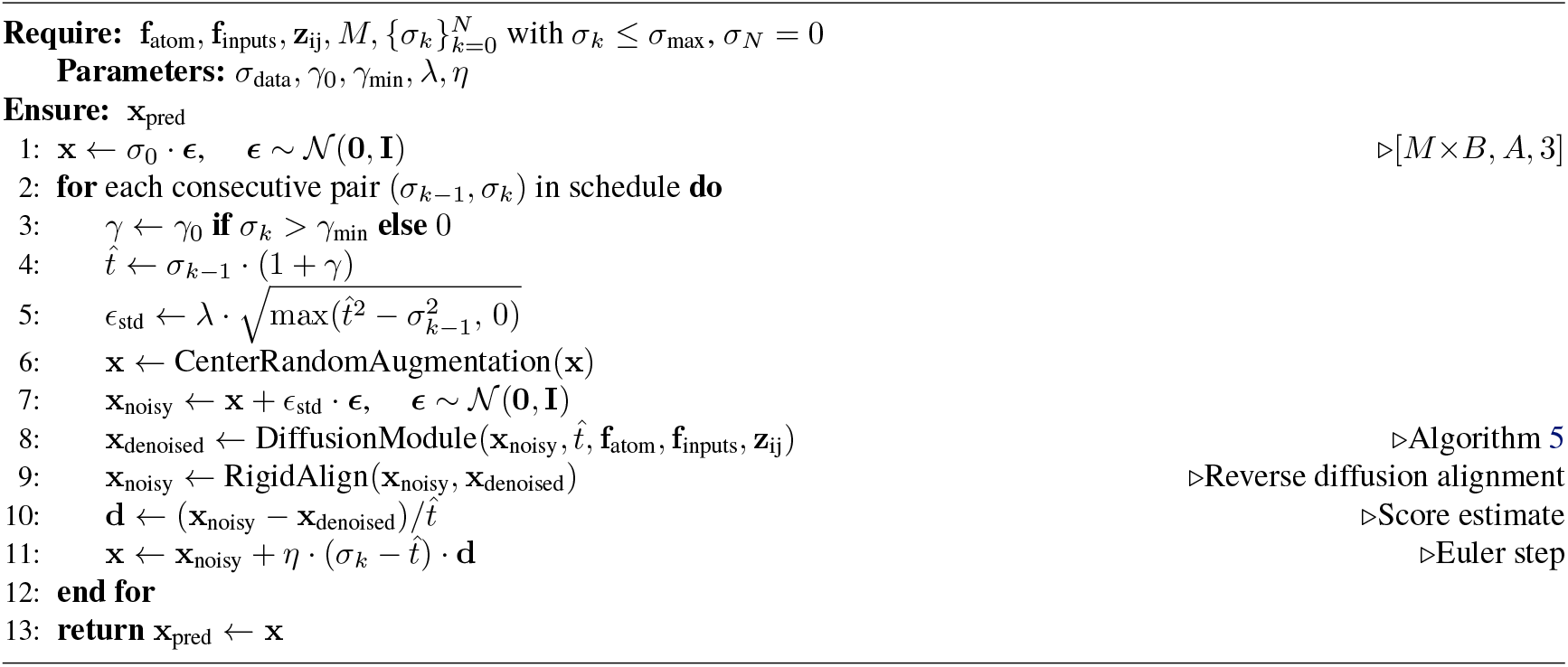

###### Algorithm 5

DiffusionModule.

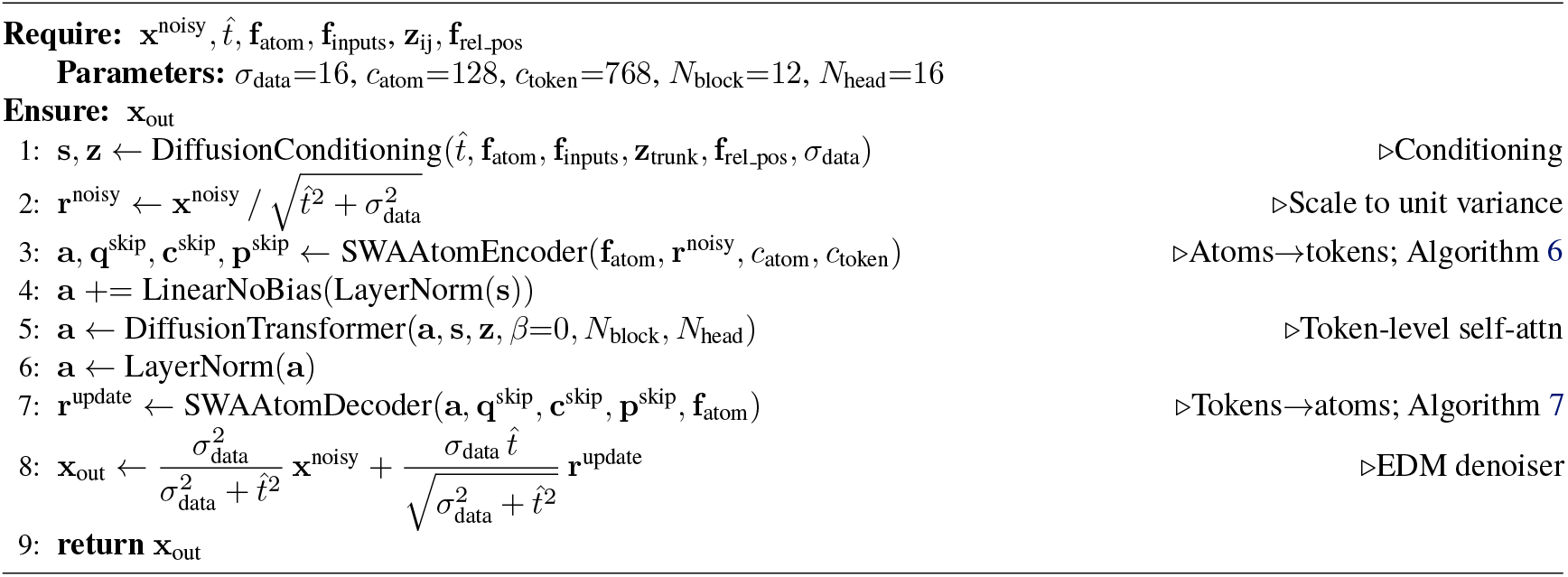

###### Algorithm 6

SWAAtomEncoder.

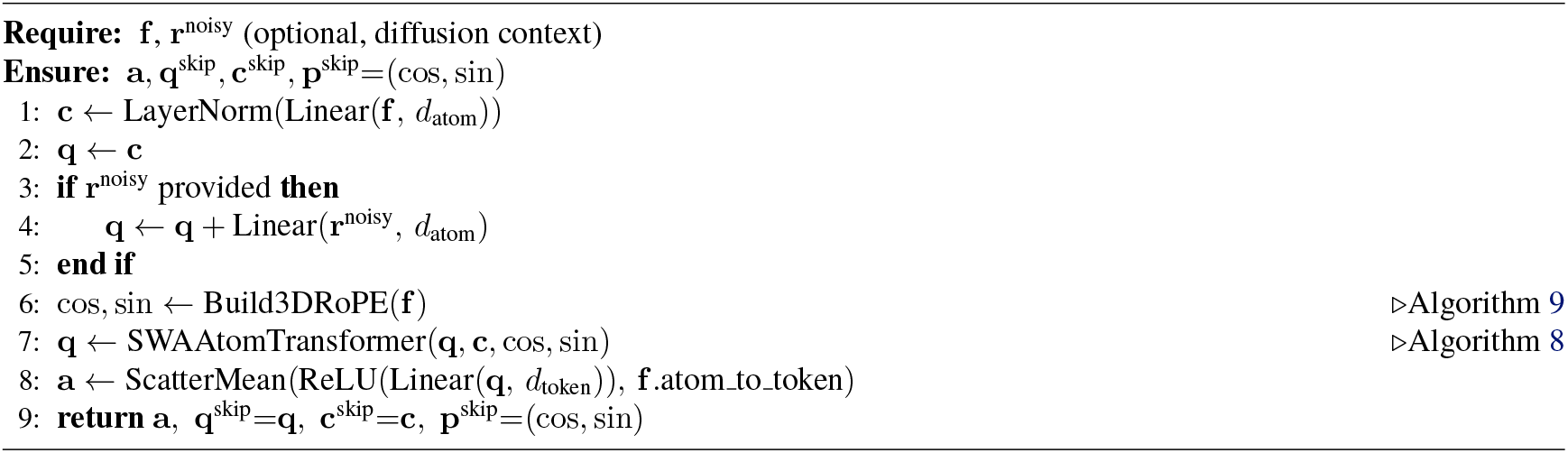

###### Algorithm 7

SWAAtomDecoder.

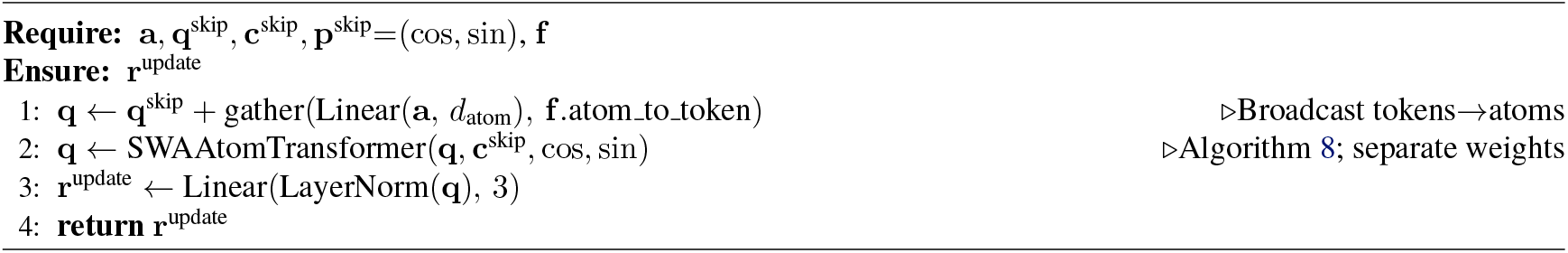

###### Algorithm 8

SWAAtomTransformer.

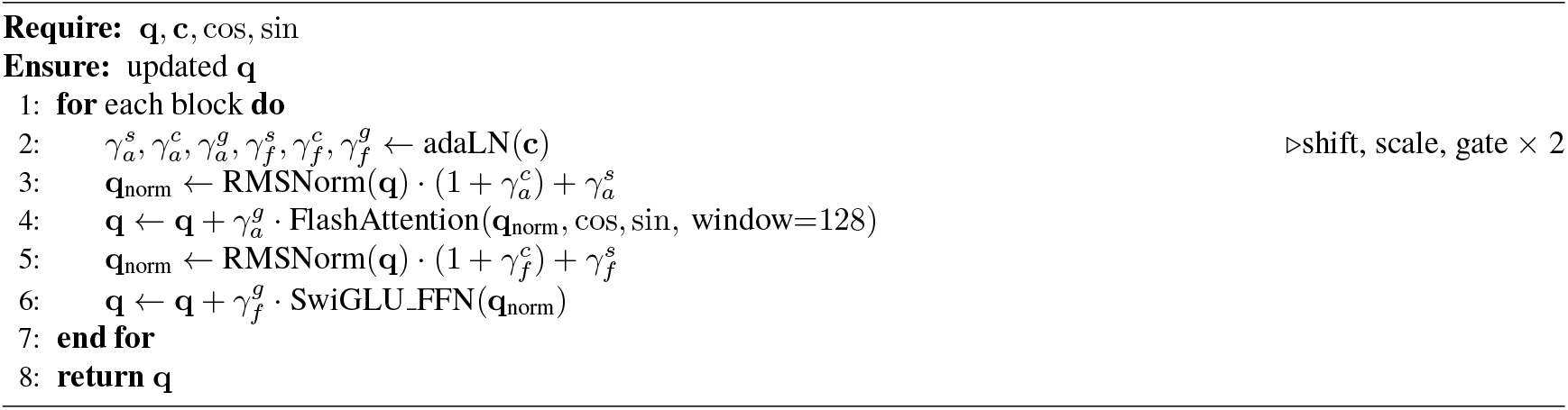

RoPE for Pairwise Features. Removing the explicit atom-pair bias means the attention has no direct view of inter-atom distances or residue membership. The signal is restored by encoding it into the queries and keys via 3D rotary position embeddings inspired by Su et al. (88) derived from two sources: 3D atom coordinates (spatial proximity) and per-residue space UIDs (which residue each atom belongs to). The resulting attention scores depend on relative atomic geometry without any explicit pair tensor. Algorithm 9 gives the construction.

#### A.2.7. Confidence Head

The confidence head predicts per-token and pairwise quality metrics: all-atom pLDDT, experimentally resolved status, predicted distance error (PDE), and predicted aligned error (PAE) (23). The confidence module is trained from the beginning of training, and the loss is added to the overall loss with weight *α*_confidence_ = 10^−4^.

The confidence head receives the final pair representation **z**_*ij*_ from the main model and residually adds a pair projection of **f**_inputs_, and the predicted coordinates **x**_pred_. The pair state *z*_*ij*_ is dropped 20% of the time during training to encourage the model to rely more on **x**_pred_ rather than learned features in **z**_*ij*_ which are agnostic to the final predicted coordinates. These representations are fed into four pair folding layers followed by linear projections for each metric (Algorithm 10). As pLDDT and resolved targets are single state features, a learnable pooling over **z** to project to a single state **s** prior to the output linear heads (RowAttentionPooling in Algorithm 10).

A mini-rollout of 20 diffusion sampling steps generates predicted coordinates, using default inference settings. Pairwise distances between representative atoms are computed and binned into a distogram with 39 bins spanning 3.25 to 50.75 Å. The distogram is embedded via a learned embedding layer and added to the pair representation.

#### A.2.8. Training Details

##### A.2.8.1 Losses

Diffusion MSE loss. The primary structural supervision is a per-atom mean squared error between the denoised diffusion sample and rigid-aligned ground-truth coordinates, with per-atom weights of 1× for protein, 5× for nucleotides, and 10× for ligand atoms. The per-sample loss is scaled by 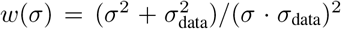, with training noise σ sampled from a log-normal distribution with *µ* = −1.2, *σ*_std_ = 1.5 (A.2.6).

###### Smooth LDDT loss

A differentiable LDDT loss term (23) computed between the denoised diffusion sample and ground-truth coordinates. For each pair of atoms within a chain-type-dependent cutoff (30 Å for nucleic-acid pairs, 15 Å otherwise), the pairwise distance deviation 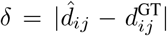 is passed through a sum of sigmoids at thresholds {0.5, 1, 2, 4} A. This loss complements the MSE by providing alignment-invariant, distance-based supervision on local geometry. We empirically observed this loss to be vital for global folding accuracy, but detrimental for all-atom quality (such as interatomic clashes). Thus, this is included only during the initial training stage.

###### Bond loss

A mean squared error polymer–ligand bond lengths (23) to encourage correct bond geometry in covalently bonded ligands to polymers (such as in post-translational modifications and cofactors). This loss is only active from stage 2.

###### Confidence losses

The four cross-entropy losses on the confidence head outputs (all-atom pLDDT, PDE, resolved, PAE; A.2.7) are summed and added to the total with weight *α*_confidence_ = 10^−4^, applied jointly from the start of training. Losses are applied only to PDB structures with resolution ≤ 4.0 Å.

###### Distogram loss

A linear head predicts inter-residue distance distributions from the symmetrized trunk pair representation, logits = Linear(**z** + **z**^⊤^), supervised with a cross-entropy loss against binned ground-truth distances (64 bins, 2.0–22 Å). The distogram loss is not active in stage 3 since the trunk is frozen.

###### Loss functions

The structural losses for all-atom structure prediction consists of the diffusion MSE, smooth LDDT, and polymer-ligand bond losses,

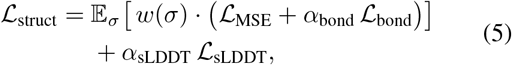

with *α*_bond_ = 1 (from stage 2), and *α*_sLDDT_ = 1 (stage 1 only) (see A.2.8.2).

The confidence head is supervised by the sum of four cross-entropy terms,

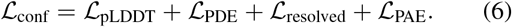

The final loss function is a weighted sum of the structural, confidence, and distogram loss,

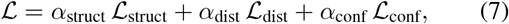

with *α*_struct_ = 4.0, *α*_dist_ = 3 × 10^−2^ and *α*_conf_ = 10^−4^. Note that ℒ_dist_ only supervises the main pair folding layers, ℒ_struct_ supervises the pair folding layers and the diffusion head, and ℒ_conf_ supervises only the confidence head due to a `stopgrad` on **z, f**_inputs_ **x**_pred_.

###### Algorithm 9

Build3DRoPE.

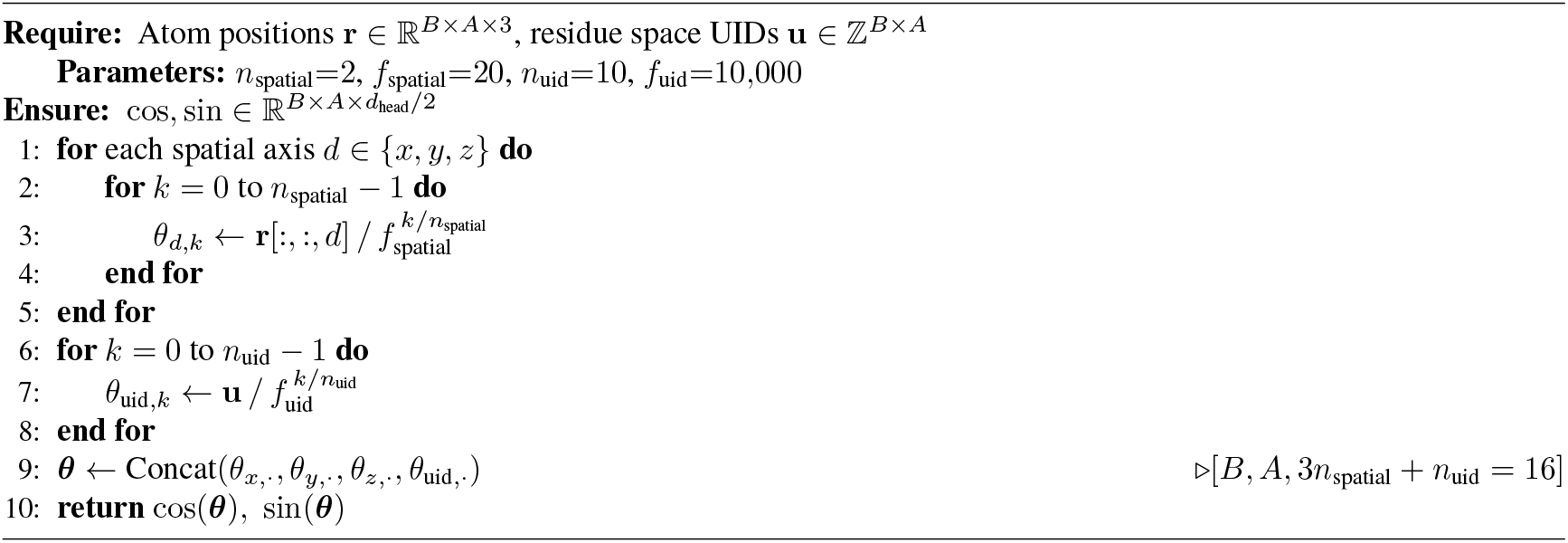

###### Algorithm 10

ConfidenceHead.

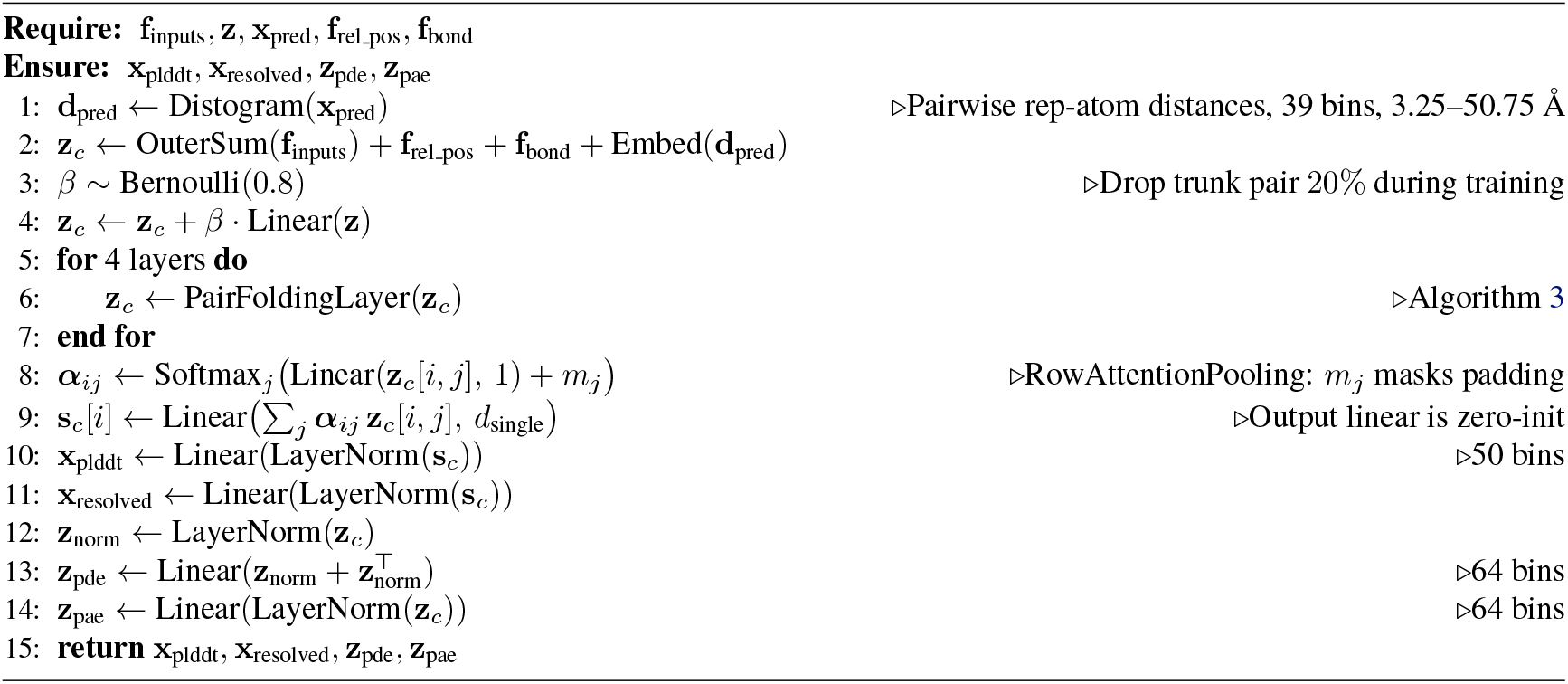

##### A.2.8.2 Training stages

ESMFold2 is trained in three progressive stages with increasing context size (Table S7). In stage 1, the full model is trained end-to-end, including the PAE head. The distogram, MSE loss and smooth LDDT losses are used to train the main structure prediction model, while the confidence heads are trained with the resolved, all-atom pLDDT, PAE, and PDE losses. In stage 2, the smooth LDDT loss is disabled (which benefits all-atom quality) and the polymer-ligand bond loss is enabled. Finally, in stage 3, the main pair folding layers are frozen using EMA weights and diffusion and confidence heads are trained, as we observed that the pair folding layers converge more quickly. This strategy improves all-atom quality while preventing overfitting. For stage 1 and 2, a step-scale learning rate schedule is used (decay factor of 0.95 every 50k steps) with a maximum rate of 1.8 × 10^−3^, including a 1000-step warmup phase. The AdamW optimizer is used with β_1_ = 0.9, β_2_ = 0.95, ϵ = 10^−8^, and weight decay 10^−4^. For the final stage 3, the learning rate is reduced to a fixed 1.0 × 10^−4^.

ESMFold2 was trained on 256 H100 GPUs for a total of 127 hours split across the three stages, and ESMFold2-Fast was trained for 85 hours. To reduce memory pressure from the ESMC language-model backbone, ESMC weights are cast to FP8. All other modules are trained in bfloat16 mixed precision, except the diffusion module which is trained in float32. Triangle multiplication use cuEquivariance fused kernels during training. The trunk is compiled with `torch.compile` with inputs padded to the crop length, so the compiled graph is reused across the batch.

#### A.2.9. Data

##### A.2.9.1 Training data

The ESMFold2 training dataset draws from two sources: (i) PDB structures and (ii) a filtered subset of the AlphaFold Database (AFDB) (80) containing AlphaFold2 predicted samples with an average pLDDT *>* 70. During all stages, samples are drawn from PDB and filtered AFDB with probabilities of 30% and 70%, respectively.

###### PDB dataset preparation

Protein chains are clustered at 40% sequence identity, while nucleic acids and short peptides (*<* 10 residues) are clustered at 100% sequence identity. Small molecules are clustered by CCD identifier. Clustering uses MMseqs2 with `easy-cluster --min-seq-id [40,100] --cov-mode 0 -c0.8`. For benchmarking, we train on PDB structures released prior to 2021-09-30. The following filters are applied to the training data: structures with resolution > 9 Å or more than 300 polymer chains are discarded, chains consisting entirely of unknown residues are removed, atoms not in the CCD-defined set for a residue or small molecule are stripped, protein chains with a *>* 10 Å gap between consecutive C*α* atoms are removed, and small molecules in the AlphaFold3 ligand exclusion list (and crystallization aids) are excluded. Clashing chains, defined as chains where *>* 30% of atoms lie within 1.7 Å of another chain, are removed; atom pairs annotated as metal coordination bonds are excluded from this filter. During training, any sample containing more than 30% unknown residues after cropping is skipped.

Cropping. Each training sample is cropped over tokens using one of three strategies, sampled per example with dataset-specific weights. *Contiguous cropping* selects, for each chain, a contiguous run of polymer residues and/or ligand atoms, with the per-chain budget proportional to chain length. *Spatial cropping* samples a reference token-center atom at random and keeps the nearest tokens by Euclidean distance to that reference. *Spatial-interface cropping* restricts the reference to interfacial token-center atoms (those within 15 Å of another chain’s token-center atom) before applying the same nearest-neighbour selection. For weighted-PDB and disordered-PDB samples a preferred chain or interface may be supplied, in which case the reference atom is drawn from that chain or interface. The default strategy weights (contiguous / spatial / spatial-interface) are (0.20, 0.40, 0.40) for multimer training data and (0.25, 0.75, 0.0) for monomer distillation, matching the values used in (104). The full contiguous and spatial selection algorithms follow Algorithms 1 and 2 of (104).

###### Synthetic data

For synthetic distillation, we use the AlphaFold Database of predicted monomer structures from UniProt. We retain entries satisfying: mean pLDDT *>* 70; sequence length between 200 and 1000; and a C*α*–C*α* contact ratio (number of pairs within 8 Å divided by polymer length) greater than 0.5, which serves as a globularity proxy. We additionally cluster the dataset at 70% and 40% sequence identity. For each cluster at 70% identity we first take the longest sequence, then within that group select the sequence with the smallest pLDDT standard deviation that passes the above filters; this favors uniformly confident predictions and avoids biasing the dataset toward easy proteins. Clustering uses `mmseqs easy-cluster --min-seq-id` $`ID --cov-mode 0 -c 0.8`. After filtering, the augmentation set contains approximately 8.8M structures.

**Table S7.** Training stages for ESMFold2. Context size and diffusion batch size increase across stages; auxiliary loss weights are adjusted between the initial stage and fine-tuning. Stage 3 freezes the main folding layers and trains the diffusion and confidence heads only.

| Stage | Training steps* | Max Tokens | Max Atoms | Diffusion batch size | Full training | Bond loss | Smooth LDDT | Distogram | Confidence |
| --- | --- | --- | --- | --- | --- | --- | --- | --- | --- |
| 1 | 50,000 | 384 | 9,216 | 48 | ✓ | 0 | 1 | 0.03 | $10^{-4}$ |
| 2 | 5,000 | 640 | 15,360 | 32 | ✓ | 1 | 0 | 0.03 | $10^{-4}$ |
| 3 | 11,000 | 768 | 18,432 | 32 | ✗ | 1 | 0 | — | $10^{-4}$ |

\* ESMFold2-Fast was trained with 60,000, 5,000, and 10,000 steps for each stage, respectively.

##### A.2.9.2 Validation data

To fairly assess the model performance for generalization, we generated three validation datasets: a general validation set, a low-homology antibody-antigen set, and low-homology protein-protein interface set. All validation sets were constructed using PDB entries released between 2021-09-30 and 2023-01-13, corresponding to the training and evaluation date cutoffs.

###### General validation set

The general validation set covers a diverse set of biomolecules including small molecules and nucleic acids to assess general performance and guide model selection.

We exclude structures with resolution worse than 4.5 Å, more than 1,000 chains, or more than 2,560 tokens. We remove any polymer chains with *>* 40% sequence-identity to training data. Polymer–ligand interfaces were removed if any training complex contained both a polymer in the same 40% cluster as the candidate polymer and a ligand with Tanimoto similarity ≥ 0.85. Interfaces with more than one ligand, ligands used as crystallization aids, or ligands in AlphaFold3’s ligand exclusion list are removed.

Surviving low-homology interfaces were grouped by molecular-type pair and capped: 600 protein–protein, 600 protein–ligand, 200 ligand–ligand, 100 protein–DNA, 100 DNA–DNA, and 50 DNA–ligand. Protein–RNA, RNA– RNA, DNA–RNA, and RNA–ligand interfaces were retained without sampling. When a type exceeded its budget, interfaces were first grouped by redundancy key (the unordered pair of polymer clusters for polymer–polymer interfaces, or the (polymer-cluster, ligand-CCD) pair for polymer–ligand interfaces), one representative was chosen uniformly at random per group to fit each budget.

Monomers were limited to 40 protein monomers with random sampling, while all low-homology DNA and RNA monomers were retained. All PDB entries contributing at least one sampled interface or monomer was used after filtering to at most 2,048 tokens. Then, we include all low-homology chains and interfaces in these PDB entries. This results in 2,309 low homology interfaces across 1,255 complexes.

###### Antibody-antigen validation set

Candidate antibody-antigen structures were identified using SAbDAb (105), filtered to entries with a heavy chain and a protein antigen. Each Fv was matched to the first bioassembly of its parent PDB entry. We remove all NMR structures, any structures with resolution higher than 4.5 Å, and more than 300 chains.

In order to reduce training homology and ensure structural quality, we:

- *Filter by homology:* Drop candidate interfaces if any antibody and antigen chain exhibits *>* 40% sequence identity and *<* 80% coverage to a chain in the training set (via MMSeqs2 with parameters `-c 0.8 -cov-mode 0 -s 7.5`).
- *Filter by chain quality:* Remove chains that have poor coverage, or where any pair of consecutive residues has a Cα–Cα distance *<* 10 Å.
- *Resolve steric clashes:* Identify inter-chain heavy atom clashes (distance *<* 1.7 Å). If 30% or more of the total heavy atom count exhibits clashes, drop the chain with the higher clashing fraction.
- *Filter by length:* Retain only complexes with a token count *c* such that 10 ≤ *c* ≤ 2560.
- *Filter by resolution:* Retain only complexes crystallographic resolution less than 2.5 Å.
- *Remove peptides:* Exclude antibody–peptide interfaces, which are defined as having an antigen length of fewer than 16 residues.

In order to deduplicate at an interface level, we additionally assign each interface chain with the MMseqs2 cluster identifier. For example, chains A, B may map to cluster X and chains C, D to cluster Y. We define redundancy at the cluster level, so that an A-C interface is mapped to a cluster level interface ID of X-Y. For each cluster level interface, a representative is chosen based on the lowest crystallographic resolution, with length serving as a tie-breaker. This results in 135 interfaces across 86 targets.

###### Protein-protein validation set

The protein–protein interaction (PPI) validation set was constructed using the same pipeline as above, with three modifications. First, candidates were drawn from the full PDB snapshot **excluding** antibody-antigen complexes defined in SAbDAb. Second, in addition to the sequence-level homology filter, we applied a structure-level homology filter using Foldseek-Multimer (106) with default settings except increased sensitivity `-s 7.5`. After per-chain quality filtering, interface-chain subsetting, and token-budget filtering, each candidate interface was structurally aligned against every multimer in the training set, discarding any interfaces with a query TM-score ≥ 0.5 against any training complex. Third, we apply a stricter resolution cutoff of 2.0Å. This results in 190 targets and 198 interfaces.

##### A.2.9.3 MSA generation

Multiple sequence alignments (MSAs) provide evolutionary context that informs structure prediction. For each query chain we search genetic databases for homologous sequences, align them, and use the resulting MSA to build pairwise representations of co-evolutionary signal.

###### PDB MSA Generation

For each unique protein chain in the training set we run JackHMMER, HHblits, or MMseqs2 against six sequence databases. JackHMMER searches use `-N 1 -E 0.0001 --incE 0.0001 --F1 0.0005 --F2 0.00005 --F3 0.0000005`; HH-blits uses `-n 3 -e 0.001 -realign max 100000 -maxfilt 100000 -min prefilter hits 1000 -p 20 -Z 500`; MMseqs2 uses `-s 7.5 --num-iterations 2 --max-seqs 1000`. From each sequence database, we take sequences from each database in the order shown in Table S8 until we reach the maximum MSA limit of 16,384 − *N*, where *N* is the number of paired MSAs constructed from UniProt, capped at *N* ≤ 8,191 paired sequences.

**Table S8.** Sequence databases and search settings used to construct training MSAs.

| Database | Search method | Sequence cap |
| --- | --- | --- |
| UniRef90 ((28)) | JackHMMER | 10,000 |
| UniProt | JackHMMER | 50,000 |
| Uniclust30 + BFD | HHblits | — |
| Reduced BFD ((107)) | JackHMMER | 5,000 |
| MGNify ((108)) | JackHMMER | 5,000 |
| Logan50 ((109)) | MMseqs2 | 3,000 |

###### MSA pairing for multi-chain complexes

For multi-chain protein complexes, MSA sequences must be paired across chains to provide co-evolutionary signal about inter-chain contacts. Pairing uses the NCBI taxonomy identifiers extracted from UniProt MSA headers: for each species (taxonomy ID) appearing in the UniProt MSAs of two or more chains, we select one sequence per chain from that species and group them into a single paired MSA row. This produces phylogenetically coherent pairings—sequences from the same organism are aligned across chains.

###### AlphaFold DB MSA Generation

For AFDB structures used in synthetic distillation, MSAs are generated with a modified version of the ColabFold (110) MMseqs2 search pipeline against UniRef30 (release `2023-02`) and the ColabFold environmental database (`colabfold envdb 202108`). The pipeline is modified to substantially reduce MSA generation time. For each query, a 3-iteration MMseqs2 profile search was run (`-e 0.1 --max-seqs 1000 --k-score seq:96,prof:80`), hits are expanded to cluster members (`--expansion-mode 0 -e inf –dif 128 --max-seq-id 0.95`), re-aligned with the profile (`-e 10 --max-accept 1000 --alt-ali 10`), and a diversity filter is applied (`--diff 128 --qid 0,0.2,0.4,0.6,0.8,1.0 --qsc 0 --max-seq-id 0.95 --filter-min-enable 1000`). The same pipeline, seeded by the UniRef profile, is then run on the environmental database and the two alignments are merged. Since AFDB structures are monomers, no paired MSAs are constructed.

###### MSA generation for evaluations

For all evaluations, we generate MSAs using ColabFold search against UniRef30 and environmental database using default parameters.

#### A.2.10. Benchmark Results

For all benchmark results, baseline models are run with 10 recycles and 200 diffusion steps. ESMFold2 uses 10 loops and a truncated diffusion schedule with 68 diffusion steps unless otherwise specified. The standard DockQ thresholds of ≥ 0.23, ≥ 0.49, and ≥ 0.8 are used to define acceptable, medium and high quality pass rates, respectively (111). To select the best sample across multiple seeds and diffusion samples, we simply use ipTM for complexes and pLDDT for monomers.

##### ESMFold2 inference settings

For all evaluations, we apply the following default settings to ESMFold2:

- max MSA depth of 1024 rows, subsampled at every loop,
- MSA column masking at rate 0.1, applied once before recurrence,
- ESMC dropout at rate 0.3, applied at every loop,

We observed MSA input diversity to be critical for inference-time scaling of ESMFold2, and simply applying ESMC dropout to ESMFold2-Fast resulted in poor scaling. We introduce another source of input diversity for ESMFold2-Fast by masking the input sequence to ESMC at rate 10%, applied once per seed before learned pooling. This significantly recovers scaling performance, and we apply this for FoldBench antibody-antigen and protein-protein interaction scaling evaluations (Figure 2D).

##### FoldBench

FoldBench selects diverse interface and monomeric structures in a low-homology dataset, providing a challenging evaluation setting for both folding of monomers and co-folding of molecular assemblies (33).

Models are evaluated on all 172 antibody-antigen targets and 278 / 279 protein-protein targets. Some models fail to produce valid predictions for all targets, in this case their performance is marked with a ^*^. We exclude the structure 8gqp from the protein-protein interaction target list as it is a complex including a D-protein binder, and passing it as a standard protein input to structure prediction methods would result in a prediction of the L-protein version. On all other Foldbench subsets, models are evaluated on the intersection of targets where all methods produced valid predictions.

Foldbench subsets are quite small; as a result, we observed high variation between different runs of the same methods at different seeds. To compensate for this, we fold each target using 25 seeds and 5 diffusion samples per seed for each method, then resample with *n*_boot_ = 200 samples to estimate the performance at 5 seeds × 5 diffusion samples. Error bars denote ±1*σ* over this bootstrap.

ESMFold2 achieves strong performance on monomer structure prediction (84%± *<* 0.1), antibody-antigen (50% ± 2), and protein-protein interactions (70% ± 1). In MSA-contioned folding, ESMFold2 is the strongest model across all tasks (89%± *<* 0.1, 53% ± 2, 76% ± 1, respectively). ESMFold2 also maintains strong performance on other Fold-bench subsets, including protein-ligand (61% ± 1), protein-DNA (79% ± 1) and protein-peptide (82% ± 4). ESMFold2-Fast is also competitive across all benchmarks, generally performing on par with the larger model in single sequence mode.

##### Runs N’ Poses

To probe generalization in receptor–ligand co-folding beyond training data, ESMFold2 is evaluated on the Runs N’ Poses (34) benchmark, comprising 2,600 systems. For each target, five seeds are co-folded with five samples each, and the top candidate is selected by ipTM score. For systems with multiple ligands, we score each ligand pose independently. Reported n is the total number of scored ligands. Ligands without defined SuCOS score are excluded from the success rate. Success is defined as lDDT-PLI *>* 0.8 and BiSyRMSD *<* 2 Å (Figure 2C). Error bars represent ±1*σ* over *n*_boot_ = 200 resamples of the test set. Results are reported on the intersection of structures where all models produced valid predictions. Performance binned by SuCOS similarity to training set is shown in Figure S8.

##### PoseBusters

ESMFold2 is assessed on protein–ligand docking using the PoseBusters (112) set of 308 receptor– ligand systems. Five samples per system are co-folded across five seeds, and results are reported for the highest predicted ipTM. For each prediction, the reference ligand conformer that minimizes the bidirectional symmetry-corrected RMSD (BiSyRMSD) is selected, and the percentage of predictions within 2 Å of deviation is reported (Figure S9). Performance is reported on the intersection of structures where all models produced valid predictions.

##### Comparison with Boltz-2

**Boltz-2** (113) is trained using PDB data released prior to 2023-06-01. Foldbench is constructed using PDB data released after 2023-01-13, which means Foldbench evaluation data is part of the Boltz-2 training set.

To construct a fair comparison we generate a dataset based on recent protein-protein complexes released in the PDB. We first apply the same procedure used to generate the training set (A.2.9) on all structures released between 2025-06-30 and 2026-01-21. We applied two futher filters:

- Structures with length *<* 40 or *>* 2048 are removed.
- All chains are clustered at 40% sequence identity alongside the training set. If any chain falls into the same cluster as a chain in the training set, the full cluster is removed. This ensures only interactions where all chains are sufficiently distant from the training set are maintained.

ESMFold2 achives a pass rate of 70% ± 4 on this dataset from a single sequence (Figure S10). Using an MSA, ESM-Fold2’s performance rises to 77% ± 3, significantly higher than Boltz-2 (70% ± 3). ESMFold2-Fast is also competitive on this benchmark, with a pass rate of 66% ± 3.5.

##### ESMC pseudo-perplexity and folding performance

Pseudo-perplexity (PPPL) (114) quantifies how unexpected a sequence is under a language model, with lower values reflecting more predictable sequences and higher values indicating unusual ones. It is defined by:

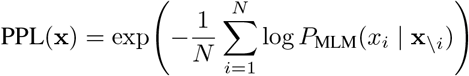

where *N* is the sequence length and *P*_MLM_(*x*_*i*_ | **x**_\*i*_) is the masked language model’s predicted probability of residue *x*_*i*_ given all other positions. Each position is masked and scored independently, then aggregated into a single perplexity value.

**Figure S6.**
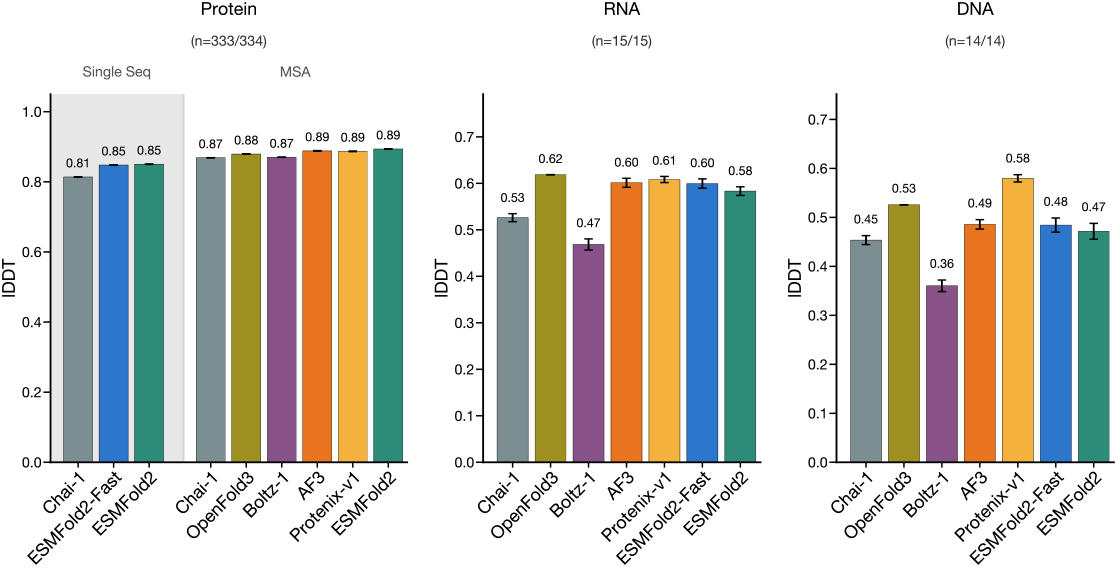
FoldBench Monomers. ESMFold2 is benchmarked against existing models across monomer modalities of FoldBench. Success rate is reported as mean lDDT.

**Figure S7.**
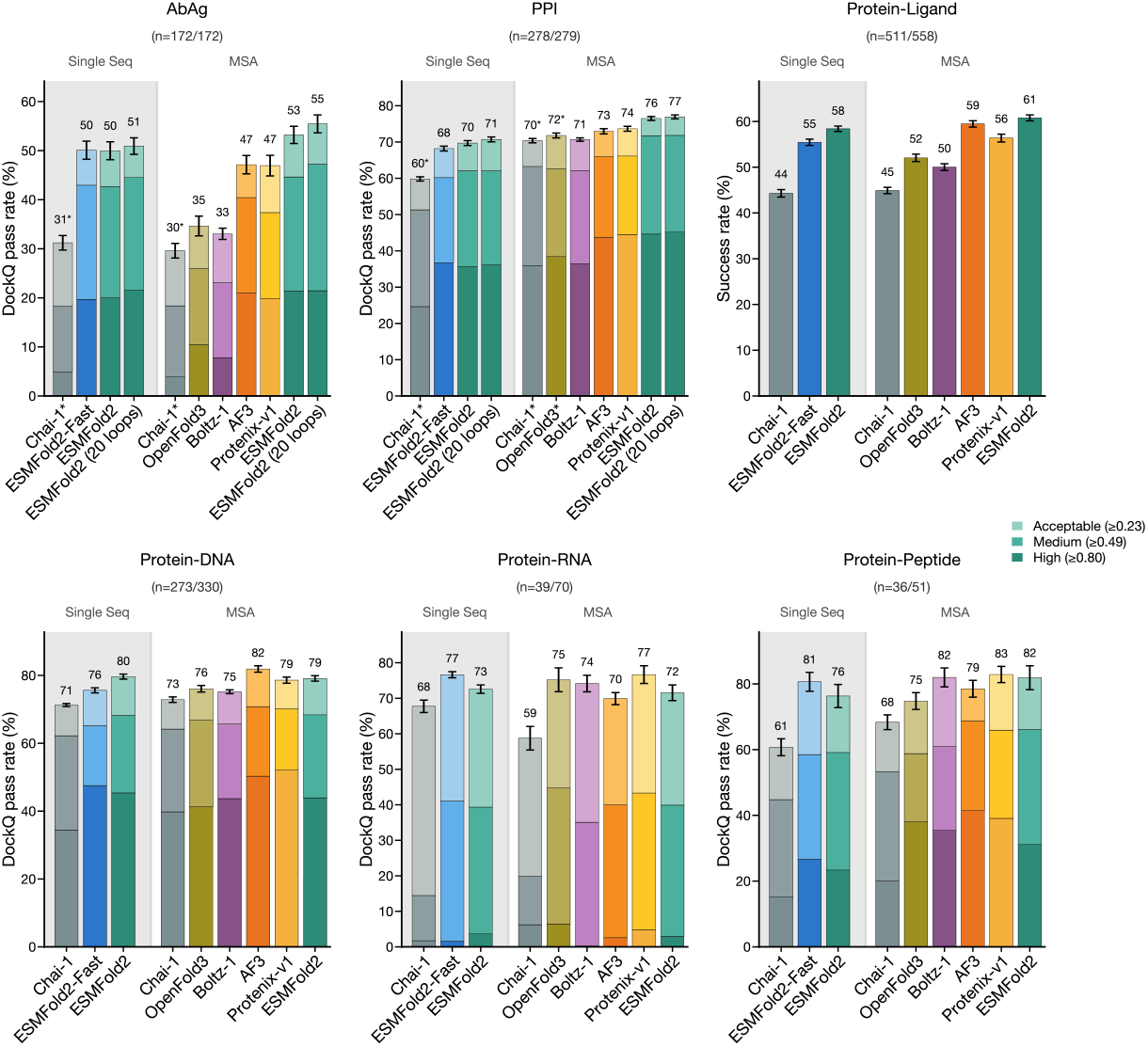
FoldBench Interfaces. ESMFold2 is benchmarked against existing models across diverse biomolecular interface modalities. Success rate is reported as the percentage of structures with predicted DockQ *>* 0.23 for interfaces, and the percentage of predictions with L-RMSD *<* 2 Å and lDDT-PLI *>* 0.8 for protein-ligand systems. ^*^ denotes the model did not make predictions for all targets. Chai-1 folds 148/172 antibody-antigen and 251 / 278 protein-protein targets. OpenFold3 folds 275 / 278 protein-protein targets.

**Figure S8.**
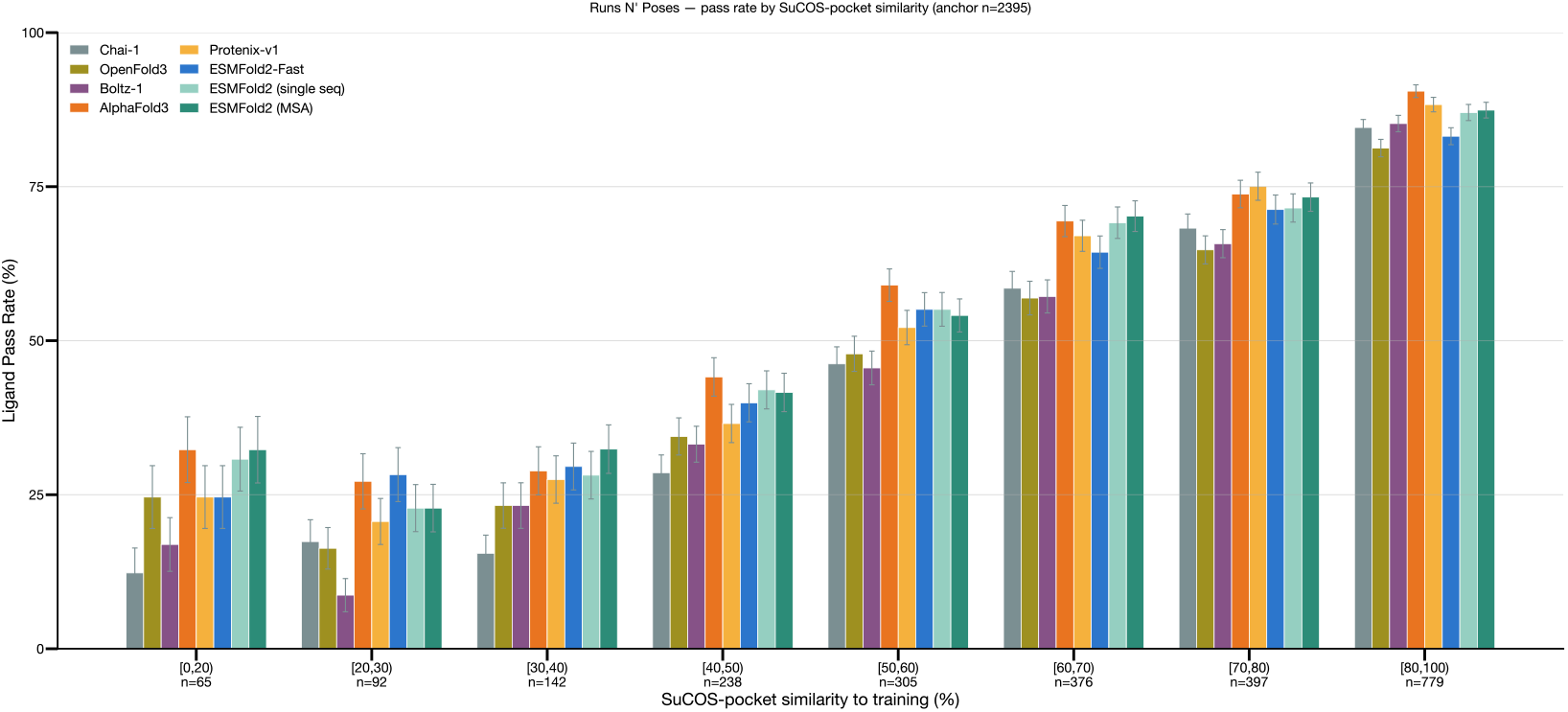
Performance on the Runs N’ Poses benchmark binned by SuCOS similarity to training set. Success rate is reported as the percentage of ligands with lDDT-PLI *>* 0.8 and BiSyRMSD *<* 2 Å. Error bars represent ±1*σ* over *n*_boot_ = 200 resamples of the test set. Results are reported on the intersection of structures where all models produced valid predictions.

**Figure S9.**
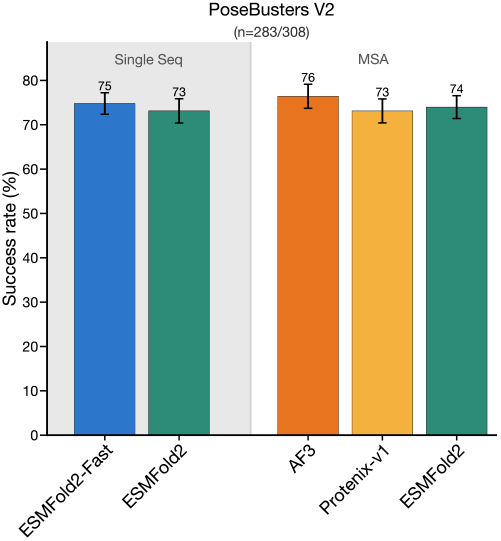
Protein–ligand docking on the PoseBusters refined set (308 receptor–ligand systems). Success rate is reported as the percentage of predictions within 2 Å BiSyRMSD of the reference. Error bars represent ±1*σ* over resamples of the test set. Results are reported on the intersection of structures where all models produced valid predictions.

We use this metric to investigate the relationship between ESMC sequence predictability and ESMFold2 structure prediction quality. For each interface, we compare the predicted DockQ against the mean ESMC PPL of the two interacting protein sequences, binning targets by PPL of the chain sequences (Figure S11). Comparing the single-sequence and MSA-conditioned regimes, we find that MSA information provides little benefit at low PPL but improves DockQ markedly for higher-PPL sequences, indicating that the MSA evolutionary context is primarily used to compensate for the language model uncertainty, and is otherwise redundant for well-characterized sequences.

**Figure S10.**
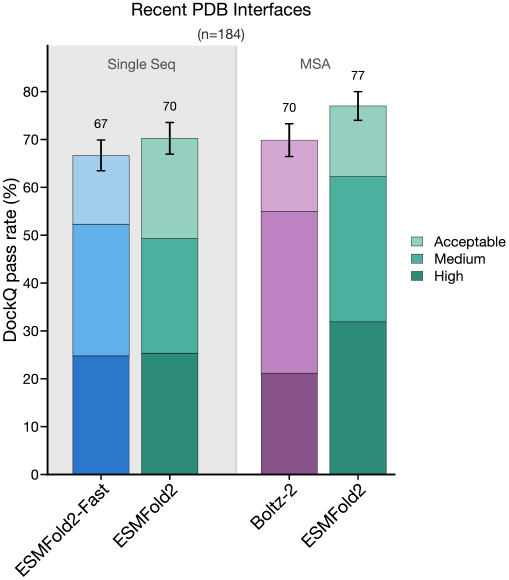
Performance of ESMFold2 and Boltz-2 on a recent protein-protein interaction evaluation dataset which is temporally held out of both models, and also held out of ESMFold2’s training set at 40% sequence identity. Error bars indicate ±1*σ* over resamples of the test set.

##### Confidence head calibration

The confidence module predicts four metrics: (1) per-atom pLDDT, (2) pairwise aligned error (PAE, from which pTM and ipTM are calculated), (3) pairwise predicted distance error (PDE), and(4) whether each atom is resolved in the deposited structure (per-atom resolved). We assessed calibration between each predicted and observed metric via Pearson *r* and Spearman ρ on the FoldBench PPI set, as well as DockQ stratified by ipTM bins (Figure S12). We observe strong correlation between the predicted and grouth truth metrics across both models.

**Figure S11.**
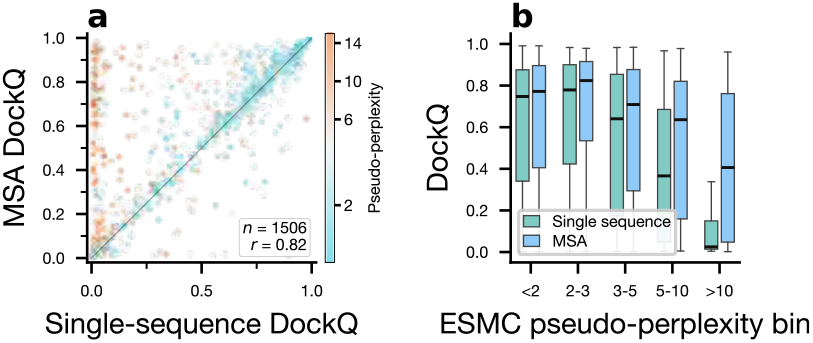
Effect of sequence predictability on ESMFold2 folding quality. (a) Single-sequence vs. MSA-conditioned DockQ per interface, colored by mean ESMC pseudo-perplexity of the interacting chains. Pearson correlation coefficient *r* = 0.82 (b) DockQ distributions binned by pseudo-perplexity, comparing single-sequence and MSA-conditioned regimes. MSA information contributes little at low PPL but markedly improves DockQ on harder, higher-PPL sequences.

#### A.2.11. Inference

##### Number of recurrence loops

Despite being trained on a maximum of 6 loops, we find that increasing the number of recurrent loops *T* at inference time sometimes improves results (A.2.5) Figure S13 sweeps *T* on FoldBench protein–protein (PPI) and antibody–antigen (AbAg) subsets at a fixed 68 diffusion sampling steps. ESMFold2 shows inference-time scaling out to 64 loops, particularly on AbAg complexes (Figure S13) On PPI complexes, scaling the number of loops shows a less significant improvement in the DockQ compared to AbAg complexes.

##### Optimal inference settings

Figure S13 shows inference parameters as a function of recurrent loops and diffusion steps. The optimal inference parameter settings depends on the prediction target, but performance is near-optimal at the default settings of ESMFold2 with 10 loops and 68 diffusion steps (truncated noise schedule described in Algorithm 4). However, to improve throughput and trade off accuracy for speed, the number of diffusion steps and recurrent loops can be reduced accordingly.

##### Mixed precision

Inference precision follows the training configuration. The language model ESMC is loaded and operates in float8. The trunk operates in bfloat16, as does the pair transition in the diffusion conditioning, while the diffusion module runs in float32 with TF32 enabled for matrix–matrix multiplications. For the confidence module, we restrict bfloat16 to the folding sub-trunk only; applying reduced precision to the full module introduces numerical instabilities.

##### Inference kernels

The pair folding layer is memory-bound: intermediate tensors are repeatedly round-tripped between High Bandwidth Memory (HBM) and on-chip SRAM through successive stages, leaving the tensor cores under-utilized. We implemented custom Triton kernels to remove these bottlenecks and fuse adjacent operations into a single launch, keeping intermediate results in registers and SRAM rather than HBM. Built on top of the Cuequivariance kernels (116), ESMFold2 kernels additionally fuse the residual addition and the row-shared dropout mask into the epilogue of the final gated GEMM — eliminating one full HBM write and one full HBM read of the trimul delta tensor. The same memory-bound pattern affects the attention with pair bias layer in the diffusion transformer. Here, a LayerNorm over the pair channel dimension, a linear projection to the per-head bias, and a key-mask addition are each executed as separate kernels — each round-tripping an intermediate tensor through HBM. A custom kernel collapses these three stages into a single Cooperative Thread Array, where the LayerNorm statistics, projection, and mask addition are all computed in registers, and only the final attention bias tensor is written to memory for scaled dot-product attention.

##### Compilation

ESMFold2 can be compiled with static compilation for specific sequence lengths. To achieve this, inputs are padded to a small set of fixed bucket sizes, which are then compiled.

##### Inference caching

In the computation of the diffusion transformer, multiple tensors are reused. These tensors can be cached to avoid re-computation (36). ESMFold2 caches the pair conditioning tensor as well as 3DRoPE tensors.

##### Inference speed vs. sequence length

Figure 2E reports inference throughput for ESMFold2 alongside four contemporary folding methods (Boltz1, Chai, Protenix-v1, AlphaFold3) as a function of sequence length, with and without MSA inputs. The benchmark uses randomized sequences of lengths 128, 256, 384, 512, 640, 768, 896, and 1024, and is run on a single H100 80 GB GPU. All methods are run with a common script using each project’s official released code. Note that AlphaFold3 is implemented in JAX, and the others in PyTorch. We also report folding time only; input and output processing, including MSA generation and processing, are not included. ESMFold2, ESMFold2-Fast, and AlphaFold3 pre-warm one compiled graph per padding bucket before any timed call, so reported times exclude compilation. For a fair comparison, all methods are run with 10 recycles and 200 inference steps.

**Figure S12.**
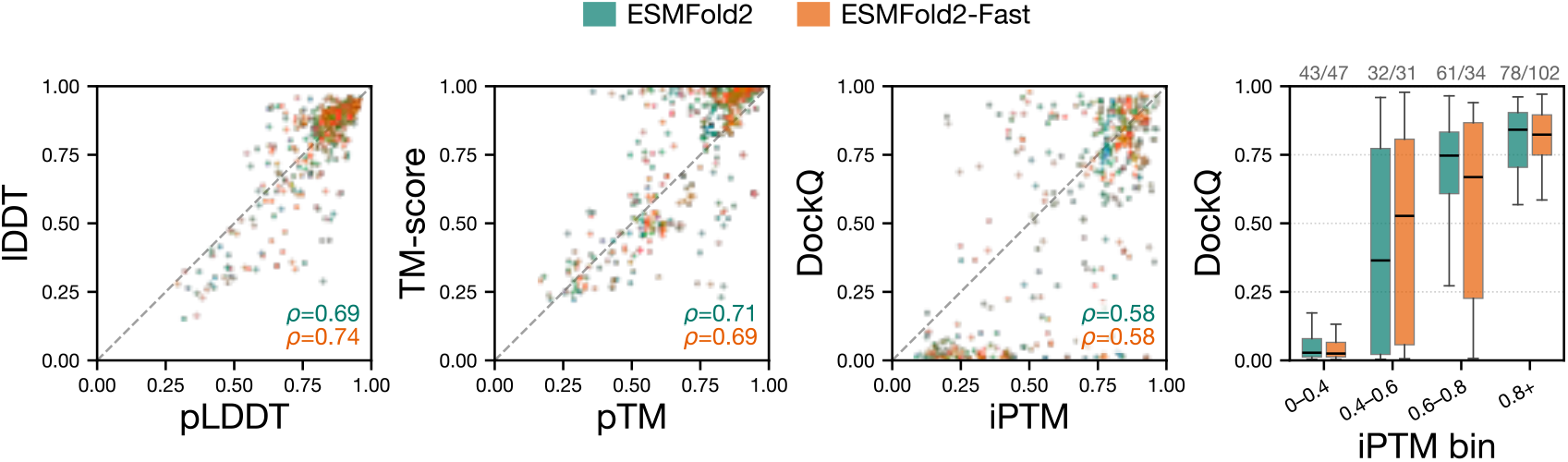
Calibration of confidence-head outputs on FoldBench PPI (*n* = 214) for ESMFold2 and ESMFold2-Fast. From left to right: ground-truth lDDT vs. predicted pLDDT; ground-truth TM-score vs. predicted pTM; ground-truth DockQ vs. predicted ipTM (each panel annotated with Pearson *r* and Spearman *ρ*); and DockQ distributions grouped by ipTM bin.

**Figure S13.**
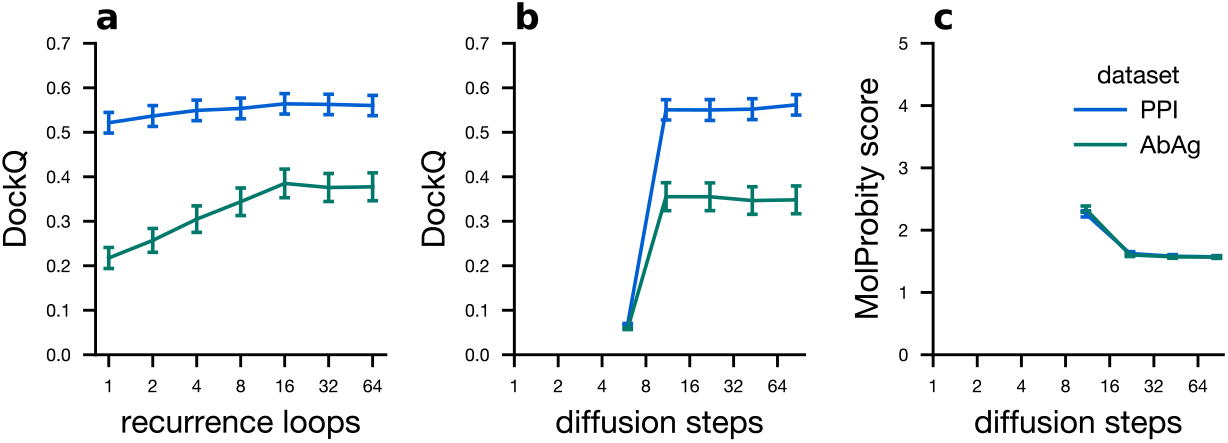
Analysis of inference hyperparameters on Foldbench antibody-antigen and protein-protein complexes. Mean score is shown over 5 seeds and 5 diffusion samples. Diffusion sampling does not converge below 11 steps. (a) DockQ score as a function of *T*, the number of inference loops, (b) DockQ score as a function of the number of diffusion sampling steps, and (c) MolProbity score (115), a metric of all-atom quality.

We observe that ESMFold2-Fast and ESMFold2 are the two fastest models, respectively. At sequence length 1024, ESMFold2 is 1.3× faster than AlphaFold3 and ESMFold2-Fast is 2.2× faster using the same inference settings. We note that in practice, ESMFold2 and ESMFold2-Fast are faster because we use fewer diffusion steps at 68 with no observable performance degradation.

##### Long context

Memory usage of ESMFold2 grows as *O*(*L*^2^), which limits the protein length that fits on a given GPU. To overcome this we use the context parallelism (CP) framework originally developed in ring attention (117) and FlashAttention (118), and specialized to protein folding components in Fold-CP (119). With these improvements, activation memory scales as *O*(*L*^2^*/g*), with *g* GPUs, enabling much longer contexts at both training and inference. While the base folding model is capped at sequence length 2k, application of fold-CP enables ESMFold2 to scale up to length 4k on 4 H100s and 6.5k on 16 H100s. Some qualitative examples of ESMFold2 on large complexes are given in (Figure S14).

#### A.2.12. Experimental Models

In addition to the final ESMFold2 models, several intermediate checkpoints are used as artifacts elsewhere in this paper:

- `ESMFold2-Experimental` and `ESMFold2-Experimental-cutoff2025` were used as a critic for protein design.
- `ESMFold2-Experimental-Fast` and `ESMFold2-Experimental-Fast-cutoff2025` were used both as critics and generators for protein design. Additionally, `ESMFold2-Experimental-Fast-cutoff2025` was used to fold ESM Atlas
- `ESMFold2-Experimental-Fast-baseXX` is a series of intermediate checkpoints saved at varying ESMC pre-training step counts, used both for scaling-law analysis and as critics for protein design.

**Figure S14.**
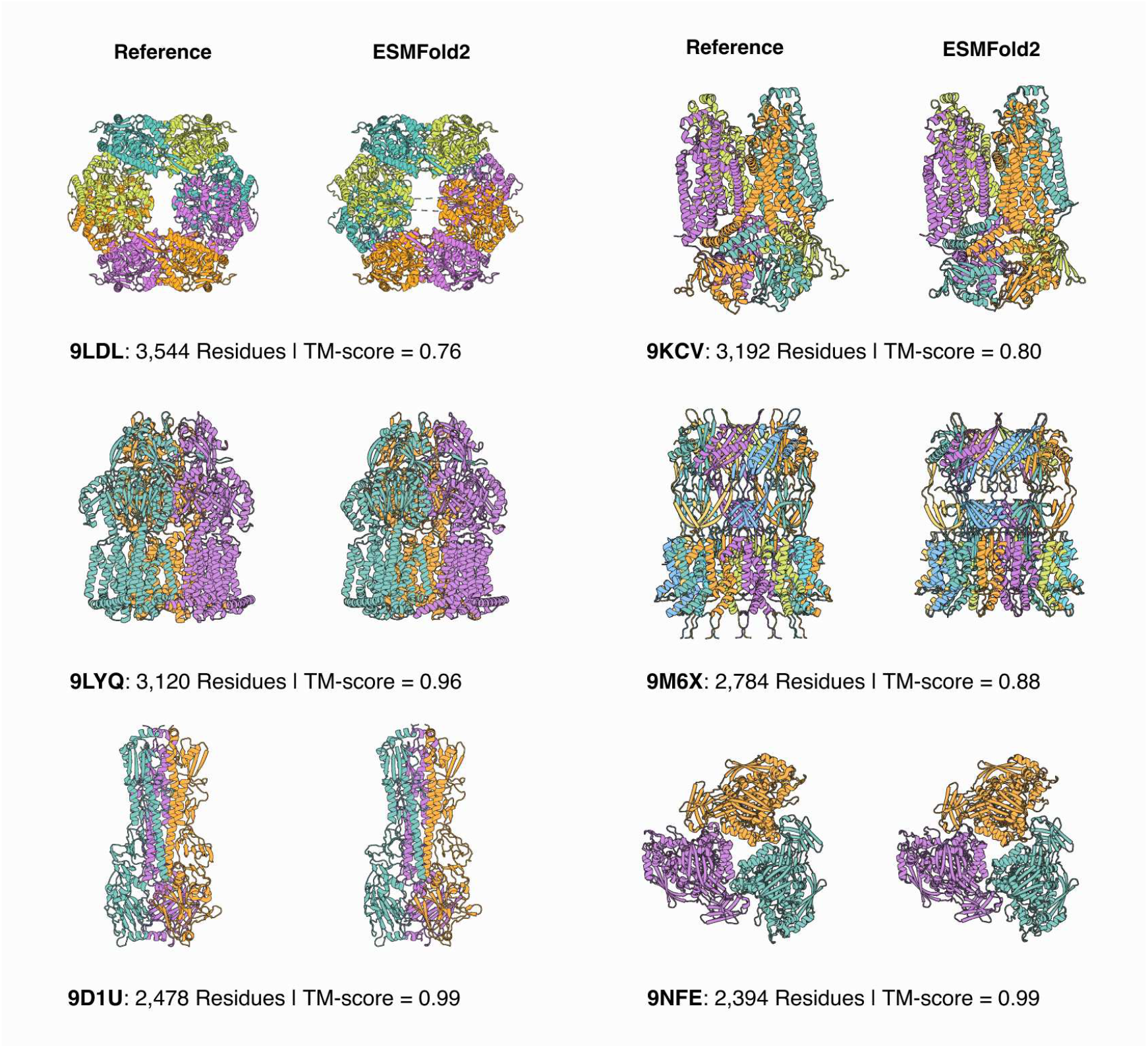
Long-context folding with ESMFold2. Using context parallelism on multiple GPUs, ESMFold2 can generalize beyond its training length to predict large biomolecular assemblies. Reference (left of each pair) and ESMFold2 prediction (right of each pair) are shown for several large complexes that lie outside the model’s training distribution (released after the training cutoff date; no chain in the complex clusters with any chain in the training set at 40% sequence identity). While four complexes (9D1U, 9NFE, 9LYQ, 9M6X) show highly accurate global folds (TM-score = 0.9-0.99), two complexes (9LDL, 9KCV) exhibit accurate local structure but large deviations in global domain arrangement (TM-score ≈ 0.7–0.8).

The main changes from the experimental models to the final ESMFold2 models are described below:

##### Model architecture

Multiple changes were made to the model during the development of ESMFold2. For the experimental models:

- Standard recycling is used. Stabilization is not applied to the recurrent update and gradients are not taken through the recurrence. The number of recycles during training is sampled between [0, 3].
- MSA encoder layers place `PairWeightedAveraging` before `OuterProductMean`. The final models reverse this.
- No folding layers are used to process ESMC inputs alone; the inputs are added directly to the pair state. The final models use 4 pair folding layers to process ESMC inputs.
- No `LayerNorm` is used prior to confidence output heads.
- Confidence heads are trained from stage 1 in the final models, supplanting a final confidence finetuning stage.
- Confidence output heads use LeCun initialization (120). In the final models, zero initialization is used.
- In the experimental models, 8 mini-rollout samples per target are used for confidence head training, increasing the effective batch size to 2048. We observed no benefit of increased diffusion samples, so the final training recipe uses one mini rollout sample.

##### Training recipe

The four experimental models (other than intermediate scaling checkpoints) are trained in four stages with progressively increasing context, plus an additional confidence-head training stage (Table S9). A batch size of 256 is used throughout. The learning rate follows a 1000-step warmup to 1.8 × 10^−3^ and is decayed to 1.8 × 10^−8^ across the final 10,000 steps (Stages 3–4).

For intermediate scaling checkpoints (`ESMFold2-Experimental-Fast-baseXX`), models were trained on the stage 1 training configuration with no MSA conditioning with 24 folding layers. The embeddings of ESMC were frozen similar to release models and folding models were fully trained out for each of the ESMC 300M, 600M, and 6B models at {250, 500, 750, 1000, 1500} thousand updates, for a total of 15 models.

**Figure S15.**
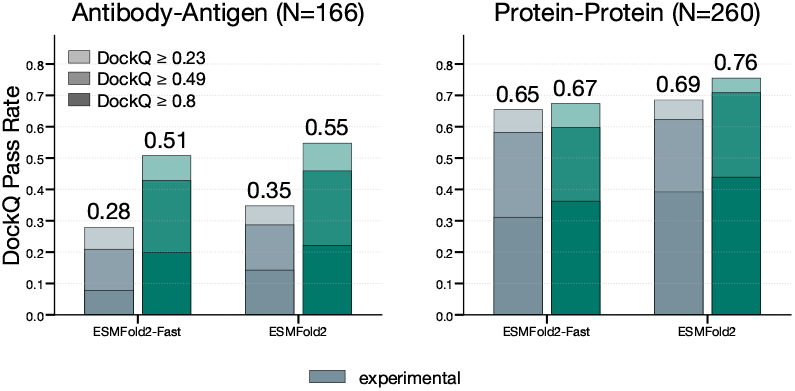
FoldBench antibody-antigen and protein-protein interface DockQ pass rates for experimental and release ESM-Fold2 models.

##### Training data

Experimental models are trained on the same data mixture as ESMFold2 and ESMFold2-Fast with one exception. For experimental models, antibody-antigen structures are sampled with higher probability. SAbDAb structures are clustered with 40% sequence identity and sampled with 1.8% probability at every training iteration. Probabilities of sampling PDB data and AlphaFoldDB (90) are adjusted to maintain the same relative ratio. No significant effect was observed from this upsampling, so for simplicity this was removed from the final training recipe.

##### Foldbench performance comparison

We compare the performance of experimental models and release models on FoldBench antibody-antigen and protein-protein interfaces Figure S15. The release models significantly outperform the experimental models across all metrics, almost doubling the DockQ pass rates on antibody-antigen interfaces.

#### A.3. Protein Design

ESMC and ESMFOLD2 define a joint model of protein sequence and structure that can be used to design new proteins that form interfaces with other proteins. A simple gradient-based optimization over the sequence prior *p*(sequence) from ESMC and the conditioned structural prior *p*(structure|sequence) from ESMFold2 is sufficient to produce high quality designs.

#### A.3.1. Design of Binders

##### A.3.1.1 Design Approach

Optimization is guided towards sequences predicted with high confidence to realize the desired interface. The end-to-end protocol is simple: generate a large number (thousands, or more) of candidate binders by searching the representations of the models, predict each in complex with the desired target using an ensemble of ESMFold2 models, and rank candidates by their average confidence over the predicted interface. The highest ranked designs are synthesized and characterized in the laboratory.

**Table S9.** Training stages for the ESMFold2 experimental models. The intermediate checkpoints for scaling-law analysis were only trained on Stage 1. SAbDab sampling is not used in released ESMFold models or intermediate scaling checkpoints.

|  | Stage 1 | Stage 2 | Stage 3 | Stage 4 | Stage 5 |
| --- | --- | --- | --- | --- | --- |
| Steps | 80,000 | 10,000 | 5,000 | 5,000 | 10,000 |
| Max tokens | 256 | 512 | 640 | 768 | 640 |
| Max atoms | 2,304 | 4,608 | 5,760 | 6,912 | 5,760 |
| Max MSA depth | 4,096 | 4,096 | 3,712 | 1,536 | 2,048 |
| Diffusion batch size | 16 | 16 | 16 | 12 | 8 |
| Smooth LDDT loss | ✓ | ✓ | ✗ | ✗ | ✗ |
| Train structure prediction | ✓ | ✓ | ✓ | ✓ | ✗ |
| Train confidence head | ✗ | ✗ | ✗ | ✗ | ✓ |
| PDB40 proportion | 28.2% | 28.2% | 28.2% | 47% | 90% |
| SAbDab proportion | 1.8% | 1.8% | 1.8% | 3% | 10% |
| AFDB proportion | 70% | 70% | 70% | 50% | 0% |

###### Search

Gradients of ESMC and ESMFold2 models are used as differential search objectives as described by Algorithm 11. Our method utilizes gradients directly optimized through the structure model’s distogram output, a method that aligns with a broader class of approaches inverting structure prediction models for the design of binders (41– 44).

A single search for a binding candidate begins by initializing a state vector *x* ∈ *L*_binder_ × 20 tensor where *L*_binder_ represents the amino acid positions of the binder chain. In cases where the identity of the amino acid is known, such as in the framework positions of an antibody, the position is initialized with logits which approach a one-hot vector after softmax. In the positions where the amino acid is to be chosen, it is initialized with 𝒩 (0, 0.01^2^).

These positions are updated iteratively with gradient descent. One update adjusts the state vector according to the gradient of the losses (which guide the design towards a binding protein) adjusted by a step-dependent learning rate which is annealed over the 150 steps of the search.

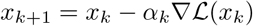

At the beginning of the search the design state is a superposition over amino acids, which corresponds to a latent representation of the model. Cysteine amino acids can be undesirable as they often form unwanted disulfide bonds. The design of cysteines is avoided by initializing their logits at −10^6^ and zeroing gradients during backpropagation.

The overall loss ℒ is a combination of four losses that utilize ESMC and ESMFold2. Three losses are computed from the ESMFold2 distogram logits. The first minimizes the entropy of predicted tertiary contacts within the binder sequence (Algorithm 12). The second minimizes the entropy of predicted cross-chain contacts between the binder sequence and target. The third penalizes the predicted packing radius of the binder (Algorithm 13). A fourth loss, computed from ESMC, minimizes the cross entropy between the binder sequence state and log probabilities output by the model (Algorithm 14). There are step-dependent transformations between the state and the inputs to the models implementing the losses, which are described in Algorithm 11.

The ESMFold2-derived losses guide optimization towards sequences that are predicted to fold with high confidence (i.e., low entropy of the distogram logits) into globular structures that engage the target. The ESMC-derived loss guides optimization towards sequences that are likely under the language model’s evolutionary prior. These losses may be implemented using multiple ESMFold2 variants to prevent adversarial solutions to a particular set of model weights.

In the final few optimization steps of a design trajectory, a forward pass through the diffusion and confidence modules is run to compute structure confidence (ipTM). The final output of a trajectory is the argmax over the amino acid logits of the state corresponding to the highest ipTM design.

This search algorithm is applied many times in parallel to generate a large set of candidate designs. Due to its embarrassingly parallel structure, this can be approached in various ways on various types of infrastructure to achieve a high level of scale.

##### A.3.1.2 Selection

Not all outputs of the search yield good candidate designs. A critical aspect of achieving high hit rates is selecting the best designs from the set of candidates. Our goal is to choose candidates for ∼1 96 plate well for evaluation out of the ∼1000’s of candidates.

Minibinder designs with an isoelectric point (pI) above 6.0, which can lead to nonspecific binding and aggregation (for near-neutral pIs), are first filtered. For scFvs, no filters are applied.

Then candidates are ranked using ESMFold2, specifically using the predicted confidence of the interface scored by ipTM or a proxy for ipTM related to the entropy of the distogram logits (Algorithm 15). One or more ESMFold2 models with different weights may be used to generate a combined score. In practice, multiple models are used and a weighted mean of scores is used for the ranking. Designs ranked the highest are selected for laboratory experiments.

#### A.3.2. Description of Experiments

##### A.3.2.1 Design Task Specification

###### Target Preparation

Binder design was performed against five targets: EGFR, PD-L1, PDGFR*β*, CTLA-4, and CD45. For each target, the amino acid sequence used in the search protocol was a crop of the full target. The sequences of the target crops and their sources are in Table S10.

###### Minibinder Design

Minibinder sequence lengths are sampled uniformly from {60, 61, …, 200} for all targets.

###### scFv Design

To generate scFvs, we utilized the framework regions of three therapeutic antibodies: trastuzumab, atezolizumab, and ocankitug. Notably, to minimize framework bias toward its native antigen, the atezolizumab scaffold was explicitly excluded from designs targeting PD-L1. Specifically, the frameworks derived from the following antibodies were used for each target: trastuzumab and ocankitug for PD-L1, trastuzumab and atezolizumab for EGFR, atezolizumab and ocankitug for CTLA-4, trastuzumab and atezolizumab for PDGFR*β*, and trastuzumab and atezolizumab for CD45.

CDRs were defined according to the Chothia numbering scheme (121). All constructs were designed in a VH-VL orientation, with the heavy and light chains connected by a (Gly-Gly-Gly-Ser)4 linker.

During generation, the lengths of the six CDRs are sampled independently from a uniform distribution. Lengths are fixed for LCDR2 (7) and LCDR3 (9) and varied for LCDR1(11–16), HCDR1 (7–9), HCDR2 (5–6), and HCDR3 (9–15 for the trastuzumab-derived and atezolizumab-derived frameworks, and 8–14 for the ocankitug-derived framework).

#### A.3.2.2 Design Experiments

We ran experiments at two scales of inference compute. Both experiments use the same targets, frameworks, and design task specifications. For each target, design modality, and scale of inference compute, 84 designs were selected for wetlab validation. For scFv designs, the selection is split evenly between the two frameworks used for design (i.e., the top 42 ranked designs were selected for each).

The two experiments differ in the amount of compute applied in candidate generation and ranking. For minibinders, the smaller and larger sets contained ∼ 15,000 and ∼ 67,000 candidates on average, requiring ∼ 500 and ∼ 2,400 H100-hours, respectively. For scFvs, the sets contained ∼ 28,000 and ∼ 117,000 candidates on average, requiring ∼ 1,800 and ∼ 7,700 H100-hours, respectively. Figure S16 provides a breakdown of expended compute over individual binder-modality pairs.

For both experiments, the following models are used in search: ESMFold2-Experimental-Fast-cutoff2021 and ESMFold2-Experimental-Fast-cutoff2025. ESMC 6B is always used for the language model term.

Both experiments use the following four models in the ensemble for ranking: ESMFold2-Experimental-Fast-cutoff2021, ESMFold2-Experimental-Fast-cutoff2025, ESMFold2-Experimental-cutoff2021, and ESMFold2-Experimental-cutoff2025, which includes the two models used in design.

In the larger compute experiment, the ensemble of ranking models was expanded from 4 to 19. Specifically, we include 15 models trained across ESMC checkpoints with varying levels of compute (Section A.2.12). The ranking score was aggregated first within groups of models with the ipTM head and with the distogram-derived confidence proxy separately (15), and then those two scores are averaged together. All folding models were run with 3 loops and 200 diffusion sampling steps.

#### A.3.3. Experimental Validation

##### A.3.3.1 Gene fragment design for affinity screening

Designed amino acid sequences were reverse translated to DNA sequences by a codon optimization algorithm for expression by *E. coli* translational machinery. The algorithm removes rare *E. coli* codons. All sequences included a T7 promoter and ribosome binding site upstream of the gene, a start codon and glycine at the *N* terminus, a *C*-terminal Twin-Strep-tag, and a T7 terminator downstream.

**Figure S16.**
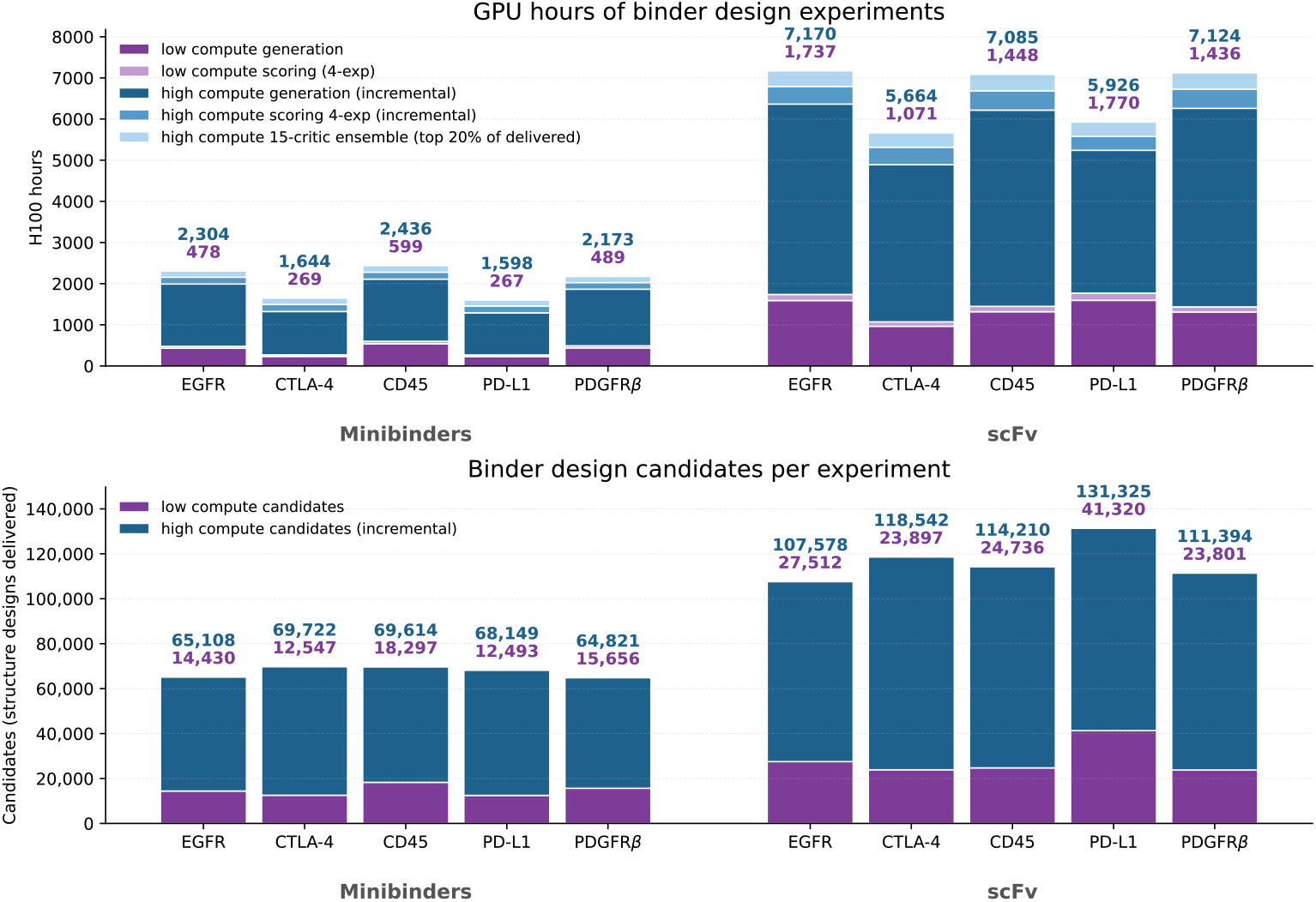
GPU hours and binder design candidates across low and high compute experiments. **Top:** H100 GPU hours allocated to each binder design experiment, decomposed by stage: low-compute generation and scoring (4-model ipTM ensemble), high-compute generation and scoring (incremental over low compute), and the 15-checkpoint distogram binder confidence ensemble applied to the top 20% of delivered designs by the 4-model ipTM ensemble. scFv pools require roughly 3× the compute of minibinders at matched targets, driven by the larger structures folded during generation and increased number of designs. **Bottom:** Number of structure designs generated for low compute and high compute experiments. The low compute design pool is a subset of the high compute pool.

###### Algorithm 11

Gradient-Guided Binder Sequence Optimization.

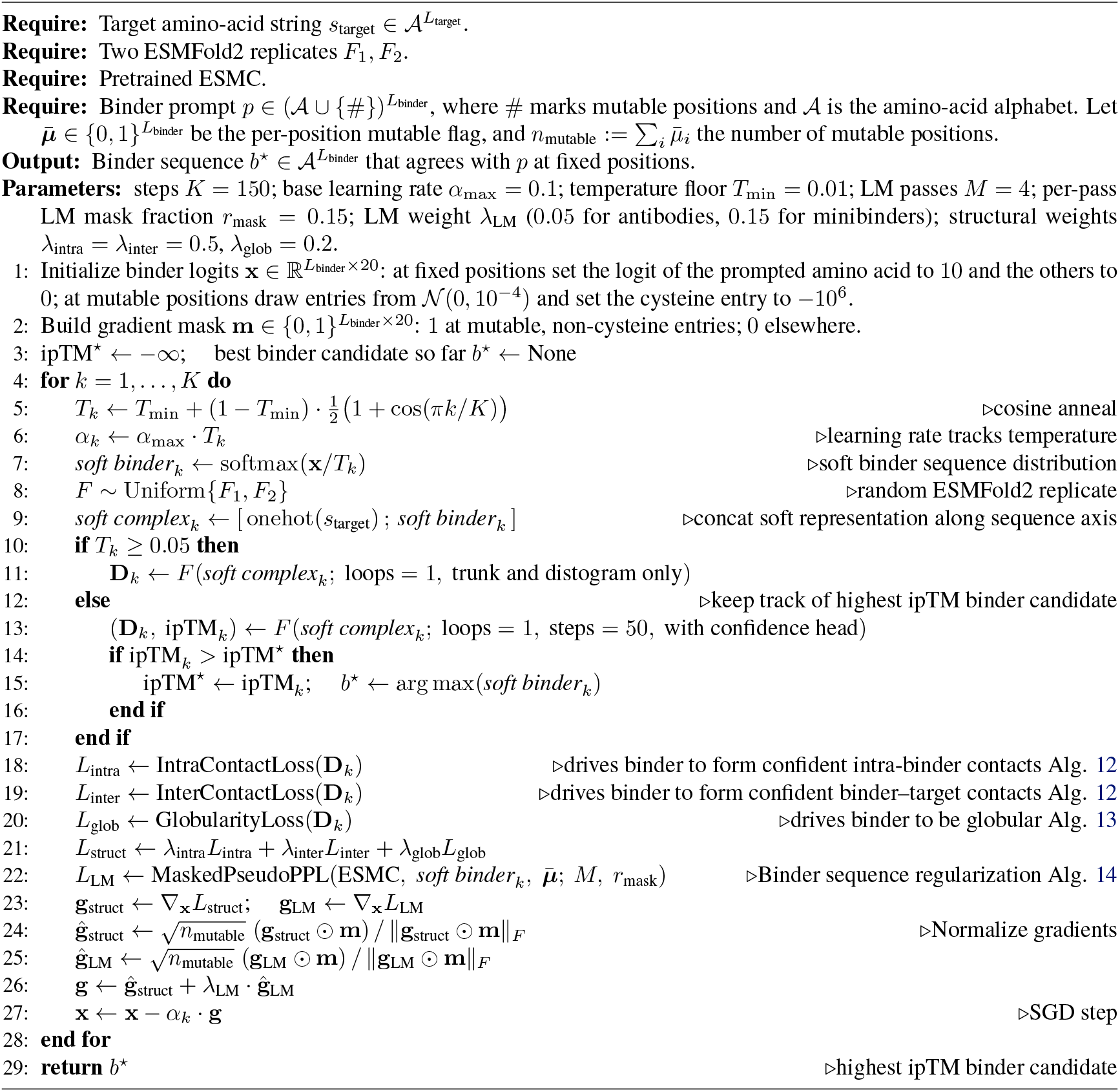

###### Algorithm 12

Contact Losses.

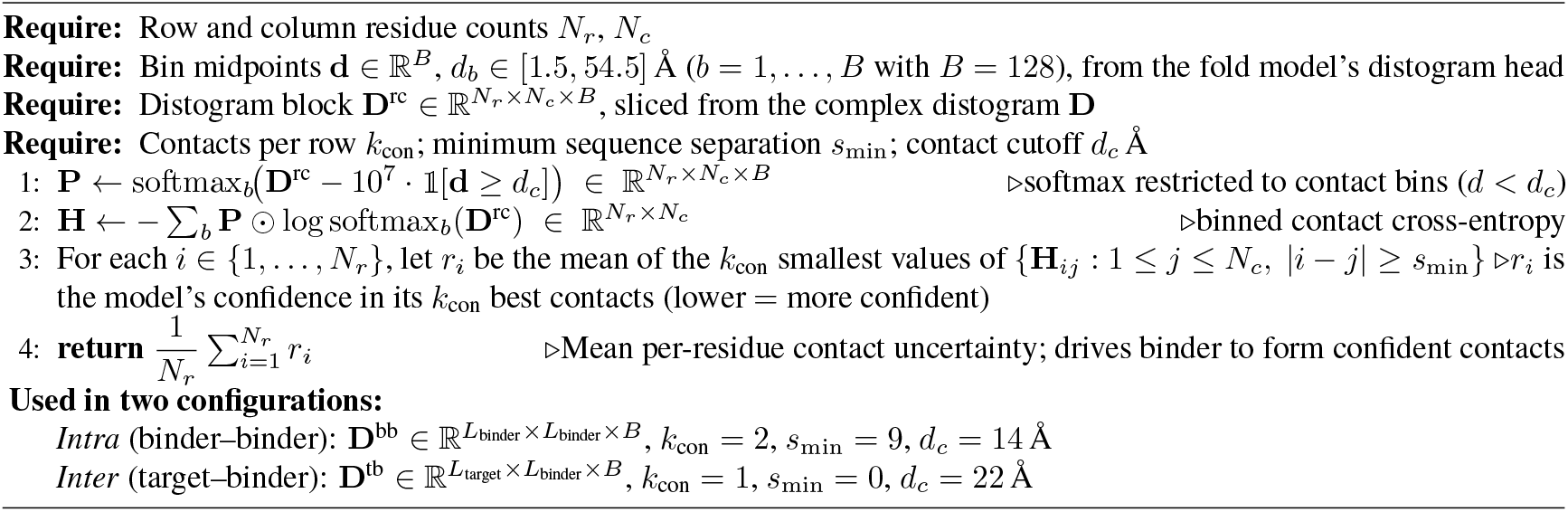

###### Algorithm 13

Globularity Loss.

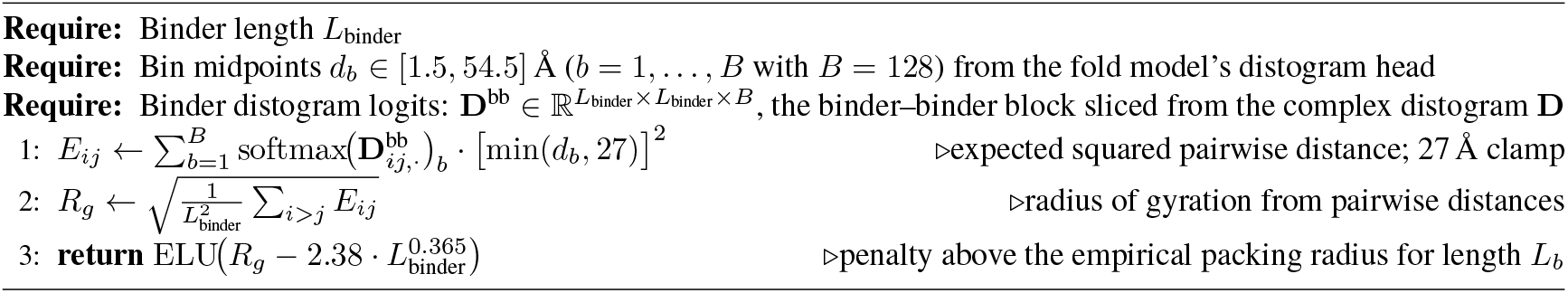

###### Algorithm 14

ESMC Pseudo-perplexity Sequence Regularization.

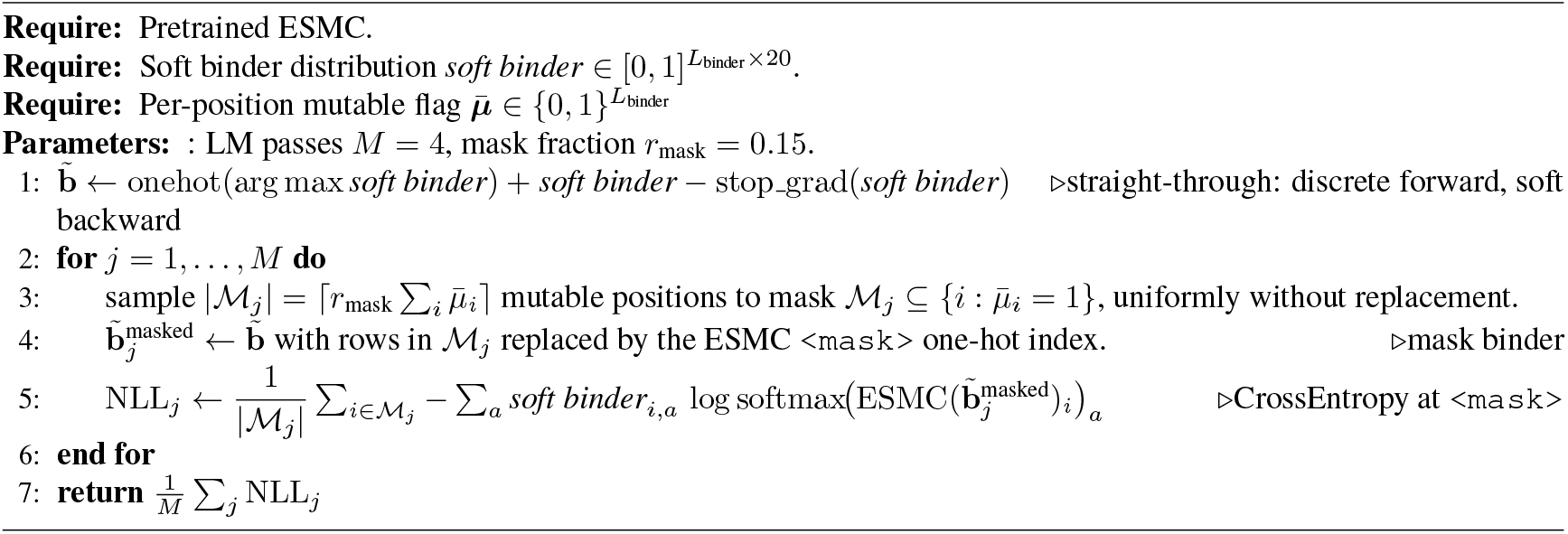

**Table S10.**
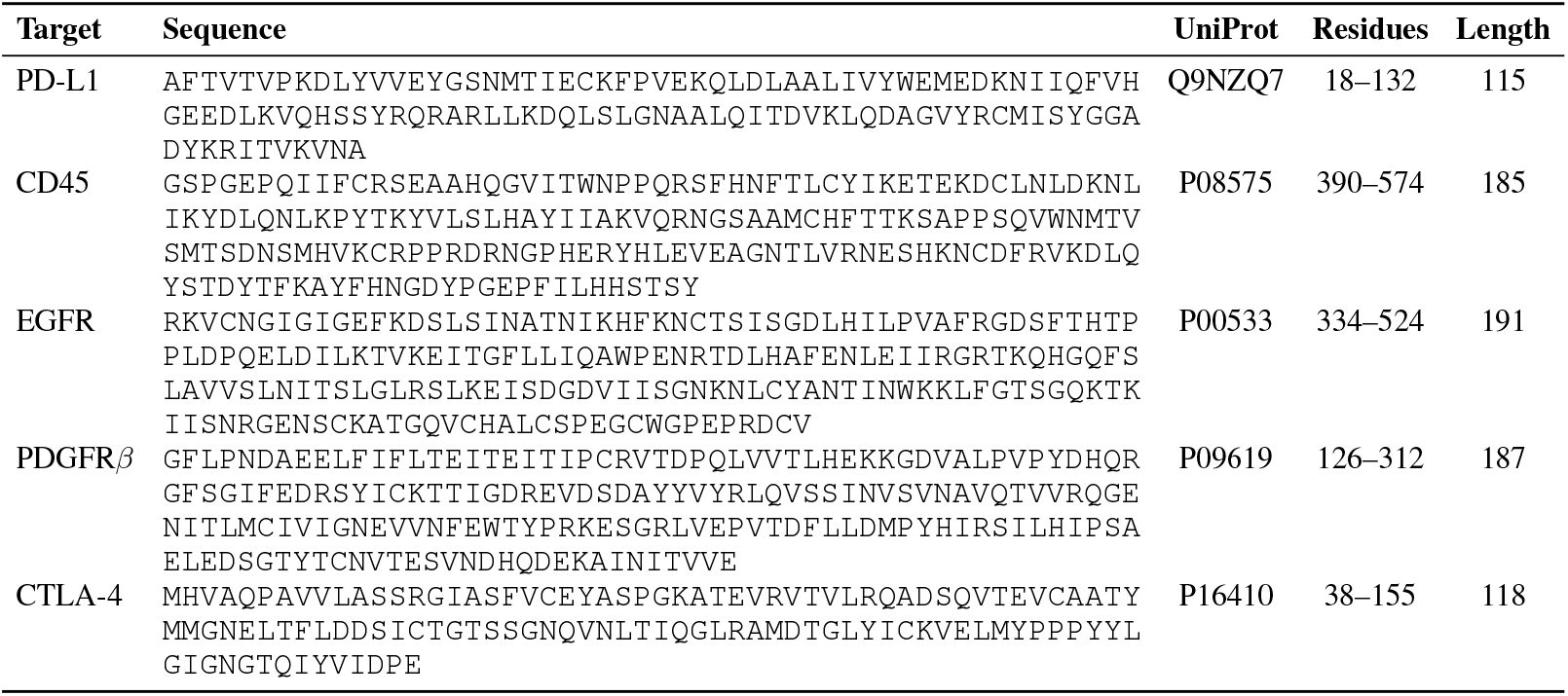
Target protein sequence crops used for binder design experiments.

###### Algorithm 15

Distogram ipTM Proxy.

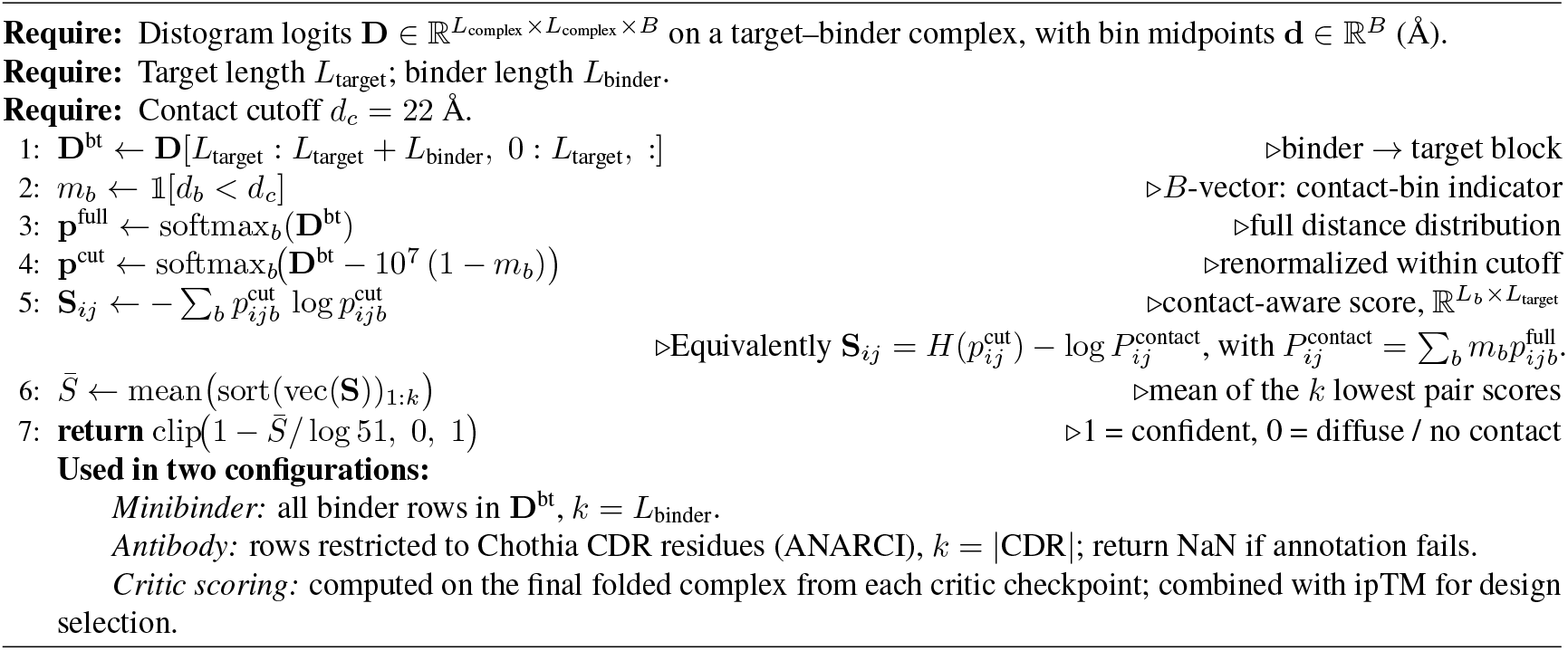

##### A.3.3.2 Minibinder and scFv expression by *in vitro* translation for antigen binding screening

Binder sequences were synthesized as linear gene fragments by Twist Bioscience. Proteins were expressed using the PURExpress *In Vitro* protein synthesis kit (New England Biolabs, NEB) in 8 *µ*L reactions according to the manufacturer’s suggested protocol. Each reaction contained 32 ng of DNA and 6.4 units of murine RNase inhibitor (NEB). For scFv constructs, 0.32 *µ*L each of PURExpress Disulfide Bond Enhancers 1 and 2 (NEB) were added to promote proper folding. Six positive and six negative control gene fragments per 96-well plate were interspersed throughout the screening plates and expressed under identical conditions (see details in Table S11). Reactions were incubated at 37^°^C for 3 hrs and subsequently stored on ice or at −20^°^C until use.

##### A.3.3.3 Binder screening by biolayer interferometry

Minibinder and scFv designs were screened for target binding by biolayer interferometry using a Gator Plus instrument (Gator Bio), as shown in Figure S17. For each design, 5 *µ*L of the *in vitro* protein synthesis reaction was diluted into 95 *µ*L K buffer (Gator Bio) and loaded onto Strep-Tactin XT probes (Gator Bio) for 120 sec to capture the Twin-Strep-tagged designs. Following loading, probes were blocked with 1 *µ*M Twin-Strep peptide (GenScript) for 30 sec to minimize nonspecific target interactions with the sensor surface. To assess nonspecific off-target binding, the immobilized designs were first dipped into 2 *µ*M ovalbumin (MP Biomedicals) for 120 sec, followed by a 120 sec dissociation phase. The immobilized designs were then screened against their respective target antigens at a concentration of 2 *µ*M (120 sec association, 120 sec dissociation). The firastotug scFv positive control on the inference-scaled CTLA-4 minibinder screen failed to bind due to the accidental omission of the disulfide bond enhancer during binder expression expression. Recombinant antigens were sourced as follows: EGFRvIII-His (Acro Biosystems EGI-H52H4), active CTLA-4-His dimer (Acro Biosystems CT4-H52H9), CD45-Llama IgG2b Fc (Acro Biosystems CD5-H5259), PD-L1-His (Acro Biosystems PD1-H5229), and PDGFRβ-His (R&D Systems 10676-PR). The EGFRvIII mutant was specifically selected over the full-length EGFR ectodomain due to its therapeutic relevance as an oncology target.

##### A.3.3.4 Screening analysis and hit characterization

Individual association and dissociation curves were aligned to the start of the association phase, inter-step corrected, and smoothed using a Savitzky-Golay filter within the Gator software. Processed data were exported to GraphPad Prism software (version 11.0.0, GraphPad Software, Boston, MA) for subsequent analysis. To account for assay drift, baseline correction was performed by subtracting the sensorgram of the nearest negative control binder from each designed binder’s sensorgram. These baseline-corrected sensorgrams were then fit using GraphPad Prism’s ‘association then dissociation’ model to estimate apparent affinities, which informed our selection of candidates for follow-up experiments. Two distinct analyses were performed to identify successful binders: an “inclusive” analysis consistent with recent literature, and a “stringent” analysis designed to isolate high-confidence candidates. Both methods required a minimum probe loading signal of *>*0.5 nm and a sensorgram line shape exhibiting positive exponential association. The inclusive analysis defined a hit using a universal binding signal cutoff of *>*0.1 nm above baseline. Conversely, the stringent analysis applied target-specific thresholds: EGFR (0.25 nm), CTLA-4 (0.25 nm), CD45 (0.4 nm), PD-L1 (0.25 nm), and PDGFR*β* (0.4 nm). These target-specific cutoffs were empirically established based on sensorgram line shape to exclude nonspecific binding and baseline artifacts. Furthermore, borderline hits passing these stringent thresholds were manually inspected to verify expected kinetic behavior and exclude atypical curvature.

**Figure S17.**
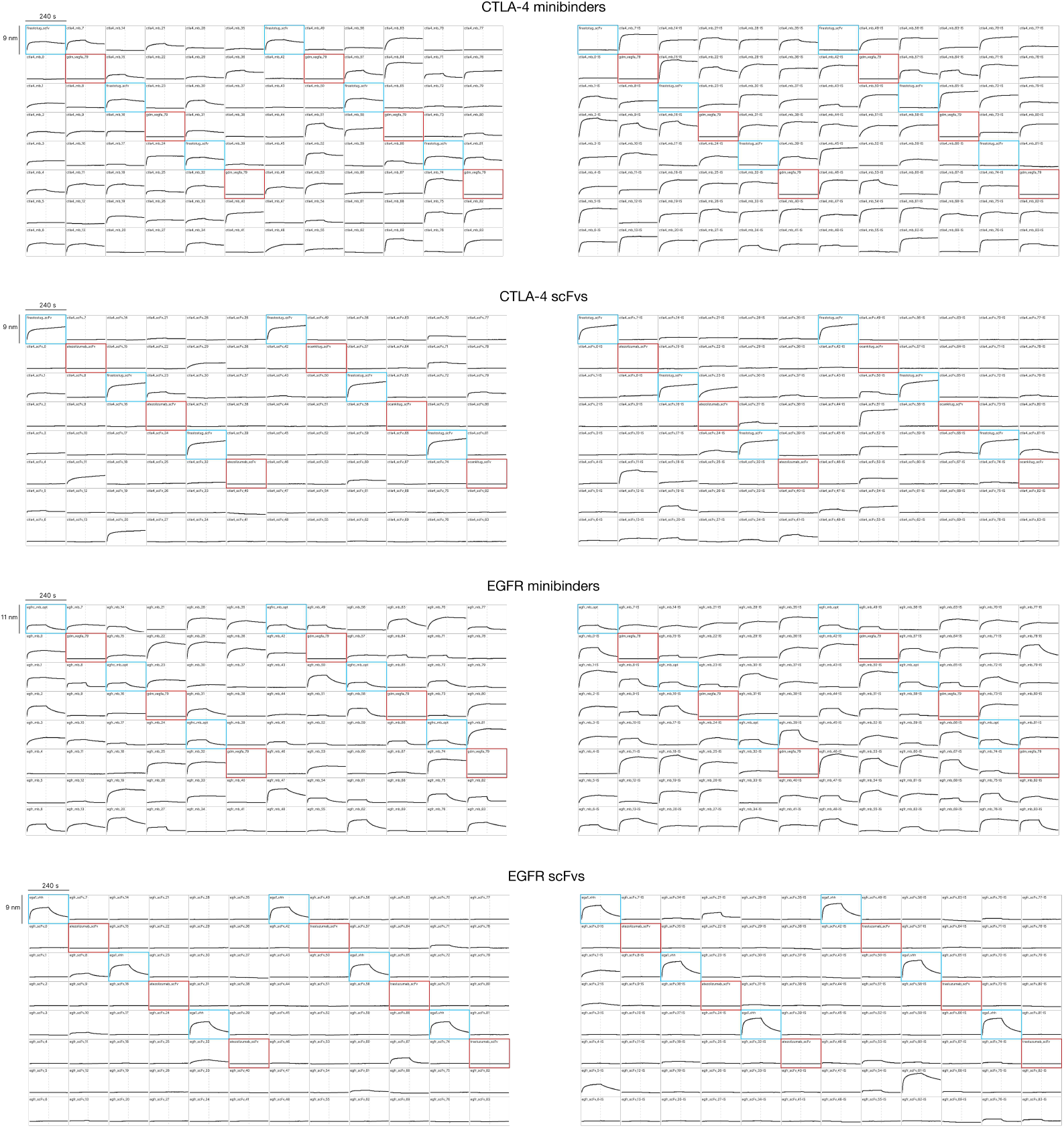
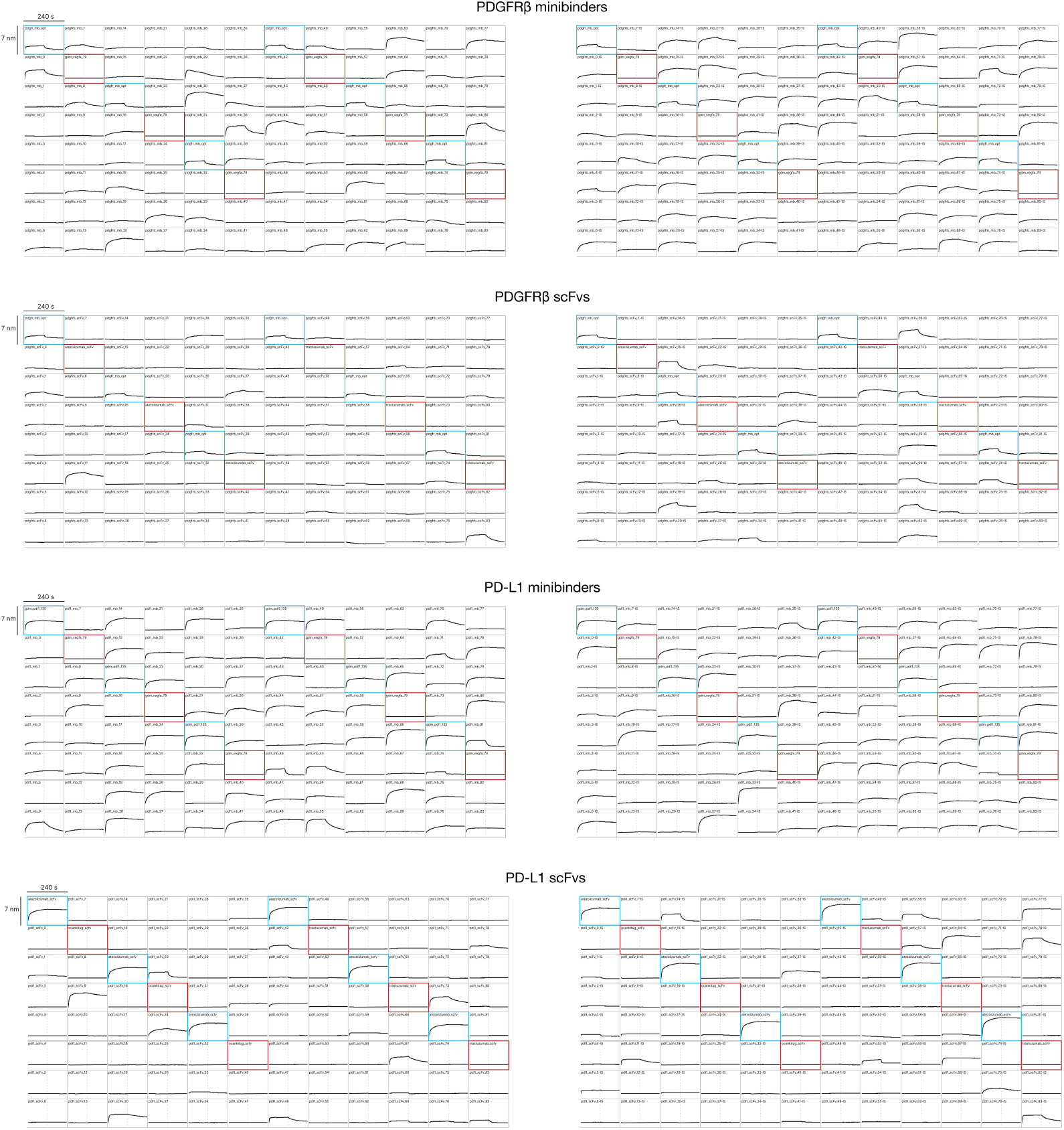
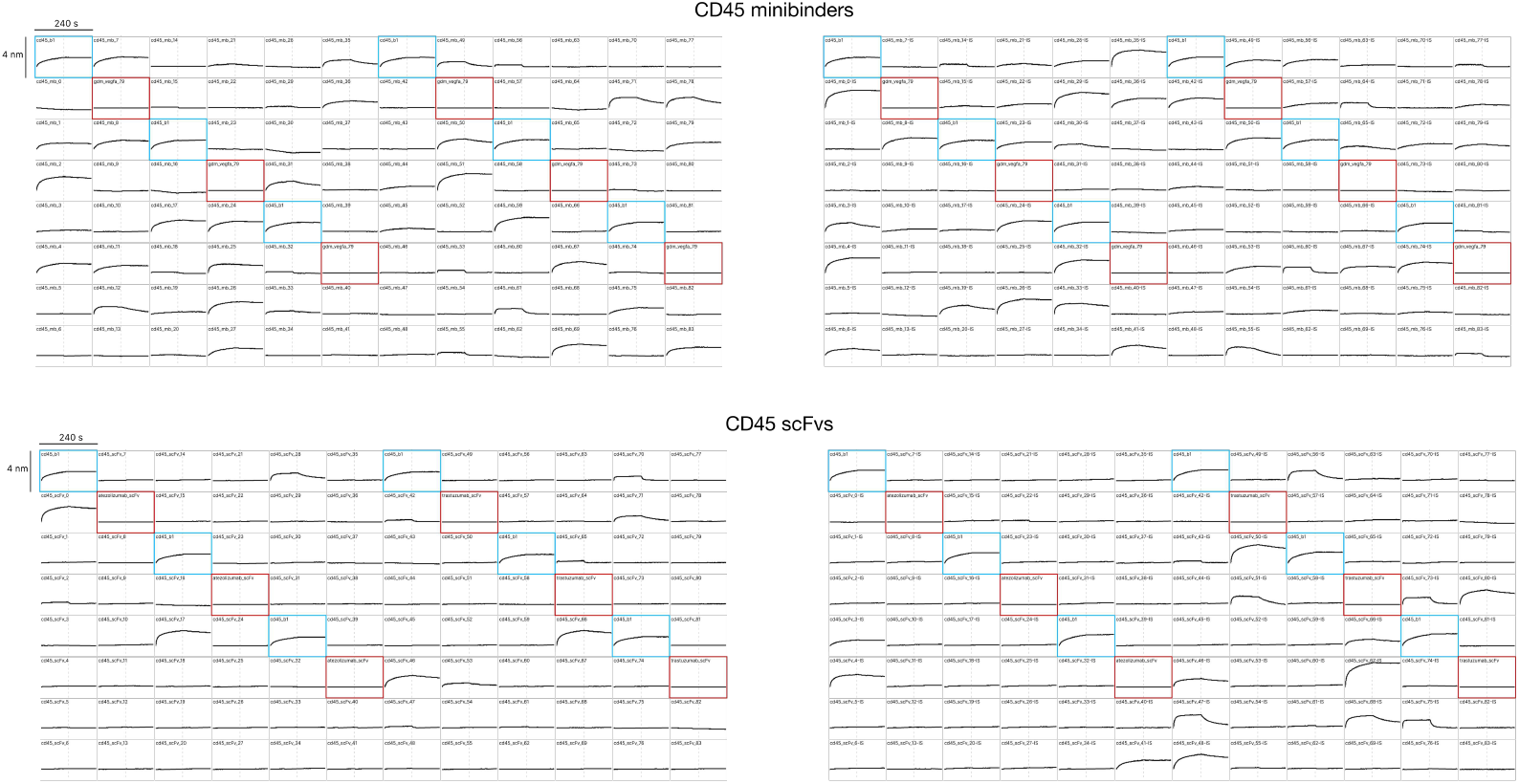
Biolayer interferometry (BLI) screening of minibinder and scFv candidate designs. Candidate designs were immobilized on Strep-Tactin XT probes and screened for binding against their respective target antigens at a concentration of 2 *µ*M, with 120 sec association and 120 sec dissociation phases. Baseline-corrected sensorgrams are displayed in a 96-well format, with low inference compute plates in the left column and high inference compute plates in the right column. “IS” in the binder name indicates that the binder was selected from the inference-scaled pool of designs. Positive controls are denoted by blue boxes and negative controls are denoted by red boxes. Details of the controls are listed in Table S11. Vertical dashed lines indicate the transition from the association phase to the dissociation phase. Scale bars for time (s) and binding response (nm) are indicated in the top left corner of each pair of plates.

**Table S11.**
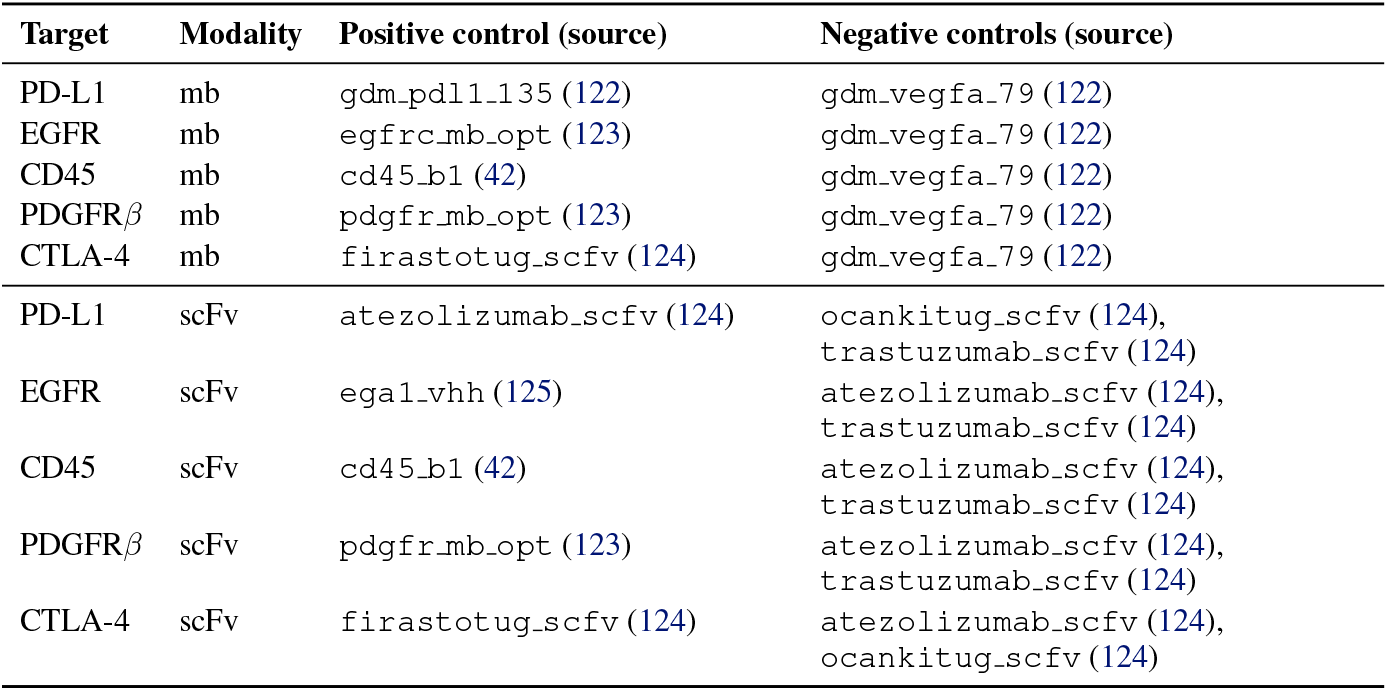
Positive and negative controls used for BLI screening. Positive and negative controls used in the BLI screening plates shown in Figure S17.

##### A.3.3.5 *In vitro* TRANSLATION & AFFINITY PURIFICATION OF BINDERS FOR BIOCHEMICAL CHARACTERIZATION

For downstream epitope competition and PD-L1/PD-1 blockade bioassays, designed binders and atezolizumab scFv (sequence obtained from TheraSabDab (124) and expressed in VH-VL orientation with a *C*-terminal Twin-Streptag) were expressed using the PURExpress *in vitro* protein synthesis kit (NEB) in scaled-up 500 *µ*L reactions according to the manufacturer’s protocol. We do not believe that there is any functional difference between binders purified from *in vitro* translation versus *E. coli* expression and generally defaulted to *E. coli*-derived proteins for biochemical characterization. In the epitope competition and blockade assays, proteins from *in vitro* translation were used in certain instances when the *E. coli* material was insufficient in amount.

Binder gene fragments (Twist Bioscience), described above, were PCR-amplified, purified with the Monarch spin PCR & DNA cleanup kit (NEB), and 2,000 ng of the resulting DNA was used as the template for each reaction. All reactions were supplemented with 400 units of murine RNase inhibitor (NEB). For scFv constructs, 20 *µ*L each of PURExpress Disulfide Bond Enhancers 1 and 2 (NEB) were added to promote proper folding. The translation reactions were incubated at 37^◦^C for 3 hrs with orbital shaking at 300 rpm on an Eppendorf Thermomixer, and subsequently stored at 4^◦^C prior to purification.

To purify the *in vitro* translated binders, 100 *µ*L of Strep-TactinXT 4Flow resin (IBA Lifesciences) slurry was added to each translation reaction and the mixture incubated at room temperature for 30 min with gentle agitation. The mixtures were then loaded onto disposable 10 mL polypropylene columns (Thermo Fisher). Liquid was drained by gravity flow and the captured resin in each sample was washed with 10 mL of high-salt phosphate buffered saline (standard PBS recipe, pH 7.4, supplemented with additional NaCl to a final concentration of 300 mM). Each protein was then eluted with 2 mL PBS containing 50 mM biotin. Biotin was removed by passing the eluate through 5 mL Zeba 7 kDa molecular weight cut-off spin desalting columns (Thermo Fisher) pre-equilibrated with PBS. Proteins were then concentrated by spin filtration through a 3 kDa molecular weight cut-off spin filter (Thermo Fisher) at 4,000 × g for 40 min at 4^◦^C. Resulting protein solutions ranged from 500-800 *µ*L in volume. Protein concentration was measured using the BCA protein assay (Thermo Fisher). Purity was confirmed by SDS-PAGE.

##### A.3.3.6 *E. coli* EXPRESSION PLASMID DESIGN AND CONSTRUCTION

To generate constructs for *E. coli* expression, binder sequences were amplified by polymerase chain reaction from the gene fragments above and cloned into the pET28a expression vector by Gibson assembly (NEB). For scFv constructs, sequences were preceded at the *N*-terminus by a pelB signal sequence for periplasmic secretion. All binders were appended with a *C*-terminal PreScission Protease cleavage site followed by a Twin-Strep-tag.

##### A.3.3.7 Minibinder and scFv expression and purification from *E. coli* FOR BINDER CHARACTERIZATION

Chemically competent BL21(DE3) *E. coli* (NEB) were transformed with the pET28a binder constructs and grown overnight at 37^◦^C on LB-agar plates with 50 *µ*g/mL kanamycin (Teknova). Overnight LB starter cultures with 50 *µ*g/mL kanamycin were inoculated from single colonies and further expanded into 0.5-1 L of Studier ZYM-5052 auto-induction medium (Teknova) in 2.5 L UltraYield flasks (Thomson Instrument Company). The cultures were grown at 37^◦^C for 4 hrs at 240 rpm, after which the temperature was lowered to 17^◦^C for 16-20 hrs. The bacteria was harvested by centrifugation at 6,000 × g and used immediately for purification, or stored at −80^◦^C for future use.

The *E. coli* pellets were lysed using a detergent-based extraction reagent for all minibinders as well as EGFR-scFv-61-IS, PDGFRβ-scFv-11, and PDL1-scFv-9, or by periplasmic lysis and osmotic shock for CTLA4-scFv-20 and CD45-scFv-0. Detergent-based extraction was performed with B-PER protein extraction reagent (Thermo Fisher) supplemented with NaCl to 150 mM, EDTA to 1 mM, and 1x cOmplete protease inhibitor cocktail tablets (Roche) or 1x HALT protease inhibitor cocktail (Thermo Fisher). Pellets were resuspended in 5 mL lysis buffer per gram of pellet and rotated for one hour at room temperature to lyse. Periplasmic lysis was performed as previously described (126), pellets were resuspended in 5 mL SET buffer (500 mM sucrose, 0.5 mM EDTA, 150 mM Tris-HCl pH 8) per gram of pellet and agitated in an orbital shaker at 100 rpm at 4^◦^C. After 30 min, 12 mL of water per gram of pellet was added to osmotically shock the cells. After 45 min of shaking, NaCl was added to a concentration of 150 mM.

The lysates were clarified by centrifugation at 15,000 × g for 30 min and vacuum-filtered with a 0.22 *µ*m bottle-top filter. StrepTrap XT pre-packed chromatography columns (Cytiva) were pre-equilibrated with binding buffer consisting of 100 mM Tris-HCl, 300 mM NaCl, and 1 mM EDTA, pH 8.0, then loaded with the clarified, filtered lysate at 1 mL/min using an automated protein purification system (127). The columns were washed with 70 mL of binding buffer and eluted with elution buffer consisting of 100 mM Tris-HCl, 150 mM NaCl, 1 mM EDTA and 50 mM biotin, pH 8.0.

The affinity-purified elution fractions were analyzed by SDS-PAGE, pooled, and spin-concentrated using 10 kDa molecular weight cut-off spin concentrators (Pierce). The binders were further polished by size exclusion chromatography (SEC) using an AKTA pure system (Cytiva) in phosphate buffered saline, pH 7.4, and a Superdex 75 Increase 10/300 column for the minibinders or a Superdex 200 Increase 10/300 column for the scFvs (Cytiva). The SEC elution fractions were analyzed by SDS-PAGE, and pure fractions were pooled and stored at 4^◦^C for less than 2 weeks for further use. The protein concentration was estimated based on absorbance at 280 nm, which was measured using a NanoDrop spectrophotometer (Thermo Fisher).

##### A3.3.8 Analytical size exclusion chromatography (SEC)

Binders purified from *E. coli* by affinity chromatography followed by preparative SEC were re-injected onto a Superdex 75 10/300 Increase column using an ALIAS Bio autosampler connected to an AKTA pure chromatography system (Cytiva). The column was run at room temperature in PBS, pH 7.4 at a flow rate of 0.8 mL/min, and absorbance at 280 nm was monitored. Binder concentrations at injection ranged from 0.1–0.25 mg/mL, with 5–20 *µ*g of protein injected per sample. Gel filtration standards (Bio-Rad) were run alongside samples as retention volume references. SEC traces were normalized to percent maximum signal and plotted in GraphPad Prism.

**Figure S18.**
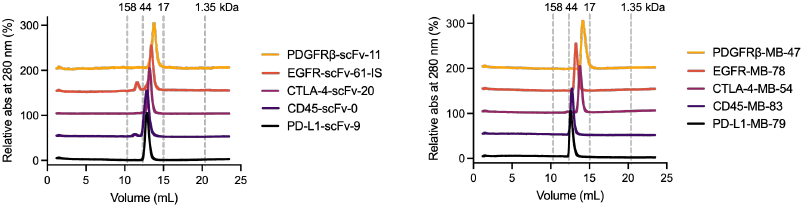
Analytical size-exclusion chromatography (SEC) of *E. coli* expressed binders. Designed scFvs (*left*) and minibinders (*right*) were sequentially purified by affinity chromatography and SEC, then a small portion of the pooled fractions were re-analyzed by SEC after storage at 4^◦^C for 24 hrs or more. The normalized 280 nm absorbance is plotted here with vertical offset. Molecular weight markers derived from size standards chromatographed under identical conditions are indicated by vertical dashed lines.

##### A.3.3.9 Structure determination by cryo-ELECTRON MICROSCOPY (Cryo-EM)

EGFR-his (ACRO Biosystems EGR-H5222) was reconstituted to 1 mg/mL in phosphate buffered saline (PBS, pH 7.4) and polished via size exclusion chromatography in PBS on an AKTA pure chromatography system with a Superdex 200 Increase 10/300 column (Cytiva). Peak fractions were concentrated to 1.25 mg/mL by spin filtration and mixed with the minibinder EGFR-MB-78, which had been purified by affinity and size exclusion chromatography from *E. coli* as described above, such that the final total protein concentration was 0.84 mg/mL and the minibinder was at a slight (1.1x) molar excess; final concentrations of minibinder and EGFR were approximately 10 *µ*M and 9 *µ*M, respectively. This complex was incubated for 14 hours at 4^◦^C prior to grid preparation.

Cryo-EM grids for the EGFR–minibinder complex were prepared using Ultrafoil R1.2/1.3 Au 300-mesh (Quantifoil Micro Tools, Germany) grids that were glow-discharged for 50 sec with a PELCO easiGlow. Grids were plunge-frozen using a Vitrobot Mark IV (Thermo Fisher Scientific) operated at 18^◦^C and 95% humidity, with 3 *µ*L of sample applied to each grid and blotted for 2 sec prior to vitrification. Cryo-EM data were collected on a Thermo Scientific Krios G4 microscope operating at 300 kV and equipped with an X-FEG electron gun, a Selectris X energy filter, and a Falcon 4i direct electron detector. Data collection was performed using Thermo Scientific EPU software, including tilted data acquisition strategies to overcome preferred orientation. Three different datasets were collected at 0^◦^, 20^◦^, and 40^◦^ stage tilts. Data were acquired at a magnification of 130,000x, corresponding to a calibrated pixel size of 0.94 Å per pixel, with a total electron dose of 74.71 e^−^/Å ^2^ and a defocus range of −1.0 to −1.8 *µ*m. A total of 14,217 micrographs were collected across the three tilt datasets.

All acquired data were processed with CryoSPARC v5.0.0 (128). Movie stacks in .eer format were imported without upsampling and split into 74 frames. Motion was corrected using PatchMotion job and CTF was estimated with PatchCTF job with default parameters. Micrographs were filtered to contain a subset with −1 to −2 *µ*m defocus range and estimated CTF *<*4.5 Å. Filtered datasets contained 991 micrographs for 0^◦^ tilt, 791 micrographs for 20^◦^ tilt and 3471 micrographs for 40^◦^ tilt. Initially particles were picked from a 0^◦^ tilt dataset using blob picker, extracted particles with a box of 420 pixels were processed in 2D classification and selected classes were used as templates for template picking of all three datasets. Template picking and subsequent 2D classification has yielded 18,113 particles for 0^◦^ tilt; 22,459 for 20^◦^ tilt and 110,547 for 40^◦^ tilt. Particles were re-extracted with recentering after 2D classification and used to generate an ab-inito model. C2 symmetry was identified in the ab-initio model and Non-uniform refinement was used to generate the final 3.83 Å map with symmetry enforced.

The central dimeric region of the map that included two copies of each EGFR domain III and minibinder were masked out and processed using local refinement, producing a 3.68 Å map of a central region. Anisotropy correction was performed with spIsoNet v1.0 (129). A directional Fourier shell correlation volume was computed from the two unfiltered half-maps and the FSC mask output by CryoSPARC. The same half-maps, mask, and 3DFSC volume were then provided to spIsoNet’s self-supervised reconstruction module, trained for 30 epochs. The corrected half-maps were combined in RELION 4.0.2 (130) to reconstruct the corrected volume, which was then sharpened in CryoSPARC with the −106 b-factor (same as the estimate from original reconstruction). The EGFR model from PDB ID 9IPB and the predicted model of the minibinder were first rigidbody docked and visualized using ChimeraX (131) and the minibinder model was further refined with Coot-1.3.1. The root-mean-square deviation (RMSD) between the refined and predicted models of the minibinder was calculated in ChimeraX, yielding an RMSD of 1.204 Å across 1,108 atom pairs; the flexible loop was excluded from the calculation due to poor resolution in the experimental map.

##### A.3.3.10 Epitope competition enzyme-LINKED IMMUNOSORBENT ASSAY (ELISA)

96-well clear MaxiSorp plates (Thermo Fisher) were coated with 80 *µ*L/well of 30 *µ*g/mL PD-L1-Fc (Acro Biosystems PD1-H5258), 20 *µ*g/mL EGFR-His (R&D Systems 11302-ER-050), or 20 *µ*g/mL CTLA-4-Fc (Acro Biosystems CT4-H5255) in phosphate buffered saline (PBS, pH 7.4) overnight at 4^◦^C with gentle rocking. 16-18 hrs later the coating solution was aspirated and the wells washed once with 200 *µ*L of PBS. The PBS was removed and replaced with 200 *µ*L/well of SuperBlock blocking buffer (Thermo Fisher) and incubated at room temperature for 2 hrs. Afterward, the SuperBlock solution was aspirated and replaced with 50 *µ*L/well of competition antibodies diluted in ELISA buffer (PBS containing 5% v/v SuperBlock) to a range of 0 to 600 nM concentration in a 3-fold dilution series. The competition antibodies were atezolizumab biosimilar (R&D Systems MAB10348), cetuximab biosimilar (R&D Systems MAB9577), and ipilimumab biosimilar (R&D Systems MAB9928), to compete with, respectively, designed binders to PD-L1, EGFR, and CTLA-4. These competition antibodies were chosen because their published, antigen-bound X-ray crystallography structures show epitopes highly similar to the predicted epitopes of ESMC-designed binders. A trastuzumab biosimilar (anti-Her2 antibody, BD Pharmingen 569599) was also diluted in ELISA buffer and used as a negative control.

Wells were incubated with competition antibodies for 30 min at room temperature with orbital shaking at 400 rpm on an Eppendorf Thermomixer. 50 *µ*L of 150 nM (for PD-L1 and EGFR) or 120 nM (for CTLA-4) designed minibinders or scFvs was then added to each appropriate well without removal of the previously added competition antibodies. scFvs in this assay were purified from *in vitro* translation while minibinders were purified from *E. coli* expression. Plates were orbitally shaken at 750 rpm for 15 sec to homogenize the mixtures, then incubated for 1 hr at room temperature with orbital shaking at 400 rpm. Afterward, contents were aspirated and each well was rinsed three times with 200 *µ*L/well ELISA wash buffer (PBS containing 0.05% v/v Tween-20), followed by the addition of 50 *µ*L/well of Strep-Tactin HRP conjugate (IBA Lifesciences) diluted 1,000-fold in ELISA buffer. Strep-Tactin HRP binds the Twin-Strep-tag on designed binders but not the competition antibodies. Plates were incubated with this reagent at room temperature for 15 min with 400 rpm orbital shaking. Afterward, the contents were aspirated and each well was rinsed five times with 200 *µ*L/well ELISA wash buffer.

Colorimetric ELISA signal was generated by the addition of 50 *µ*L of TMB substrate (3,3’,5,5’-tetramethylbenzidine, Thermo Fisher) per well, then quenched with 50 *µ*L of 1 M hydrochloric acid 30 - 120 sec afterward depending on the binder and before signal saturation. 450 nm absorbance was measured on a BioTek Synergy H1 plate reader. Background absorbance from wells with coating antigen omitted was subtracted from the raw measurements, and the adjusted measurements plotted with GraphPad Prism.

**Figure S20.**
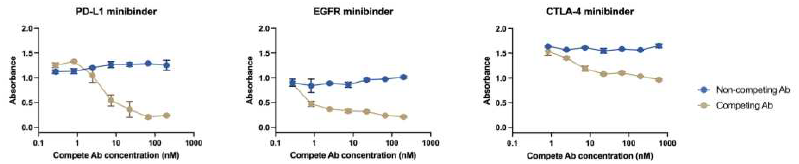
Competition enzyme-linked immunosorbent assay (ELISA) to characterize minibinder epitopes. Designed minibinders bound surface-anchored antigens in the absence or presence of therapeutic biosimilar antibodies overlapping the designed epitopes: atezolizumab (anti-PD-L1), cetuximab (anti-EGFR), ipilimumab (anti-CTLA-4), or non-competing trastuzumab (anti-HER2). Error bars indicate 1 SD across duplicate measurements.

##### A.3.3.11 Immunofluorescence staining of cultured mammalian cells with scFvs and minibinders

To promote cell adhesion, 8-well chamber slides (Cellvis) were coated with 150 *µ*L/well of Poly-D-Lysine for 1–4 hrs at 37^◦^C. Following aspiration of the coating solution, wells were washed twice with 200 *µ*L of sterile phosphate buffered saline (PBS, pH 7.4) and dried in a biosafety cabinet for 10 min. Human embryonic kidney 293T cells (ATCC) were then seeded at a density of 40,000 cells/well in Dulbecco’s modified Eagle’s medium (DMEM, Gibco) supplemented with 10% v/v FBS and allowed to recover overnight in a humidified incubator at 37^◦^C with 5% CO_2_.

**Figure S19.**
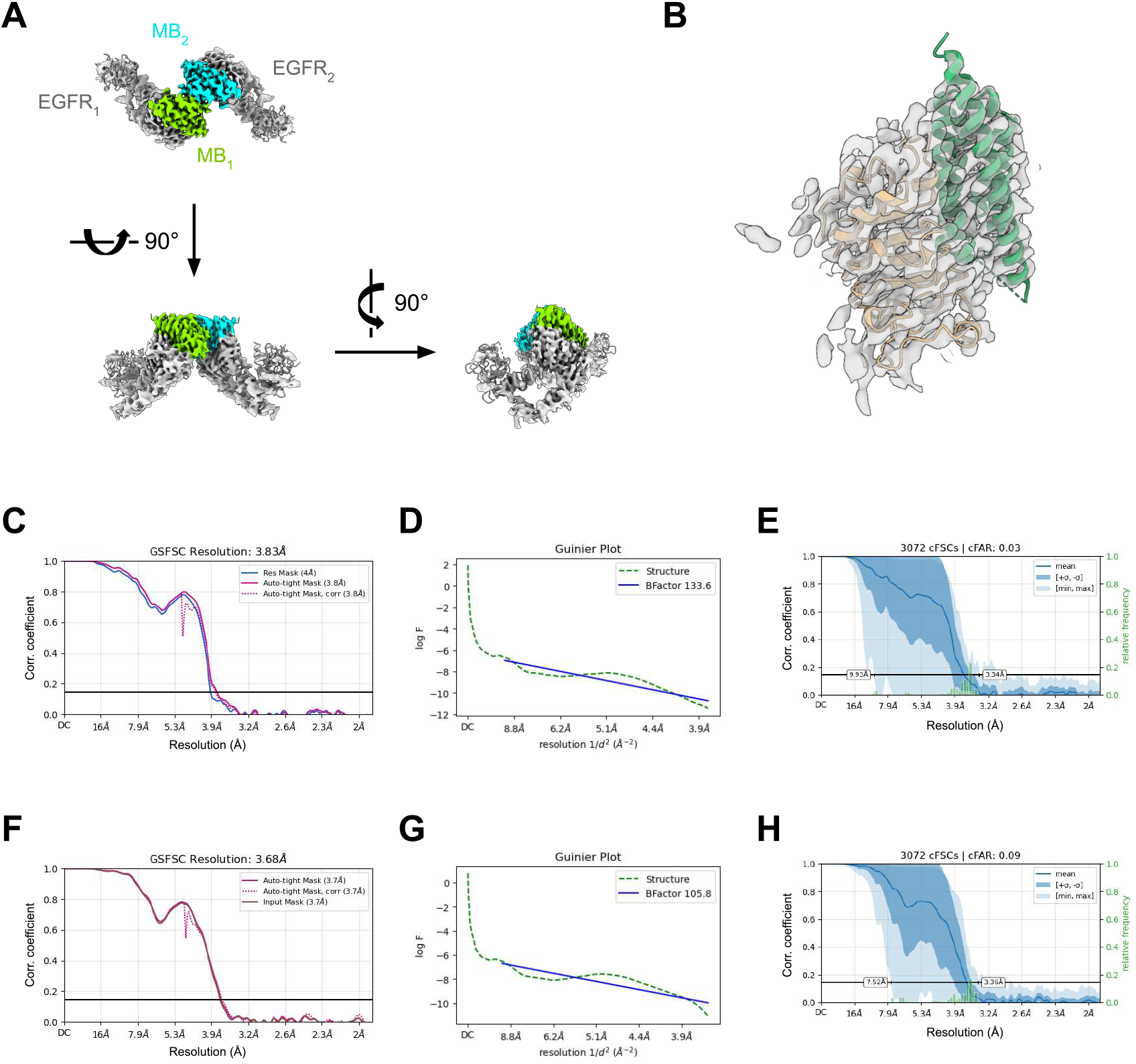
Cryo-EM structure and validation of the EGFR-minibinder complex. (A) Cryo-EM density map of the full complex from different orientations, demonstrating a 2:2 (EGFR:minibinder) stoichiometry mediated by minibinder dimerization. EGFR_1_ and EGFR_2_ represent individual EGFR protomers each consisting of domains I, II, and III; MB_1_ and MB_2_ represent the respective minibinders. (B) Cryo-EM density map of the locally refined region containing EGFR domain III (tan) and the minibinder (green), with the refined atomic model docked into the density. (C) Gold-standard Fourier shell correlation (GSFSC) plot for the global reconstruction, indicating a nominal resolution of 3.83 Å at the FSC_0.143_ threshold. (D) Guinier plot for the global reconstruction with an estimated B-factor of −133.6 Å ^2^. (E) Directional 3DFSC plot for the global map (conical area FSC plot), highlighting the presence of directional resolution anisotropy. (F–H) Corresponding GSFSC plot, Guinier plot, and directional 3DFSC plot for the locally refined map shown in (B), yielding a nominal resolution of 3.68 Å at the FSC_0.143_ threshold and an estimated B-factor of −105.8 Å ^2^.

At ∼ 70% confluency, cells were transfected with constructs expressing EGFR-GFP (132), EGFP-PD-L1 (133), PDGFRβ-IRES2-EGFP (134), or CTLA-4-mStayGold. EGFR-GFP was a gift from Alexander Sorkin (Addgene plasmid #32751; http://n2t.net/addgene:32751; RRID:Addgene 32751). pEGFP-N1/PD-L1 was a gift from Mien-Chie Hung (Addgene plasmid #121478; http://n2t.net/addgene:121478; RRID:Addgene 121478). pCAGGS-PDGFRb-IRES2-EGFP was a gift from Hirofumi Nakatomi (Addgene plasmid #204354; http://n2t.net/addgene:204354; RRID:Addgene 204354). The CTLA-4-mStayGold construct consists of CTLA-4 amino acids 1-200 tagged with mStayGold at the *C*-terminus spaced by a (Gly-Gly-Ser)_2_ linker, synthesized and cloned into the pTwist CMV vector by Twist Bioscience. For each well, 200 ng of DNA and 0.8 *µ*L of Lipofectamine 2000 (Invitrogen) were separately diluted into 25 *µ*L aliquots of Opti-MEM (Gibco). The diluted DNA and Lipofectamine were combined and incubated at room temperature for 10 min to allow complex formation. This 50 *µ*L transfection mixture was then diluted with 150 *µ*L of DMEM containing 10% v/v FBS. The existing cell culture medium was aspirated and replaced with the 200 *µ*L transfection medium. After 4-6 hrs, the medium was aspirated and replaced with 200 *µ*L of fresh DMEM supplemented with 10% v/v FBS.

Cells were stained 25-30 hrs after transfection. Prior to staining, 2X Bright-Live Cell Imaging Buffer (MP Biomedicals) was diluted to 1X with ultrapure water. Binders were diluted to 0.045–10 *µ*M in staining buffer (1X imaging buffer supplemented with 2% w/v bovine serum albumin, Fisher Scientific). To form fluorescent complexes, each binder was pre-incubated with Strep-Tactin XT DY-649 (IBA Life-sciences) at either a 1:1 molar ratio or with the Strep-Tactin in molar excess. This mixture was incubated at room temperature for 15–30 min, protected from light, and subsequently diluted with staining buffer to a final binder concentration of 45–100 nM. Cell culture medium was aspirated and replaced with 200 *µ*L/well of the diluted binder complex. Cells were incubated with the complex at room temperature for 30 min, protected from light. Cells were then washed gently with 200 *µ*L/well of wash buffer (PBS supplemented with 0.9 mM CaCl_2_ and 0.5 mM MgCl_2_). For nuclear counterstaining, cells were incubated for 10 min at room temperature in imaging buffer containing either 1 *µ*g/mL Hoechst 33342 (Thermo Fisher Scientific) or 2 drops/mL NucBlue Live ReadyProbes Reagent (Invitrogen). Finally, the nuclear stain was aspirated, and cells were washed gently 1–2 times with wash buffer before being maintained in 1X imaging buffer for visualization.

Images were acquired using a Leica DMi8 THUNDER inverted epifluorescence microscope equipped with a 63x/1.40 oil immersion objective and a K8 scientific CMOS camera (Leica Microsystems). Fluorescence was excited using a Leica LED5 engine and captured using standard filter cubes: DAPI (Ex: 355/56 nm, Em: 460/50 nm) for Hoechst 33342, FITC (Ex: 480/40 nm, Em: 527/30 nm) for GFP-fusion targets and reporters, and Cy5 (Ex: 620/60 nm, Em: 700/75 nm) for Strep-Tactin XT DY-649 labeled binders. Image acquisition was controlled via LAS X software (Leica Microsystems). Following acquisition, images were processed using ImageJ/FIJI.

**Figure S21.**
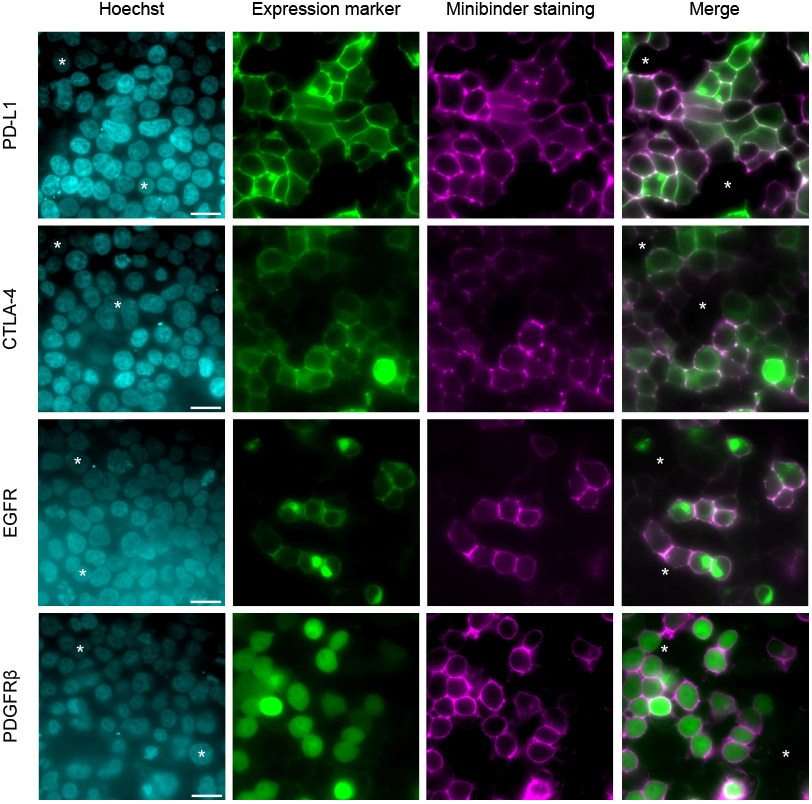
Live-cell immunofluorescence staining by designed minibinders. Live-cell immunofluorescence staining by designed minibinders for characterization of binding specificity. Human embryonic kidney 293T cells were transfected with constructs harboring the indicated targets tagged with green fluorescent protein (GFP, green). Cells were then stained with Hoechst 33342 (cyan) and Twin-Strep-tagged minibinders complexed with Strep-Tactin XT-DY649 (magenta), followed by imaging on an epifluorescence microscope. The GFP variant is fused to the target of interest in all cases except for PDGFR*β*, where it is translated separately from the same transcript via an internal ribosome entry site. The fourth column is a merge between the green and magenta channels. White asterisks denote untransfected cells or cell clusters, which show no detectable minibinder staining, indicating target-specific binding. Scale bars, 20 *µ*m.

##### A.3.3.12 Affinity measurement by biolayer interferometry

Multi-concentration kinetic binding measurements of *E. coli*-purified binders were collected by biolayer interferometry using a Gator Plus instrument (Gator Bio). Binding was performed in K buffer (Gator Bio), in 384-well tilt-bottom plates at 25^◦^C, 1000 rpm, and 5 Hz, with the exception of the CTLA-4 binders, for which binding was performed at 10 Hz. All reagents and consumables were obtained from Gator Bio unless otherwise specified. The antigen reagents (Acro Biosystems, R&D systems) were buffer exchanged into PBS before dilution. These antigens matched those described in the initial single-concentration screening experiments, with the exception of CD45, for which the Fc-tagged antigen was replaced with the his-tagged antigen to avoid avidity artifacts.

For EGFR, PD-L1, and PDGFRβ binders, the Twin-Strep-tagged binders were immobilized on Strep-Tactin XT probes (Gator Bio) and antigen association and dissociation was monitored. For CD45 binder affinities, the assay was flipped to circumvent nonspecific binding by his-tagged CD45 to the probes at high concentrations: His-tagged CD45 (Acro Biosystems CD5-H52H8) was immobilized on Anti-His probes (Gator Bio) and the association and dissociation of binders was monitored. For CTLA-4 binder affinities, the assay was flipped to minimize the effect of increased avidity of the CTLA-4 homodimer: his-tagged CTLA-4 dimer (Acro Biosystems CT4-H52H9) was immobilized on Anti-His probes (Gator Bio) and the association and dissociation of binders was monitored.

Binders (or CD45/CTLA-4 protein) were diluted to 25-50 nM and loaded simultaneously onto 8 probes to a threshold of 1 ≤ nm. Strep-Tactin XT probes were blocked for 30s with 1 *µ*M of Twin-Strep peptide (GenScript). Association to a half-log dilution series of antigen or binder, or buffer alone, was then monitored in parallel for 60-3600 sec, followed by dissociation in buffer for the same time frame.

Data was analyzed in GraphPad Prism (version 11.0.0, GraphPad Software, Boston, MA). Drift correction was performed by subtracting the buffer-only reference trace from each sensorgram. For local kinetic analysis, individual association traces were fit without constraining Y_0_. The resulting observed rate constants (*k*obs) were verified to be linear with antigen concentration, across an antigen concentration range spanning at least 1.5 orders of magnitude (≥ 4 concentrations) and extending to at least 10-fold over the KD calculated below. The fitted Y_0_ value from each local association fit was subtracted from the corresponding trace to correct association-phase baseline offsets. For all binders except for the CTLA-4 scFv, drift- and baseline-corrected sensorgrams from each concentration series were globally fit using Prism’s ‘Association then dissociation’ model, with shared ‘Kon’, ‘Koff’, and ‘Bmax’ values, concentrations (‘HotNM’) defined as dataset constants, and nonspecific binding (‘NS’) fixed to 0. For the CTLA-4 scFv, non-ideal, biphasic dissociation was observed, so affinity was obtained from the association phase only, using Prism’s ‘Association kinetics - Two or more conc. of hot.’ model with shared ‘Kon’, ‘Koff’, and ‘Bmax’ values and concentrations (‘HotNM’) defined as dataset constants.

##### A.3.3.13 Cellular assay for PD-L1/PD-1 blockade by designed binders

Purified PD-L1 minibinder and scFv were assessed for their ability to relieve inhibition of T-cell receptor (TCR) signaling via the PD-1/PD-L1 Blockade Bioassay (Promega), according to manufacturer instructions with minor modifications. All materials were obtained from Promega unless otherwise specified.

CHO-K1 artificial antigen presenting cells (aAPCs) expressing human PD-L1 and an artificial TCR activator were thawed at 37^◦^C, diluted into 14.5 mL of pre-warmed Ham’s F-12 medium with 10% fetal bovine serum (FBS), then seeded into two white 96-well flat-bottom cell culture plates (Corning). Cells recovered in a humidified incubator at 37^◦^C and with 5% CO_2_. Culture medium was removed 18 hrs later and replaced with RPMI 1640 medium containing 1% FBS and a designed binder (purified from *E. coli*), an scFv fragment of atezolizumab (purified from *in vitro* translation), or a trastuzumab biosimilar (569599, BD Pharmingen). Each binder concentration was assayed in duplicate. Immediately after, Jurkat T cells expressing PD-1 and a luciferase reporter driven by the nuclear factor of activated T cells (NFAT) response element were thawed, diluted into prewarmed RPMI with 1% FBS, and added to wells containing aAPCs and binders. Cell mixtures were then incubated for 6 hrs in a humidified incubator at 37^◦^C and with 5% CO_2_.

After 6 hrs, plates were equilibrated to room temperature and Bio-Glo reagent in Bio-Glo luciferase assay buffer was added to each well containing cells. Luminescence was measured after 15 min of incubation, on a BioTek Synergy H1 plate reader at a height of 1 mm and a gain of 150. Data was fit with four-parameter logistic curves in GraphPad Prism (GraphPad Software, Boston, MA).

#### A.3.4 Novelty and Diversity of Validated Binders

Structural novelty. Designed binders are predicted to exhibit binding modes not represented in known structures deposited within the PDB. We use a union of `mmseqs easy-search -s 7.5 --max-seqs 1000 and foldseek easy-search -s 7.5 --alignment-type 1 --max-seqs 1000` against the PDB to find targets which are similar in either sequence or structure. All protein chains with 1 or more heavy-atom contacts (*<* 5 Å) to the target-similar chain are selected.

**Figure S22.**
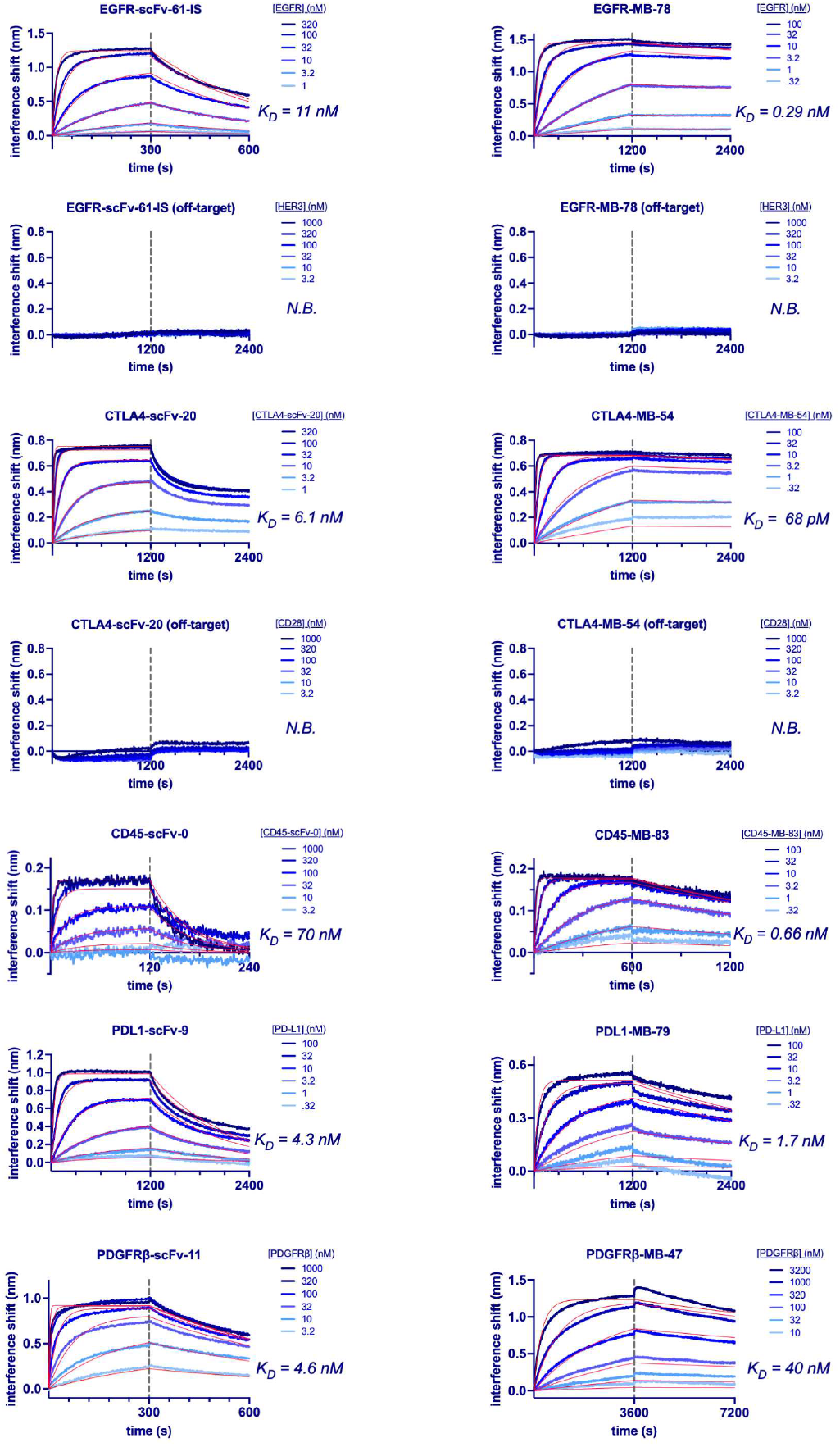
Multi-concentration kinetic binding measurements of selected binders. Binding affinities of representative scFv (left column) and minibinder (MB, right column) designs against each target were measured using biolayer interferometry (BLI) at several concentrations. Twin-Strep-tagged binder or His-tagged recombinant target antigen was immobilized on Strep-Tactin XT or anti-His probes, respectively, and bound to varying concentrations of target antigen or binder (indicated in the legends). Binders targeting EGFR and CTLA-4 showed no detectable interaction with the corresponding homologous domains of HER3 and CD28, respectively, demonstrating high target specificity. Blue traces show drift- and baseline-corrected sensorgrams, and overlaid red lines represent kinetic global fits of the association and dissociation phases (a 1:1 binding model) used to determine affinity, with the exception of the CTLA-4 scFv, whose dissociation appeared biphasic for unknown reasons, so affinity was determined using a global fit of the association phase only. Vertical dashed lines indicate the transition from the association phase to the dissociation phase. Calculated equilibrium dissociation constants (K_D_) are displayed for each target interaction. ‘IS’ in the binder name indicates that the binder was selected from the inference-scaled pool of designs. N.B. denotes no binding.

Structural divergence is measured as the nearest-neighbor Cα RMSD of the binder after CEAlign of the target chains (135). For each binder, we report the minimum target-aligned Cα RMSD over all interactions with target-similar chains in Figure 3C.

##### Sequence novelty of antibody CDRs and designed minibinders

The sequence novelty of antibodies reported in Figure 3C was measured using the minimum CDR edit distance to SAbDab (105). Specifically, the edit distance reported is the minimum between either of the designed variable domains’ concatenated CDRs and any such concatenated CDRs in SAbDab. For minibinders, we queried NCBI NR with BLASTP using an E-value cutoff of 10 and `max target seqs=500`. The percentage identity and E-value of the lowest E-value hit are reported in Figure S23 for each confirmed binder.

##### Diversity of binding poses

Where novelty measures deviation of designs from structures deposited within the PDB, diversity measures deviation of designs from each other. The same structural-distance definitions, applied between pairs of designs, are used to compute the pairwise distance matrix between all designs. A dendrogram is constructed from this matrix using UPGMA average-linkage clustering (`scipy.cluster. hierarchy.linkage(squareform(D),method= “average”`)) and the number of clusters at a given RMSD threshold is obtained by cutting the dendrogram at that height (`scipy.cluster.hierarchy.fcluster, criterion=“distance”`). The number of clusters as a function of the RMSD threshold is shown in Figure S24.

##### Diversity of minibinder structures

The low and high compute experiments yielded 473 total minibinder hits (after de-duplicating for designs selected for both experiments). These hits correspond to 288 unique structural clusters after structural clustering of designs against a shared target using FoldSeek with a threshold of 0.5 TM (93).

##### Diversity of minibinder and scFv sequences

We cluster sequences with MMseqs2 `easy-cluster` (136) on the concatenated heavy- and light-chain CDRs (scFv) or the full binder sequence (minibinders), and report the number of clusters as a function of the `--min-seq-id` threshold in Figure S24C,D.

### A.4 ESM Cambrian Sparse Autoencoders

#### A.4.1 Models and Training Details

Sparse autoencoders (SAEs) are encoder-decoder networks trained with a reconstruction loss objective and an additional sparsity constraint. Specifically, let 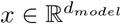 be a single token’s hidden-state activation from the frozen base model. The SAE maps *x* to a sparse vector 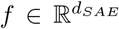 with *d*_*SAE*_ ≫ *d*_*model*_. The SAE is trained to reconstruct 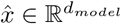. Each input to the encoder is Z-score normalized.

The encoder is defined as

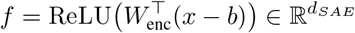

and

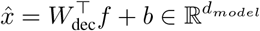

where 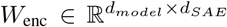 and 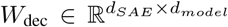. The bias *b* is tied across both encoder and decoder. Rows of the decoder matrix are normalized to unit vectors after each training step to maintain an interpretable dictionary.

To enforce sparsity of SAE representations, SAEs are trained with a TopK sparsity constraint, where only the *K* highest activations of the feature vector are kept in the intermediate representation before decoding, and all other activations are set to zero. At training time, this TopK is applied at batch level by flattening a batch of size *B* and keeping the top *K* · *B* entries across the batch. At inference time, the top *K* per token are selected to ensure that the active features per token are not dependent on other tokens in the batch.

SAEs are trained with a typical reconstruction mean squared error loss plus an auxiliary loss term to prevent “dead features” (features with zero activation that no longer receive gradient updates). Features that have had zero activation for five consecutive forward passes are denoted as dead, and the set of dead features D are trained to reconstruct the residual 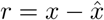 left by the non-dead features. This enables dead features to keep receiving gradient updates and reactivate during training.

Define the dead-only sparse code as

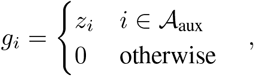

where 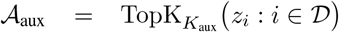 and 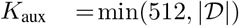, then the residual reconstruction is

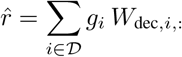

In total, the loss function is

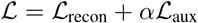

where 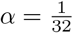 and

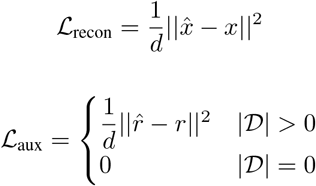

**Figure S23.**
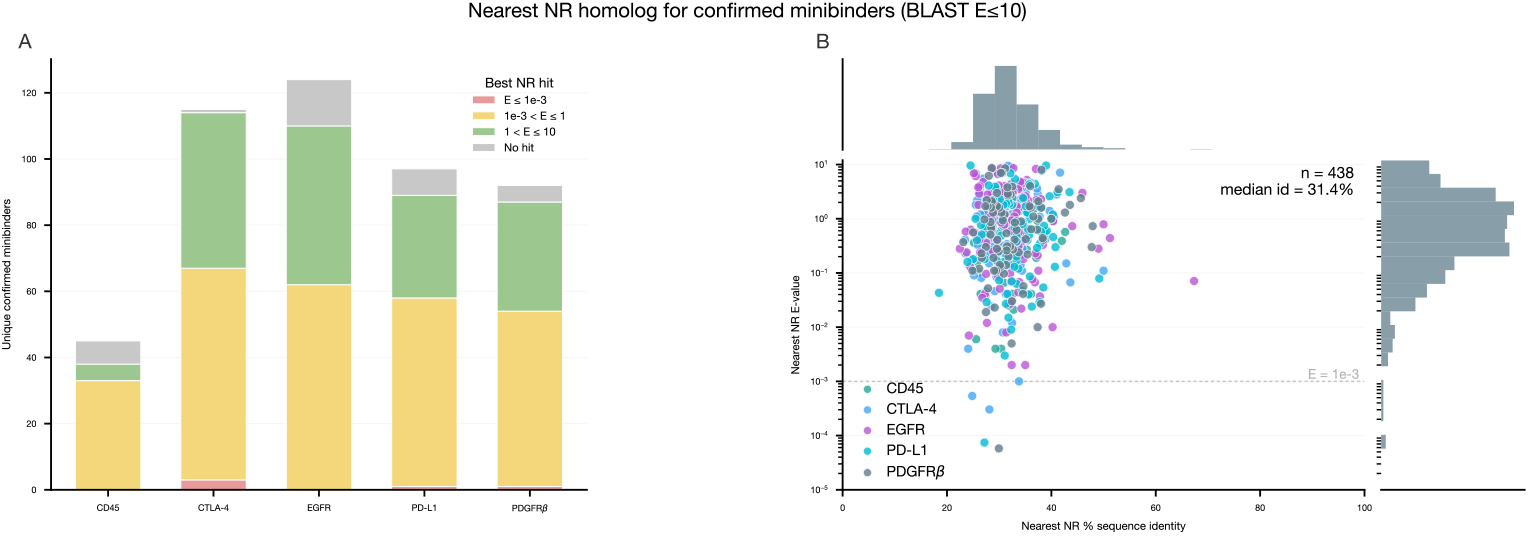
Sequence novelty of designed minibinders against NCBI NR. Most significant hit in the non-redundant (NR) protein database for each experimentally validated minibinder sequence, as reported by BLAST (search E-value ≤ 10). A. Counts of unique confirmed minibinders per target, stratified by the E-value of the most significant NR hit (10^−3^ *< E* ≤ 1, 1 *< E* ≤ 10, 10 *< E* ≤ 10^3^, or no detectable hit). No hits meet the conventional significance threshold (*E* = 10^−3^). B. Nearest NR % sequence identity versus E-value for BLAST-detectable hits (*n* = 438), colored by target.

**Figure S24.**
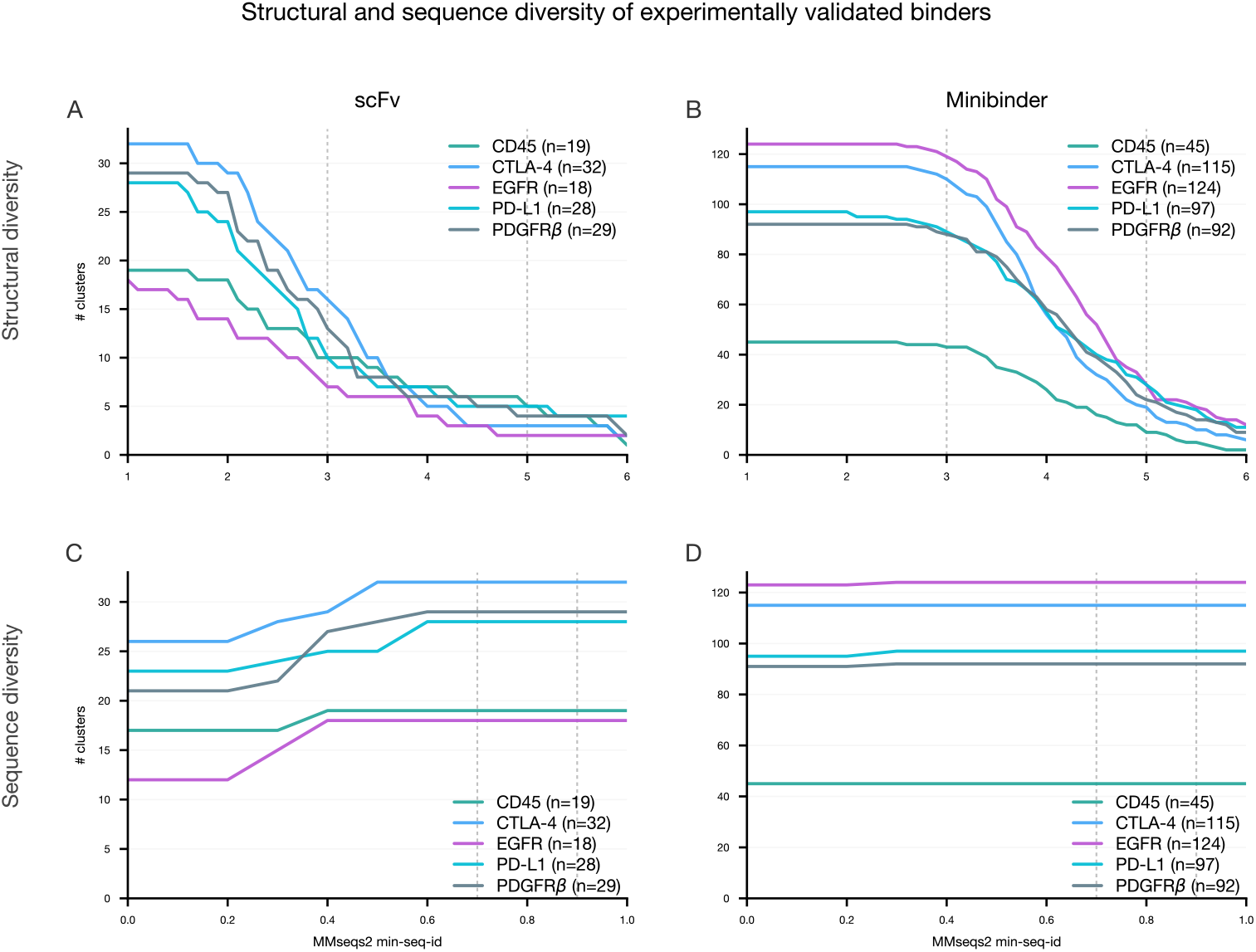
Structural and sequence diversity of designed minibinders and antibody scFvs. Number of clusters as a function of clustering threshold for each experimentally validated (target, modality) cohort. A. B. Structural clusters as a function of target-aligned binder C*α* RMSD threshold for scFv and minibinder hits, respectively. Structural clustering of scFvs resolves multiple distinct clusters at 5Å for all targets, consistent with multiple binding poses rather than redesigns of a single solution. C, D. Sequence clusters as a function of MMseqs2 minimum sequence identity threshold for concatenated scFv CDRs and minibinders sequences.

Our training data consists of UniRef90 sequences clustered at 70% sequence identity (35.7% of the training dataset), MGnify clustered at 70% sequence ID (10.7%), and the JGI dataset (53.6%). All SAEs are trained on a total of 8 billion tokens drawn from the same dataset of clustered Uniref, MGnify, and JGI as the base ESMC models with proportions 36%, 11%, and 54%, respectively. Training hyperparameters are listed in Table S12.

**Table S12.** SAE training hyperparameters.

| Parameter | Value |
| --- | --- |
| $d_{\text{model}}$ | 2560 |
| $d_{\text{SAE}}$ | $\{2^{13}, 2^{14}, 2^{15}, 2^{16}, 2^{17}\}$ |
| $K$ | $\{8, 16, 32, 64, 128\}$ |
| max train tokens | $8 \times 10^9$ |
| max train steps | 61,035 |
| batch size | 16 (8 for $2^{17}$ codebook) |
| max sequence length | 2048 |
| learning rate | $7 \times 10^{-5}$ |
| optimizer | AdamW |
| $\beta_1$ | 0.0 |
| $\beta_2$ | 0.95 |
| warmup steps | 1000 |
| $K_{\text{aux}}$ | 512 |
| $\alpha$ | $\frac{1}{32}$ |
| dead feature threshold | 5 forwards |
| precision | bf16-mixed |

SAEs were trained on hidden states of each layer of the ESMC 300M, 600M, and 6B parameter models with codebook sizes of 16, 384, 65, 536, and 131, 072. Exact SAEs used in each analysis are listed in the respective methods— most analyses use layer 60 of ESMC 6B.

#### A.4.2 Sparse Autoencoder Evaluations

##### A.4.2.1 Feature Activation Distributions

Feature activations for each protein were computed on a 90% clustered, high-quality subset of SwissProt for three SAEs trained on representations from ESMC 6B with *k*=64 sparsity and codebook sizes of 16,384, 65,536, and 131,072 features. The maximum activation of each SAE feature across its the residues of each protein was recorded, yielding a matrix of shape number of proteins times number of features for each SAE.

For each SAE and each of three activation thresholds (*>*1.0, *>*5.0, and *>*10.0), the number of proteins whose max activation exceeded the threshold was counted per feature. These counts were converted into a frequency by dividing by the total number of proteins considered. Features were sorted by ascending frequency within each SAE, threshold pair and the empirical cumulative distribution function of the proportion of features is shown in Figure S25.

**Figure S25.**
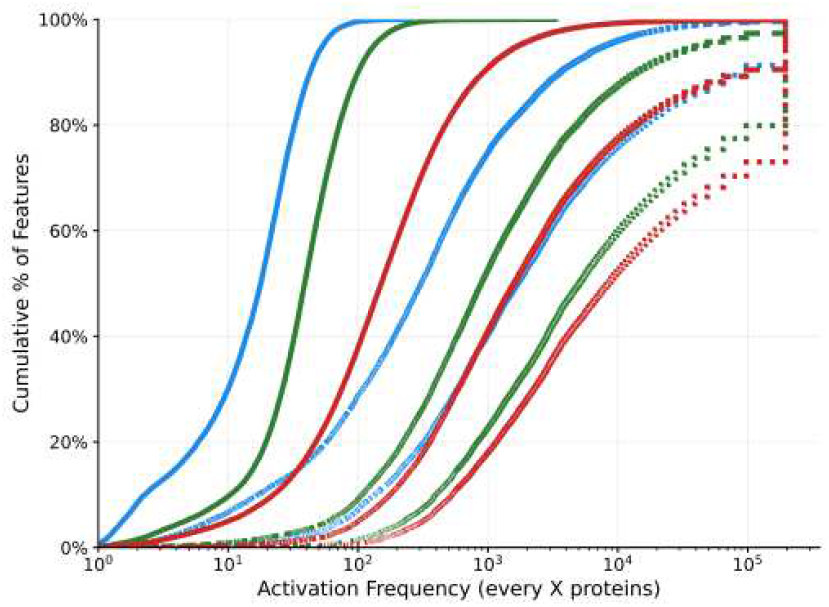
SAE feature activation distributions. Proportion of features with activation *>*1, 5, and 10 for codebook sizes 16, 384, 65, 536 and 131, 072. For a given activation threshold, features in larger codebooks appear more sparsely than features in small codebooks.

##### A.4.2.2 Layerwise Reconstruction and Perplexity Analysis

To determine the information density at different depths in the model, SAEs were trained on each layer of the final trained checkpoint of ESMC 6B. Two metrics were chosen as proxies for information density: layerwise SAE reconstruction loss (computed on the hidden state after passing the SAE for each layer) and layerwise perplexity degradation (change in full-sequence perplexity after reconstruction of the hidden state with each layer’s SAE). The motivation for both of these metrics is similar, as more information-rich representations are assumed to be both “harder” to reconstruct and also lose essential information needed for masked token prediction. SAEs with 16, 384 and 131, 072 dimensional codebooks were used for reconstruction loss analysis, and only 16, 384 was used for PPL degradation.

From the plots shown in Figure S26, it is clear that layers 60–70 contain the most information, with gradual increases in information density in layers 0–50 and a steep dropoff in the final layers of the model. This suggests the final layers are specialized for logit prediction rather than rich representations. One could hypothesize that a representation that combines information across all layers of the model would be more powerful than any individual layer’s representation, but such speculation is beyond the scope of this appendix.

##### A.4.2.3 Feature Specificity and Granularity

SAE features capture a wide range of different types of concepts. Even without specific knowledge of what biological concepts are being represented, the landscape of the concept space that the model uses to represent proteins can be characterized based on summary statistics across a reference database, in this case ∼ 208M proteins from Uniref90 (release 2025 03). We define two metrics for summarizing the residue-level activation pattern of each feature, one that captures its specificity for specific protein families (an interprotein metric) and one that captures its local granularity within the sequences it activates on (an intra-protein metric).

For specificity, we compute Pfam domain boundaries using pyHMMER’s HMMscan implementation (137, 138) against the Pfam-A database (81) and cross-reference with the residue-level feature activations. For residues which are annotated within a Pfam domain, all features at that position are assigned the domain’s accession, while residues outside any domain are assigned to the nearest annotated domain in the same protein (to account for features which are consistently just outside the HMM-annotated region). Features can be associated with multiple unique domains, but each domain only counts once per protein. Proteins with no annotated domains contribute to a separate unassigned bin. For each feature, the number of proteins *n*(*f, d*) where the activation of *f* falls in domain *d* is tabulated, a protein-level domain probability *p*(*f, d*) = *n*(*f, d*)/ Σ_*d*′_ *n*(*f, d*)′ is computed, and the final metric is derived from the Shannon entropy of that distribution:

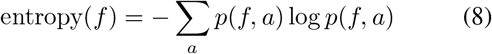

Low entropy indicates the feature concentrates in a small number of Pfam families (family-specific), while high entropy indicates broad distribution across many domain contexts (universal). The entropy value itself is not particularly interpretable, so we then z-score the entropies over all 16, 384 features to produce a normalized specificity value.

For granularity, for each pair of feature *f* and protein *p*, we identify the residue positions *x*_*i*_ where *f* has non-zero activationPand compute its mean contiguous run length, mcrl = ∑_*r*∈ℛ_ |{*x*_*i*_}_*i*∈*r*_|*/*|ℛ|, where ℛ is the set of all contiguous runs *r* of activated positions (a contiguous run is defined as a span of residues in which no more than one consecutive position has zero activation). To focus on the primary concept the feature represents and suppress noise from sub-threshold activations, which tend to be more sparsely active across the sequence, mcrl values are accumulated into per-feature histograms binned by the protein-level max activation, then apply Otsu’s method (139), originally developed for image processing, to each feature’s activation histogram to find a feature-specific threshold *t*_*f*_ that maximizes between-class variance, and the mean mcrl is computed over only pairs of *p, f* whose max activation exceeds the threshold.

Individual features represent conserved concepts that are reused across a large number of proteins, with even the more specific features spanning many unique Pfam domains. Figure S27 shows the distribution of the number of unique domains represented for each feature in the two major modes of the distribution: universal local features (Mode 1, bottom left) and more specific broadly activating features (Mode 2, top right). Considering only families that have at least 10 examples for each feature, features in Mode 2 still activate on a median of 189 unique domains, with a minimum of 19 and a maximum of 1,865. These features, while still more broadly conserved than a single family, are nonetheless much more specific than those in Mode 1, which activate on a median of 2,295 families with count *>*10 (min 109, max 7,463).

##### A.4.2.4 Feature Normalization

Features represent concepts at different scales and levels of specificity, meaning that some are more directly informative for understanding the function of a protein than others. Additionally, features have different dynamic ranges in their activations, spanning two orders of magnitude from as low as 1 to as high as 50, and more universal features tend to have a larger range of activation magnitudes. When comparing the activation of individual features across proteins this is not an issue, but in order to assess the relevance of each feature relative to others within a protein or set of proteins, it is useful to normalize the features to a scale that is consistent and biologically meaningful. To account for both the variation in specificity and activation range across features, we normalize the activations using a feature-specific normalization factor derived using the maximum activation for the feature and the number of proteins on which it activates in UniRef90. Unless otherwise stated, any features referred to with specific activation values use these normalized activation values.

Specifically, we compute the following two quantities given a reference dataset *D* of proteins *p*:

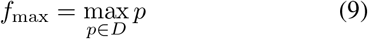

**Figure S26.**
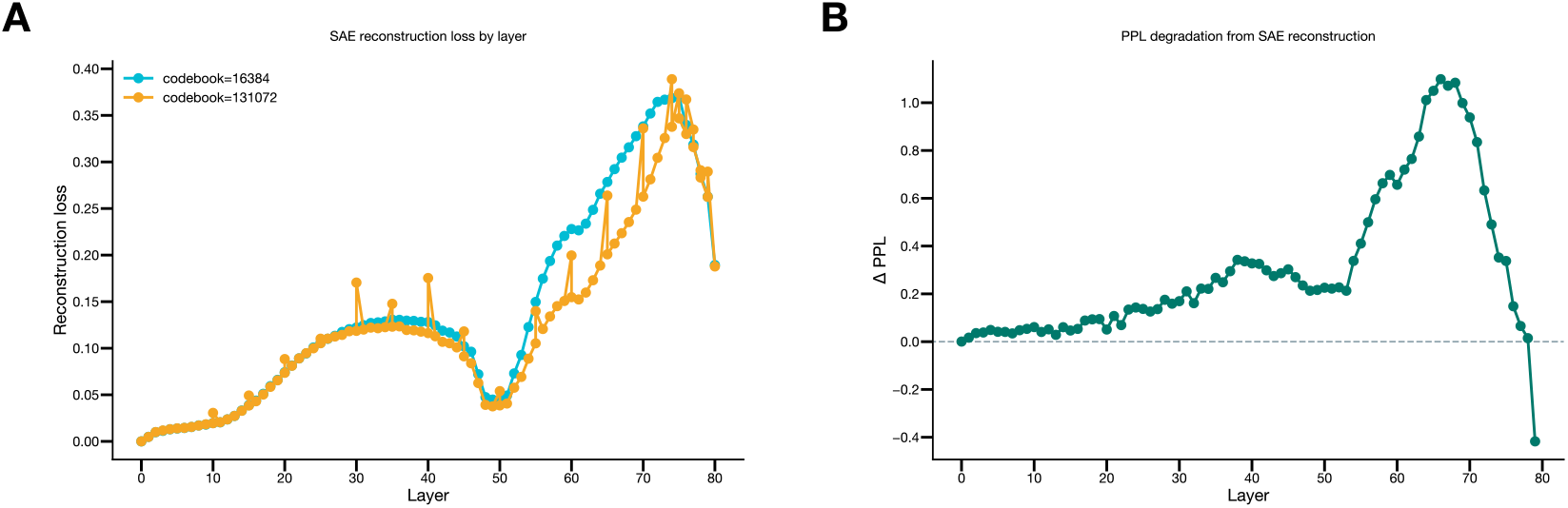
Layerwise SAE evaluation. (A) SAE reconstruction loss across layers for 16k and 131k codebooks. (B) Perplexity degradation after SAE reconstruction at each layer.

**Figure S27.**
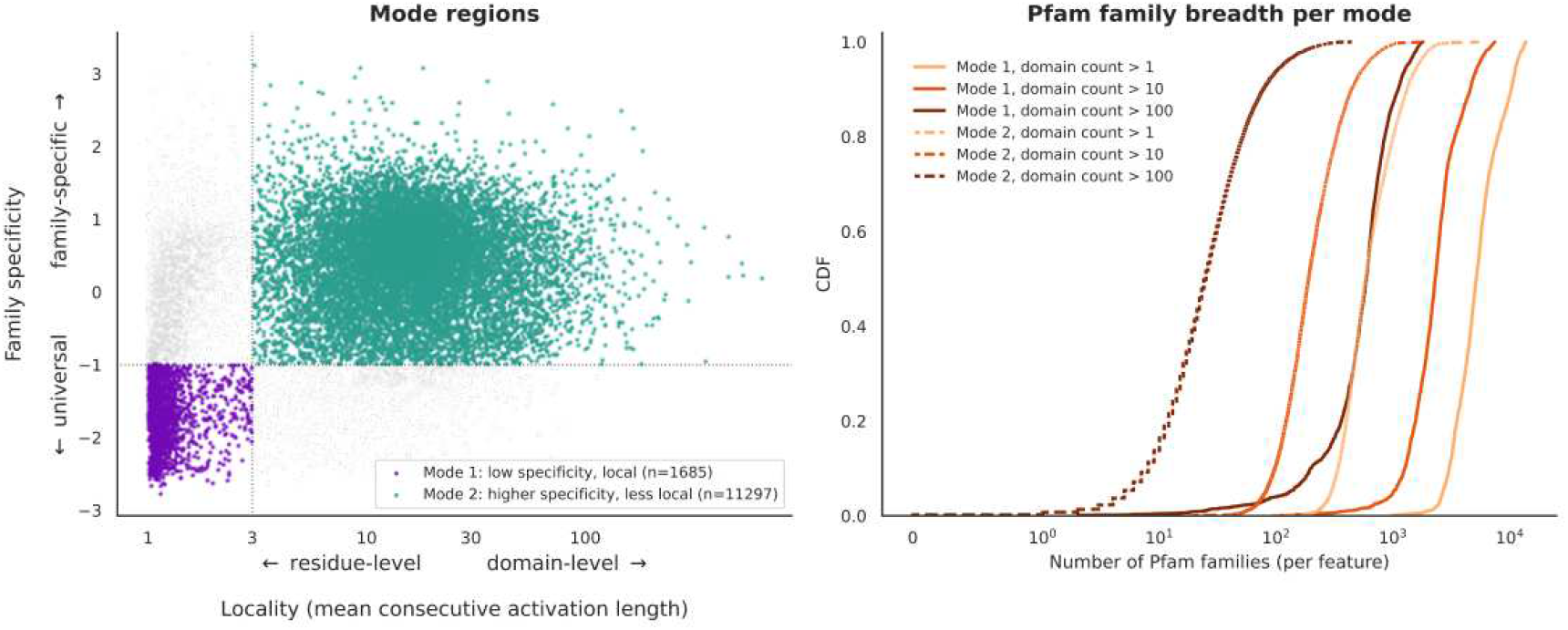
Feature specificity and granularity modes. Left: Division of specificity-granularity space into quadrants, defining two primary modes. Right: Number of unique Pfam domains associated with features in each mode, considering only domains with count >1, *>* 10, or *>* 100 for each feature.

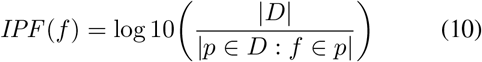

The normalized feature activation *f* ^*^ can then be derived from its raw activation *f* by:

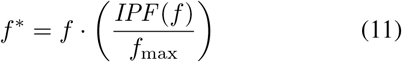

#### A.4.3. Biological Concepts in the SAE Feature Space

##### A.4.3.1 Automated Feature Descriptions

We develop a multi-agent system to describe features in natural language, including information on the concept the feature represents as well as examples of proteins in which it activates and a description of its activation pattern. Figure S28 shows a schematic of this system, which works through an iterative hypothesis-verification cycle with multiple data splits and holdouts. We find that such a system produces more generalizable descriptions of the concept a feature represents than a single agent provided with a set of top-activating proteins and annotations. The system consists of the following steps.

###### Step 1. Get top-activating proteins

The first step is to look up the top 200 proteins with the highest activation for the feature in a well-annotated protein dataset derived from SwissProt. SwissProt assigns an “annotation score” from 1–5 for each protein based on the amount of functional annotations contained in a particular protein entry. We first subset the full SwissProt database to only proteins with an annotation score of 3 or higher, representing a set which are of highest confidence and which will provide the most functional context to the analysis agents. This dataset is then clustered at 90% sequence identity using MMSeqs2 (136) to ensure that descriptions are not biased by a large number of very similar proteins. SAE feature representations for the resulting reference dataset of ∼ 195k proteins were used to create a lookup table from each feature to a ranked list of proteins and activation values. This dataset is used as a reference dataset throughout this work.

###### Step 2. Generate initial observations

The full list of 200 SwissProt proteins with their names, descriptions, organisms / taxonomic information, and a list of residue positions where the feature is active (e.g. “M1 R2 G3 … “) is provided to an agent which generates a set of initial observations about the feature. This can include general global patterns of activations (e.g. always activates on the N-terminus, only activates on 2–3 acidic residues per protein), taxonomic information (e.g. activates only on fungal proteins), and functional patterns (e.g. all are signaling proteins, all are localized to mitochondria). In later steps, the agents see mostly information from specific proteins, so this information provides some global context and regularizes against over-specific descriptions. It is also the agents’ main source of information such as taxonomy which may span multiple groups of proteins.

###### Step 3. Group proteins by function

In this step, the same list of 200 proteins, including their names and SwissProt descriptions, are provided to a GPT-5 (140) agent with high reasoning which is prompted to group the proteins into 2–10 groups based on their biochemical function. This ensures that the descriptions are generated based on a wide range of proteins and do not end up overly influenced by the most common families, or the families with the highest total activation values. At this stage, we also hold out a random protein from each group to use for validation.

###### Step 4. Generate initial feature description in the context of each functional group

For each of the groups identified in the previous step, a single protein is randomly selected to analyze in depth. The goal of this step is to describe the concept on which the feature activates in this specific class of proteins. Since features can represent concepts at different scales, from single residues to domains, the residue-level activation pattern is provided at this stage. A table is constructed where each row contains the residue-level information at each sequence position in the selected exemplar protein. At each position, the table includes the activation value of the feature, the secondary structure computed from the structure predicted by ESMFold2 (an earlier experimental version, in single sequence mode with three recycles), and all functional annotations at the corresponding position in SwissProt. These annotations include domain annotations from databases such as Pfam as well as functional annotations such as ligand binding, catalytic sites, functional regions, and disorder. This table is provided to GPT-5 in JSON format to produce a detailed description of the feature’s activation in the context of the protein provided.

###### Step 5. Validate initial description with the other proteins in the group

To make sure the description isn’t overly specific to the analyzed protein, the description and the list of proteins not selected as exemplars is provided to a validation agent (GPT-5 with medium reasoning) to assess whether the description applies to all proteins. This agent is not given the full residue-level context that is given for the exemplar protein, but high-level information about the proteins is generally sufficient to confirm that the description generalizes to other proteins in the group. If the validation agent rejects the description, a new exemplar is selected and Steps 4–5 are repeated (up to 3 times, after which the description is accepted as is).

###### Step 6. Unify descriptions from all functional groups into a single description

Once Steps 4–5 have been completed for all functional groups, all descriptions are provided to an agent (GPT-5 with high reasoning) to discern the unifying concept which connects all descriptions and would enable the feature description to generalize across classes of proteins. We also provide this agent with the initial observations produced in Step 2 so that it can incorporate cross-family context. The LLM is then prompted to provide the following information about the feature:

**Figure S28.**
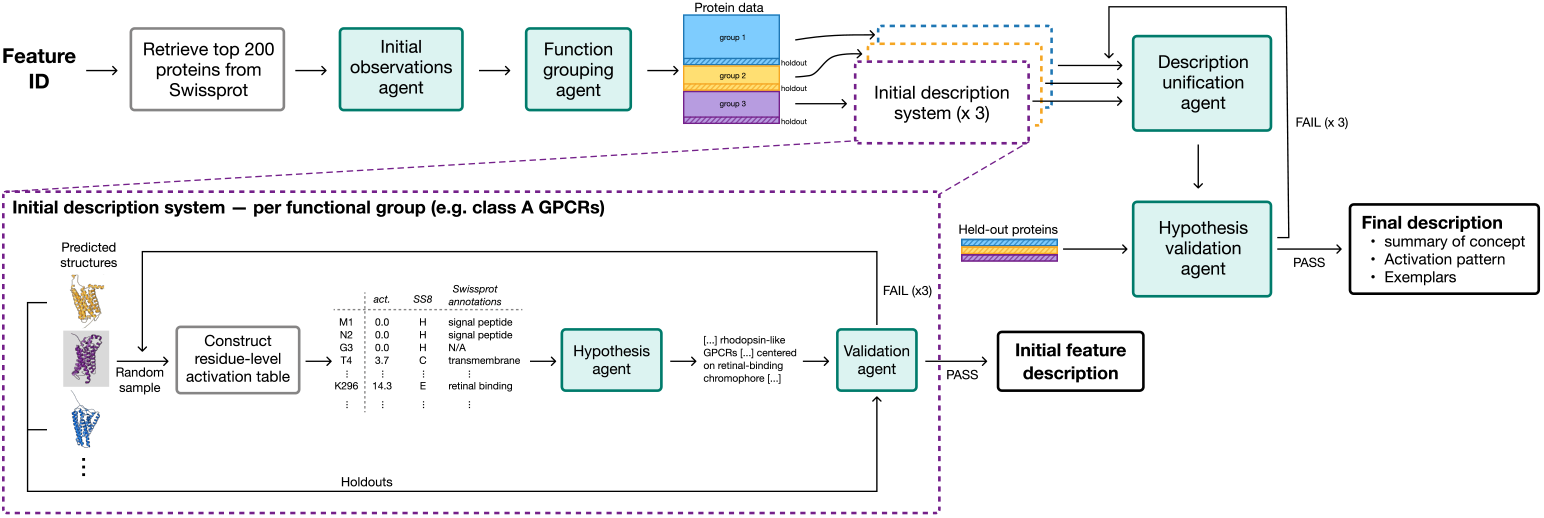
Multi-agent system for generating feature descriptions based on top-activating proteins from Swissprot. The top shows the overall flow, and the bottom shows a detailed view of the process for creating an initial description for the feature’s activation in a single protein group. This basic block is repeated for each protein group and then aggregated to create the final description. At each stage, held out proteins are used to validate the description to improve generalizability of the final description. Deterministic data or code steps are shown in empty black boxes, and agentic steps are shown in teal boxes.

- **Summary:** what biological concept does this feature activate on?
- **Activation pattern:** what is the typical activation pattern (e.g. domain-level, regional around a functional site, sharp peaks on specific residues)
- **Exemplars:** what are key examples of protein families that this feature activates on, and what does the feature specifically represent in each?

Separating the feature description into these categories encourages the LLM to consider specific evidence from the candidate descriptions in its final summary, and provides a canonical format that can be consistently interpreted in analysis of new proteins in future analysis.

###### Step 7. Validation of final description with held out proteins

In the final step, the unified description is provided to a validation agent (again GPT-5 with medium reasoning) along with the set of proteins held out from each group in Step 1. The validation agent is prompted to assess whether the final feature description is applicable to all held-out proteins. If it is, the description is accepted; if not, Steps 6– 7 are repeated up to three times until the description passes.

##### A.4.3.2 Feature Classification

The feature descriptions generated by the above procedure were used to classify the features into a set of eight broad categories spanning structural, functional, and biochemical concepts at different levels of complexity. These categories were chosen manually based on inspection of the feature descriptions in the context of proteins from SwissProt. The categories and category descriptions in Table S13 were provided along with the feature description to GPT-5 with medium reasoning effort, which returned simply the name of the category which best matches the feature description.

These categories are not mutually exclusive, but are illustrative of the distribution of concepts within the model’s representation space at each level of the biological hierarchy. It is also important to note that both the feature descriptions and the resulting classifications are biased by the type of information available in SwissProt proteins with extensive annotations of binding sites, catalytic sites, and domain information will contribute more to the description than those without any annotations. Many features which appear in proteins or domains with no annotations will simply be annotated with descriptions such as “low-complexity region” or “region enriched in polar residues”. Some of these concepts may in fact be associated with well-defined functions that are simply not well characterized yet, and the features could provide a path to improving our understanding of these lesser-known areas of biology.

Features from all categories are used in the representation of individual proteins. Figure S29 shows the fraction of features in each category that are used in each of the ∼ 195k proteins in our 90% clustered Swissprot reference dataset. The categories are roughly organized in a hierarchy according to their complexity, and their usage patterns reflect that—every category is represented in every protein, and proteins tend to activate a large number the features in more universal categories (residue identity, secondary structure, biochemical microenvironment) and a smaller number in more specialized categories (functional sites, domains, localization).

**Table S13.**
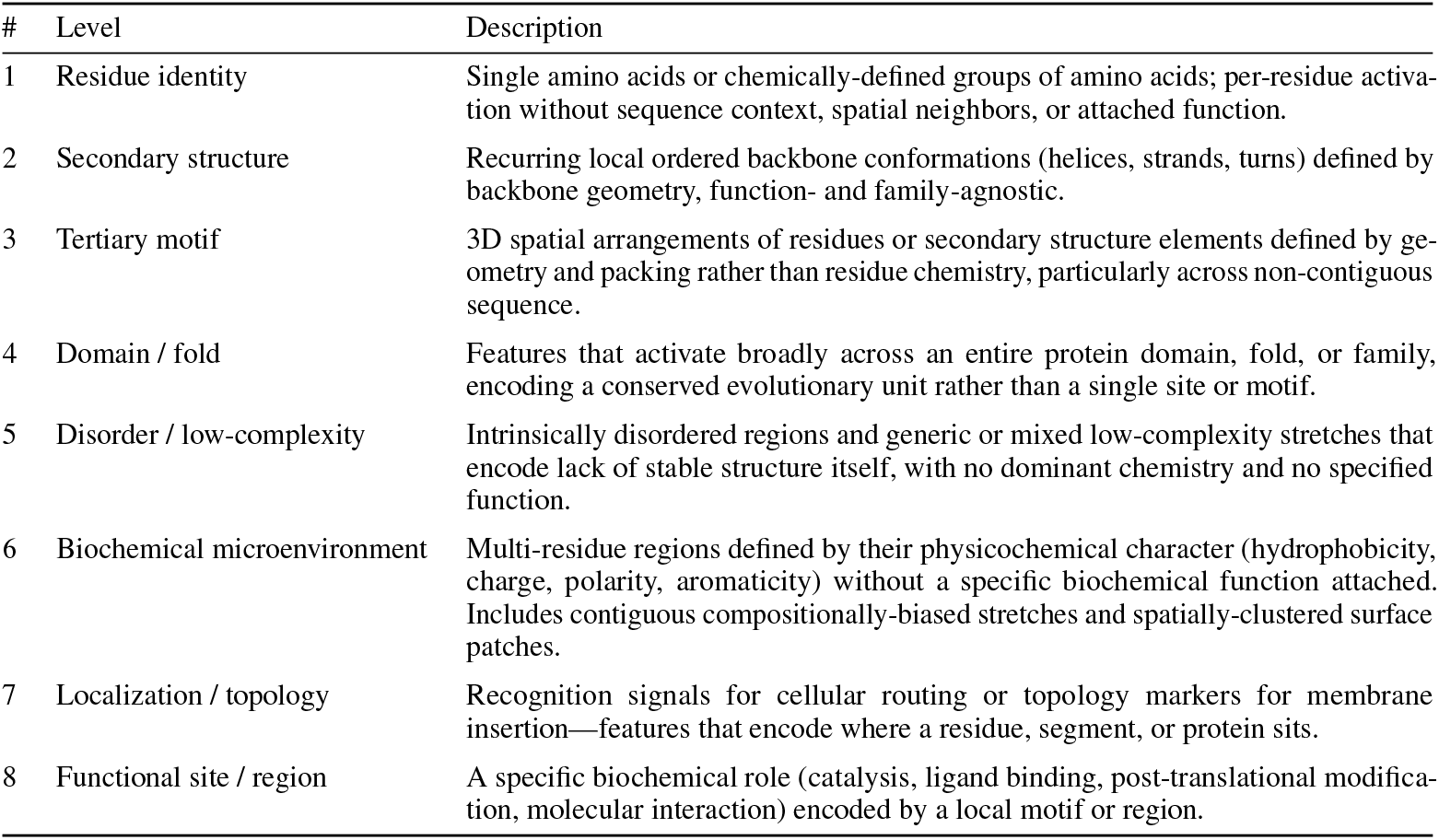
Feature classification categories, ordered from lowest (most universal, least context-dependent) to highest (most specific, most context-dependent) complexity.

| # | Level | Description |
| --- | --- | --- |
| 1 | Residue identity | Single amino acids or chemically-defined groups of amino acids; per-residue activation without sequence context, spatial neighbors, or attached function. |
| 2 | Secondary structure | Recurring local ordered backbone conformations (helices, strands, turns) defined by backbone geometry, function- and family-agnostic. |
| 3 | Tertiary motif | 3D spatial arrangements of residues or secondary structure elements defined by geometry and packing rather than residue chemistry, particularly across non-contiguous sequence. |
| 4 | Domain / fold | Features that activate broadly across an entire protein domain, fold, or family, encoding a conserved evolutionary unit rather than a single site or motif. |
| 5 | Disorder / low-complexity | Intrinsically disordered regions and generic or mixed low-complexity stretches that encode lack of stable structure itself, with no dominant chemistry and no specified function. |
| 6 | Biochemical microenvironment | Multi-residue regions defined by their physicochemical character (hydrophobicity, charge, polarity, aromaticity) without a specific biochemical function attached. Includes contiguous compositionally-biased stretches and spatially-clustered surface patches. |
| 7 | Localization / topology | Recognition signals for cellular routing or topology markers for membrane insertion—features that encode where a residue, segment, or protein sits. |
| 8 | Functional site / region | A specific biochemical role (catalysis, ligand binding, post-translational modification, molecular interaction) encoded by a local motif or region. |

**Figure S29.**
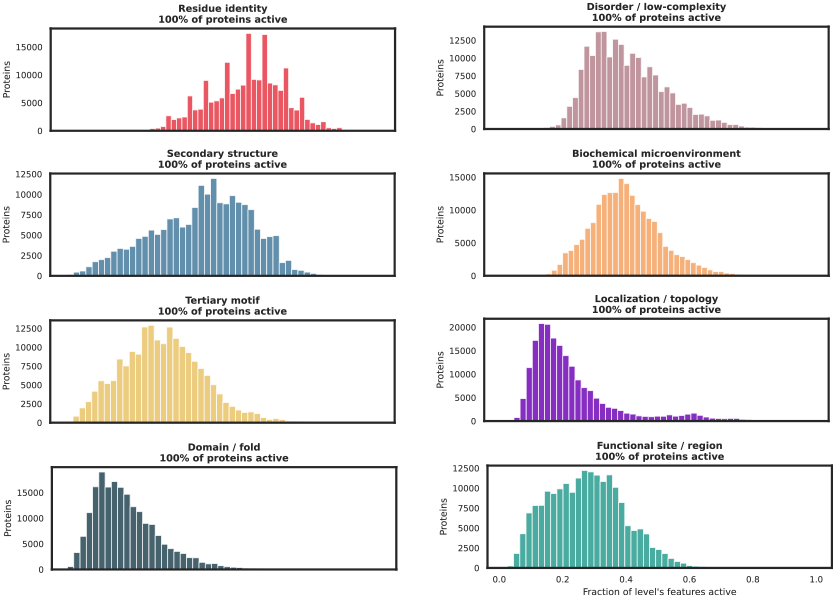
Per-protein usage of features at each complexity level. Histograms of the fraction of the features in each category that are active in each protein in a non-redundant dataset of 195k proteins from Swissprot. Every protein uses at least one feature from every category, illustrating the hierarchical nature of the feature space across levels of complexity.

##### A.4.3.3 Nucleophilic Elbow

One catalytic motif that has arisen several times over the course of evolution is the nucleophilic elbow (71), which consists of a sharp turn sandwiched between secondary structure elements in the catalytic site. In enzymes bearing this motif, the catalytic residue (such as serine, threonine, or cysteine) is present at the “elbow” of this turn and initiates catalysis through nucleophilic attack of the electrophilic substrate.

To identify whether there were features specific to the nucleophilic elbow motif, we used the dataset curated by Du *et al*., 2025, which contains 119 enzymes which perform nucleophilic attack on carbonyl substrates, of which 99 contain the nucleophilic elbow motif. These enzymes span 32 distinct structural folds and a range of enzyme classes, including serine and cysteine proteases but also other hydrolases, transferases, and oxidoreductases that act on diverse substrates. The fact that this motif is conserved across such a diversity of enzymes suggests that it has evolved independently many times, providing an excellent case study in how ESMC learns to represent functional features that are not strictly tied to evolutionary similarities.

To identify candidate features, we compute the SAE features for all proteins in the dataset and compute the precision and recall of each feature for detection of the nucleophile residue relative to other residues in the sequence. The top feature in this search was F6716, which activates directly on the nucleophile on 75 of 99 enzymes (76%; Figure S30), spanning 25 of the 32 unique folds. In an additional four (1a4s, 2esd, 1nba, 1im5), it activates on an annotated oxyanion hole H-bond donor. In 17 of 21 enzymes (81%) which perform similar nucleophilic reactions but without the elbow motif, there is no activation of feature 6716, suggesting it is specifically detecting the structural and chemical environment around the nucleophilic elbow.

Apparent off-target activations tend to occur on catalytically relevant residues in the active site. For example, in proteinase K (1ic6), there is a strong activation on the nucleophilic serine S224 and a smaller peak at the unannotated serine S132. However, the OG–OG distance between S224 and S132 is only 4.2 Å in 3D space, suggesting that the side chains of the two serines are in direct contact in the active site region. Similarly, in fatty acid amide hydrolase (1mt5) and peptide amidase (1ocl), there is no activation on the nucleophile but a single peak on two consecutive serines that are 23 and 24 residues away in sequence but very close in the active site (2.9/5.4 Å and 2.9/5.1 Å from the nucleophile, respectively) and directly involved in the catalytic mechanism according to the Mechanism and Catalytic Site Atlas (141). This reinforces the hypothesis that the feature is capturing important residues that form the catalytic machinery surrounding the nucleophilic elbow rather than a specific sequence-level nucleophile motif.

##### A.4.3.4 Feature Neighborhoods

Each concept in a sparse autoencoder is represented as a direction in the representation space the SAE was trained on. Specifically, the SAE uses encoder and decoder ma-trices 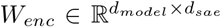 and 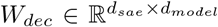 to project the hidden states of the model into the sparse *d*_*dsae*_-dimensional feature space. Each row 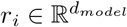 in *W*_*dec*_ represents the direction in the latent space corresponding to feature *i*; that is, the contribution along each dimension of the model’s representation space that is used to reconstruct the representation when feature *i* is active. Features that correspond to similar rows in the decoder matrix therefore correspond to similar directions in the representation space, and one might expect the biological concepts they represent to also be similar.

To visualize feature neighborhoods in this SAE decoder space, we identify all decoder directions that have a cosine similarity of greater than 0.1 with a selected anchor feature (in high-dimensional space, this is significantly denser organization than random chance) and project these *d*_*model*_-dimensional vectors into two dimensions using UMAP (72) with n neighbors = 15 and min dist = 0.1. Since all of the SAEs are trained to reconstruct the same input representations, directions corresponding to features in SAEs with different feature dimensions are directly comparable and can be projected into the same space. All neighborhoods shown here are for the SAEs trained on the activations of layer 60 in the ESMC 6B-parameter model, and we show SAEs with different latent dimensions from 2^13^ − 2^17^. Figure S31 shows the neighborhoods around several features, with one selected from each feature category in Figure 4B. Some neighborhoods, like the ones around F13801 (generic α-helix, i–i+4 positions) and F14444 (hydrophobic assembly/anchoring), are very dense while others are more sparse. Some have a high density of feature that are very tightly clustered: F189 (tryptophans) clusters closely with eight other generic residue-detecting features, while features that correspond to broader residue classes are slightly more separated within the same neighborhood.

##### A.4.3.5 Feature Splitting

Concepts which are represented by single features in SAEs with smaller latent dimension are often represented by several features in larger SAEs, each feature capturing the broader concept in a different context. This phenomenon is known as feature splitting (59). As a case study of feature splitting in our models, we construct a dataset of convergently evolved serine proteases, a diverse enzyme superfamily which share a conserved catalytic mechanism (serine proteases are just one of the protein families that contain the nucleophilic elbow motif from A.4.3.3). This dataset is split based on protein family: the training dataset consists of 332 enzyme sequences from the S1 peptidase family, and the evaluation dataset consists of 60 sequences from the S8 peptidase family. These two families both use the characteristic catalytic triad of serine, histidine, and aspartic acid to perform hydrolysis but in a different order, and the proteins have significant differences in overall fold and active site structure. Because of this divergence, a feature which activates on both catalytic sites is not simply recognizing a sequence or fold, but has captured a concept related to the potential for a specific catalytic chemistry.

Features activating on catalytic residue positions in the S1 family were identified using the differential activation on catalytic residues compared with non-catalytic residues in all S1 sequences, using the SAEs trained on layer 60 of ESMC 6B with the smallest (2^13^) and largest (2^17^) latent dimensions. Cases of feature splitting are identified by isolating features that fire on catalytic residues from *both* families at a small dimensionality, and *each family independently* at a large dimensionality. Figure S32 shows an example where a single feature activates on the catalytic triad in both S1 and S8 families in the smallest SAE (F6960), while the larger SAE model uses separate features to represent the active site in the two families (F77290 and F109350).

**Figure S30.**
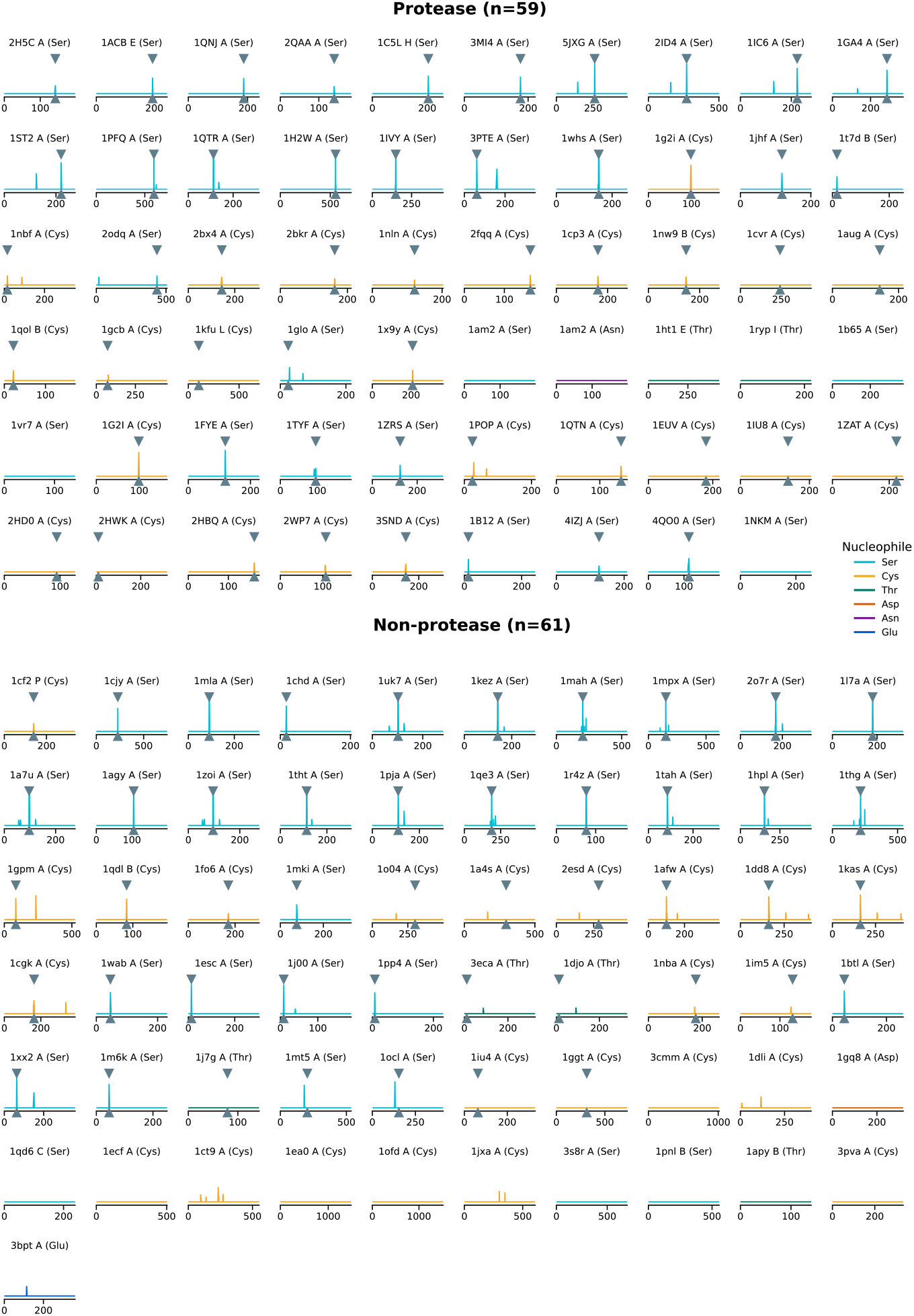
Activation of feature 6716 on nucleophilic enzymes from Du *et al*. 2025. The activation of F6716 is shown along the sequence dimension for each protein, with the color of the line given by the nucleophilic residue. The nucleophilic residues are annotated with triangles for enzymes that possess the nucleophilic elbow motif. The feature shows high specificity for the nucleophilic elbow across diverse nucleophiles and both protease and non-protease enzymes.

**Figure S31.**
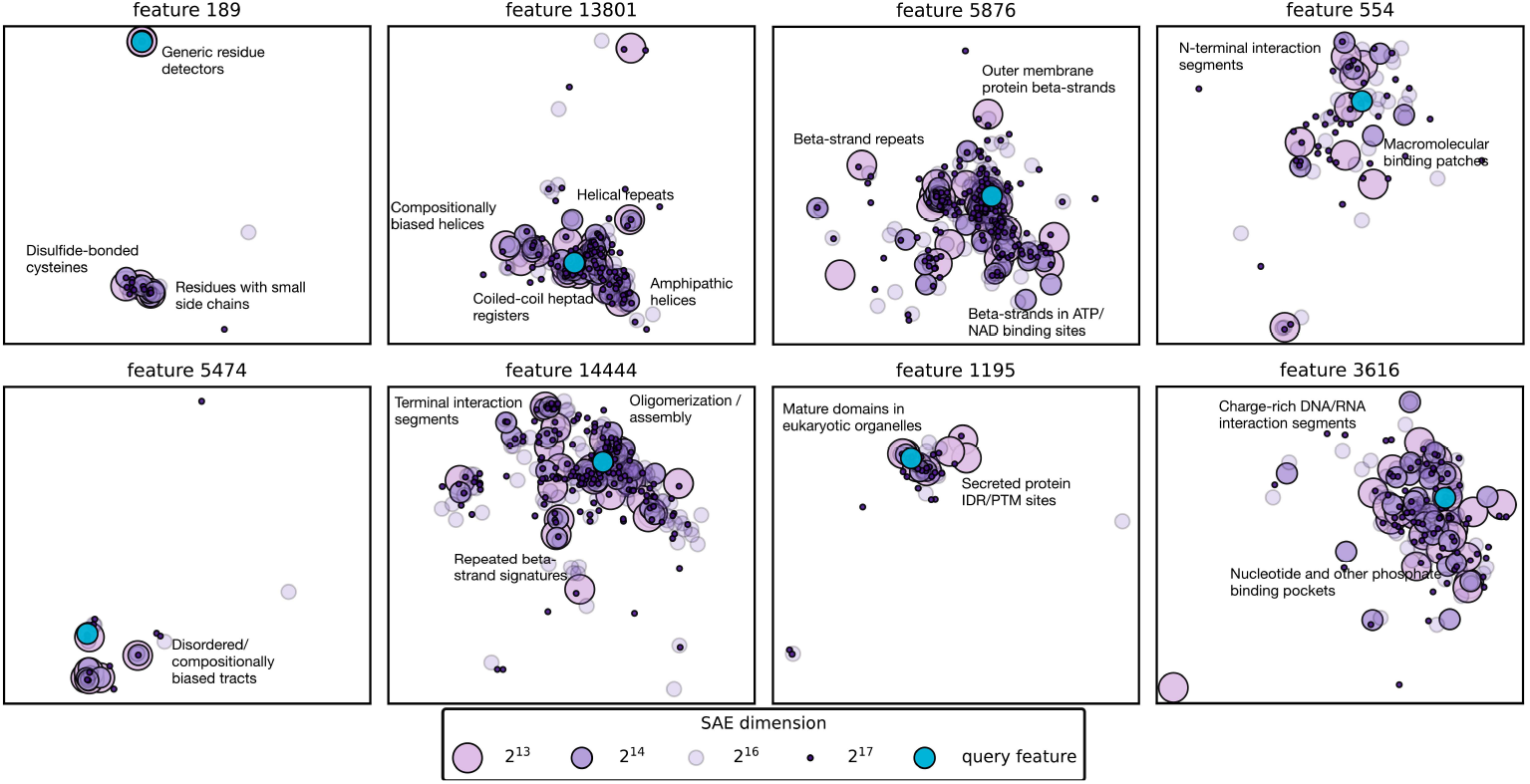
Selected feature neighborhoods. Feature neighborhoods constructed from the SAE decoder directions for eight example features, one from each category in Figure 4. Features from SAEs with different latent dimensions are shown as different point sizes, with larger points representing features from smaller-dimensional feature spaces. In larger feature spaces, more features are used to represent similar concepts.

##### A.4.3.6 Other Selected Sae Features

Figure S33 shows 50 additional examples of SAE features selected from the ESMC 6B layer 60 SAE with a 16,384-feature codebook. These features are arranged by the type of functional concept they encode: local structural motifs, local sequence motifs, domain-specific signatures, repeats, catalytic motifs, membrane-associated patterns, ligand-binding sites, post-translational modification sites, molecular interaction sites, or regions associated with intrinsic disorder. The activation of some features capture meaningful concepts across different chains or subunits within a macromolecular complex. Figure S34 shows three such examples, visualized on the multi-chain PDB structures. In these cases, the features capture concepts in which the functional site or region is defined by the interaction with other molecules or subunits.

#### A.4.4 Compositionality of SAE Features

##### A.4.4.1 Kinase Compositional Grammar

We use eukaryotic protein kinases to showcase how SAE features compose together to represent complex functions because while they are one of the largest and most diverse enzyme families in biology, their core catalytic machinery is highly conserved. We use this to highlight how different kinase subfamilies share key functional features and how they differ in key ways which reflect their specific biological roles. For this analysis, we choose a representative selection of kinases from each of the seven major kinase families and computed SAE features based on the sequence retrieved from UniProt:

- *Calcium/calmodulin-dependent kinase (CAMK):* CAMK1 (Q14012), CAMK2 (Q13554), CAMK4 (Q16566), DAPK1 (P53355), MARK2 (Q7KZI7)
- *PKA/PKG/PKC kinase (AGC):* PRKACA (P17612), AKT1 (P31749), PRKCA (P17252), RPS6KB1 (P23443), SGK1 (O00141)
- *Tyrosine kinase (TK):* SRC (P12931), EGFR (P00533), ABL1 (P00519), JAK2 (O60674), MET (P08581)
- *Tyrosine kinase-like (TKL):* TGFBR1 (P36897), BMPR1A (P36894), BRAF (P15056), RAF1 (P04049), IRAK4 (Q9NWZ3)
- *Casein kinase 1 (CK1):* CSNK1A1 (P48729), CSNK1D (P48730), CSNK1E (P49674), CSNK1G1 (Q9HCP0), TTBK1 (Q5TCY1)
- *STE kinase (STE):* PAK1 (Q13153), STK4 (Q13216), PAK4 (O96013), MAP2K1 (Q02750), MAP3K1 (Q13233)
- *CMGC kinase (CMGC):* CDK1 (P06493), CDK2 (P24941), MAPK1 (P28482), MAPK3 (P27361), GSK3B (P49840)

**Figure S32.**
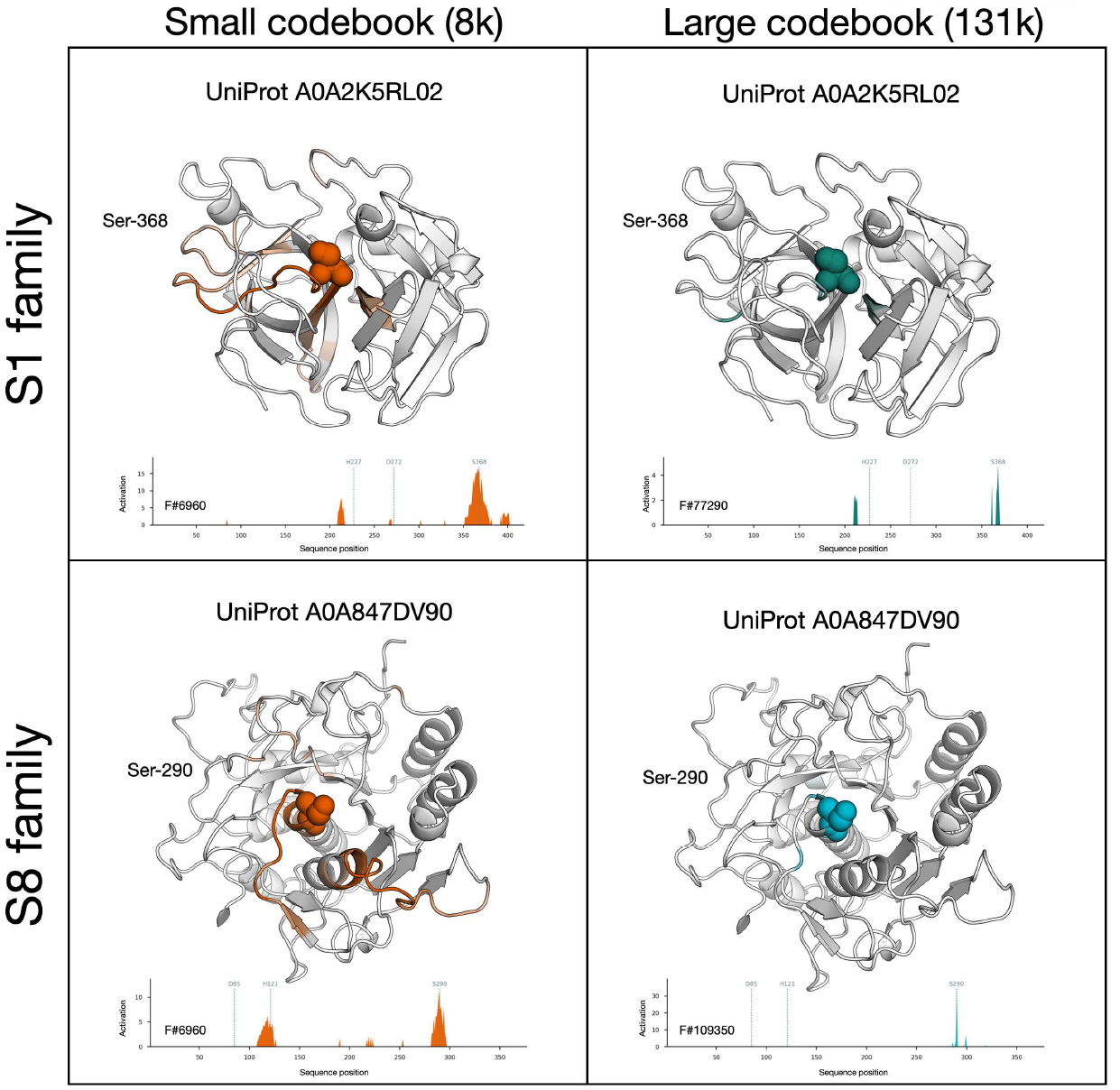
Feature splitting in serine protease SAE features. Feature splitting example in S1 and S8 proteases. Top panels: S1 protease UniProt A0A2K5RL02 showing the catalytic nucleophile in spheres. Bottom panels: S8 protease UniProt A0A847DV90 showing the catalytic nucleophile in spheres. Note that the two enzymes do not share a fold. Left panels: at small codebook size, the model uses feature F6960 which strongly activates on both families’ nucleophilic catalytic residue. Right panels: at large codebook size, the model uses distinct features to represent the active site nucleophile of S1 proteases and S8 proteases, namely for S1 the feature is F77290, and for S8 the feature is F109350.

**Figure S33.**
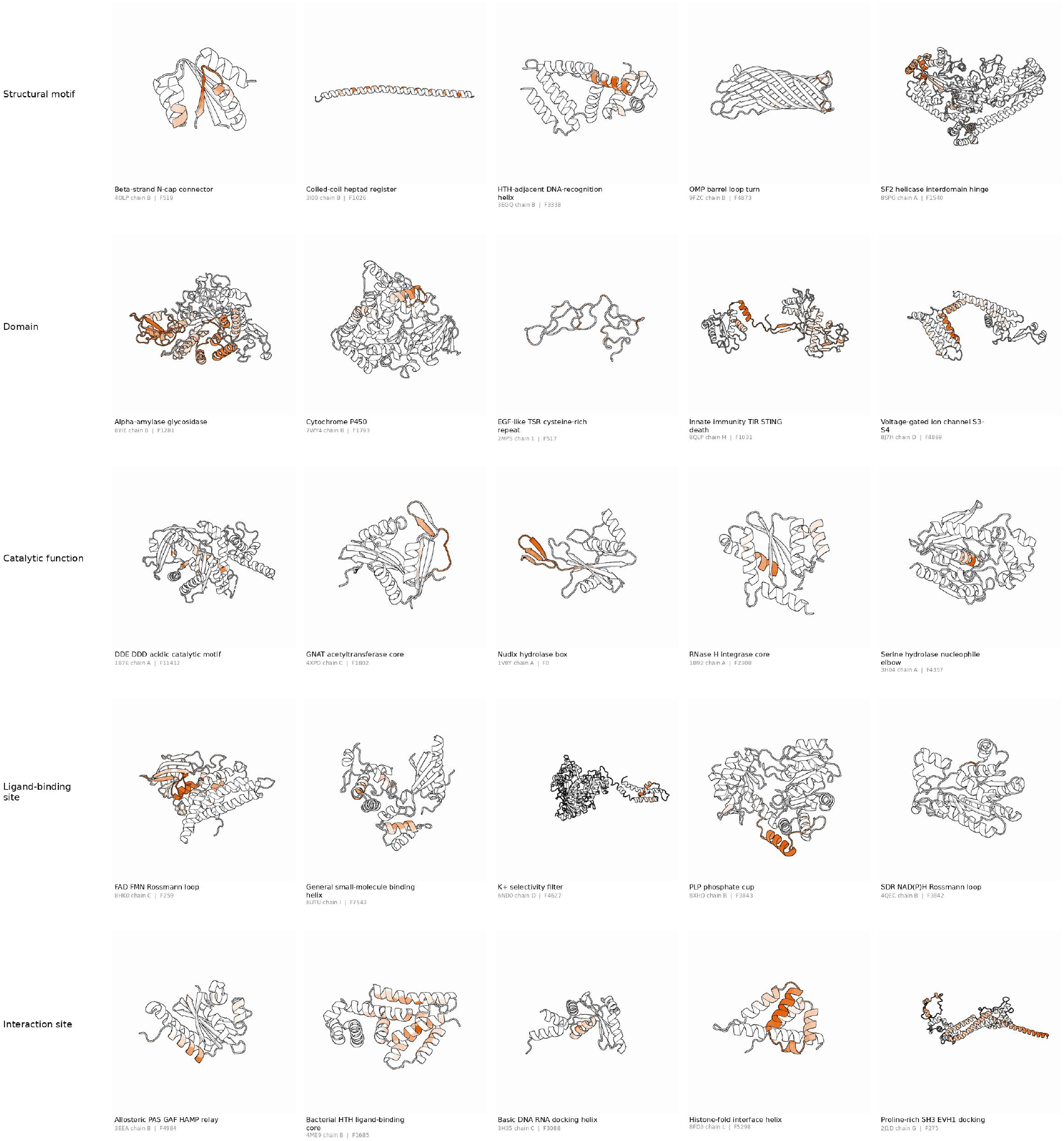
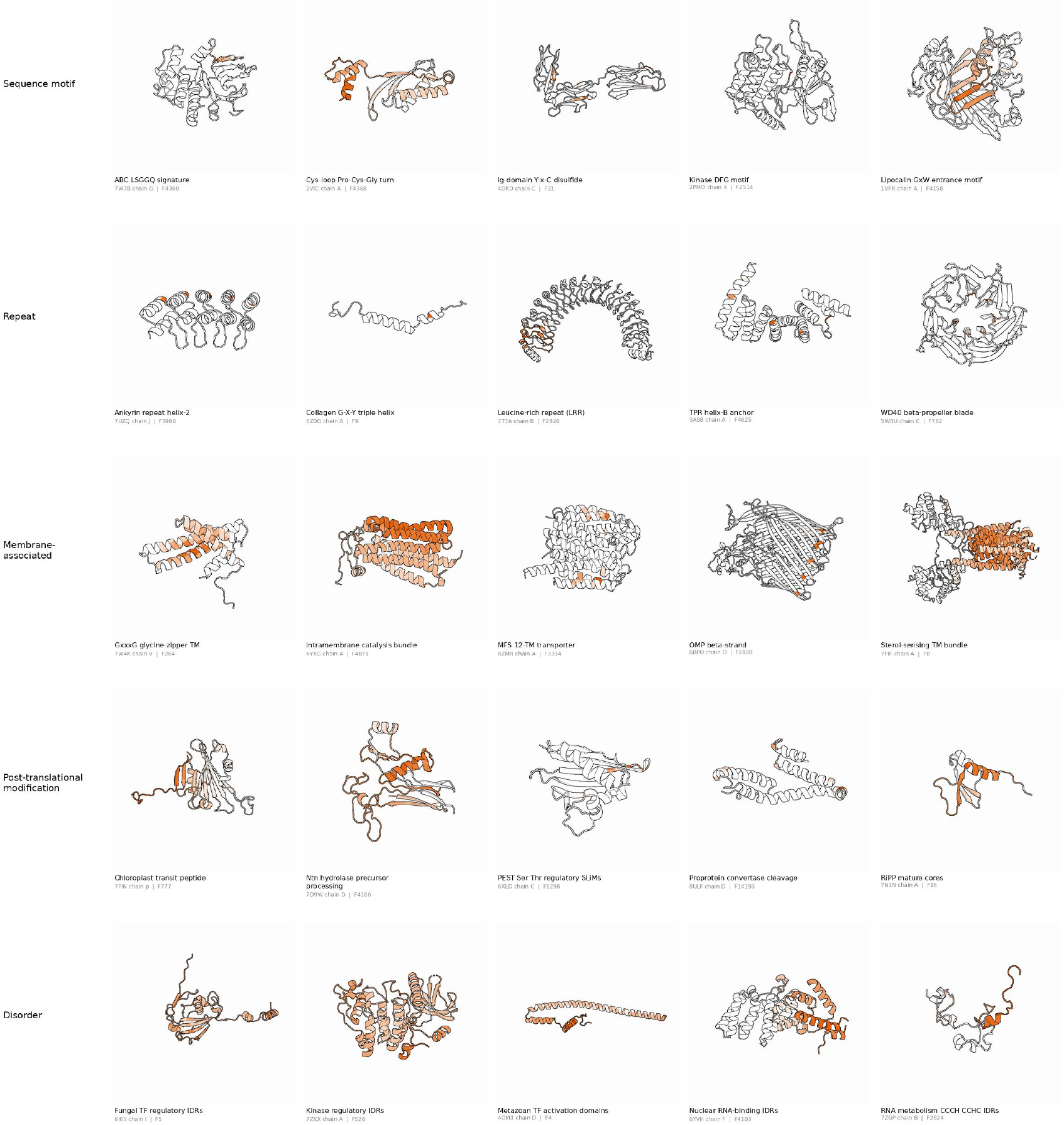
Gallery of 50 SAE features, grouped into 10 biological categories. For each feature, one representative PDB chain is rendered as a cartoon colored by per-residue activation (orange). Activations localize to catalytic triads, ligand-binding loops, transmembrane segments, sequence motifs, and intrinsically disordered tracts.

**Figure S34.**
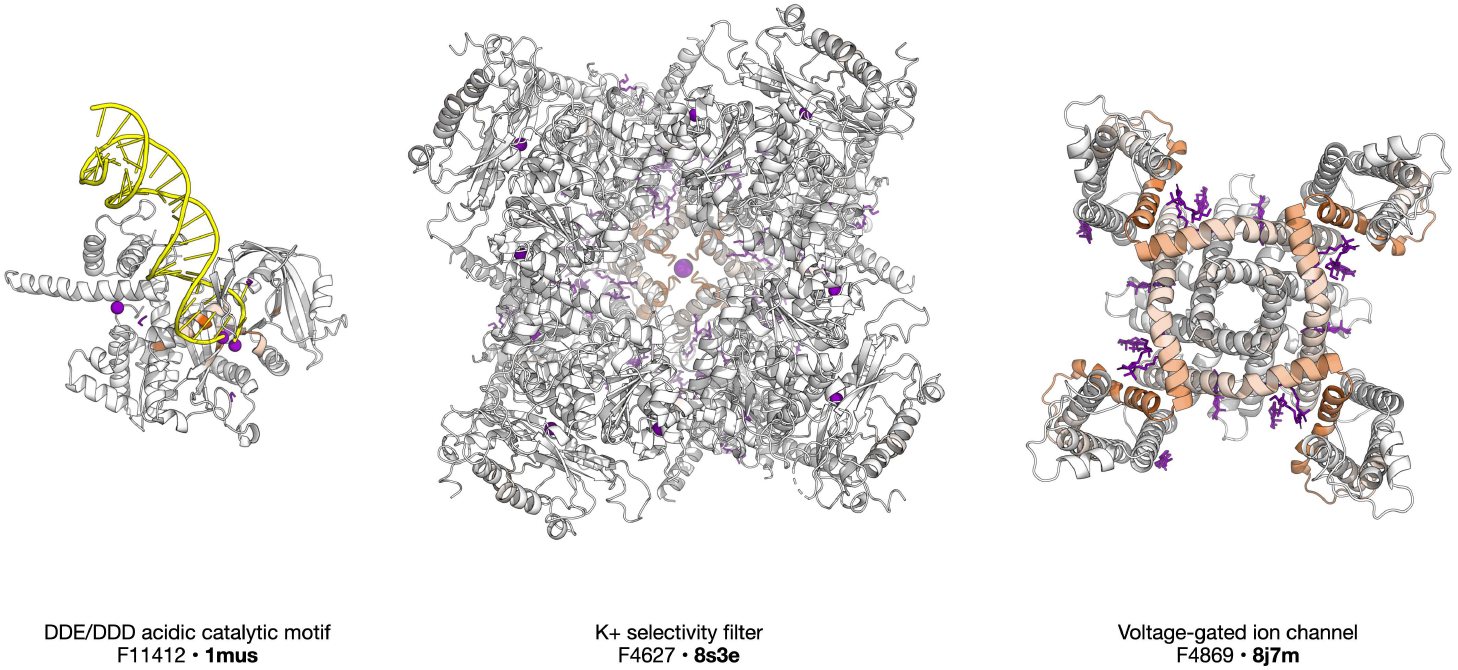
Selected feature activations in molecular complexes. From left, F11412 activates on the DDE/DDD acidic catalytic motif in 1mus. In the middle panel, F4627 (K+ selectivity filter), fires on the key residues of 8s3e that form a selective pore region. At right, voltage-gated ion channel 8j7m is shown with the activation of F4869 (voltage-sensor/pore module of 6-TM cation channels).

To select the subset of kinase-related features for this analysis, we filtered all 16,384 features down to only those which contain the string “kinase” anywhere in their descriptions (either in the feature summary or as an exemplar). For each kinase family, we then identify all of the features in this kinase-related subset that activate on *all members* of the family. Finally, we classify each feature by how many families it is associated with: a feature is considered “kinase-universal” if it is found in all seven families, and it is considered “family-specific” if it is found in only one. This resulted in 531 features found in all kinases, and 254 family-specific features (10 CAMK, 42 AGC, 51 TK, 24 TKL, 8 STE, 32 CMGC, 87 CK1).

We select CAMK1 (UniProt Q14012) as the central example as it is a canonical kinase with a monomeric structure. We identify the key catalytic regions in the structure—the phosphate-binding P-loop, the *α*C-helix, the HRD and DFG motifs, the activation segment, and the autoinhibitory segment. The density of kinase-universal features is computed in each region as the total number of active features divided by the number of residues in the region and reported in the legend of Figure 4E. A single family-specific feature is chosen for representatives of the TK, AGC, TKL, and CK1 families.

##### A.4.4.2 Fitness Landscapes AND Mutation Sensitivity

We investigate how SAE features represent protein functional activity landscapes using 25 deep mutational scanning (DMS) datasets from ProteinGym (142) which measure the function of enzymes and a few transporter proteins (ignoring non-functional readouts such as stability). Linear probing experiments suggest that a compact subset of SAE features (median 79 per dataset; range 11–288) is sufficient to predict the local fitness landscape with high accuracy (mean test Spearman *ρ* = 0.63, range 0.27–0.83). Many of the selected features activate in mechanistically important regions and their descriptions specifically reference core biological machinery. For some features, the change in maximum activation magnitude is correlated with changes in protein activity due to mutations both at the site of the activation and many residues away. Similar patterns are observed across datasets, but for illustration purposes we highlight the findings for c-Src kinase, a dataset consisting of 3,372 mutations in the kinase domain (143) for consistency with Figure 4E, A.4.4.1.

###### SAE Features Predict Fitness in DMS Data

For each variant in the DMS dataset, we extract the layer 60 representation of ESMC 6B and compute (1) the max-pooled 16, 384-dimensional sparse vector of SAE features, which have ∼1,500–5,000 non-zero features per dataset, as well as (2) the mean-pooled 2560-dimensional layer 60 representation itself.

ElasticNet regression with L1 ratio *α*_elastic_ = 0.6 is used to select the subset of features that are predictive of DMS function score. A separate Ridge regression (*α*_ridge_ = 1000) is then fit on the function-selected subset, with per-feature z-score standardization (computed on the training fold), to identify feature-specific contributions. We conduct a sweep on *α*_elastic_, which controls the sparsity of the ElasticNet solution, to estimate the performance of different sizes of SAE feature subset. Following ProteinGym’s standard 5-fold random split, each outer fold sweeps *α*_elastic_ and chooses the value whose ElasticNet-selected features give the best downstream Ridge Spearman under 5-fold inner CV on the training set; we then refit on the outer training split and report mean number of features and mean test Spearman *ρ* across folds for each *α*_elastic_ (Figure S35A). As baseline, we apply the identical pipeline to mean-pooled ESMC embeddings from the same layer.

Fitness prediction performance on Src kinase rises steeply within a few hundred features (Figure S35A) and then plateaus around *ρ* ≈ 0.74, indicating that information encoding the local functional landscape around the wild-type protein is captured by a small fraction of all SAE features active across the DMS dataset (*n* = 3, 805).

To further prioritize the most functionally relevant features, the five fold-specific sparse models are collapsed into a single robust feature set. We choose one *α*_elastic_ that maximizes the median sparsity across folds, such that each fold is at least 95% of the optimum, and retain features present in four out of five folds that cover 70% of cumulative coefficient weight. For Src kinase this results in a set of 108 robust SAE features whose activations are predictive of activity. Each of the 108 features had the same coefficient sign across all cross-validation splits. Fitting a ridge regression on those 108 features alone gives *ρ* = 0.74, showing that this subspace contains most of the predictive signal (Figure S35A).

###### Predictive features are mechanistically interpretable

Figure S35B-C plot the predicted structure of Src kinase alongside the wild-type per-position activations for seven representative features from the 108-feature subset. Key catalytic regions and regulatory features from SwissProt annotations are also shown along with the activations. The features cover all levels of biological specificity, from generic residue detectors (F2319: threonines; F12861 aromatics) to kinase-specific catalytic features that activate on individual residues (F847: kinase helix-to-catalytic-loop junction; F3809: kinase catalytic core residues) to broad regions (F8785: kinase activation segment). The residue detectors F2319 and F12861 are generically active across proteins, but they peak at strongly conserved functional residues in the activation segment: F12861 peaks at Y419, the canonical autophosphorylation site, and F1635 activates on both of the major glycine-rich flexible loops that are critical for kinase function (DFG, P-loop). The description of the more specific features closely matches the corresponding element of the kinase regulatory machinery.

###### Feature activations are sensitive to mutations and often correlate with fitness

The absolute changes of the normalized SAE feature activations are highly variable, from small but significant changes to turning the feature off completely. Z-score standardization on the training set controls for the difference in feature magnitudes and changes across the mutation landscape. This enables the linear model to extract meaningful fitness signal from each feature’s max-pooled activations.

Comparing the maximum activation of each feature to the measured DMS fitness across all 3,372 mutations of the dataset, we find that 57% of the 108 features have a non-trivial Spearman correlation (|*ρ*| *>* 0.1). The largest positive spearman correlation is observed for F1635 (glycine-rich loops; +0.35), and the largest negative for F847 (catalytic loop junction; −0.24). This matches the sign of the respective ridge coefficients. A further 6% of features have a purely binary signal: half are not active in WT but are activated in mutants with no kinase activity, and half activate in wild-type but turn off completely in non-functional mutants. The remaining 37% have no individual correlation and contribute only through combination with other features. The linear model combines the localized functional signals of all 108 features to reach *ρ* = 0.74.

The above analysis considers only the univariate relationship between max-pooled activation and activity, but does not resolve how much a feature changes locally in response to mutations. To investigate this, we compute the position-wise activations for every mutant and define the per-mutation *peak sensitivity* 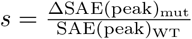. This reveals that some features respond to mutations only in their local context, while others are sensitive to mutations far away.

Local residue detectors like F2319 and F12861 are useful for the linear fitness model because they peak at functionally important positions. Mutations at these peak positions drops the respective feature to zero locally (*s* = −1), but mutations away from the peak do not affect activation. That signal is not seen in the max-pooled representation as the feature value there falls back to the next-highest position: for example, the max-pooled F2319 activation only drops 11% to the next threonine in the sequence.

The activation of features such as F278 and F1635, which both activate on flexible loops, is sensitive to mutations at the peak position and at neighboring positions, with the magnitude of response often correlating with the mutation’s fitness effect. Figure S35D,E illustrates this by plotting *s* along the sequence for F278 (peak at K275 in the P-loop) and F1635 (peak at G409 in the DFG motif), with each substitution colored by its measured kinase activity. The feature activation at its wild-type peak drops by up to *s* = −0.35, and close to the peak the sign of *s* mirrors the sign of the fitness effect. At some neighboring positions the feature activation closely reflects the effect of individual subsitutions on fitness — at G277, where substitution reduces F278 activation by 9–22%, the sensitivity *s* is highly correlated with measured kinase activity (within-position *ρ* = 0.73). This is much higher than the F278’s max-pooled univariate *ρ* = 0.25 across all mutations; focusing on nearby positions reduces noise from mutations elsewhere in the protein. In these cases, the activation of the feature appears to reflects the integrity of the local neighborhood that defines the functional motif.

**Figure S35.**
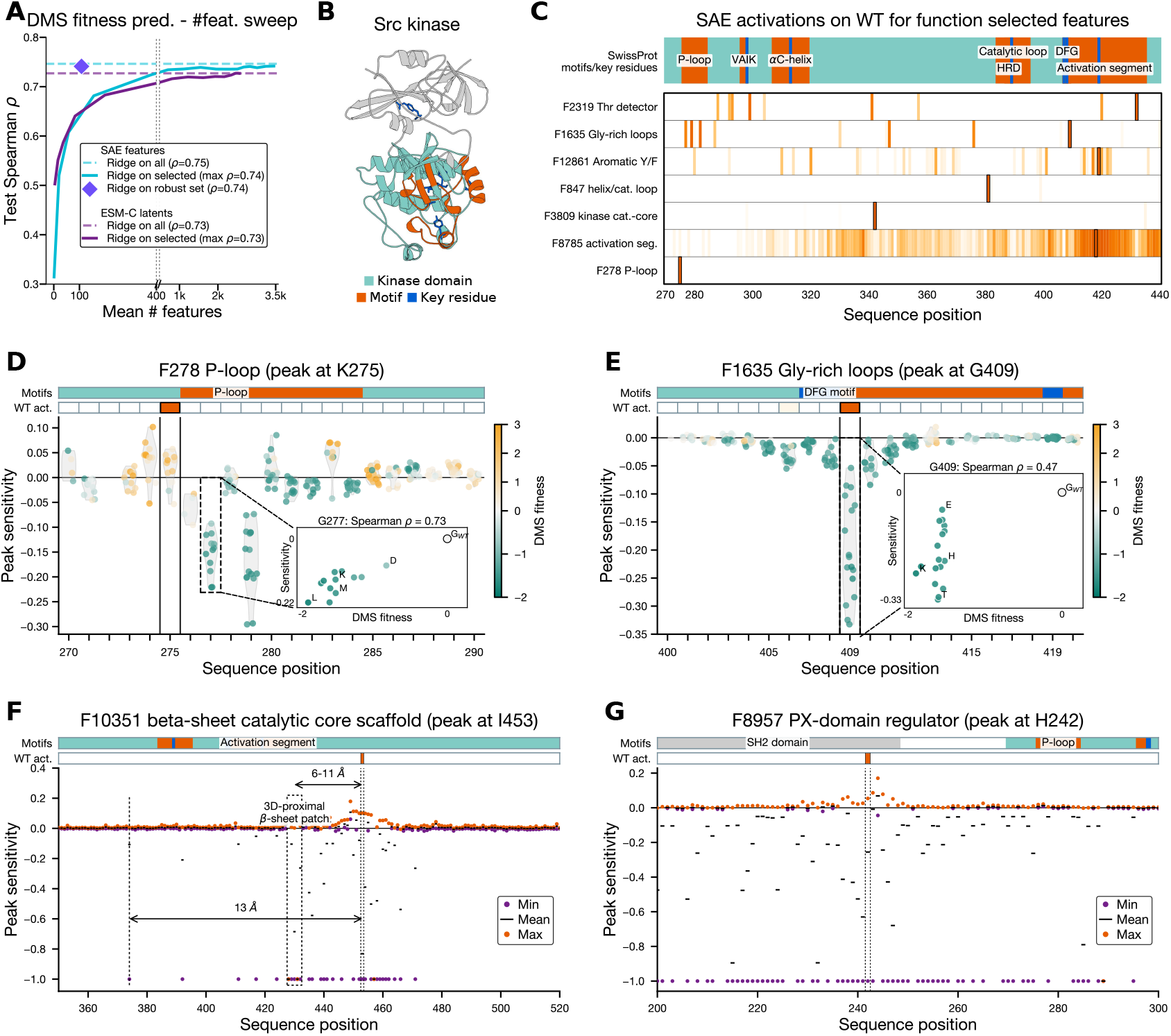
A compact set of SAE feature activations predict Src kinase activity and localizes to the kinase regulatory machinery. (A) Test Spearman *ρ* of a simple Ridge model as a function of the size of the function-selected feature subspace, mean across 5-fold random split CV, for the SAE feature space and the mean-pooled ESM-C embedding baseline. The x-axis is broken at 400; positions **>** 400 are compressed. The diamond marker indicates the 108-feature robust set of features investigated in the rest of the figure. (B) Src kinase structure with key functional motifs and regulatory residues highlighted. (C) Wild-type per-position activation of seven features from the selected set across the kinase domain, with key motifs shown along the top. Each row is one feature, normalized by its peak activation (boxed cell). (D,E) Key feature activations rank correlates with fitness. Per-mutation feature *peak sensitivity* (ΔSAE(peak)_mut_*/*SAE(peak)_WT_) at each sequence position are shown as dots, with each dot representing a variant colored by its measured kinase activity. (F,G) Some feature activations are sensitive to mutations many residues away. Minimum, mean, and maximum peak sensitivity across the 19 substitutions at each position is shown. For many positions there are mutations that reduce the activation to zero. For F10351, the 3D-proximal *β*-sheet patch is highlighted, along with A374.

###### Features can be sensitive to distant mutations

Some feature activations are sensitive to mutations many residues away. Figure S35F,G illustrate two examples, showing the min, mean, and max peak sensitivity *s* at each position across the 19 amino-acid substitutions. F10351 is a structural feature described as “conserved beta-strand/loop elements in beta-sheet-dominated catalytic cores”, and it peaks at I453. The activation of this feature is reduced to zero by many mutations that are clustered in 3D but scattered along the sequence, with some up to 79 residues away in sequence from I453. However, in structure space these positions are close together, with median *Cα* distances of 11 A◦, suggesting that the feature is sensitive to changes in the surrounding structural environment. In another case, F8957, which peaks at H242 in the SH2 regulatory domain, is highly sensitive to mutations not only in the same domain but also in the kinase domain over a range of over 100 residues, many of which are structurally far away. The signal that this feature is capturing is not clear, but it is possible that it captures changes in the allosteric regulation of the catalytic domain by the SH2 domain.

##### A.4.4.3 Protein-Protein Interaction Features

Patterns in SAE feature co-occurrence across proteins are predictive of known protein-protein interactions (PPIs), despite ESMC being trained only with the context of a single chain at a time. By analyzing how pairs of SAE features co-occur across known PPI interfaces in the PDB, we develop a method for predicting PPIs and find that features that co-occur more often than expected by random chance at interaction interfaces often correspond to known interaction motifs. For example, in a TCR-pMHC complex, the method’s prediction is largely driven by the activation of F14846 (MHC-like beta sandwich) and F12376 (Ig-like domains) (Figure S36). On a held-out test set, the method achieves an AUROC of 0.855. As a comparison, we fold the protein pairs using ESMFold2 and use ipTM as a predicted interaction score. The folding method achieves an AUROC of 0.828, which is similar to the SAE-based approach (*p* = 0.64). These results suggest that information about protein-protein interaction is encoded in the ESMC world model, and is accessible through both the folding model and sparse representations.

**Figure S36.**
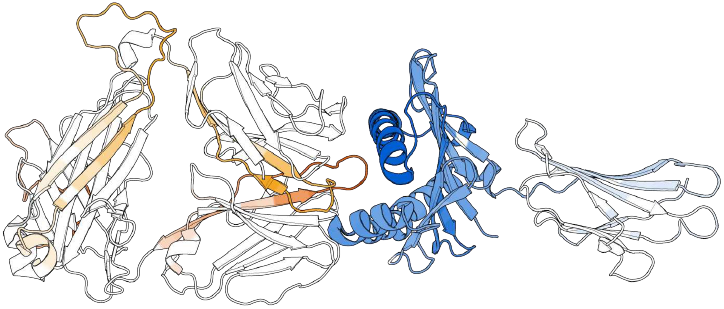
PPI feature co-occurrence visualization. Visualization of PDB 6MTM, chains A, D, and E. F14846 is visualized in blue on chain A (MHC-I) and detects “MHC/Ig-superfamily– like beta-sandwich folds”. F12376 is visualized in orange on chains D and E (B2M; different shades of orange distinguish the two chains) and detects “the conserved beta-strand framework of immunoglobulin-like domains.” Darker color intensity indicates higher activation.

###### Learning feature co-activation patterns from PPI3D

We obtained a dataset of 140,276 protein-protein interfaces (representing 52,650 unique PDBs) by downloading interface data from PPI3D (144) clustered at 40% sequence identity. For each interface, we took the protein sequences, removed any unresolved residues at the beginning or end of the sequence, and computed IDF-normalized, per-residue SAE features. Features with IDF-normalized activations *>* 0.5 were considered active.

All SAE feature pairs that co-occur at contacting residues in each interface were identified and the proportion of contacts at which each pair co-occurs were computed. For each feature pair (*i, j*), we define:

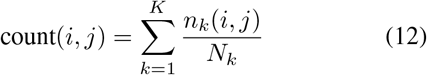

where *K* is the number of total interfaces in the dataset, *n*_*k*_(*i, j*) is the number of contacts (between two protein chains) in interface *k* where features *i* and *j* co-occur, and *N*_*k*_ is the total number of contacts in interface *k*. This normalization ensures that each interface contributes equally, regardless of size.

Normalized pointwise mutual information (NPMI) is used to measure the degree to which features co-occur with each other over expected. NPMI is computed based on the joint probability of the two features with smoothing:

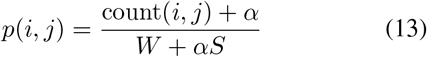

where *W* is the total sum of all weights, *S* is the number of observed feature pairs and *α* is a smoothing parameter (set to 0.1). We also compute the marginal probabilities with smoothing:

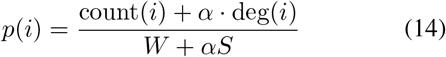

where count(*i*) is the sum of count(*i, j*) across all features *j*, and deg(*i*) is the number of unique partner features with which feature *i* co-occurs.

The PMI is then calculated as

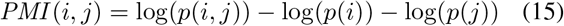

and NPMI as

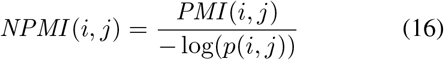

The NPMI ranges from −1 to +1, with higher values indicating features that co-occur more frequently within interfaces. The final NPMI lookup table contained ∼1 million feature pairs with NPMI *>* 0.

###### Scoring algorithm

For a query pair of protein sequences and their SAE features, a feature is considered to be active if its IDF-normalized activation is greater than 0.5. For each residue, the top-*k* active features with the highest activation magnitudes are extracted.

Residue pairs are scored by looking up NPMI values in the precomputed table. For each residue *i* in protein *A* and residue *j* in protein *B*, we compute:

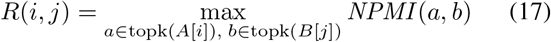

where topk(*A*[*i*]) are the top-*k* most active features for residue *i* in protein *A* and topk(*B*[*j*]) are the top-*k* most active features for residue *j* in protein *B*. This identifies the maximum NPMI across all feature combinations between the two residues.

We then select the top-*T* highest-scoring residue pairs, where:

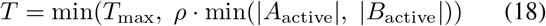

where |*A*_active_| and |*B*_active_| are the number of residues with at least one active feature in proteins *A* and *B*, respectively.

The final interaction score is the mean NPMI of the selected top-*T* residue pairs.

###### Hyperparameter Tuning

Three hyperparameters were tuned on the PPI3D training set:

- top_*k*_ : Number of top features per residue (values: [3, 5, 7, 9, 11, 13, 15])
- *ρ*: Fraction of smaller protein’s active residues to consider (values: [0.05, 0.10, 0.15, 0.20, 0.25])
- *T*_max_: Maximum number of residue pairs to score (values: [30, 40, 50, 60, 70])

The optimal hyperparameters, based on the greatest AUROC in the training set, were top_*k*_ = 3, *ρ* = 0.25, *T*_max_ = 70.

###### Results

We evaluated our method on the PPI prediction benchmark set from Zhang *et al*. (145). The benchmark was created by taking a randomly sampled subset of 3, 000 positive interactions from the intersection of STRING PPIs (score *>* 700), UniProt PPIs, and high-confidence BioGRID PPIs (multi-validated). The benchmark set also contains 30, 000 negative pairs that were randomly sampled with exclusion of known interactions: pairs with evidence from UniProt, BioGRID, or STRING were removed, along with homologs of known interacting pairs. We filtered this benchmark to remove any sequence with more than 30% sequence identity (using MMseqs2) to any sequence in the PPI3D training set. After this filtering, the final test set contains 778 positive pairs and 12, 355 negative pairs. The SAE-based method has an AUROC of 0.855 and an AUPRC of 0.467.

##### A.4.4.4 Feature Space Factorization

###### Non-negative matrix factorization on sparse feature representations

We use non-negative matrix factorization (NMF) with a Bernoulli–Poisson link (146) to factor SAE data into latent topics that represent biologically coherent topics. Each protein is represented by a binary vector representing the presence or absence of each SAE feature across all residues. The latent Poisson variable represents the number of residues on which a given feature is active. We only observe the Bernoulli variable: whether the feature is active in the protein (i.e. the latent Poisson is ≥ 1) or not (i.e. the latent Poisson is 0).

Let us denote *N* := number of proteins, *M* := number of SAE features, and *T* := number of topics. Under our parameterization, we represent each element *X*_*ij*_ ∼ Bernoulli(*P*_*ij*_) of our binary data matrix ***X*** ∈ ℝ^*N*×*M*^ as a draw from a Bernoulli random variable *P*_*ij*_ = ℙ[*Z*_*ij*_ ≥ 1] whose success probability equals the probability that the underlying latent Poisson random variable *Z*_*ij*_ ∼ Poisson(Λ_*ij*_), is non-zero.

In our formulation of NMF, the underlying Poisson parameter matrix Λ is decomposed into non-negative matrices ***W*** (with shape *N* × *T*) and ***H*** (with shape *T* × *M*) and a feature-specific bias column vector ***b*** (with shape *M* × 1):

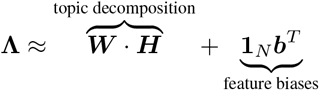

***W*** and ***H*** are initialized with uniform random noise between 0 and *q*_*W H*_, where *q*_*W H*_ is chosen such that the expected value of each (*WH*)_*ij*_ is approximately proportional to the density *p*_*W H*_ of nonzero features in X:

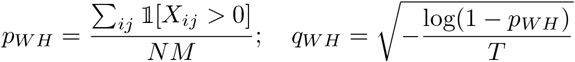

We initialize ***b*** such that each 1 − exp(−*b*_*j*_) equals the marginal frequency (*p*_*b*_)_*j*_ of each feature *j* in ***X***.

***W*** and ***H*** are then updated using a set of multiplicative update rules (147), modified for our Bernoulli–Poisson formulation so that at convergence, ***W*** and ***H*** minimize the total negative log-likelihood of observing *X*_*ij*_:

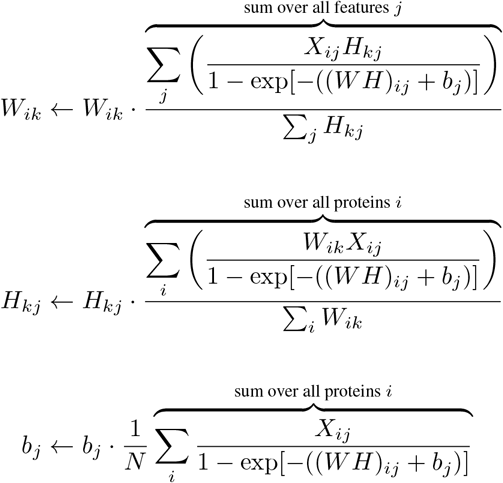

For each ***H*** update, we derive ***W*** _*batch*_ (from random initialization) using the current ***H*** and 30 iterations of the multiplicative update rule for ***W***. For the ***H*** update, the terms in the numerator and denominator are accumulated separately across all proteins in ***X***, then an update is performed with the accumulated terms.

In order to jointly optimize the NMF for both the negative log-likelihood and internal biological coherence of the topics, a second step is added after each H update to improve the biological coherence of the topics. First, the shifted, positive pointwise mutual information (SPPMI) (148) is computed between every pair of features. For a pair of features *i, j*, this value is computed as SPPMI(i, j) = max(0, PMI(*i, j*) − 1).

For each topic *k* and feature *j*, we compute a coherence score over all other features using the SPPMI matrix 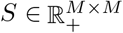

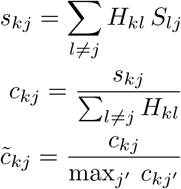

and update the weight of feature *j* within topic *k*:

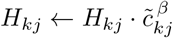

where *β* ≥ 0 is a hyperparameter that controls the filter strength. We use *β* = 0.1, which empirically balances biologically coherent topics with good reconstruction. The rows of *H* are normalized at the end of each epoch so that they sum to 1 to keep *H* bounded and makes topics more interpretable.

###### Feature combinations in reference proteomes

To compute the feature combinations (topics) that form a basis for representing a diverse space of protein functions, we use the protein sequences from 81 Quest for Orthologs (QfO) (74) reference proteomes (UniProt release 2025 04), comprising 7 Archaea, 23 Bacteria, and 51 Eukaryota species (1, 058, 822 proteins total).

For each protein, we computed binary SAE feature vectors by max-pooling activations across residues and thresholding at zero (presence/absence). The resulting matrix (1,058,822 proteins × 16,384 features) served as input to an NMF trained for a maximum of 200 multiplicative updates of ***H***.

**Figure S37.**
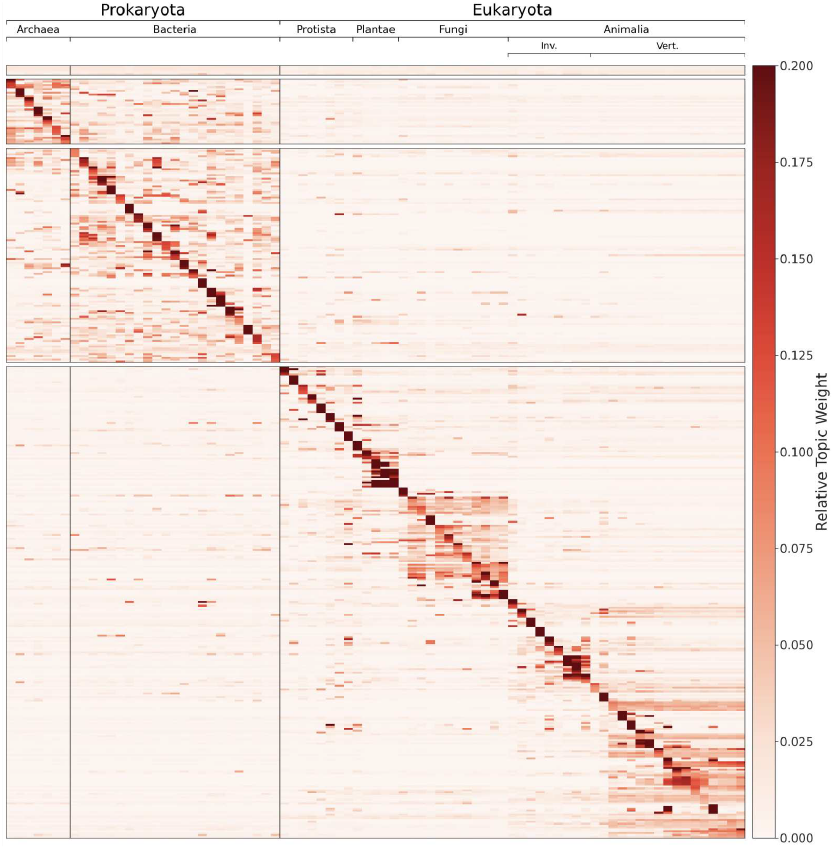
Topic weights for NMF on 81 reference proteomes. Universal topics (top) are selected as the 5 highest-weight topics where all three domains contribute to ≥80% of the topic’s mass; lineage-specific topics (per-kingdom panels) are the 5 topics per proteome with highest dominance and specificity. Columns are organized phylogenetically within each kingdom.

Topics capture shared biological signal across proteomes. Figure S37 shows a heatmap of the top topic weights across the 81 proteomes. The profile of each protein is computed by averaging the topic weight vector ***W*** across all proteins within each proteome, yielding an 81 × 3,000 matrix. For visualization, we selected topics using two criteria: (1) universal topics, defined as the five highest-weight topics where all three domains (Archaea, Bacteria, Eukaryota) are required to account for 80% of the topic’s mass; and (2) lineage-specific topics, defined as the five topics per proteome where that proteome has the highest normalized weight (dominant) and greatest specificity (max − min weight across proteomes). This yielded 395 topics for display. Heatmap values were normalized per topic (columns sum to 1) to show relative distribution across proteomes rather than absolute magnitudes. Proteomes are ordered phylogenetically within each kingdom based on the NCBI taxonomy. A clear block diagonal structure emerges where related organisms share similar sets of feature combinations.

###### NMF topics in the SAE decoder space

Each NMF topic can also be represented as a direction in the SAE decoder space by taking the weighted sum of the respective feature vectors in the decoder space (i.e. ***H* · *W*** _dec_). For each topic, we compute its nearest neighbor in H-space (i.e. for each topic, we find the topic that is most similar in terms of the SAE features that compose the topics) and compute the cosine similarity between these two topics in the SAE decoder space. Figure S38A visualizes this joint distribution among these topic pairs. Looking at the marginal distribution along the x-axis, we can see that the majority of topics do not have close neighbors in the SAE space. This suggests appropriate parameterization of our H matrix. Furthermore, we see from the high correlation (Pearson *ρ* = 0.95) that the H-space broadly reflects the functional organization of the decoder space.

As shown in A.4.3.4, neighborhoods of individual features in the decoder space are generally coherent, with nearby features exhibiting similar semantics. A natural question is whether such coherence is preserved as these features are combined into topics. For each feature, we compute the cosine similarity to its nearest neighbor in the decoder space. We also compute the cosine similarity between this pair of features in the topic-composition space (i.e. the columns of H, which capture how the feature is used across all topics). As shown in Figure S38B, these similarities are largely uncorrelated, suggesting that topic semantics are not determined by individual feature identities alone, but emerge from how features are combined.

This finding motivates further investigation into the role individual features play in the topic semantics. To probe whether decoder-space neighbors of individual topics are substitutable under compositional semantics, we replaced the feature with the greatest weight in a particular topic with its nearest neighbor in the decoder space. We then recompute the topic vector in the decoder space and compute the cosine similarity to the original topic vector. Figure S38C shows that many topics change substantially under this substitution, consistent with the hypothesis that features act as distinct elements in a compositional grammar.

As expected, the magnitude of this change is strongly correlated with the weight of the substituted feature (Figure S38D). This behavior is well explained by the underlying decoder geometry—substituting a dominant feature moves the topic along a nearby direction in decoder space, but does not generally preserve the original topic representation. Taken together with the lack of correspondence between decoder-space and feature-space neighborhoods, this result reinforces that decoder-space similarity reflects alignment in the standalone semantics of individual features, while topic meaning depends on the specific combination of features used to realize that direction.

#### A.4.5. Protein Similarity in the SAE Feature Space

SAE features create an alternative space in which similarity between proteins can be computed. Specific features represent specific functional characteristics, and when pooled over a sequence, the similarity in this space directly compares proteins according to the presence or absence of these functional characteristics. This allows for interpretable searches over protein structure and function, even when the characteristics driving the comparison are present in only a local region or even single residues. SAE similarity can be used to cluster proteins by function or to assign functions to unannotated proteins in a way that is not possible by comparing sequences alone.

**Figure S38.**
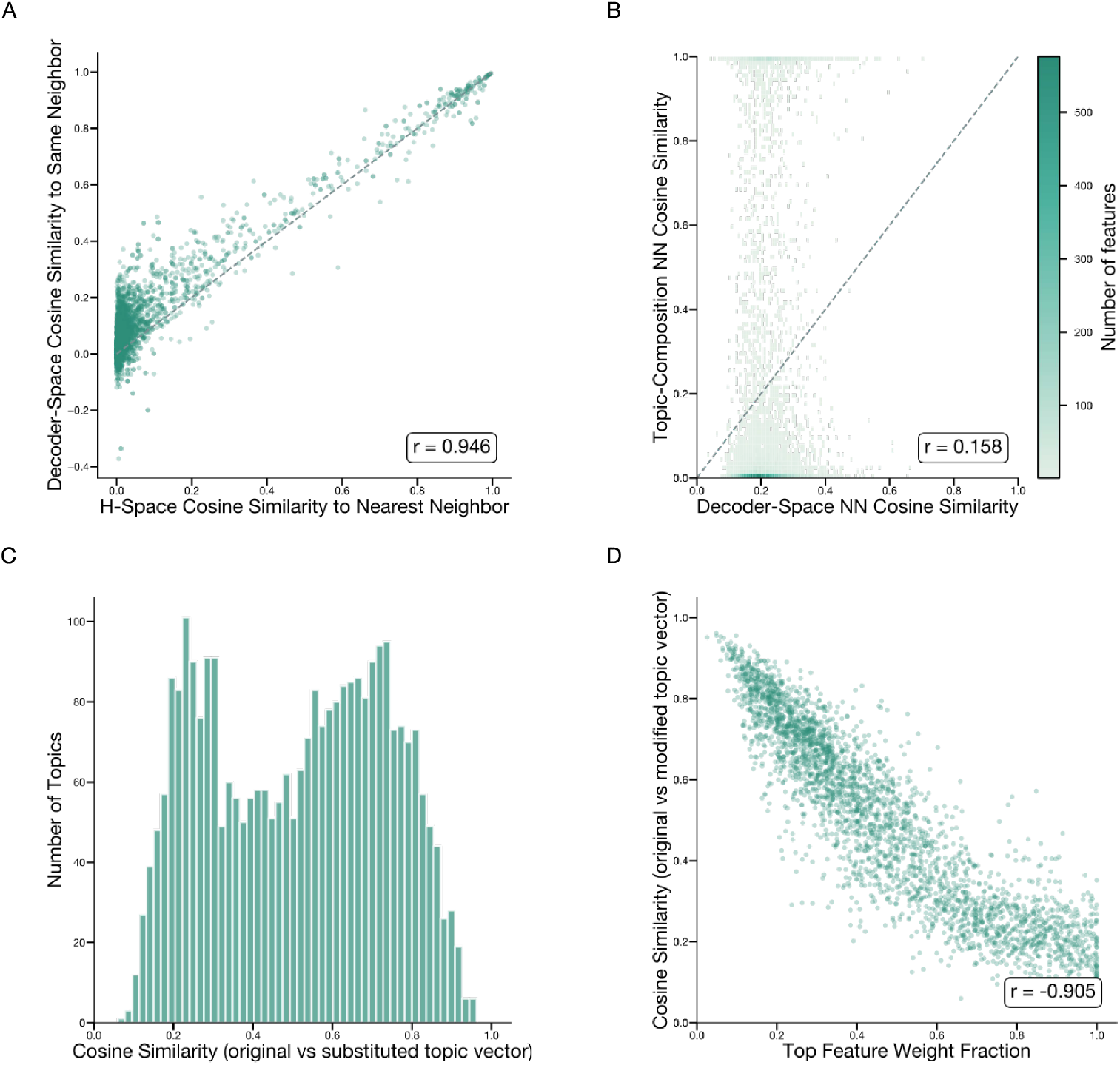
Decoder-space and feature-space representations of NMF topics. (A) Cosine similarity in H-space (rows of ***H***) vs decoder-space (***H*** · ***W*** _dec_) between each topic vector and its nearest neighbor in H-space. (B) For each SAE feature, decoder-space nearest-neighbor cosine similarity (***W*** _dec_) versus topic-composition cosine similarity (columns of ***H***) to the same neighbor. (C) Cosine similarity between original and modified topic vectors after replacing each topic’s highest-weight feature with its decoder-space nearest neighbor. (D) Top-1 feature weight fraction versus cosine similarity between original and substituted topic vectors.

##### A.4.5.1 Protein-Level Representations

Pooling residue embeddings across the sequence dimension is commonly used to generate a single, fixed-dimension embedding for a protein in order to easily compare proteins of different lengths. Mean pooling has been commonly used throughout the protein language model literature, but may not be the best strategy for the sparse, high-dimensional space produced by sparse autoencoders. We investigate this question in the context of a curated dataset from Ding *et al*. (149), which contains sets of homologous proteins across many different organisms. Specifically, we examine a pair of distant ATP8 homologs: human ATP8 and *Candida albicans* ATP8.

When computing the cosine similarity between two protein representations x and y, we compute the relative feature importance of dimension *i* as abs(*x*_*i*_*y*_*i*_)*/* ∑_*j*_ abs(*x*_*j*_*y*_*j*_). Figure S39A shows the cumulative relative importance for the top 100 features for different representations and different pooling strategies. All pooling is done over the full sequence, with BOS and EOS tokens stripped.

The SAE feature with the greatest importance (for both mean pooling and max pooling) is F9211, which represents “core, highly hydrophobic multi-pass alpha-helical transmembrane subunits of bioenergetic inner membranes […]”

The feature is highly active across most of both proteins; the activation for the human ATP8 and *C. albicans* ATP8 proteins are shown in Figure S39B. Because the mean activation is quite high, it dominates the SAE cosine similarity when mean pooling is used: with mean pooling, 70.98% of the relative feature importance comes from this one feature, while with max pooling, its relative feature importance is 8.55%.

An individual feature that spikes on a few residues can still drive a significant amount of cosine similarity when features are aggregated with max pooling. The third most important feature with max pooling is F6331, representing a membrane–cytosol interface motif. With max pooling, the relative importance of this feature is 3.7%. With mean pooling, F6331 is the 7th most important feature and its relative importance is only 0.52%. Figure S39C shows the activation for this feature across the proteins.

Figure S39D shows the distribution, across the entire dataset, of the max-pooled values for F9211 and F6331. The discriminative ability of F6331 is much stronger than that of F9211, but with mean pooling, F9211 dominates the cosine similarity. With max pooling, other features can contribute more meaningfully to the overall signal.

Another consideration when comparing protein representations, regardless of pooling strategy, is the choice of similarity metric. Cosine similarity is sensitive to the magnitude of the activation, which is critical when activations are polysemantic but in a sparse and monosemantic space it becomes less important and can even cause challenges. Figure S39D also shows the discriminative ability of F12752, which has a greater AUPRC than F9211. However, the activation magnitude of F12752 is quite low across both proteins (max of 0.10 in human ATP8 and max of 0.11 in *C. albicans* ATP8). Consequently, F12752 only contributes 0.003% to the mean-pooled cosine similarity score and 0.19% to the max-pooled cosine similarity score. Feature normalization, which increases the relative weight of features with higher information content, is one effective way to address this. Another is to use Jaccard similarity, which considers the *set* of active features and is not dominated by high-magnitude features. Since individual feature magnitudes are not correlated with ability to discriminate function (Figure S39E), Jaccard similarity can be a useful way to compare proteins based on simply the presence or absence of specific features.

Finally, to illustrate the benefits of using a high-dimensional SAE space relative to the original dense representations, Figure S39F shows the contribution of each latent dimension to F6331 as defined by the corresponding weights in the encoder. This distribution confirms that the information encoded in this feature is not recoverable from any individual dimension in the latent space.

##### A.4.5.2 Pfam Family Analysis

###### Dataset

The Pfam protein families database (150) is a comprehensive collection of protein families and domains which is widely used for protein annotation. Each domain in Pfam is defined by a hidden Markov model (HMM) trained on a set of representative sequences known as a “seed alignment”. By construction, Pfam seed alignments are made up of proteins with similar sequences and therefore similar functions. For this reason, we would expect proteins from similar Pfam domains to contain similar sets of SAE features.

To confirm this, we downloaded the Pfam-A seed alignments (Release 38.0) and extracted UniProt accessions and residue ranges for each domain in the alignments. We then filtered to proteins in UniRef90, for which we had computed SAE features at each position in the full-length protein. The resulting dataset contained features for 1, 404, 531 unique protein domains representing 24, 343 Pfam families. The feature representation for each domain is computed using the set of features only within the domain boundaries specified in the alignment.

**Figure S39.**
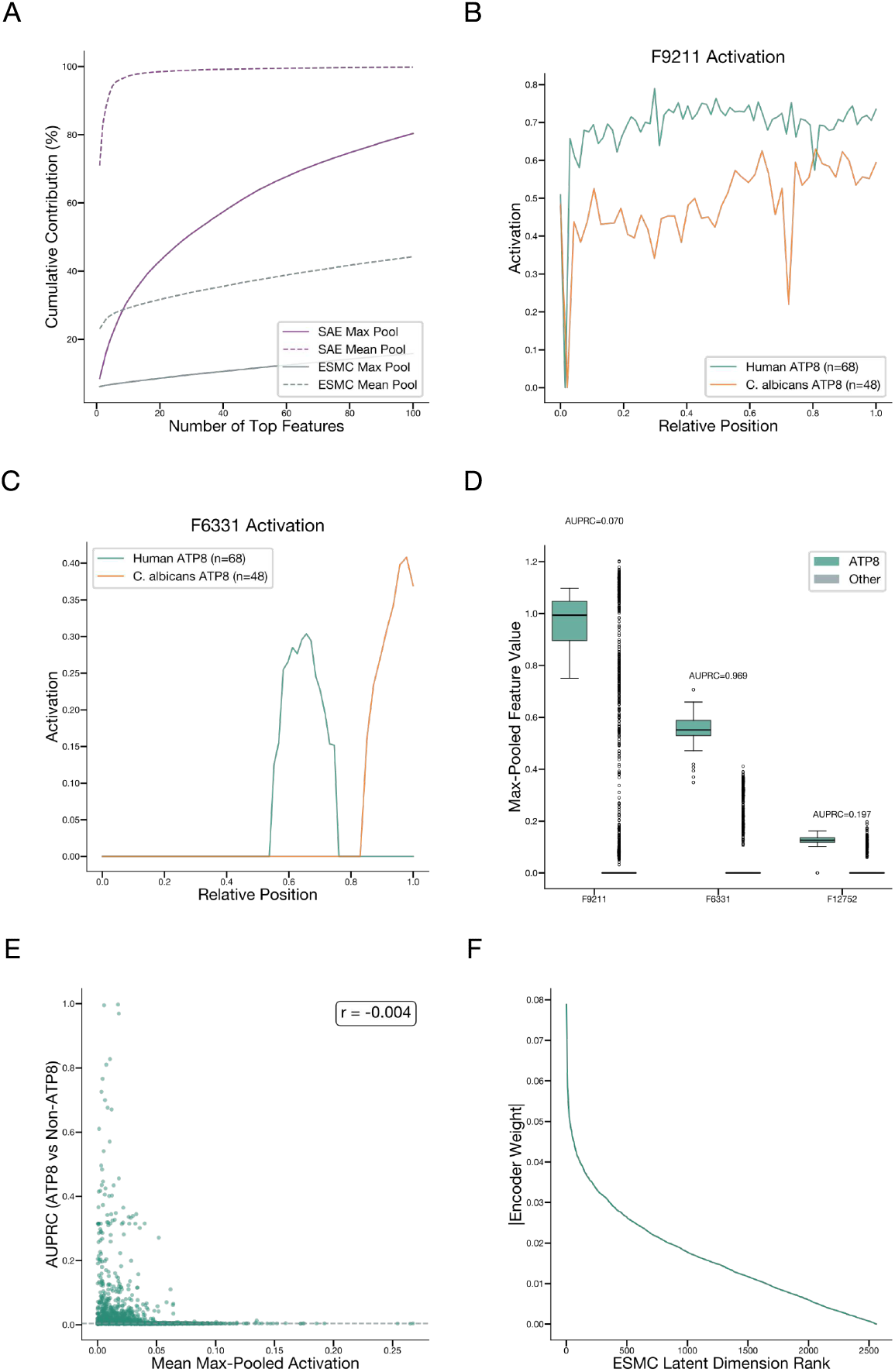
Max pooling preserves discriminative SAE features lost by mean pooling. **(A)** Cumulative relative feature importance for cosine similarity between human ATP8 and *C. albicans* ATP8, comparing SAE and ESMC representations under max and mean pooling. **(B)** Per-residue activation of the top-ranked feature F9211 across human ATP8 and *C. albicans* ATP8. **(C)** Per-residue activation of F6331 across human ATP8 and *C. albicans* ATP8. **(D)** Distribution of maximum value of F9211, F6331, and F12752 for ATP8 proteins and all other proteins. **(E)** Mean of max-pooled activations across all proteins compared to AUPRC for ATP8 classification. **(F)** Sorted absolute encoder weights mapping ESMC latent dimensions to F6331.

###### SAE feature similarity

To measure the degree to which domains from the same Pfam family share SAE feature representations, we compute the pairwise Jaccard similarity between the active (activation *>* 0) features in all pairs of matching domains, considering only the 14,841 Pfam families which have at least ten examples. In Figure S40A, we plot the median and minimum pairwise similarity, showing that there is high concordance between the active feature sets of domains from the same family—over 88% of families have a median pairwise Jaccard of greater than 0.6, and over 50% of families exceed this threshold for every pair of domains. We compared these results against a background distribution of 100,000 random pairs of domains belonging to different Pfam families. In this background distribution, the median Jaccard similarity was only 0.177, with not a single randomly sampled pair exceeding 0.6 similarity.

###### Family characterization

As protein sequence databases have expanded, many more sequence clusters have been discovered and proposed as potential new Pfam families. These include both families with and without characterized function, and domains with unknown function represent a significant fraction of the Pfam database. To classify a family as uncharacterized, we used string matching on the Pfam family description with the following conditions:

The strings `uncharacter`, `unknown function`, `hypothetical`, `predicted`, or `putative` anywhere in the description; any of the following strings followed directly by a number: `DUF[number]`, `UPF[number]`, `FAM[number]`; open reading frame notation: `C[number]orf[number]`

This captures slightly more families than the standard set of DUFs (domains of unknown function) and UPFs (unknown protein functions), but it is still not complete—many families with unknown function are given names which do not map to one of these categories, and some DUFs have been characterized since their addition to the Pfam database but the name or description has not been updated. Nonetheless, this criteria offers a reasonable estimate of the proportion of families which remain functionally unknown. Figure S40B shows that uncharacterized families have a similar distribution of seed alignment size to characterized families, but that the majority of large families (seed alignment size *>* 100) are characterized. We estimate that ∼25% of families are functionally uncharacterized, representing 18.2% of all seed alignment proteins. This characterization scheme is used to estimate the characterization level of clusters in the ESM Atlas (A.5.6).

##### A.4.5.3 Structure and Function Retrieval at Low Sequence Similarity

To quantitatively assess how well max-pooled SAE features can identify structural and functional similarities between proteins of low sequence identity, we conducted low-homology retrieval experiments on two datasets. The CATH S95 (95% nonredundant) dataset assesses structural retrieval at the fold (T) level (*n* = 65,534 CATH domains), and EC numbers from our 90% nonredundant SwissProt dataset assess functional retrieval at EC level 4 (*n* = 91,470 proteins with a single full EC number; length ≤1024 residues; and a matching AlphaFold-DB structure). On each dataset we compare five methods: three use the ESMC 6B layer-60 representations (SAE cosine, SAE Jaccard, and mean-pooled ESMC cosine), one is structure-based (Foldseek (93), paper settings: bit-score with −*s*, 9.5, − *e*, 10, `--max-seqs` 2000), and one is sequence-based (MM-seqs2 (136), all-vs-all and ranked by bit-score; −*e*, 1.0, default sensitivity 7.5, `--max-seqs` 2000 for CATH and `--max-seqs`5000 for EC). In both datasets, SAE-based similarity outperforms all other methods at detecting proteins with shared structure and function when the sequence similarity is below 40%.

We evaluate each method’s retrieval performance by measuring how well it ranks related proteins above unrelated ones. For each method, an all-vs-all pairwise similarity is computed across the dataset, giving each query a similarity-ranked list of every other protein. The protein at rank *r* in *q*’s ranked list (denoted *p*_*r*_) is a true positive (TP) if it shares the query protein *q*’s fold (CATH T tier) or full EC number, otherwise a false positive (FP). For each TP pair (*q, p*_*r*_) we compute the fraction of proteins ranked at or above *p*_*r*_ that are themselves true positives:

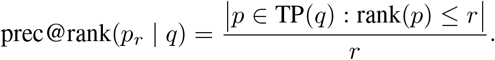

We stratify pairs by global sequence identity between *q* and *p*_*r*_ into sequence-identity bins. Within each bin we visualize the distribution of per-pair prec@rank(TP) values — intuitively, how well a query’s homologs at this identity level rank above false positives. Each pair is weighted so that every query contributes equally within its class and every class contributes equally, preventing heavily populated CATH folds or EC groups from dominating. Dots mark the resulting class- and query-weighted mean per method.

Figure S41 shows the result of this search experiment on each dataset. All methods perform very well above a sequence identity of 50%, where there is significant sequence homology. Below this, there is separation between the average performance, with SAE-based methods performing best on both tasks. On the CATH dataset, Foldseek is more competitive as a structure based method, but still falls short at very low sequence similarities. At low sequence identity, retrieval is notably bimodal for a (query, true positive) pair: either the true positive is near the top (hit) or buried among false positives. This means that for measuring performance on an individual query, the average over all classes is not as important as how many TP a query retrieves at the top of the list. For proteins below 30% identity, we therefore also compute the fraction of true-homolog pairs ranked at near-perfect positions (precision@rank(TP) ≥ 0.9). For an average CATH query with SAE cosine similarity, 70% of true homologs at *<*30% sequence identity are ranked correctly — 5–6 percentage points above ESMC cosine similarity (64%) and Foldseek (65%) (Figure S41B top). The bimodality also holds on the per-query level and the class-level (Figure S41B bottom). For 51% of CATH topologies, the average query with SAE cosine similarity retrieves nearly all true homologs correctly (avg hit rate ¿ 0.9). This is higher than SAE with Jaccard similarity (43%), ESMC cosine similarity (43%), Foldseek (39%), and MM-seqs (3%). Taken together, these experiments demonstrate that the ESMC latent space, particularly when projected into the sparse feature space with SAEs, identifies structural and functional connections between distant proteins that would otherwise be very difficult to discover with standard methods for large-scale search in sequence or structure space.

**Figure S40.**
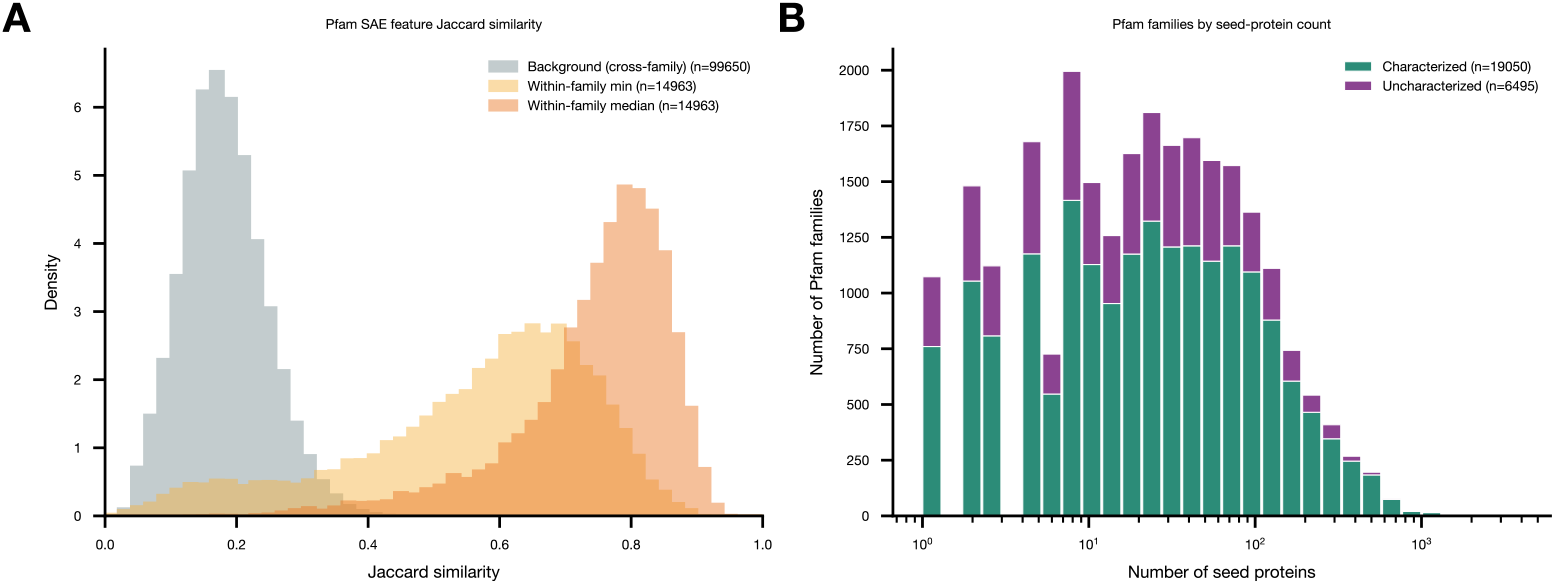
Pfam domain similarity in SAE space. (A) Jaccard similarity between domains within the same Pfam family and randomly sampled pairs of domains from different families. Both median and minimum pairwise similarity are shown for within-family pairs. (B) Number of protein domains in the seed alignment for Pfam families with characterized and uncharacterized function. The inset shows the relative proportion of characterized vs. uncharacterized families in the Pfam-A seed dataset, and the corresponding proportions of proteins.

##### A.4.5.4 Taxonomic Bias Evaluation

SAE features lend themselves to sensitive functional search. One might speculate that the properties of the sparse SAE feature space might also help to mitigate a common failure mode of functional search using protein language model embeddings—taxonomic bias. ESMC’s representation space learns to encode everything that is relevant about a sequence for predicting masked amino acids, including features that are correlated with the organism the sequence is from. It has been shown that phylogeny is a confounder for functional search with language model embeddings, as proteins from closely related species may share, for example, amino acid composition biases that obscure their true functional relationships (151). If these features are present across the entire sequence, then local, functionally specific features, may be drowned out by more broadly active taxonomic signal. Under this hypothesis, max pooling in SAE space may be less biased by taxonomy than mean pooling in the dense representation space (A.4.5.1), particularly for capturing functional signal across large evolutionary distances.

To validate this hypothesis, we first investigated whether SAE features have higher sensitivity for identifying functional homologs compared to sequences that are spuriously “similar” because they are derived from similar taxa. To evaluate this, we use the “Proteinbias” dataset curated by Ding and Steinhardt(149) to highlight the effects of uneven sampling across the tree of life on protein language models. This dataset consists of approximately 8,000 sequences curated to contain functional homologs across multiple species, with each protein associated with 15 to 100 homologs from other species. This enriches the dataset for broadly conserved functions, such as ribosomal subunits and ATP metabolism. Each protein is annotated with a functional label derived from its UniProt protein name.

**Figure S41.**
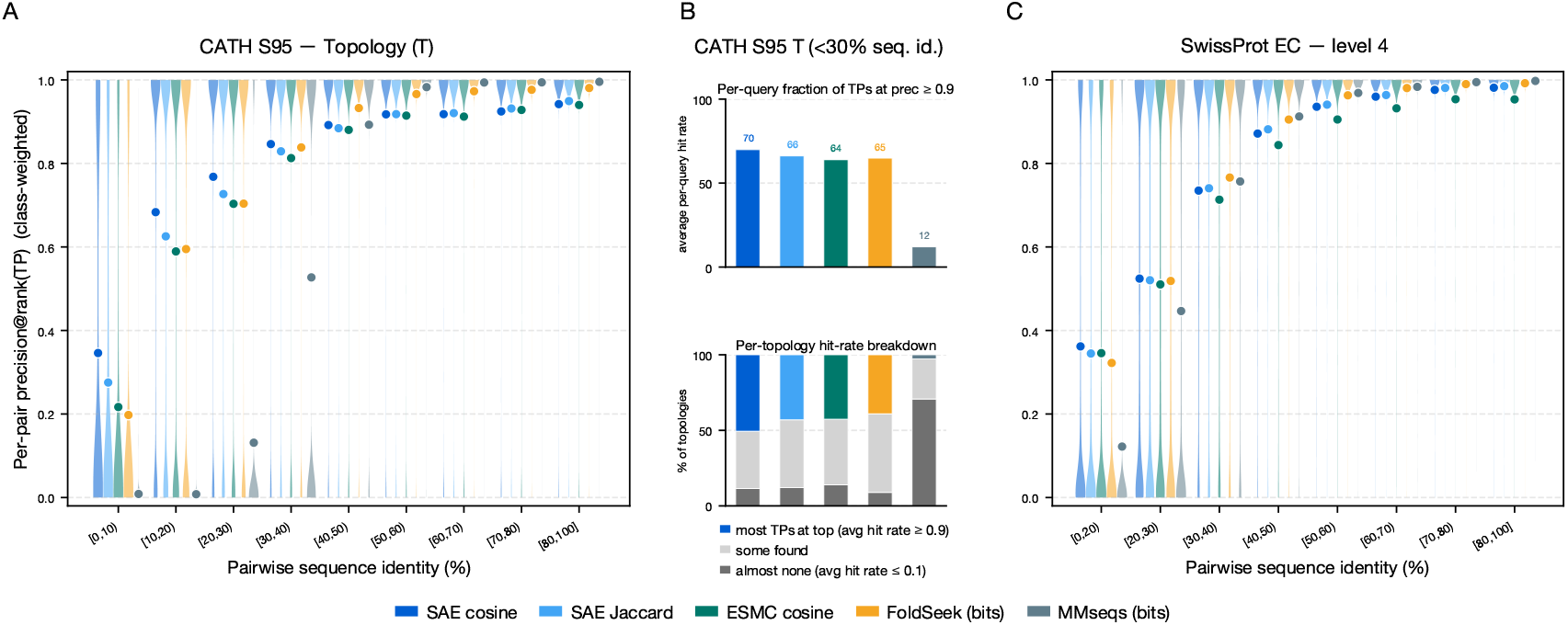
Remote structural and functional retrieval in SAE space. (A) Class- and query-weighted per-pair precision@rank(TP) distributions across pairwise sequence-identity bins (computed with global Needleman–Wunsch alignment) for CATH topology retrieval. Dots mark the class+query-weighted mean per method, and violin plots show the full distribution over pairs. (B) Direct comparison for CATH topology queries at *<*30% sequence identity: (top) per-method average per-query hit rate; (bottom) fraction of topologies broken down by average hit rate across classes. Hit rate(q) = fraction of q’s TPs with precision@rank(TP) ≥ 0.9, i.e., how many of a query’s TPs are ranked at the top of the retrieval list. (C) Class- and query-weighted precision@rank(TP) for SwissProt EC level 4 retrieval, analogous to (A).

For benchmarking functional homology search on this dataset, each protein in the dataset is embedded with ESMC 6B. In order to establish parity between the ESMC cosine search and SAE search, the 60th layer from ESMC was selected for comparison between the SAE (*K*=64, codebook size 16, 384) and the dense representations. As in prior analyses, max pooling is used for SAEs and mean pooling for ESMC embeddings. For each query protein, all 8,000 proteins in ProteinBias are ranked using three similarity metrics: (1) Jaccard similarity on the SAE max-pooled features, converted to a binary feature vector indicating whether each feature activation *>* 0; (2) cosine similarity on the ESMC mean-pooled embeddings; and (3) MMSeqs similarity, using `mmseqs search -e 1.0 --max-seqs 2000`. The recall at 1 FP is then computed for each query. This reflects a common annotation-transfer setting, in which a function is assigned to a new protein by searching against a database of proteins with known functions.

Figure S42 shows that a higher proportion of queries achieve 80% recall at 1 FP using Jaccard similarity of SAE features than for ESMC representations, with both outperforming sequence similarity search on this dataset. In particular, when restricting to queries for mammalian proteins, feature similarity substantially outperforms ESMC representation similarity, increasing the proportion of queries achieving 90% recall from 78% to 92%. This improvement is driven in large part by reduction in false positive matches to proteins from the same taxonomic class, as indicated by the close correspondence between dotted and solid curves in Figure S42.

**Figure S42.**
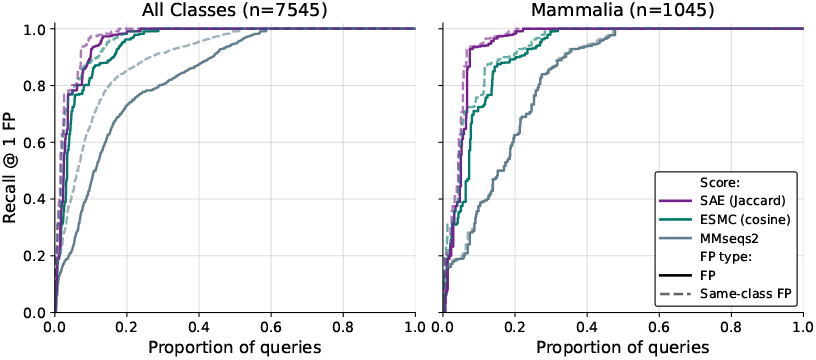
Performance on retrieving proteins with the same function across different species. Recall up to the first false positive (FP) is plotted for both any FP (different function or different taxonomic class; solid lines) and specifically for FPs due to shared taxonomic class (dotted lines).

##### A.4.5.5 Biological Pathway Assignment

Proteins can be organized into pathways based on the specific biological processes they participate in. For example, some mitochondrial proteins may participate in the oxidative phosphorylation (OXPHOS) pathway, which is responsible for the production of ATP. Assigning a protein to a particular pathway can illuminate a protein’s role in cellular mechanisms. We use pathway assignment as a test case for evaluating how well similarity in the SAE feature space can associate proteins with shared functional roles.

Given a reference database of proteins where each is associated with a set of one or more biological pathways, pathway assignments for query proteins with unknown function are predicted based on similarity in SAE space. Each protein in the reference database is embedded into the SAE feature space, as well as into the ESMC representation space as a baseline. Residue-level embeddings are pooled over the sequence dimension to produce a single embedding vector for each protein (SAE features are max pooled, ESMC latents are mean pooled).

The same procedure is applied to query proteins and Jaccard similarity is computed to each protein in the reference database. We evaluate pathway prediction performance using the top-*k* reference proteins by Jaccard similarity. For held-out proteins with known pathway assignments, we take the union of all pathways assigned to the top-*k* reference proteins as the set of predicted pathways and all known pathways for the query protein as the set of true pathways. Using these two pathway sets, we compute a precision and recall score for each (protein, *k*) pair. The final scores reported here are the maximum F1 score achieved over *k* ∈ {1 … 5}. For query proteins without known pathway assignments, all pathways from the top five nearest reference proteins are considered as candidate pathway assignments.

This procedure is agnostic to the specific pathway database used, as long as the database defines a mapping from protein sequences to a set of labels. We focus primarily on the MitoCarta dataset (152), an inventory of 1136 human genes encoding proteins with strong support for mitochondrial localization, including annotations for sub-mitochondrial compartment and biological pathway, which has been curated as part of an effort to create a functional map of human mitochondrial proteins.

###### MitoCarta reference dataset

Data for all 1136 human genes with high confidence of mitochondrial localization in MitoCarta 3.0 were downloaded from the Broad Institute website. Uniprot IDs were used to retrieve sequences for each human protein. After removing proteins with sequences over the 2048 context length, this produced a reference dataset with 1126 unique proteins mapped to 137 unique mitochondrial pathways (also termed MitoPathways). The number of pathways per sequence ranges from 1–7 with a median of 1 and a mean of 1.58.

For the retrospective pathway assignment analysis, we held out proteins that were not present in the previous version of MitoCarta (v2.0) but were added in MitoCarta 3.0. For this evaluation, we created a new reference database containing only MitoCarta 2.0 proteins (*n* = 1051 proteins). We do not include proteins which were removed from the entire database in MitoCarta 3.0 because they did not have sufficient evidence of mitochondrial localization. The remaining 60 proteins that were contained only in MitoCarta 3.0 and had at least one MitoPathway annotation were used as our held-out evaluation dataset.

###### MMSeqs2 baseline

Sequence search was performed using MMSeqs2 easy search against the MitoCarta 2.0 reference sequences:

~~~
     mmseqs search -e 1.0 --max-seqs 2000
~~~

###### Foldseek baseline

Structural search was performed using Foldseek using predicted structures for both reference and query proteins from ESMFold2. MSAs were generated using the ColabFold pipeline using parameters:

~~~
     colabfold search -s --use-env
     1 --filter 1 --expand-eval inf
     --align-eval 10 --diff 3000 --qsc
     -20 --max-accept 1000000
~~~

After structures were predicted, a Foldseek search was performed:

~~~
     foldseek easy-search -s 9.5
     --alignment-type 1
~~~

###### Retrospective evaluation results

We predict the biological pathway annotations for these 60 mitochondrial proteins using a similarity search against proteins in MitoCarta 2.0 based on shared features. Each query protein is compared to proteins in MitoCarta 2.0 using Jaccard similarity between the binarized feature vectors, and pathway annotations are assigned by aggregating the labels of the k nearest neighbors. As in the original dataset, a protein may have multiple pathways assigned to it. We perform the same nearest-neighbor search procedure using similarity measures based on the ESMC representation (embedding cosine similarity), sequence similarity computed with MMSeqs2 (bit-score), and structural similarity computed with Foldseek. Both the SAE features and ESMC representations recovered more correct pathway annotations than sequence and structure-based searches alone, with SAE features performing best at a max F1 score of 0.391 (Figure S43).

**Figure S43.**
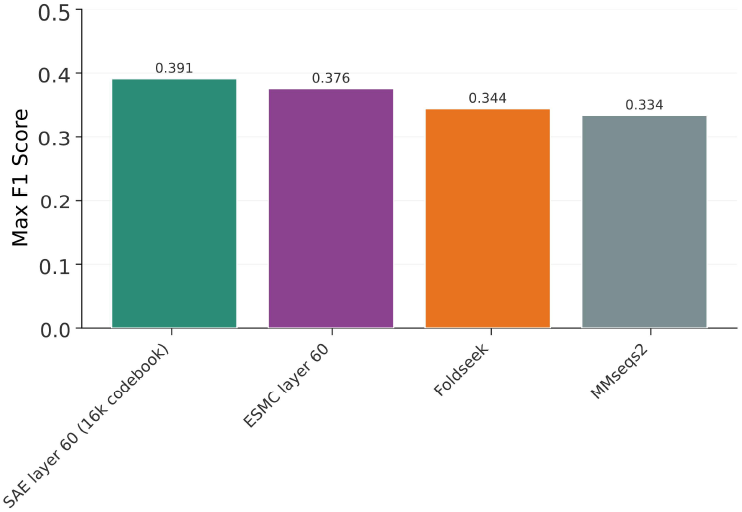
MitoCarta 3.0 pathway assignment results. (B) Mitochondrial pathway assignment results for retrospective analysis of 60 proteins in Mitocarta 3.0. Results shown are the average over the best F1 score achieved by aggregating the top k search results, for *k* ∈ 1, 5.

###### Literature validation case study

As of the publication of Human MitoCarta 3.0 in 2021, there were 99 proteins which did not have any MitoPathways assigned. Some of these have been experimentally characterized since then, including the inner mitochondrial membrane protein transmembrane protein 65 (TMEM65).

TMEM65 is highly expressed in the heart and brain, and while it was known to localize to the inner mitochondrial membrane, its function was until recently unknown. Since 2025, several papers have revealed new roles for TMEM65 as a Na^+^/Ca^2+^ exchanger which is essential for mediating calcium balance and cellular homeostasis in the mitochondria (153–155). This provides a natural retrospective evaluation for how SAE features can be used for functional annotation transfer.

Measuring similarity against the MitoCarta reference dataset, the top neighbors of TMEM65 group into multi-pass inner membrane machinery (e.g. OXPHOS assembly factors and TIM translocases), driven heavily by F9059 (multimeric membrane-pore ring architecture). Since the MitoCarta database limits this approach to annotations from mitochondrial proteins only, we hypothesized that it may be useful to expand the search to a broader set of pathways. Any pathway database can be used in this prediction pipeline, including the Kyoto Encyclopedia of Genes and Genomes (KEGG) (156), which maintains a set of curated canonical pathway maps for the entire proteomes of a wide range of organisms. We construct a reference database for all human pathways recorded in KEGG, considering only those focused on biochemical function (removing “Human Diseases” and “Drug Development” categories, which capture correlations with broader phenotypes), resulting in a dataset consisting of 8,141 human proteins assigned to 262 unique pathways.

Similarity search against the KEGG dataset provides a more precise mechanistic hypothesis for TMEM65, one which matches closely with the experimental evidence. The nearest neighbors to TMEM65 in the KEGG-annotated human proteome are SLC-family sodium-dependent transporters and rotary-pump ring subunits, centering the predicted pathways on ion homeostasis. This was driven by features for multi-pass transporter bundles (F131) and divalent metal cation binding (F9143). These results demonstrate how similarity in SAE space can be useful for identifying similarities between an uncharacterized protein and selected reference datasets containing known functional annotations, enabling the generation of explainable hypotheses for protein function from sequence alone.

### A.5. ESM Atlas

#### A.5.1. Source datasets

Protein sequences were assembled from eight public source repositories, summarized in Table S14: UniProt’s UniParc archive (an aggregation of EMBL-Bank/GenBank/DDBJ, RefSeq, Ensembl, PDB, and other public repositories) (75); IMG/VR (79), IMG/M NR, and the prokaryotic and eukaryotic MAG-binned UMAG subsets from the JGI Integrated Microbial Genomes and Microbiomes resource (78); the MGnify protein catalog (30) and the UHGG human gut microbiome catalog (77) from EBI MGnify; and the SPIRE metagenomic gene catalog from EMBL (76). FASTA headers from each source were standardized to encode the MD5 hash of the amino-acid sequence, a data-source label, and a unique source-side identifier. Across the eight sources the input contained 7,824,094,326 sequences before deduplication.

The full ESM Atlas dataset (6,824,676,938 sequences) was produced by exact deduplication on sequence MD5 across all eight sources with no length filter. All of these sequences are embedded using ESMC 6B and SAE features are computed using the layer 60, *k* = 64, 16, 384-dimensional sparse autoencoder.

#### A.5.2. Sequence clustering and structure prediction

Sequence-based clustering was performed to derive a representative non-redundant database for computing predicted structures in the ESM Atlas. This subset comprises sequences of length 60–1000 amino acids clustered at 70% sequence identity. Sequence clustering was performed with MMseqs2 v18.8cc5c (136, 157). Seven of the eight sources (IMG/M, IMG/VR, SPIRE, UHGG, UMAG-prokaryotic, UMAG-eukaryotic, and UniParc) were independently clustered at 90% sequence identity using Linclust with `mmseqs linclust --min-seq-id 0.9 -c 0.8 --cov-mode 1`. Coverage mode 1 was used throughout all clustering operations, requiring the shorter sequence to align across at least 80% of its length to the cluster representative. MGnify proteins were not clustered at this step because the upstream MGnify release is already clustered at 90% identity (30).

**Table S14.** Sequence database sources in the ESM Atlas.

| Database | Dataset (release) | Source of sequences | Reference |
| --- | --- | --- | --- |
| UniProt | UniParc | Aggregation of public protein and nucleic-acid databases | UniProt Consortium 2025 (75) |
| JGI IMG | IMG/M NR (snapshot Sept 16, 2024) | Non-redundant proteins from microbial isolate genomes | Chen *et al.* 2023 (78) |
| JGI IMG | IMG/VR (release 7, Dec 19, 2022) | Cultivated and uncultivated viral genomes | Camargo *et al.* 2023 (79) |
| JGI IMG | UMAG-prokaryotic (file dated 2025-04-30) | MAG-binned prokaryotic sequences | Chen *et al.* 2023 (78) |
| JGI IMG | UMAG-eukaryotic (file dated 2024-02-25) | MAG-binned eukaryotic sequences | Chen *et al.* 2023 (78) |
| EBI MGnify | MGnify proteins (release 2024_04; pre-clustered at 90% identity upstream) | Metagenomic proteins from multiple environments | Richardson *et al.* 2023 (30) |
| EBI MGnify | UHGG v2.0.2 | Human gut microbiome protein catalog | Almeida *et al.* 2021(77) |
| EMBL SPIRE | SPIRE genes (SPIRE v1) | Metagenomic sequences from multiple environments | Schmidt *et al.* 2024 (76) |

The per-dataset 90% representative sets, together with the MGnify 90% representatives, were then merged into a single MMseqs2 database after restricting sequences to lengths between 60 and 1000 amino acids. The database was clustered at 70% sequence identity using Linclust with `--min-seq-id 0.7 -c 0.8 --cov-mode 1`. The resulting representative set of 1,095,527,764 sequences defines the atlas-clustered collection used to compute predicted structures. Structures were predicted with `esmfold2-exp-2026-03`, 3 loops, 22 diffusion steps, `step scale=2.0`, `noise scale=0.3`.

#### A.5.3. Pfam annotations

Protein domain annotations were generated by searching all sequences in the ESM Atlas against the Pfam-A profile HMM library (release 38.1, derived from UniProtKB reference proteomes 2025 03; 27,481 families) (81) using pyhmmer.hmmsearch (pyhmmer v0.12.0 (138), built against HMMER ≥ 3.4 (137)) with Pfam gathering thresholds (`bit_cutoffs=`“`gathering`”). For each protein the resulting per-domain hits were retained as the list of matching Pfam family accessions and descriptions and the per-domain envelope residue ranges, together with a domain-coverage value defined as the fraction of amino-acid positions covered by the union of all domain envelope coordinates.

For each SAE-based cluster (A.5.4) Pfam annotation statistics were derived by joining per-protein annotations to SAE cluster membership through the MD5 sequence identifier the member proteins. For each SAE cluster, the fraction of SAE cluster members carrying at least one characterized Pfam domain, the mean per-member domain coverage, and per-family member counts were reported. Pfam accessions were version-stripped for cluster-level counting, and multiple occurrences of the same Pfam family within one protein were counted as a single observation for that protein. Characterization status of each family was computed as described in Section A.4.5.2 and used to compute the level of characterization in each cluster.

#### A.5.4. SAE clustering

Section A.4.5 shows that Jaccard similarity over binarized, max-pooled SAE feature sets (essentially treating each protein as a “bag-of-features”) is effective for recapitulating known biological relationships, and that similarities in this representation space can surface connections that sequence similarity alone cannot. This motivates organizing the atlas according to the model’s latent space directly. This would provide a complementary view on protein space to existing databases and provide more interpretable insights into the relationships between proteins through the model’s latent representations. Ultimately, this could reveal biological insights that could not be gained from analyzing only existing protein cluster databases, classification systems, or ontologies.

We aimed to cluster all 6.8B proteins in the Atlas such that the Jaccard similarity of all proteins in a cluster is at least 0.6 to its cluster center, a threshold which has high precision for clustering proteins from the same Pfam domain together (A.4.5.2). Computing an all-vs-all Jaccard similarity over the entire 16k-dimensional feature space is infeasible, so we developed an efficient linear-time algorithm for approximate Jaccard-based clustering using a hash-based approach. Hash-based algorithms such as locality-sensitive hashing (LSH) provide a mechanism for approximate similarity search and clustering across very large datasets in linear time (158). The MinHash LSH family in particular is effective for set-based comparison of massive databases (159), and has been applied for linear-time sequence clustering of large protein and genomic databases (157, 160, 161).

In the MinHash LSH approach, *d* independent hash functions are applied to the elements of each set, and the minimum hash value produced by each is collected into a *d*-dimensional signature, or *sketch*. Comparing the number of hashes shared between the signatures of two samples (which represent uniform random permutations of the full input sets) produces an unbiased estimate of the true Jaccard similarity 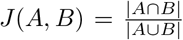, with uncertainty that scales with the number of hash functions *d* (162).

For estimating sequence similarity, methods such as Linclust (157) convert proteins to sets using *k*-mers (for sufficiently sized *k*, proteins with matching *k*-mers are highly likely to be homologous). SAE features convert sequences into sets in a fundamentally different way—two proteins that share SAE features may be related by sequence, structure, or functional properties. We use the sets created by binarized, pooled feature activations over each sequence to implement a distributed analog to Linclust that clusters proteins based on their SAE feature representations.

##### A.5.4.1 Clustering algorithm

In principle, linear-time hash-based clustering methods work by assigning all proteins to a set of “buckets” based on their hash signatures, grouping by bucket, selecting a single representative per bucket, and then comparing all members of the bucket to the representative to identify pairs that are above the specified clustering threshold. The selection of a single center within each bucket is what enables the algorithm to scale linearly (avoiding all-vs-all comparisons within a bucket), but it also reduces recall—sequences that have high similarity to each other but not to the chosen representative will not be clustered together. In general for protein clustering, such false negatives are more acceptable than false positives, as they simply produce more redundancy in the clustering without affecting the integrity of the clusters themselves.

###### Stage 1: MinHash sketch computation

We compute sketches based on the set of unique active SAE feature indices across the entire sequence. One challenge that SAE features present for identifying meaningful functional relationships using fixed-length sketches is their universality; many features are found across a very large fraction of protein space (A.4.2.3). This property, which makes features useful for generalization across families, also means that the false positive rate for grouping proteins based on a random subset of features is very high (the median Jaccard similarity between full feature sets across randomly sampled protein pairs is 0.25). This produces buckets that are very large and heterogeneous, increasing verification costs and reducing sensitivity of the final clustering. Two mitigations reduce this false positive rate: IDF-based feature pruning and LSH banding.

To reduce the effect of non-discriminative features, we compute hashes only on features with inverse protein frequency (A.4.2.4) above a threshold of 0.87. This filters out 3,635 features (22%) and reduces the median Jaccard on the background distribution to 0.02. Using the remaining high-information features, we compute a *m*-dimensional MinHash sketch in which the *k*^th^ hash is computed by

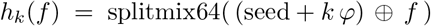

where *φ* = `0x9E3779B97F4A7C15` is the 64-bit fractional part of the golden ratio. The *k*^th^ signature entry is the feature minimizing *h*_*k*_ over the protein’s feature set.

To further improve the specificity of the hash buckets, we use an LSH banding strategy in which we split the full *m*-dimensional sketch into *b* bands of width *r*. In order for two proteins to be considered candidates for grouping, they must agree on *all r* hashes in a band, meaning effectively that proteins are grouped by combinations of *r* features rather than individual features. Each band hash *b*_*i*_ is the iterated `splitmix64` fold of the *r* hashes within the band.

In addition to the band hashes, we extract a fixed-size sketch of size 256 evenly-spaced signature entries which we use for downstream Jaccard estimation in the verification step. The standard deviation of the sketch similarity at *J* = 0.6 is 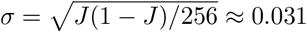.

###### Stage 2: Candidate pair generation and verification

Pairs of proteins that collide in at least one band hash (i.e. share a `(band idx, band hash)` bucket) are considered candidates. We materialize the full `(protein, band index, band hash)` table, partition by band index, and process the table in batches. Within each batch we perform the following steps:

*Bucketing*. For every bucket we deterministically select a bucket representative (minimum protein index). Buckets larger than 10^6^ are dropped to prevent quadratic explosion in “heavy-hitter” hashes. All (rep, member) edges from each bucket are generated and deduplicated (generating a maximum of *k* − 1 edges rather than 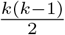).

###### Verification

Each candidate pair is verified using the checkpointed `(protein, sketch256)` table, counting element-wise sketch agreements to estimate Jaccard similarity. A pair is retained if the number of matches exceeds the target Jaccard threshold times the sketch size, i.e. ⌈0.6 · 256⌉ = 154. Verified edges from each batch are written to disk, and after all batches the union is deduplicated to form the final verified edge set.

###### Stage 3: Distributed greedy clustering

We partition proteins into clusters from the (symmetrized) verified edge set using a distributed adaptation the Linclust’s greedy set cover algorithm. In each round, every node that is a strict local minimum among its still-active neighborhood becomes a cluster center, claims its unclustered neighbors, and is removed from the active set together with those neighbors.

Concretely, every protein *u* is assigned a fixed priority π(*u*) = xxhash64(md5(*u*)) (ties broken by `protein_idx`). In round *t* with active node set *V*_*t*_ and active edge set *E*_*t*_, do the following until |*V*_*t*_ | = 0:

1. For every *u* ∈ *V*_*t*_, compute the minimum-priority active neighbor (π_min_(*u*), *v*_min_(*u*)).
2. A node is a center iff it has no active neighbors, or its own priority is strictly less than π_min_(*u*), or it equals π_min_(*u*) and ties on `protein_index`.
3. Each non-center node that has at least one center neighbor is assigned to the lowest-priority center it has an edge to.
4. Assignments are written as `(protein_index, rep protein_index)` and removed from both *V*_*t*_ and *E*_*t*_, with unassigned nodes remaining active for the next round.

##### A.5.4.2 Parameter selection

The choice of band width *r* and number of bands *b* determine the tradeoff between sensitivity and computational cost of the algorithm. For a target Jaccard threshold τ and a sensitivity *S* (probability that a true pair produces at least one verified edge within a bucket), we can model the estimated work done in Stage 2 as a function of *r* and *b*. Let *J*_0_ denote the empirical background distribution of pairwise Jaccard similarities across random proteins, and let *N* be the target database size (6.8 × 10^9^ for the full atlas). Because a single representative is selected per bucket, a pair at similarity *J* becomes a candidate in a given band only when the two proteins share the same hash at that band (probability *J*^*r*^) *and* one of them is the bucket representative, which inside a bucket of size *k*(*r*) (bucket size is dependent on *r*) occurs with probability 2*/k*(*r*):

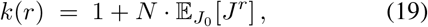

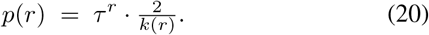

Across *b* independent bands, the achieved sensitivity is 1 − (1 − *p*(*r*))^*b*^, so the minimum number of bands attaining *S* is

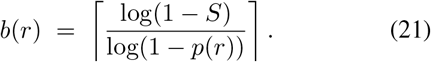

Once candidates are generated, the cost scales linearly in the total number of candidate edges, which is ∼ *N* · *b* regardless of *r* (each band contributes one edge per non-representative protein), so *b*(*r*) alone can be used as a proxy for
 the total cost. Therefore, we can select the value of *r* that minimizes this cost and then compute *b*(*r*) directly.

Intuitively, larger *r* exponentially suppresses background bucket size but also exponentially suppresses the per-band hit rate. For τ = 0.6, *N* ∼ 10^10^, and a median filtered background Jaccard of ∼0.02, the minimum lies at approximately *r* = 6, at which *b* = 10^3^ bands achieves a sensitivity of ∼ 83%. We use these defaults for clustering the full atlas.

##### A.5.4.3 Implementation and Runtime Scaling

The clustering algorithm is implemented in a fully distributed manner in Spark (163). Each step writes a disk-backed partitioned parquet table to enable checkpointing, reduce the repetition of intermediate computation steps, and limit overall memory pressure.

To estimate the runtime scaling, we clustered five random subsamples of UniRef90 from 5M to 100M proteins on a Spark cluster with 64 `i4i.8xlarge` CPU nodes (2,048 cores). As shown in Figure S44, the algorithm scales effectively, clustering up to 100M proteins in ∼1 hour. A power-law fit to these data estimate sub-linear scaling, with *α* = 0.73, which projects to just over 24 hours for the full 6.8B atlas. We find that this extrapolation is an over-optimistic estimate when scaling up a further two orders of magnitude due to bandwidth and memory pressure at scale and the increased iterations required for the greedy clustering stage; the full run took ∼2 days on a dynamically sized cluster with 64-128 `i4i.8xlarge` nodes.

**Figure S44.**
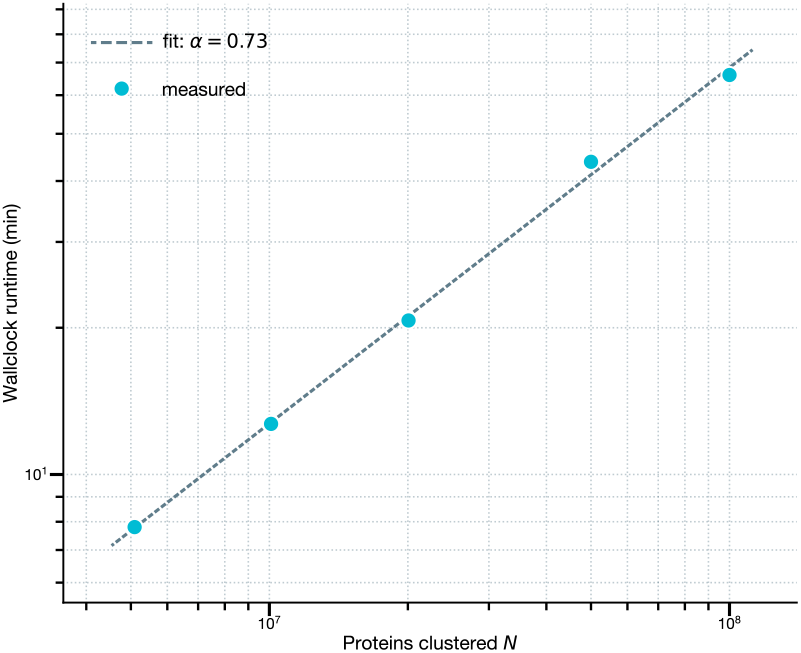
Runtime scaling of SAE clustering. Wall-clock run-time (min) for different data subsets of UniRef90 from 5M to 100M proteins. Power-law fit on this data: 9.836 × 10^−5^ ∗ *N* ^0.730^, with *R*^2^ = 0.99.

##### A.5.4.4 Singleton Rescue

The combination of the single-representative selection procedure and the imperfect recall of LSH candidate generation keeps the algorithm tractable at scale, but also produces a large number of singletons which could be assigned to existing cluster centers. We therefore run a single-pass rescue step that attempts to assign each singleton to an existing cluster.

Let 𝒮 denote the set of singletons from Stage 3 of the clustering pipeline and ℛ the set of representatives of clusters with ≥2 members. Using only the first 150 bands from the band-hash table computed in Stage 1, we calculate all candidate pairs between 𝒮 and ℛ. Because we no longer select only the minimum hash per bucket, every pair is preserved: at *r* = 6, *J* = 0.6, *b*_rescue_ = 150 this achieves almost perfect per-singleton recall relative to the LSH limit. Candidate pairs are verified against the checkpointed 256-dimensional MinHash sketches. For each singleton, we retain the edge to the representative with the highest sketch similarity and update the cluster mapping table. This procedure does not merge clusters or create new representatives, so cluster representatives are preserved.

This procedure rescued 245, 561, 415 singletons (∼ 10% recovery rate). The number of large clusters (≥ 50 members) increased from 7.5M to 7.7M, representing the addition of 28.5M proteins.

##### A.5.4.5 Final clustering statistics

After singleton rescue, the 6, 824, 676, 938 proteins in the ESM Atlas clustered into 3, 049, 430, 104 unique clusters, with 2, 339, 584, 725 of these remaining unassigned singletons (34%). The cluster sizes generally follow a scale-free distribution Figure S45, with a maximum cluster size of 90, 019 and a much larger number of small clusters than large ones. Clusters with more members are more likely to represent conserved proteins (as opposed to low-confidence sequences or gene-calling artifacts). There are 230.8M unique clusters with at least five members, and 7.7M with at least fifty. These large clusters, totaling over 817M proteins, form the basis for much of our exploratory analysis in this paper.

**Figure S45.**
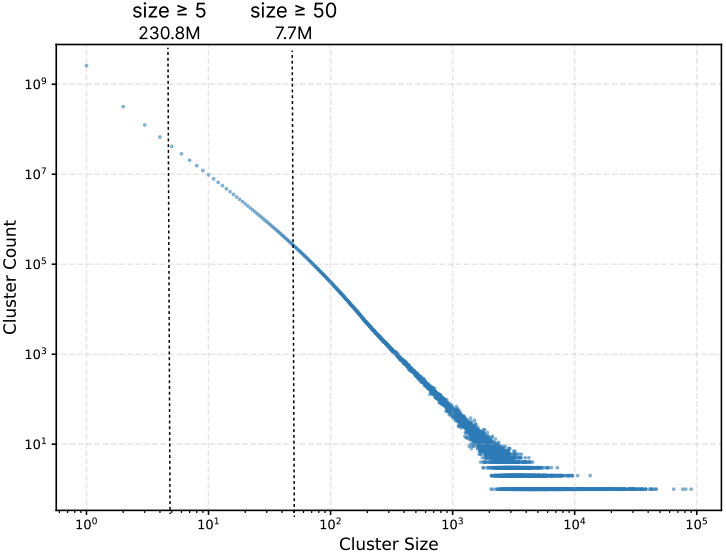
Size distribution of SAE-based clusters. Log-log plot showing the number of clusters at each cluster size. Cluster sizes exhibit a scale-free distribution, with many small clusters and relatively few large ones.

#### A.5.5 Organization of Nucleases in the Atlas

To generate the UMAP over RNase H (Pfam CL0219) members, we defined the similarity between every pair of clusters as the Jaccard similarity over max-pooled SAE features of their representatives. The 2D UMAP was computed on only cluster representatives containing CL0219 with parameters `n_neighbors=15, min_dist=0.1`. Each cluster was assigned a coarse family label by mapping its dominant Pfam domain through a hand-curated mapping covering canonical RNase H subfamilies (Table S15).

**Table S15.** Class-to-Pfam mapping used for RNase H cluster coloring. A cluster is assigned to the plurality Pfam class if ≥ 30% of cluster members hit members of the class.

| Class | Pfam IDs |
| --- | --- |
| Cas9 | PF22702, PF13395, PF16593, PF16592, PF16595, PF18470, PF21069, PF18525, PF18061, PF17893, PF17894, PF18070 |
| Cas12 | PF22126, PF07282, PF18516, PF18501, PF18510, PF22222, PF21918 |
| TnpB | PF01385, PF14279, PF14239 |
| Bridge recombinase | PF01548, PF02371 |
| Mu-like transposase | PF02914 |
| Tn3 transposase | PF01526 |
| RAG1 | PF26100 |
| Argonaute / Piwi | PF02171, PF22474 |
| Spliceosome | PF12134 |
| Maelstrom (piRNA) | PF13017 |
| KDZ + Plavaka | PF18758, PF18759 |
| NurA | PF09376 |
| Med13_C | PF06333 |

For the Cas12 deep-dive, we chose canonical exemplars from the literature: Cas12a from *Acidaminococcus sp*. (AsCas12a), CRISPR-associated transposon from *Scytonema hofmanni* (ShCAST), and the first uncharacterized Cas12f1 (Un1Cas12f1). Features were selected based on fold-change enrichment between activation in each group and manually inspected for biological significance.

The phylogenetic tree for the *lipomyces* Fanzor cluster was constructed by computing a high-quality MSA with MAFFT (164) and performing maximum likelihood tree reconstruction with IQ-TREE 2 (165) with an evolutionary model selected automatically based on the Bayesian information criterion (ModelFinder Plus) with branch-support calculated with 1,000 ultrafast bootstrap replicates.

We performed a search for uncharacterized Cas12/TnpB proteins in the 1.5 million clusters in the Atlas with no annotated Pfam families by computing the pairwise Jaccard similarities between the unannotated representatives against the 1,927 cluster representatives in the Cas12/TnpB section of the RNase H UMAP. This search yielded 315 “dark” clusters with ≥ 0.6 similarity to any Cas12/TnpB. We also computed the nearest five fully unannotated clusters to *S. punctatus* Fanzor (UniProt ID A0A0L0H5U9). All sequences from these clusters were compared to a BLAST database containing every Fanzor protein previously listed in the supplementary information files of Saito *et al*. and Jiang *et al*..

#### A.5.6. Biological Themes in the Atlas

Figure 5E shows the organization of the Atlas according to several biological themes, including those defined by sequence, structure, taxonomy, database annotation, and SAE feature information.

##### A.5.6.1 Pfam Family Distribution

Considering Pfam family annotations as a proxy for function, we can also map the distribution of different functional themes across the atlas. We downloaded the full list of 812 Pfam clans (curated groups of Pfam families defined by shared structural or functional characteristics) as of April 2026, and mapped them to Atlas clusters. A cluster is considered associated with a clan if its dominant Pfam family (family found in the highest number of unique members) is a member of the clan. We sample 70 random clans and visualize them on the 2D UMAP of large clusters in Figure S46. Clans with different functions generally localize to different regions of the UMAP projection, illustrating how the ESMC latent space organizes the protein universe coherently according to shared structural and functional characteristics. The RNase H clan (CL0219) is also visualized separately and shown in Figure 5D.

Clusters with no Pfam annotations at all are combined with those classified as *uncharacterized* by the method described in A.4.5.2 in order to identify clusters with no characterized family annotation for display in Figure 5E.

##### A.5.6.2 Sequence and Structural Diversity Within SAE Clusters

To assess the sequence and structural diversity of the atlas, for the clusters with *>*50 members (7.7M clusters), the within-cluster sequence identity was computed using MM-Seqs2 (136). If a cluster contains more than 1K sequences, it is randomly subsampled to 1K members to compute average pairwise identity. In total, 80,646 clusters were identified which have, on average, *<*30% sequence intra-cluster pairwise sequence identity (`pident` from MMSeqs). This represents around 1% of the atlas. These clusters are shown in the third plot of Figure 5E.

These sequence-dissimilar clusters were then further investigated for coherence with respect to shared predicted structures.

Intra-cluster TM scores were computed using Foldseek (93) easy-search (options `-s 9.5 -e inf --alignment-type 2 --tm-score-threshold 0.0`) on the folded proteins in the Atlas (derived from 70% seq-id clustered versions of the atlas, clustering detailed in A.5.2). Clusters with intra-cluster TM scores all 0.5 were considered to have a single consensus fold. At this threshold, 87.5% of the sequence-diverse clusters (∼70K clusters) were found to have a single consensus fold. The remaining 12.5% (∼10K clusters) have either multiple folds or no consensus folds. This category includes, for example, disordered phage portal families that have no known Pfam annotations.

**Figure S46.**
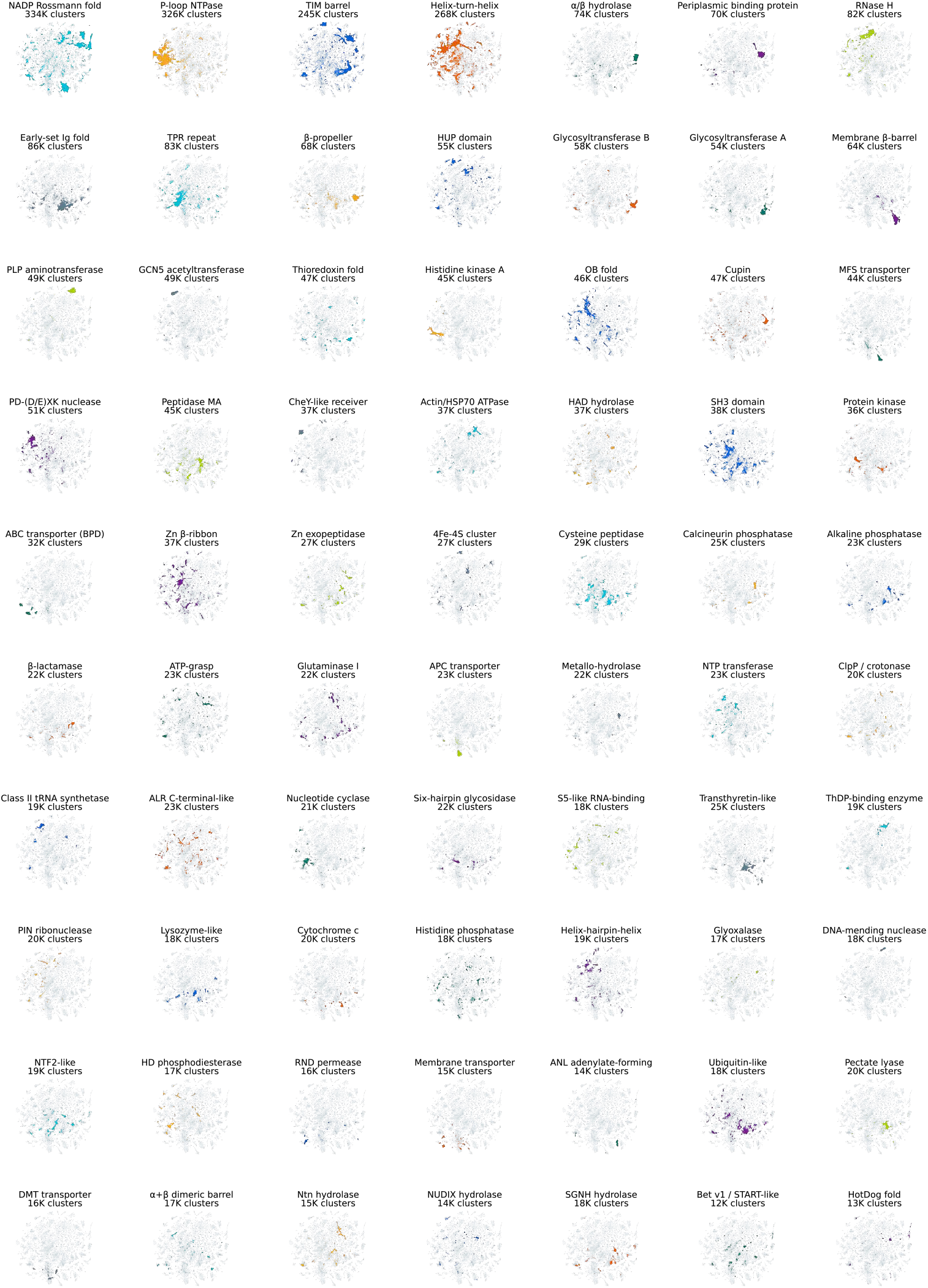
Distribution of 70 Pfam clans in the latent space. UMAP of the 7.7M large clusters in the ESM Atlas, with clusters associated with 70 sampled Pfam clans highlighted. A cluster is highlighted if the most-represented Pfam family in the cluster belongs to the clan.

##### A.5.6.3 Biome and Taxonomy Analysis

###### Biome mappings

To compute biome membership for metagenomic samples from SPIRE and MGnify, we downloaded the biome metadata as of April 2026 (mgy_biomes.tsv.gz, spire_v1_microntology.tsv.gz). For the largest biome categories such as human gut, marine, and soil, we created a unified mapping from each database’s individual categorization scheme to a universal set of categories, shown in Table S16. These categories are not comprehensive but have high enough coverage to calculate trends across the dataset. Clusters for which halophilic biomes are the most common biome in the cluster were selected for display in Figure 5E.

###### Taxonomy mapping

For taxonomy mapping in the remaining databases (IMG/M, IMG/VR, UMAG prok., UMAG euk., Uniparc), we use the included metadata tables from the source databases and convert each taxonomic lineage into NCBI taxonomy IDs using ETE3 (166), resolving to the most specific taxon in the lineage that maps to a TaxID. Lowest common ancestor was then computed with reference to the NCBI taxonomy tree using ETE3. For taxonomic level aggregation in Figure 5F, we combine viruses and cellular organisms at the `root` and `domain` levels: {`cellular root`, `acellular root`} → `root`; {`domain`, `realm`} → `domain`.

##### A.5.6.4 Biome-specific Feature Enrichment

We compute feature enrichment in each of the biome categories defined in Table S16 relative to a background distribution comprising the remainder of the Atlas dataset. First, we build a count matrix of shape |ℬ| × |ℱ|, where each element *b* ∈ ℬ, *f* ∈ ℱ is the number of proteins assigned to biome *b* whose cluster representative has nonzero activation for feature *f* anywhere in its sequence, and compute the marginal total counts in each biome *N*_*b*_ and feature *N*_*f*_. For this analysis, we consider all clusters with at least five members (∼ 230M clusters). To compute enrichment, we build a 2 × 2 contingency table for each combination of biome *b* and feature *f*. Given *A* = count[*b, f*], *B* = *N*_*b*_ − *A, C* = *N*_*f*_ − *A, D* = *N*_*total*_ − *N*_*b*_ *N*_*f*_ + *A*, then the odds ratio is 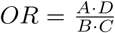. To compute significance, a conventional volcano plot would show the negative log of the corrected p-value (− *log*(*q*)), but for a hypergeometric test with *N*_*total*_ ∼ 10^9^ this blows up rapidly. Instead, we use a standardized z-score on the observed count distribution (assuming that *z* is approximately normal for large *N*):

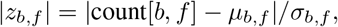

where we assume count[*b, f*] ∼ Hypergeom(*M, K*_*f*_, *N*_*b*_) and *µ*_*b,f*_ = *N*_*b*_*N*_*f*_ */N* and 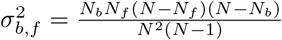 are the mean and variance of the hypergeometric distribution.

In the volcano plots, we filter on a Bonferroni-corrected p-value of 0.01 and then highlight the top five features by combined score |*z*| · log_2_(*OR*). Volcano plots for all biomes are shown in Figure S47.

Some patterns in enrichment are shared across multiple biomes; for example, there is strong enrichment for fungal interaction and regulation signatures in skin, food, and the built environment, and human gut and respiratory microbiomes share similar enrichment for gram-positive bacterial cell envelope signatures. Animal gut samples seem to combine human gut-like bacterial features with signal from mobile genetic elements and bacterial defense systems, suggesting more complex phage-bacteria interactions in these samples than in human gut. This may be driven by the fact that a large fraction of animal gut samples come from livestock, whose gut microbiomes are heavily influenced by antibiotic use and therefore are a major reservoir for novel antimicrobial resistance genes. There are many features related to retroelements, particularly in host-associated metagenomes, which could be interesting sources of new gene editing or delivery proteins. There is also evidence that host-associated proteins drive some of the signal for certain biomes—in other human-associated biomes we see features associated with keratin, suggesting proteins from hair, nails, or skin are being sequenced alongside microbial samples, and in skin we see enrichment in features found in proteins involved in melanin biosynthesis, such as type-3 copper tyrosinases. Extremophilic biomes (other than halophiles, which are highly enriched for acidic tracts), tend to show enrichment for generic C-terminal chain termination motifs, terminal IDRs, and initiator methionines. This likely reflects less a positive enrichment for these features and more a negative enrichment for structured domain-related features; extremophilic samples are disproportionately short hypothetical proteins and ORFs with few known annotations.

##### A.5.6.5 Biological Process Annotation

The SAE feature space provides a mechanism for mapping aspects of biological function without requiring sequences to map directly to existing annotation systems. Proteins can be assigned to function at various levels of specificity, from their precise molecular mechanism at the lowest level to the broad biological processes they are involved in, such as metabolic processes, transcription and translation, or energy production. The Gene Ontology (GO) (87) provides a standardized set of high-level biological processes that are conserved across organisms through their `go-slim` collection, which collects many specific process and pathway annotations into more general groups. These processes are general enough (i.e. contain enough proteins with shared functional characteristics) that it is possible to identify individual features which are highly specific to certain processes above a certain activation. By mapping GO biological processes to features and then using the features to search in the Atlas, it may be possible to identify potential functional connections in proteins that are distant in sequence space.

**Table S16.** Mapping from MGnify and SPIRE biome tags to unified categories. Matching is case-insensitive and in the case of multiple matches, the higher in the list takes priority. ^∗^Extremophile categories are computed only for feature enrichment (Figure 5G); for overall biome counts in Figure 5A these are skipped and assigned to marine/soil/etc. ^†^Collapsed to “Other minor biomes” in Figure 5A.

| Display category | MGNify tags | SPIRE tags |
| --- | --- | --- |
| Human gut | host-associated:human AND (digestive, fecal, intestine, rumen, stomach) | human host AND (intestine, digestive tract, rumen) |
| Animal gut | (digestive, fecal, intestine, rumen, stomach) without human | (intestine, digestive tract, rumen) without human host |
| Oral | host-associated:human AND (mouth, oral, saliva) | saliva, :mouth |
| Skin | host-associated:human AND skin | :skin |
| Respiratory | host-associated:human AND (respiratory, lung, airways) | airways, lung, respiratory |
| Human (other) | host-associated:human (fallback) | human host (fallback) |
| Thermophilic* | thermal spring, hydrothermal, hot spring | hot spring, hydrothermal, very high temperature |
| Psychrophilic* | ice AND aquatic | aquatic:ice, very low temperature |
| Halophilic* | hypersaline, saline, brine | briny, very high salinity |
| Acidic pH* | — | ph:acidic |
| Alkaline pH* | — | ph:alkaline |
| Plant-associated | host-associated:plants, phyllosphere, rhizosphere, plant litter | rhizosphere, phyllosphere, plant litter, plant host |
| Marine | aquatic:marine, ocean, pelagic | marine, ocean, pelagic |
| Freshwater | aquatic:freshwater, lacustrine, freshwater, lake, river, stream | fresh water, lacustrine, stream |
| Soil | terrestrial:soil, :soil, permafrost | soil, permafrost, subsurface |
| Other terrestrial | terrestrial (fallback) | wetland, forest, grassland, tundra, arid, OR terrestrial (fallback) |
| Wastewater / sludge | wastewater, sewage, sludge | wastewater, sewage, sludge |
| Built environment | built environment, hospital | built environment, hospital |
| Food | food production, fermented, dairy | fermented food, dairy |
| Other host-associated (animal) | — | host-associated:animal (collapsed to “Other host-associated” on display) |
| Other host-associated | host-associated | host-associated (fallback) |
| Other aquatic<sup>†</sup> | aquatic (fallback) | aquatic (fallback) |
| Other anthropogenic<sup>†</sup> | engineered | anthropogenic |
| Air | air | air |
| Unknown | no match | no match |

**Figure S47.**
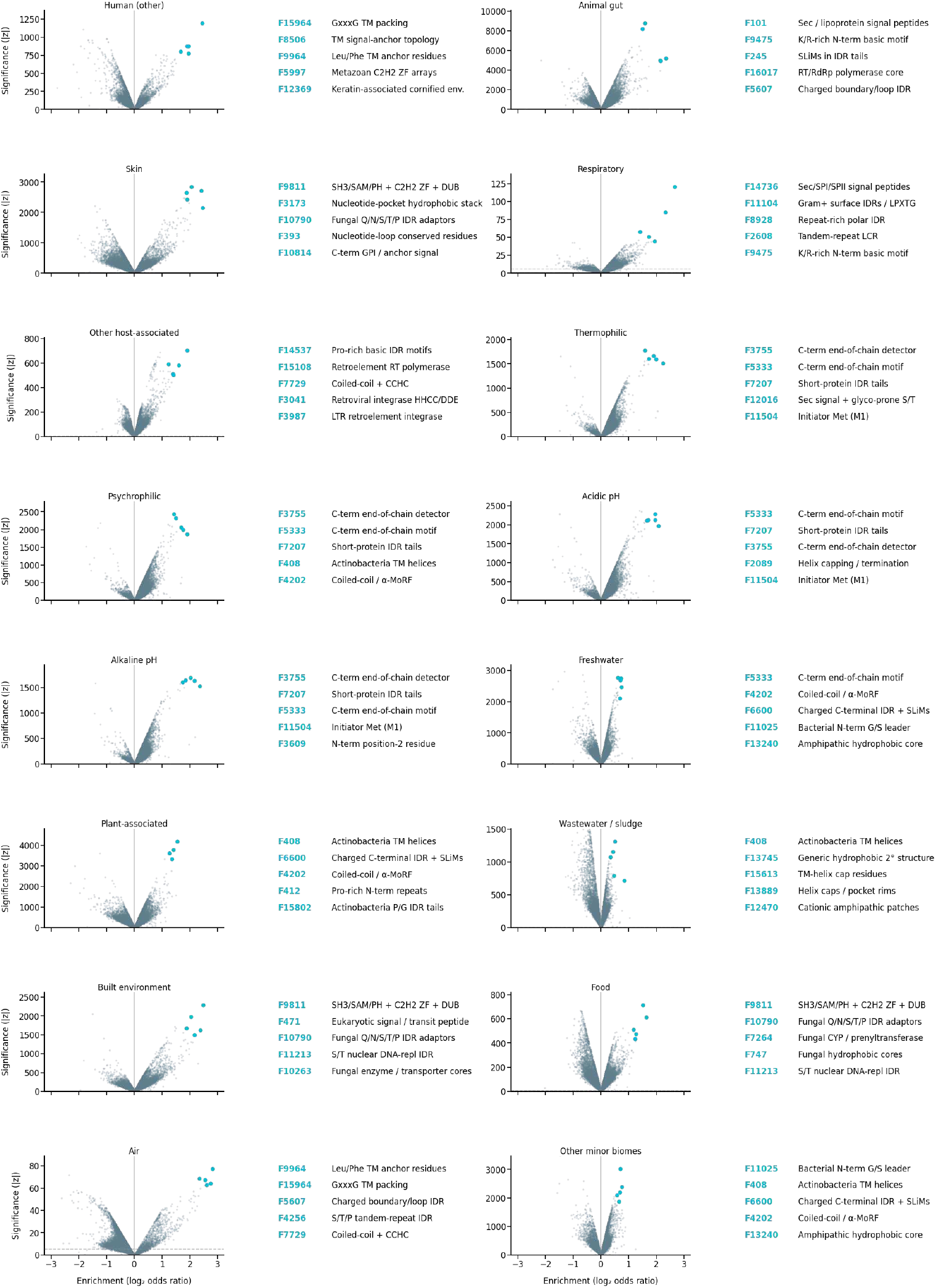
Biome-specific feature enrichment. Biome-specific feature enrichment for the major biome categories in Table S16, not including those highlighted in Figure 5. Effect size (log_2_ *OR*)) and significance |*z*| are computed against a full-atlas background using hypergeometric test and plotted for each biome category. Top five features by combined score are highlighted with a short summary description.

High-precision features for GO biological process terms are identified using the GO annotations downloaded from Swissprot for the 90%-clustered nonredundant Swissprot subset used throughout this work. The dataset is split in half, with 50% of proteins used to identify features and select activation thresholds and 50% used for validation. For each pair of feature and GO term, a threshold is selected from [0.15, 0.3, 0.45, 0.6, 0.75, 0.9, 0.93, 0.96, 0.99] on max-normalized activation to maximize the F1 score on the train proteins. Pairs are then evaluated on the held-out proteins and precision, recall, and F1 score are computed. Sets of feature-process pairs with precision of greater than or equal to 0.5, 0.7, and 0.9 (filtered to have at least 10 true positives) are stored for use in identifying candidate proteins for each biological process. This procedure optimizes for precision, limiting false positives, and thus finds features associated with only a subset of the proteins involved in a process; an approach focused on recall would identify more candidate proteins but would also surface many false positives due to structural homology, shared ligand/cofactor binding, or other spurious similarities. This procedure produces 5, 955, 2, 258, and 491 feature-process pairs at 50%, 70%, and 90% precision, respectively (representing 61, 57, and 49 unique processes). The number of high-precision features per process ranges from 1–37 (median 6) at 90% precision and 1–578 (median 35) at 50% precision.

The feature sets produced by this procedure are used to search against cluster representatives of the 7.7M large clusters in the Atlas, recording the clusters which activate the feature above the threshold identified in the tuning stage. The number of proteins in these clusters are summed up to estimate how many proteins could potentially be related to each process at different levels of precision. To highlight how features can be used to connect known proteins to previously uncharacterized proteins, we identify clusters that activate features with ≥90% precision but have no existing Pfam annotations, a proxy for them being functionally uncharacterized. Depending on the family, up to 70% of clusters are uncharacterized. The processes with the highest fraction of uncharacterized clusters are `mitochondrion organization` (69.5%), `photosynthesis` (54.5%), `mitochondrial gene expression` (33.5%), `snRNA metabolic process` (24.3%), `cell motility` (20.5%).

These uncharacterized clusters are ranked by the SAE vector dot product similarity to clusters with Pfam annotations that were assigned to the same biological process. The top hits in each were used for a structural search against the PDB using Foldseek. Figure 5H shows three examples of the results of this procedure, with the high-precision features used to connect the two proteins highlighted on the structures. In general, these features are highly interpretable, representing concepts that are closely related to the process they are associated with. This provides a mechanistic explanation for the high SAE similarity in terms that are grounded in both the ESMC latent space and also in known biological annotations, increasing confidence in potential functional hypotheses.

#### A.6. Safety and Model Release

Biohub has established a safety team which assesses the benefits and potential risks of our models and tools prior to release, and develops mitigations where necessary. We follow a structured approach that includes assessing both biosafety and biosecurity risks, as well as existing comparable open-source models and tools. We actively engage with the scientific community, stakeholders, and domain experts to advance innovation as well as best practices for responsible development. Risk assessment was conducted for each of the components of this release, including our ESMC models, ESMFold2, ESMC SAEs, ESM Atlas, and binder design system.

Informed by our risk assessments, we are releasing the source code and model weights for ESMC 6 billion, ESMFold2, and ESMC SAEs. (ESMC 600 million and 300 million parameter model weights have already been open to the scientific community.) We are also releasing our ESM Atlas dataset and binder design system openly. Biohub values open science, and we share our research with the scientific community so that others can evaluate, reproduce, and build upon our work.

##### A.6.1. Evaluations

Prior to release, we conducted evaluations to inform our understanding of capability uplift for specific misuse-relevant functional tasks. We measure the ability of the ESMC models to understand the effects of mutations in viral proteins. The evaluation consists of 25 Deep Mutational Scanning (DMS) datasets collected from ProteinGym, all of which are from viral proteins (142). We evaluate the correlation (Spearman *ρ*) between the predicted variant effect and the measured variant effect. We compare the performance of the ESMC models to existing open source protein language models ESM2 650M (3) and ESM3 1.4B Open (4). We also compare to a BLOSUM62 baseline, which is based on amino acid substitution frequencies, and to EVMutation, which is a Markov Random Field model trained on a multiple sequence alignment of the target protein (167). The results show ESMC models perform worse than or similar to existing tools for variant effect prediction in viral proteins (Figure S48).

**Figure S48.**
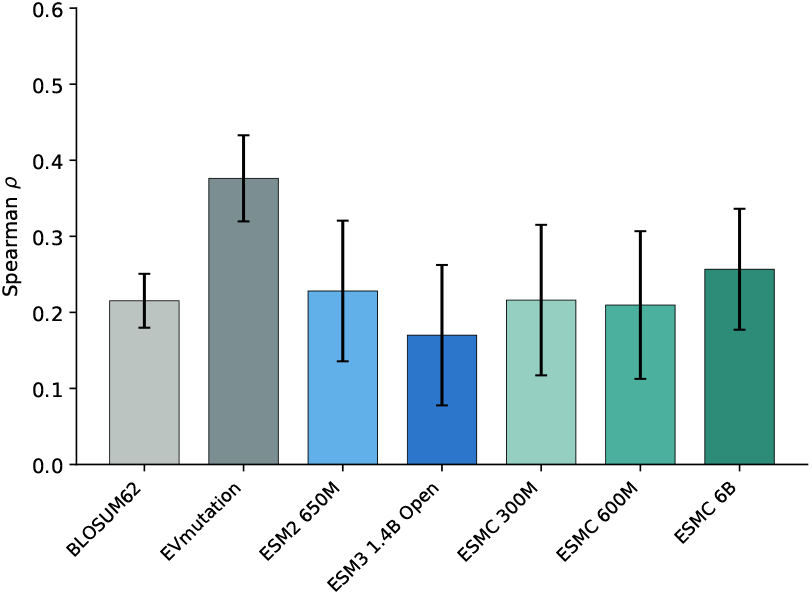
Fitness prediction of open source and new models. Correlation (Spearman *ρ*) across 25 deep mutational scanning datasets from viral taxa, collated in ProteinGym. Error bars indicate a 95% confidence interval of the mean estimate.

